# Sex-Dependent Shared and Non-Shared Genetic Architecture Across Mood and Psychotic Disorders

**DOI:** 10.1101/2020.08.13.249813

**Authors:** Gabriëlla A. M. Blokland, Jakob Grove, Chia-Yen Chen, Chris Cotsapas, Stuart Tobet, Robert Handa, Schizophrenia Working Group of the Psychiatric Genomics Consortium, David St Clair, Todd Lencz, Bryan J. Mowry, Sathish Periyasamy, Murray J. Cairns, Paul A. Tooney, Jing Qin Wu, Brian Kelly, George Kirov, Patrick F. Sullivan, Aiden Corvin, Brien P. Riley, Tõnu Esko, Lili Milani, Erik G. Jönsson, Aarno Palotie, Hannelore Ehrenreich, Martin Begemann, Agnes Steixner-Kumar, Pak C. Sham, Nakao Iwata, Daniel R. Weinberger, Pablo V. Gejman, Alan R. Sanders, Joseph D. Buxbaum, Dan Rujescu, Ina Giegling, Bettina Konte, Annette M. Hartmann, Elvira Bramon, Robin M. Murray, Michele T. Pato, Jimmy Lee, Ingrid Melle, Espen Molden, Roel A. Ophoff, Andrew McQuillin, Nicholas J. Bass, Rolf Adolfsson, Anil K. Malhotra, Bipolar Disorder Working Group of the Psychiatric Genomics Consortium, Nicholas G. Martin, Janice M. Fullerton, Philip B. Mitchell, Peter R. Schofield, Andreas J. Forstner, Franziska Degenhardt, Sabrina Schaupp, Ashley L. Comes, Manolis Kogevinas, José Guzman-Parra, Andreas Reif, Fabian Streit, Lea Sirignano, Sven Cichon, Maria Grigoroiu-Serbanescu, Joanna Hauser, Jolanta Lissowska, Fermin Mayoral, Bertram Müller-Myhsok, Beata Świątkowska, Thomas G. Schulze, Markus M. Nöthen, Marcella Rietschel, John Kelsoe, Marion Leboyer, Stéphane Jamain, Bruno Etain, Frank Bellivier, John B. Vincent, Martin Alda, Claire O’Donovan, Pablo Cervantes, Joanna M. Biernacka, Mark Frye, Susan L. McElroy, Laura J. Scott, Eli A. Stahl, Mikael Landén, Marian L. Hamshere, Olav B. Smeland, Srdjan Djurovic, Arne E. Vaaler, Ole A. Andreassen, Major Depressive Disorder Working Group of the Psychiatric Genomics Consortium, Bernhard T. Baune, Tracy Air, Martin Preisig, Rudolf Uher, Douglas F. Levinson, Myrna M. Weissman, James B. Potash, Jianxin Shi, James A. Knowles, Roy H. Perlis, Susanne Lucae, Dorret I. Boomsma, Brenda W. J. H. Penninx, Jouke-Jan Hottenga, Eco J. C. de Geus, Gonneke Willemsen, Yuri Milaneschi, Henning Tiemeier, Hans J. Grabe, Alexander Teumer, Sandra Van der Auwera, Uwe Völker, Steven P. Hamilton, Patrik K. E. Magnusson, Alexander Viktorin, Divya Mehta, Niamh Mullins, Mark J. Adams, Gerome Breen, Andrew M. McIntosh, Cathryn M. Lewis, Sex Differences Cross-Disorder Analysis Group of the Psychiatric Genomics Consortium, The Lundbeck Foundation Initiative for Integrative Psychiatric Research (iPSYCH), David M. Hougaard, Merete Nordentoft, Ole Mors, Preben B. Mortensen, Thomas Werge, Thomas D. Als, Anders D. Børglum, Tracey L. Petryshen, Jordan W. Smoller, Jill M. Goldstein

## Abstract

**BACKGROUND:** Sex differences in incidence and/or presentation of schizophrenia (SCZ), major depressive disorder (MDD), and bipolar disorder (BIP) are pervasive. Previous evidence for shared genetic risk and sex differences in brain abnormalities across disorders suggest possible shared sex-dependent genetic risk.

**METHODS:** We conducted the largest to date genome-wide genotype–by–sex (GxS) interaction of risk for these disorders, using 85,735 cases (33,403 SCZ, 19,924 BIP, 32,408 MDD) and 109,946 controls from the Psychiatric Genomics Consortium (PGC) and iPSYCH.

**RESULTS:** Across disorders, genome-wide significant SNP-by-sex interaction was detected for a locus encompassing *NKAIN2* (rs117780815; *p*=3.2×10^−8^), that interacts with sodium/potassium-transporting ATPase enzymes implicating neuronal excitability. Three additional loci showed evidence (*p*<1×10^−6^) for cross-disorder GxS interaction (rs7302529, *p*=1.6×10^−7^; rs73033497, *p*=8.8×10^−7^; rs7914279, *p*=6.4×10^−7^) implicating various functions. Gene-based analyses identified GxS interaction across disorders (*p*=8.97×10^−7^) with transcriptional inhibitor *SLTM*. Most significant in SCZ was a *MOCOS* gene locus (rs11665282; *p*=1.5×10^−7^), implicating vascular endothelial cells. Secondary analysis of the PGC-SCZ dataset detected an interaction (rs13265509; *p*=1.1×10^−7^) in a locus containing *IDO2*, a kynurenine pathway enzyme with immunoregulatory functions implicated in SCZ, BIP, and MDD. Pathway enrichment analysis detected significant GxS of genes regulating vascular endothelial growth factor (VEGF) receptor signaling in MDD (*p*_FDR_<0.05).

**CONCLUSIONS:** In the largest genome-wide GxS analysis of mood and psychotic disorders to date, there was substantial genetic overlap between the sexes. However, significant sex-dependent effects were enriched for genes related to neuronal development, immune and vascular functions across and within SCZ, BIP, and MDD at the variant, gene, and pathway enrichment levels.

## Introduction

Sex differences are pervasive in psychiatric disorders, including major depressive disorder (MDD), schizophrenia (SCZ), and bipolar disorder (BIP). There is a significantly higher risk for MDD in women (1) and SCZ in men (2). BIP prevalence is approximately similar, but age at onset, course, and prognosis vary considerably by sex (3, 4), as they do in SCZ and MDD (5–7). Additionally, certain brain regions share structural and functional abnormalities and dysregulated physiology across disorders that are sex-dependent (8, 9).

The majority of twin studies have not detected sex differences in heritability of these disorders (10), or differences in twin intra-pair correlations between same-sex and opposite-sex dizygotic pairs (11, 12). However, specific disease risk variants may not be the same in both sexes (i.e., “sex-specific” effects) or variants may have different effect sizes in each sex (i.e., “sex-dependent” effects). Sex-dependent modification of allelic effects on the autosomes and X chromosome may contribute to sex differences in disease prevalence, similar to other complex human traits (e.g., blood pressure, waist-hip ratio) (13, 14). Aside from sex-specific variants, incidence differences may result from a female or male protective effect, whereby one sex may require a higher burden of genetic liability to cross the threshold to disease manifestation. This suggests quantitative risk differences (i.e., “sex-dependence”), a notion supported by an early observation that female SCZ cases were more likely to come from multiplex families (15).

Regarding SCZ, there is a long history of examining sex differences in familial/genetic transmission (16), given differences in incidence, brain abnormalities, and course (17, 18). Recently, large genetic cohorts of SCZ and autoimmune disorders identified greater effects of complement component 4 (C4) alleles in SCZ men than women (19, 20). Compared with SCZ, sex differences in incidence of MDD are greater, with a 2:1 female predominance, and there is some evidence for stronger sex differences in recurrent MDD (rMDD) compared with single-episode MDD, although inconsistent (7, 21–23). With increased interest in examining the genetics of sex differences in psychiatric disorders and related phenotypes (24–32), transcriptomics studies are beginning to provide insights into mechanisms underlying sex differences in risk. Notably, >10% of autosomal genes exhibit sexually dimorphic gene expression in the brain, predominantly genes related to synaptic transmission, dopamine receptor signaling, and immune response, suggesting potential mechanisms mediating sex differences in psychiatric disorders.

In order to test for sex differences in genetic risk, it is essential to have adequate power to test for interaction effects (33). Given sample size limitations, genome-wide association studies (GWAS) of psychiatric disorders have typically not examined genotype-by-sex (GxS) interactions. Here, we capitalized on a unique opportunity to utilize cohorts from the PGC and iPSYCH consortia (n = 195,681) to assess interactions between sex and genetic risk of MDD, SCZ and BIP within and across disorders.

## Methods and Materials

### Participants

The Psychiatric Genomics Consortium (PGC) (34–36) included 43 SCZ (30,608 patients, 38,441 controls), 28 BIP (18,958 patients, 29,996 controls), and 26 MDD cohorts (15,970 patients, 24,984 controls; **Supplementary Table 1**). The iPSYCH cohort in Denmark (37) included 2,795 SCZ patients and 2,436 controls, 966 BIP and 551 controls, and 16,438 MDD and 13,538 controls (**Supplementary Table 2**). Primary analyses used the PGC and iPSYCH datasets. Secondary PGC-only analyses (see **Supplementary Materials**) were performed to facilitate comparison to other PGC studies and ensure that different diagnostic criteria in PGC and iPSYCH (DSM-IV and ICD-10, respectively) were not impacting results. All cohorts were European ancestry, except three East Asian SCZ cohorts.

### Quality Controls and Analytics

Quality control (QC) and imputation to the 1000 Genomes Phase 3 reference panel were performed using PGC’s Rapid Imputation for COnsortias PIpeLIne (RICOPILI) (38) and previously described filtering thresholds (34–36). An overview of subsequent QC and analytic steps is provided in **Supplementary Figure 1**. Identity-by-descent (IBD) filtering is described in **Supplementary Methods**. At MAF=0.05, this study had 83%-99% (within-disorder) and 88% (cross-disorder) power to detect interaction effects at an odds ratio of >= 1.2, and >= 1.1, respectively (**Supplementary Table 3; Supplementary Figure 2**).

Sex-stratified GWAS summary statistics were obtained by logistic regression of men and women separately within each cohort using PLINK (39), followed by standard-error weighted meta-analysis across cohorts using METAL (40). Summary statistics were entered into Linkage Disequilibrium (LD) Score Regression (LDSC) (41, 42) to estimate autosomal sex-specific SNP-based heritability 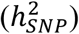 for each disorder (**Figure 1)** and bivariate genetic correlations (*r*_*g*_) within and across disorders.

**Figure 1.**
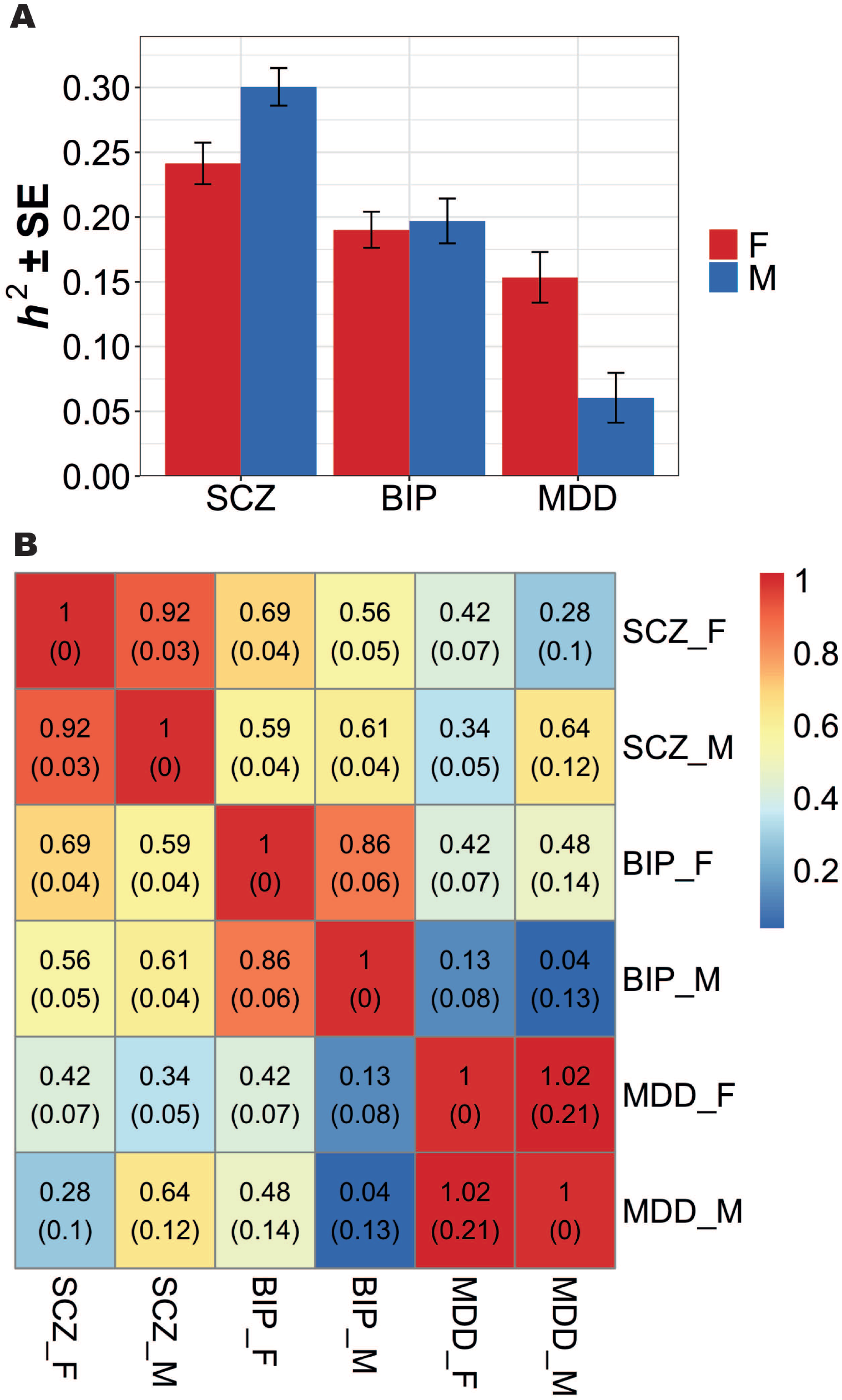
LD Score Regression estimates of sex-specific SNP-based **(a)** heritability, *h*^*2*^ (±SE), and **(b)** genetic correlations, *r*_*g*_ (SE).This graph shows *h*^*2*^ and *r*_*g*_ estimates for MAF > 0.01. a) Heritability estimates were substantially different between the sexes for SCZ (*p*_FDR_ = 0.019) and MDD (p_FDR_ = 0.005), but not BIP (*p*_FDR_ = 0.381). b) SNP-based genetic correlations (*r*_*g*_) between males and females within each disorder ranged between 0.86 and 1 and were significantly different from 1 for SCZ (*p*_FDR_ = 0.039) and BIP (*p*_FDR_ = 0.039), but not MDD (*p*_FDR_ = 0.397). No significant differences in the cross-disorder genetic correlations between males and females, with the exception of *r*_*g*_ between BIP and MDD (*r*_*gF*_ = 0.42; *r*_*gM*_ = 0.04; *p*_FDR_ = 0.044). Abbreviations: BIP = bipolar disorder; MDD = major depressive disorder; SCZ = schizophrenia; F = females; M = males; LD = linkage disequilibrium; SE = standard error.

PLINK (39) was used to perform a genome-wide GxS interaction analysis in each study cohort, followed by standard-error weighted meta-analysis of GxS interactions using METAL (40). GxS interaction analyses were performed using linear regression with a main effect for each SNP, main effect of sex, and SNP-by-sex interaction terms, using an additive model for SNPs (controlling for 10 ancestry principal components [PC]). Secondary regression models included additional controls using 10 SNP-by-PC and 10 sex-by-PC interaction terms in addition to the above (43). Adding too many covariates can destabilize the effect estimates and lead to increased dropout of SNPs due to estimation problems, especially in smaller cohorts, thus the first model is our primary model. Secondary analytic model *p*-values are included in brackets.

GxS interactions with X-linked SNPs were tested using two models. Model A assumed complete and uniform X-inactivation in women and similar effect size between the sexes by assigning 0, 1, or 2 copies of an allele to women and 0 or 2 copies to men. As these assumptions often do not hold, Model B assigned 0 or 1 copy to men.

A three-degrees-of-freedom test omnibus test (44) was performed by summing the χ^2^ values for individual disorder GxS interaction meta-analyses in order to identify SNPs with opposing GxS effects across disorders (see **Supplementary Methods** for details).

LD-independent SNPs (r^2^ < 0.1) with suggestive or genome-wide significant GxS interactions (*p*<1×10^−6^) were used as index SNPs for fine-mapping to obtain credible SNPs (i.e. likely causal) using FINEMAP (45) and CAVIAR (46) (see **Supplementary Methods**). Regions for fine-mapping were defined as all SNPs in LD (r^2^ > 0.6) with the index SNP.

SCZ and cross-disorder analyses of autosomes and X chromosome were conducted with and without inclusion of the East Asian cohorts to evaluate population effects. Findings were not significantly different and therefore all subsequent analyses utilized only European ancestry cohorts (see **Supplementary Methods**).

Gene-based analyses were conducted using MAGMA (47) (significant *p*-value=2.6×10^−6^; see **Supplementary Methods)** Gene set enrichment tests were performed (47) to determine whether (near-)significant SNPs (*p*<1×10^−4^) clustered into particular biological pathways, to explore functional similarity of genes implicated by GxS interactions. Hypothesis-free analyses were performed for 10,353 gene sets from the Molecular Signatures Database (MSigDB). Data-driven enrichment analyses were performed for nine gene sets/ pathways implicated in prior studies: immune/neurotrophic, synaptic, and histone methylation gene sets enriched across PGC SCZ, BIP, and MDD cohorts (48), and central nervous system (CNS) pathways enriched in the CLOZUK+PGC SCZ cohort (49). Pathway enrichment *p*-values were FDR-corrected based on number of pathways tested.

Gene expression and expression quantitative trait locus (eQTL) data from several publicly available resources were evaluated to validate and interpret SNPs with a GxS interaction *p*-value < 1×10^−6^ (see **Supplementary Methods** for details).

Finally, GxS interaction results were compared to previously reported sex-dependent or sex-specific effects on psychiatric risk (*p*<5×10^−8^) (see **Supplementary Methods and Tables**).

## Results

### Sex-stratified LD Score Regression

Based on sex-stratified GWAS results displayed in **Supplementary Figure 3**, men and women 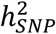 were estimated. Within each disorder, the 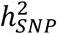 for men and women (**Figure 1a**) was significantly greater than 0 (mean 0.19; all *p* < 0.001) (**Supplementary Table 4**), indicating adequate power to detect broader polygenic signals. Estimates of 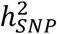 increased minimally across a range of MAF cutoffs (MAF>1%, 2%, 5%), indicating rarer variants contributed little (**Supplementary Table 4**). Heritability estimates were substantially different between the sexes for SCZ (*p*_FDR_ = 0.019; 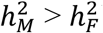) and MDD (*p*_FDR_ = 0.005; 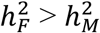), but not BIP (*p*_FDR_ = 0.381) (**Supplementary Table 4**). Although correlations between male and female GWAS *p*-values were low (**Supplementary Figure 4**), SNP-based genetic correlations (*r*_*g*_) between men and women within disorders ranged between 0.86 and 1 and were significantly different from 1 for SCZ (*p*_FDR_ = 0.039) and BIP (*p*_FDR_ = 0.039), but not MDD (*p*_FDR_ = 0.397) (**Figure 1b; Supplementary Table 5a**). Additionally, we observed no significant differences in cross-disorder genetic correlations by sex, except *r*_*g*_ between BIP and MDD (*r*_*gF*_ = 0.42; *r*_*gM*_ = 0.04; *p*_FDR_ = 0.044) (**Figure 1b; Supplementary Table 5b**). However, within-sex, SCZ and BIP women were more highly correlated than SCZ with MDD women, and MDD women correlated similarly to both SCZ and BIP. In contrast, SCZ with BIP and MDD men correlated similarly, but MDD and BIP men were uncorrelated. Findings suggest there may be different within-sex genetic differences that need further understanding and raise the complexity of investigating the genetics of sex effects.

### Genome-wide SNP-by-Sex Interactions

Quantile-quantile plots indicated no systematic inflation of test statistics (**Supplementary Figure 5**). Genomic control lambda (λ_GC_) revealed no significant evidence of population stratification in the meta-analysis of the cross-disorder European ancestry (λ_GC_=0.9828), cross-disorder European + East Asian (λ_GC_=0.9838), SCZ European ancestry (λ_GC_=0.9991), SCZ European + East Asian (λ_GC_=1.002), BIP (λ_GC_=0.9879), or MDD (λ_GC_=0.9833) cohorts.

Analyses within disorders did not detect genome-wide significant interactions for SCZ, BIP, or MDD, however suggestive evidence (*p*<1×10^−6^) was obtained for several loci (**Table 1, Supplementary Table 8**). Overall, there was little overlap between the strongest interactions for each disorder (**Supplementary Figure 6)**. The most significant results were obtained for SCZ for a locus in the 5’ UTR of the *MOCOS* gene (rs11665282: *p*=1.48×10^−7^ [secondary model *p*_*ext*_=2.53×10^−5^]; **Supplementary Figures 6-8)** and an intergenic locus near the non-coding RNA gene *LINC02181* (rs12445424: *p*=3.52×10^−7^ [*p*_*ext*_ =2.28×10^−4^]; **Supplementary Figures 6-8)**. The top GxS interaction locus for BIP was located on chromosome 9 near the *TUSC1* gene (rs12341335: *p*=2.29×10^−7^ [*p*_*ext*_ =7.91×10^−7^]; **Supplementary Figures 6-8)**. Suggestive evidence for GxS effects in MDD risk was detected for chromosome 1 locus in and around *SPAG17* (rs9428240: *p*=1.64×10^−7^ [*p*_*ext*_ = 3.31×10^−7^]), which remained in rMDD (*p*=1.40×10^−7^ [*p*_*ext*_=1.05×10^−7^])), and chromosome 17 locus spanning multiple genes including *ZNF385C* (rs147515485: *p*=4.61×10^−7^ [*p*_*ext*_ = 4.76×10^−6^]; **Supplementary Figures 6-8)**. Post-hoc analysis of rMDD did not reveal additional loci at *p* < 1×10^−6^. Secondary analyses of the PGC SCZ cohort identified a noteworthy locus in an intergenic region between the *IDO2* and *C8orf4* genes (rs13265509: *p*=1.09×10^−7^ [*p*_*ext*_ =1.23×10^−6^]; **Supplementary Table 15a**). Meta-analysis of GxS interactions across cohorts from all 3 disorders (in contrast to omnibus tests) revealed suggestive evidence for three additional intergenic loci (*p*<1×10^−6^) (**Table 1, Supplementary Table 6f-i**).

**Table 1.** Single-disorder and Cross-disorder Autosomal SNP-by-sex interaction results. Cross-disorder and within-disorder meta-analyses were carried out using METAL, incorporating cohort-level summary statistics from PLINK. Listed are SNPs with interaction *p*-values < 1×10^−6^ in SCZ, BIP, (r)MDD, and cross-disorder. Loci were clumped using ‘plink --bfile 1kgp_ref_file --clump metal_output --clump-p1 1e-4 --clump-p2 1e-4 --clump-r2 0.6 --clump-kb 3000’. Extended results (p < 1×10^−4^), including eQTL data for the variants highlighted in this table, and including secondary extended model statistics, are available in **Supplementary Table 6**. Abbreviations: SNP, Variant rs ID; p**GxS**; *p*-value for GxS interaction in combined PGC + iPSYCH datasets (p-value for secondary extended model, p_ext_, in parentheses); CHR, Chromosome; BP, Base Pair Position; A1/A2, Allele 1/Allele 2; Freq1, Frequency of Allele 1; MAF, Minor Allele Frequency; Beta_GxS_, Beta (Standard Error) for GxS interaction; Beta_F_ (SE), Beta (Standard Error) for female-stratified association; p_F_, *p*-value for female-stratified association; Beta_M_, Beta (Standard Error) for male-stratified association; p_M_, *p*-value for male-stratified association; Z_FM_, Z-score heterogeneity females-males; p_FM_, *p*-value heterogeneity females-males

| SNP | CHR | BP | A1/A2 | Freq1<br>MAF | Compartment | Gene<br>(Distance in kb) | N Cases<br>(%Female) | N Controls<br>(%Female) | Beta<sub>GxS</sub><br>(SE) | p<sub>GxS</sub> (p<sub>ext</sub>) | Beta<sub>F</sub><br>(SE) | p<sub>F</sub> | Beta<sub>M</sub><br>(SE) | p<sub>M</sub> | Z<sub>FM</sub> | p<sub>FM</sub> |
| --- | --- | --- | --- | --- | --- | --- | --- | --- | --- | --- | --- | --- | --- | --- | --- | --- |
| **Schizophrenia (European only)** |  |  |  |  |  |  |  |  |  |  |  |  |  |  |  |  |
| rs11665282 | 18 | 33767479 | A/G | 0.69<br>0.31 | UTR5 | MOCOS | 21,581<br>(35.18%) | 24,250<br>(48.62%) | -0.156<br>(0.030) | 1.48E-7<br>(2.53E-5) | -0.081<br>(0.023) | 3.98E-4 | 0.072<br>(0.019) | 2.16E-4 | -5.09 | 3.50E-7 |
| rs12445424 | 16 | 87063374 | A/G | 0.26<br>0.26 | intergenic | LINC02188 (291.9);<br>LINC02181 (280.2) | 29,467<br>(36.04%) | 34,519<br>(48.33%) | 0.140<br>(0.028) | 3.52E-7<br>(2.28E-4) | 0.097<br>(0.021) | 5.80E-6 | -0.050<br>(0.018) | 4.67E-3 | 5.30 | 1.19E-7 |
| **Schizophrenia (European + East Asian)** |  |  |  |  |  |  |  |  |  |  |  |  |  |  |  |  |
| rs11665282 | 18 | 33767479 | A/G | 0.69<br>0.31 | UTR5 | MOCOS | 22,060<br>(35.39%) | 24,674<br>(48.26%) | -0.149<br>(0.03) | 3.74E-7<br>(4.46E-5) | -0.077<br>(0.023) | 6.74E-4 | 0.070<br>(0.019) | 2.53E-4 | -4.96 | 6.89E-7 |
| **Bipolar Disorder** |  |  |  |  |  |  |  |  |  |  |  |  |  |  |  |  |
| rs12341335 | 9 | 25649145 | T/C | 0.90<br>0.10 | intergenic | TUSC1 (27.2) | 7,730<br>(57.72%) | 13,635<br>(51.28%) | 0.373<br>(0.072) | 2.29E-7<br>(7.91E-7) | 0.176<br>(0.048) | 2.59E-4 | -0.201<br>(0.054) | 2.11E-4 | 5.20 | 2.03E-7 |
| rs17651437 | 2 | 106055684 | T/C | 0.52<br>0.48 | upstream | FHL2 | 16,365<br>(60.18%) | 28,140<br>(50.75%) | 0.155<br>(0.031) | 3.72E-7<br>(1.04E-5) | 0.079<br>(0.020) | 9.97E-5 | -0.069<br>(0.023) | 3.08E-3 | 4.79 | 1.63E-6 |
| **Major Depressive Disorder** |  |  |  |  |  |  |  |  |  |  |  |  |  |  |  |  |
| rs9428240 | 1 | 118831676 | T/C | 0.59<br>0.41 | intergenic | SPAG17 (103.8) | 14,232<br>(68.63%) | 21,846<br>(50.63%) | -0.181<br>(0.035) | 1.64E-7<br>(3.31E-7) | -0.087<br>(0.022) | 6.41E-5 | 0.094<br>(0.028) | 8.41E-4 | -5.08 | 3.70E-7 |
| rs147515485 | 17 | 40182099 | T/C | 0.02<br>0.02 | intronic | ZNF385C | 31,149<br>(61.17%) | 35,385<br>(50.89%) | -0.472<br>(0.094) | 4.61E-7<br>(4.76E-6) | -0.190<br>(0.060) | 1.55E-3 | 0.303<br>(0.074) | 4.39E-5 | -5.17 | 2.39E-7 |
| **Recurrent Major Depressive Disorder** |  |  |  |  |  |  |  |  |  |  |  |  |  |  |  |  |
| rs61138090 | 1 | 118832069 | D/I2 | 0.59<br>0.41 | intergenic | SPAG17 (104.2) | 7,685<br>(70.59%) | 15,976<br>(51.71%) | -0.240<br>(0.046) | 1.40E-7<br>(-) | -0.109<br>(0.028) | 1.03E-4 | 0.142<br>(0.038) | 2.08E-4 | -5.28 | 1.30E-7 |
| **Cross-Disorder SCZ-BIP-MDD (European only)** |  |  |  |  |  |  |  |  |  |  |  |  |  |  |  |  |
| rs7302529 | 12 | 77321581 | T/C | 0.26<br>0.26 | intergenic | CSRP2 (48.8);<br>E2F7 (93.4) | 34,638<br>(51.36%) | 34,696<br>(50.15%) | 0.145<br>(0.028) | 1.60E-7<br>(5.35E-7) | 0.087<br>(0.019) | 5.09E-6 | -0.051<br>(0.020) | 1.15E-2 | 4.98 | 6.51E-7 |
| rs73033497 | 7 | 2910659 | A/T | 0.86<br>0.14 | intergenic | GNA12 (26.7);<br>CARD11 (35.0) | 14,916<br>(49.21%) | 17,547<br>(47.81%) | 0.246<br>(0.050) | 8.82E-7<br>(2.24E-6) | 0.116<br>(0.036) | 1.09E-3 | -0.128<br>(0.035) | 2.69E-4 | 4.89 | 1.03E-6 |
| **Cross-Disorder SCZ-BIP-MDD (European + East Asian)** |  |  |  |  |  |  |  |  |  |  |  |  |  |  |  |  |
| rs7914279 | 10 | 122161890 | T/G | 0.89<br>0.11 | intergenic | MIR4682 (44.3);<br>PLPP4 (54.6) | 78,640<br>(49.95%) | 71,790<br>(49.70%) | 0.146<br>(0.029) | 6.39E-7<br>(4.78E-6) | 0.064<br>(0.020) | 1.86E-3 | -0.077<br>(0.021) | 2.27E-4 | 4.82 | 1.43E-6 |
| rs73033497 | 7 | 2910659 | A/T | 0.86<br>0.14 | intergenic | GNA12 (26.7);<br>CARD11 (35.0) | 14,916<br>(49.21%) | 17,547<br>(47.81%) | 0.246<br>(0.050) | 8.82E-7<br>(2.24E-6) | 0.116<br>(0.036) | 1.09E-3 | -0.128<br>(0.035) | 2.69E-4 | 4.89 | 1.03E-6 |
| rs7302529 | 12 | 77321581 | T/C | 0.25<br>0.25 | intergenic | CSRP2 (48.8);<br>E2F7 (93.4) | 35,114<br>(50.69%) | 36,707<br>(50.72%) | 0.133<br>(0.027) | 9.37E-7<br>(2.69E-6) | 0.082<br>(0.019) | 1.35E-5 | -0.044<br>(0.020) | 2.37E-2 | 4.64 | 3.51E-6 |
| **Cross-Disorder SCZ-BIP-rMDD (European only)** |  |  |  |  |  |  |  |  |  |  |  |  |  |  |  |  |
| rs73033497 | 7 | 2910659 | A/T | 0.86<br>0.14 | intergenic | GNA12 (26.7);<br>CARD11 (35.0) | 13,497<br>(47.22%) | 14,619<br>(48.26%) | 0.267<br>(0.054) | 6.22E-7<br>(2.22E-6) | 0.142<br>(0.039) | 2.55E-4 | -0.129<br>(0.037) | 4.89E-4 | 5.05 | 4.37E-7 |
| rs7302529 | 12 | 77321581 | T/C | 0.26<br>0.26 | intergenic | CSRP2 (48.8);<br>E2F7 (93.4) | 31,541<br>(49.75%) | 31,377<br>(50.42%) | 0.144<br>(0.029) | 7.43E-7<br>(2.32E-6) | 0.094<br>(0.020) | 4.48E-6 | -0.048<br>(0.021) | 2.13E-2 | 4.86 | 1.18E-6 |
| **Cross-Disorder SCZ-BIP-rMDD (European + East Asian)** |  |  |  |  |  |  |  |  |  |  |  |  |  |  |  |  |
| rs8040598 | 15 | 71857368 | A/G | 0.86<br>0.14 | intronic | THSD4 | 41,001<br>(45.92%) | 43,732<br>(50.94%) | 0.183<br>(0.036) | 3.90E-7<br>(8.25E-7) | 0.084<br>(0.026) | 1.18E-3 | -0.093<br>(0.025) | 2.18E-4 | 4.89 | 9.90E-7 |
| rs73033497 | 7 | 2910659 | A/T | 0.86<br>0.14 | intergenic | GNA12 (26.7);<br>CARD11 (35.0) | 13,497<br>(47.22%) | 14,619<br>(48.26%) | 0.267<br>(0.054) | 6.22E-7<br>(2.22E-6) | 0.142<br>(0.039) | 2.55E-4 | -0.129<br>(0.037) | 4.89E-4 | 5.05 | 4.37E-7 |

Omnibus tests of autosomal SNP GxS effects across disorders revealed a significant locus in *NKAIN2* (rs117780815; *p*=3.2×10^−8^ [*p*_*ext*_ =4.67×10^−7^]; **Figure 2**) driven by BIP and SCZ (**Table 2**, **Supplementary Table 7**). The effect was in opposite directions, with the minor allele increasing risk in BIP women and decreasing risk in BIP men, and vice versa in SCZ women and men (see **Table 1**, **Supplementary Table 6a-e**, disorder-specific sex-stratified effects). The second strongest omnibus signal was for the *AMIGO1/GPR61* gene locus (rs12141273; *p*=4.16×10^−7^ [*p*_*ext*_ =1.95×10^−6^]), common to BIP and MDD, though in opposite directions. Of note, omnibus tests of the PGC dataset detected a second strong signal in the *IDO2/C8orf4* gene locus (rs13270586; *p*=1.55×10^−7^ [*p*_*ext*_ = 4.62×10^−7^]), common to BIP and SCZ in opposite directions (**Supplementary Table 16**). Overall, all results from the secondary analytic model supported the primary model.

**Figure 2.**
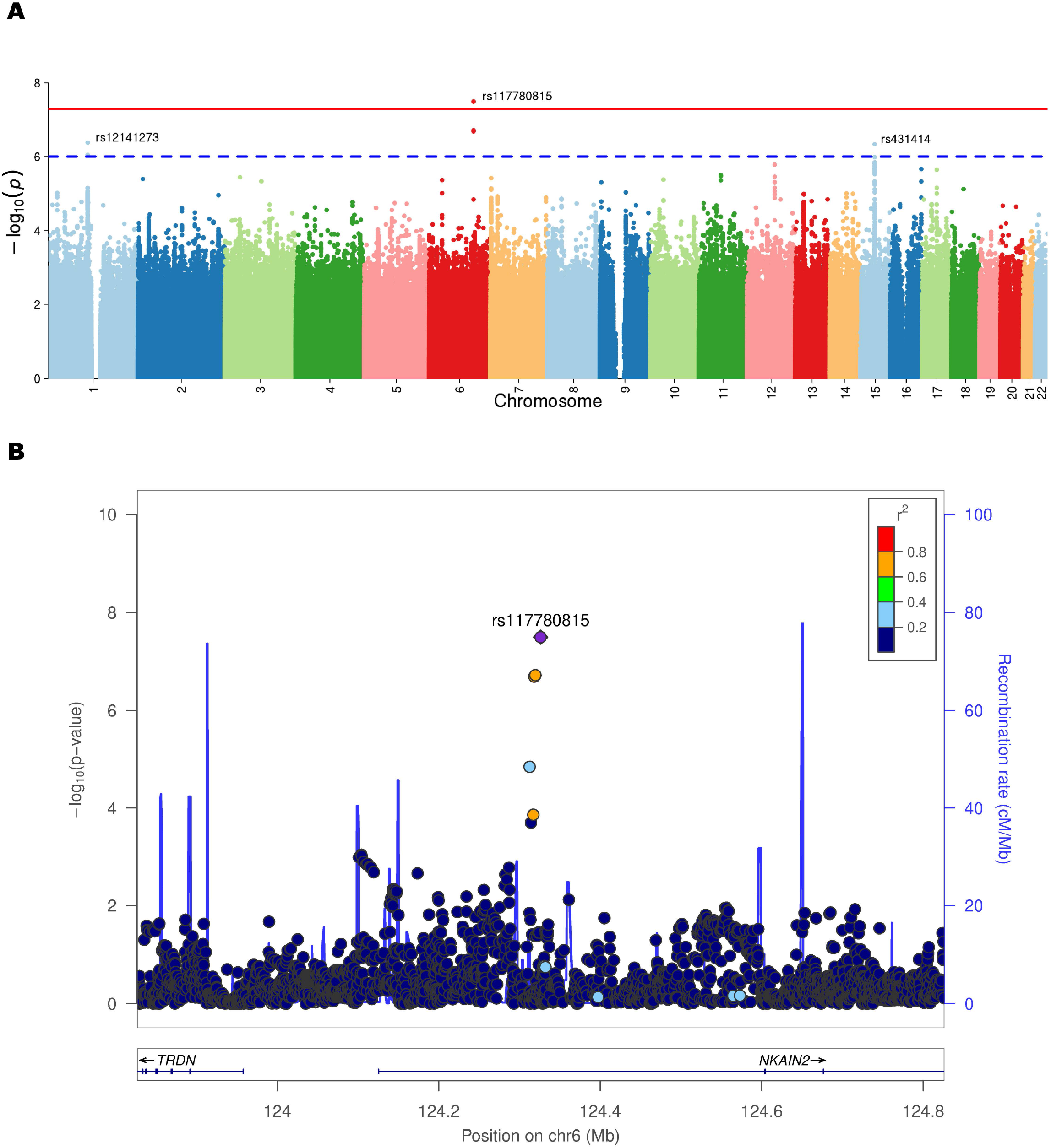
Cross-disorder Manhattan plot of SNP-by-sex interaction p-values **(a)** and LocusZoom plot for the *NKAIN2* gene locus exhibiting a significant SNP-by-sex interaction effect on cross-disorder risk **(b)**. This graph shows the genome-wide significant result from the cross-disorder omnibus test in ASSET (primary model). Negative log10-transformed p-values for each variant (each dot) (y-axis) are plotted by chromosomal position (x-axis). The red and blue lines represent the thresholds for genome-wide significant association (p = 5×10^−8^) and suggestive association (p = 1×10^−6^), respectively. The strongest GxS interaction was found for SNP rs117780815 on chromosome 6 (*p*=3.2×10^−8^) driven by BIP and SCZ. The effect was in opposite directions, with the minor allele increasing risk in BIP women and decreasing risk in BIP men, and vice versa in SCZ women and men (**Table 2, Supplementary Table 7**). Abbreviations: chr = chromosome; cM = centimorgans; Mb = megabases; r^2^ = linkage disequilibrium level; *NKAIN2* = Sodium/Potassium Transporting ATPase Interacting 2

**Table 2.**
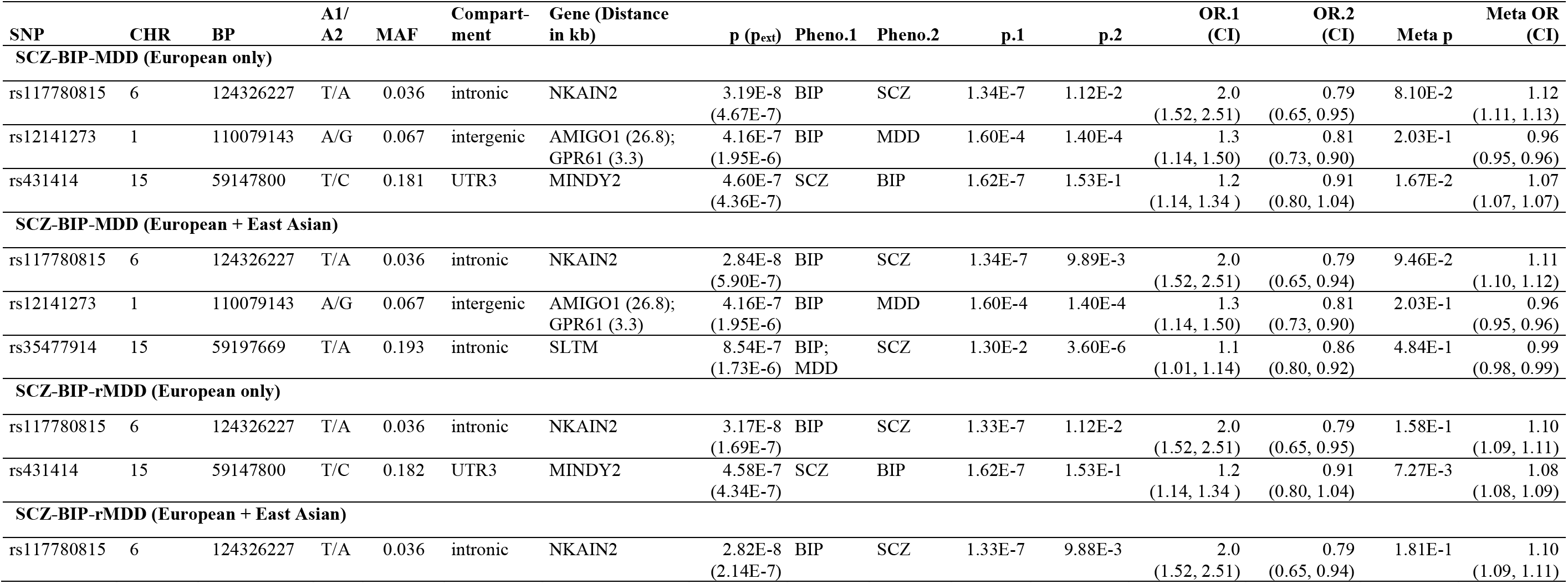
Cross-Disorder Omnibus tests. Omnibus tests were carried out using ASSET, incorporating the within-disorder meta-analysis summary statistics from METAL. Listed are SNPs with cross-disorder interaction *p*-values < 1×10^−6^. Loci were clumped using ‘plink --bfile 1kgp_ref_file --clump asset_output --clump-p1 1e-4 --clump-p2 1e-4 --clump-r2 0.6 --clump-kb 3000’. Extended results (p < 1×10^−4^), including eQTL data for the variants highlighted in this table, and including secondary extended model statistics, are available in **Supplementary Table 7**. Abbreviations: SNP, Variant ID; A1/A2, Allele 1 (reference allele)/Allele 2; CHR, Chromosome; BP, Base Pair Position; p, Omnibus p-value in combined PGC+iPSYCH datasets (p-value for secondary extended model, p_ext_, in parentheses); Pheno.1, Phenotype(s) associated in direction 1; Pheno.2, Phenotype(s) associated in direction 2; p.1, Phenotype(s) 1 p-value; p.2, Phenotype(s) 2 p-value; OR.1 (CI), Phenotype(s) 1 Odds Ratio (Confidence Interval); OR.2 (CI), Phenotype(s) 2 Odds Ratio (Confidence Interval); Meta p, Basic Meta-Analysis p-value; Meta OR (CI), Basic Meta-Analysis Odds Ratio (Confidence Interval)

SNP-by-sex interactions of X chromosome SNPs using model A or B detected only modest effects within/across disorders (lowest *p* = 6.89×10^−6^; **Supplementary Table 8a,b)**, similar regardless of model (**Supplementary Figure 8**). Omnibus tests of X chromosome SNPs detected no significant interactions (lowest *p* = 1.67×10^−5^; **Supplementary Table 9)**.

### Identification of credible SNPs

Loci displaying evidence for GxS interactions (index SNP *p*<1×10^−6^) (**Tables 1–2, Supplementary Tables 6-9**) underwent fine-mapping to identify those SNPs most likely to be causal. Sixteen loci had a mean of 75 (± 68) SNPs. In ~50% of the loci, the index SNP was among the three most credible SNPs, and >70% of clumps had a “simple” model (<=3 causal variants). We summarize the posterior probabilities of all SNPs in fine-mapping loci (**Table 3**, **Supplementary Table 10**) and highlight SNPs with likely causal effects in our disorders. Together, CAVIAR and FINEMAP indicated that genome-wide significant SNP rs117780815, with posterior probability >0.90 (FINEMAP), was the most likely causal variant in the *NKAIN2* locus (see **Table 3**).

**Table 3.** Credible SNP results for genome-wide significant locus *NKAIN2*. CAVIAR and FINEMAP results for the genome-wide significant locus observed in the omnibus test of SCZ, BIP, and MDD (European ancestry). There were four SNPs, including genome-wide significant *NKAIN2* SNP rs117780815, with posterior probability higher than 0.90. These SNPs are the most likely variants to have a causal effect on mood and psychotic disorders from that locus. Abbreviations: Index SNP, genome-wide significant SNP; SNP, all SNPs in locus; A1/A2, Allele 1 (reference allele)/Allele 2; CHR, Chromosome; BP, Base Pair Position; MAF, Minor Allele Frequency; PP(_*ext*_), posterior probability (extended secondary model); SE, Standard Error

| Index SNP | SNP | FINEMAP | CAVIAR | Compartment | Gene | CHR | BP | A1/A2 | MAF | Beta | SE | Z |
| --- | --- | --- | --- | --- | --- | --- | --- | --- | --- | --- | --- | --- |
|  |  | PP causal<br>(PPext) | PP causal<br>(PPext) |  |  |  |  |  |  |  |  |  |
| rs117780815 | rs117780815 | 1<br>(1) | 0.83<br>(0.88) | intronic | NKAIN2 | 6 | 124326227 | T/A | 0.04 | 0.670 | 0.127 | 5.27 |
| rs117780815 | rs4574657 | 1<br>(1) | 5.9E-03<br>(7.2E-03) | intronic | NKAIN2 | 6 | 124319710 | A/G | 0.04 | 0.283 | 0.089 | 3.17 |
| rs117780815 | rs4895382 | 1<br>(1) | 8.0E-02<br>(7.8E-03) | intronic | NKAIN2 | 6 | 124312658 | G/A | 0.02 | 0.736 | 0.171 | 4.29 |
| rs117780815 | rs73557075 | 1<br>(1) | 1.4E-02<br>(5.6E-03) | intronic | NKAIN2 | 6 | 124313730 | A/G | 0.04 | 0.195 | 0.114 | 1.71 |
| rs117780815 | rs7748718 | 6.7E-02<br>(3.5E-02) | 8.8E-03<br>(1.6E-02) | intronic | NKAIN2 | 6 | 124317132 | C/A | 0.05 | 0.358 | 0.108 | 3.33 |
| rs117780815 | rs7754419 | 2.9E-02<br>(5.4E-01) | 6.1E-02<br>(1.6E-01) | intronic | NKAIN2 | 6 | 124318348 | G/A | 0.04 | 0.541 | 0.118 | 4.58 |
| rs117780815 | rs7761506 | 3.7E-02<br>(4.8E-05) | 6.8E-03<br>(7.2E-03) | intronic | NKAIN2 | 6 | 124314413 | G/A | 0.02 | 0.493 | 0.159 | 3.09 |

### Gene- and pathway-based analyses

Gene-based tests within and across disorders detected near-significant GxS interaction of the SLTM gene within SCZ (*p*=4.22×10^−6^ [*p*_*ext*_ =7.28×10^−6^]; **Supplementary Figure 10a**) and genome-wide significant cross-disorder interaction (omnibus *p*=8.97×10^−7^ [*p*_*ext*_ = 6.64×10^−7^]**; Supplementary Figure 10g-h**). No other results approached significance (**Supplementary Table 11; Supplementary Figure 10b-f)**.

Gene set enrichment tests showed that within MDD, GxS SNPs were significantly enriched in genes regulating vascular endothelial growth factor (VEGF) receptor signaling (*p*_FDR_= 3.90×10^−4^ [*p*_FDR*ext*_ = 2.70×10^−2^]; **Supplementary Table 12c)**. SNPs showing GxS interactions within SCZ or BIP were not significantly enriched for any MSigDB pathway (**Supplementary Table 12a,b)**. Across disorders, the ‘wang_barretts_esophagus_and_esophagus_cancer_dn’ pathway showed enrichment (*p*_FDR_ = 0.035 [*p*_FDR*ext*_ = 0.065]; **Supplementary Table 12f)**.

### Brain expression analysis

Brain expression data were examined for genes located adjacent to or encompassing SNPs with evidence for GxS interactions (*p*<1×10^−6^). Most of these genes were expressed in multiple brain regions (**Supplementary Figure 11-13**), particularly prefrontal, anterior cingulate, pituitary, and hypothalamus (**Supplementary Figure 14**) from prenatal development (*C8orf4* [= *TCIM*], *CRSP2*, *GNA12*, *MOCOS, SPAG17*), through puberty (*IDO2)* (**Supplementary Figure 12**), **and** through adulthood. **12-13)**. Genes were expressed in various brain cell types (**Supplementary Figure 15**), with high relative expression of *NKAIN2* and *GNA12* in oligodendrocytes, and *CSRP2, C8orf4* and *MOCOS* in endothelial cells. (**Supplementary Results** reports other genes.)

### eQTL overlap with GxS loci

Examination of eQTL data for SNPs with evidence for GxS interactions (*p*<1×10^−6^; **Supplementary Tables 6-7)** found the highly significant SCZ SNP (rs11665282) in *MOCOS* was a cis-eQTL in several brain regions (**Supplementary Table 6a**) associated with transcriptional elongation and chromatin remodeling in the *ELP2* gene in cerebellum and DLPFC. The most significant cross-disorder SNP (rs7302529) was an eQTL for *CSRP2* (**Supplementary Table 6f**), although the top omnibus cross-disorder SNP (rs117780815) in *NKAIN2* was not an eQTL. Finally, genome-wide SNP rs12141273, intergenic between *AMIGO1* and *GPR61*, is a cis-eQTL for *AMIGO1* in non-brain tissues and associated with expression of glutathione-S-transferase genes *GSTM1* and *GSTM5* and microtubule regulator gene *PSRC1*, in DLPFC (**Supplementary Table 7**).

Overall, consistency of our significant GxS effects with previous GWAS of sex differences in MDD, BIP, and SCZ is described in **Supplementary Results, Table 14**.

## Discussion

Sex differences in incidence, age of onset, symptomatology, and/or brain abnormalities and physiology in SCZ, BIP, and MDD are pervasive (1–7). Previous work demonstrated the impact of gonadal hormones on some of these phenotypic differences. Here, we hypothesized sex differences may, in part, be due to genetic variation, either sex-specific or sex-dependent, and that risk variants may be shared among the disorders.

Heritability estimates were significantly different between the sexes for SCZ and MDD, but not BIP, partly reflecting significant sex differences in incidence for SCZ and MDD, but not BIP. Male-female SNP-based genetic correlations ranged between 0.86 (BIP) and 1 (MDD), significantly <1 for SCZ and BIP but not MDD, with by-sex cross-disorder correlation differences suggesting further complexity. Thus, although the majority of common variant genetic effects were shared between the sexes, there were sex-specific and sex-dependent effects on risks, with modest effect sizes (27).

Significant sex effects, primarily sex-stratified associations, were reported previously in GWAS studies (25–32, 34), implicating neurodevelopmental mechanisms and immune pathways (26–28, 30). However, sex-stratified analyses are only equivalent to GxS interaction tests when there are no interactions between covariates and sex, and the trait variances are equivalent in the two sexes. As this is unlikely, GxS interaction tests are ultimately necessary to identify significant sex differences, and sex-stratified analyses may fail to detect or spuriously report differences.

GxS interaction findings in our study implicate neuronal excitability and inhibitory regulation of brain development and functioning and immune and vascular pathways. Omnibus tests across disorders detected genome-wide significant evidence for GxS emanating from the *NKAIN2* gene, expressed in brain implicating potassium sodium ATPases regulating neuron membrane potential, transmembrane fluxes of Ca^2+^ and excitatory neurotransmitters, and CNS differentiation (50). *NKAIN2* has previously been associated with cognitive ability (51) and SCZ risk (52, 53). The second most significant omnibus GxS result was a SNP adjacent to *AMIGO1*, which regulates activity of the Kv2.1 voltage-dependent potassium channel (54), again important for regulating neuronal excitability in brain (55). Other support for GxS interaction was obtained from gene-based analyses across disorders that detected a genome-wide significant GxS interaction with the *SLTM* gene, a general inhibitor of transcription, highly expressed in cerebellum and putamen, among other regions. Taken together, these findings suggest a sex-dependent genetic contribution to the balance between excitatory and inhibitory regulation of neuronal development and functioning, a hypothesis worthy of further functional “omics” investigations.

In fact, the strongest locus identified in GxS analyses for SCZ (PGC-only; rs13270586) was near *C8orf4* (aka *TCIM*), which functions as a positive regulator of the Wnt/ß-catenin signaling pathway with a central role in fundamental neuronal processes—including synaptogenesis, axon guidance, and dendrite development (56)—and a pathway implicated previously in SCZ, BIP, and MDD (57–60). Interestingly, recent transcriptomic work identified female-biased genes enriched for expression in Cajal-Retzius cells that play a major role in neural migration, whereas male-biased genes were enriched for neural progenitor cells (61). This is consistent with our earlier work in mice with impaired GABA-B receptor signaling and demonstrating sex differences in developmental migration of neurons containing estrogen receptor (ER)-α into the hypothalamus paraventricular nucleus that impacted depressive-like behaviors, particularly in females (62).

Several genes that implicated neuronal excitability and immune functions had opposite effects on disorder risk by sex. For example, the *NKAIN2* SNP GxS effect was opposite in SCZ and BIP, with the minor allele increasing risk in SCZ women and decreasing risk in SCZ men, and opposite effects on risk in BIP women and men. Similarly, the *AMIGO1/GPR61* GxS effect was opposite in BIP and MDD, with the minor allele having stronger effects in BIP women and weaker effect in MDD women versus men.

Immune pathway dysregulation, shared across disorders, also demonstrated some evidence of opposite genetic effects by sex. The strongest GxS interaction for SCZ was in a locus between *IDO2* and *C8orf4* (rs13270586; *p*=1.55×10^−7^), with opposite risk effects by sex. *IDO2* is involved in catabolism of tryptophan in the kynurenine pathway. An end metabolite of the kynurenine pathway, kynurenic acid (KYNA), is elevated in the cerebrospinal fluid (63, 64) and postmortem brains (65, 66) in SCZ and BIP, while reduced plasma levels were associated with depressive symptoms (63). Given recent evidence implicating the kynurenine pathway as a link between brain immune activation and disorder risk (67, 68) and sex differences in immune mechanisms (69), it is plausible that *IDO2* has different effects on SCZ risk in men and women through differential KYNA expression between the sexes. This is consistent with recent findings implicating the complement system (C4) as a source of sexual dimorphisms in vulnerability to SCZ and autoimmune disorders (20). Further, among the strongest results for MDD was a locus spanning *ZNF385C*, associated with transcriptional regulation (70) and immune-related phenotypes via transcriptional enhancers (71, 72).

Our sex-biased genes implicating immune mechanisms at the population level complement recent transcriptomic work in healthy brain development (73), population work in SCZ (19), and MDD (74). Sex-by-diagnosis interactions were seen in the rearrangement of brain transcriptional patterns in MDD (74), an effect also seen in stressed mice (75). In MDD, cell type–specific analyses revealed MDD men exhibited transcriptional increases and MDD women transcriptional decreases in oligodendrocyte- and microglia-related genes (74).

Consistent with this, animal studies demonstrated sex differences in microglia density and morphology in key brain regions beginning in prenatal development (e.g., hypothalamic preoptic area (POA), hippocampus, amygdala). In males *in utero*, there is heightened activation of POA microglia that may result in a priming effect leading to sex-dependent vulnerability for disorders such as SCZ (76). In contrast, while males appear to have a prolonged period of enhanced immune sensitivity *in utero* in *preclinical* studies, the period of immune sensitivity for females is shifted toward the end of prenatal development continuing into early postnatal life in rodents (76), a critical period analogous to human sexual brain differentiation (2^nd^ and 3^rd^ trimesters). This suggests that timing is critical in identifying sex-by-gene effects, which may have opposing effects at different developmental periods, a fact that must be considered in transcription studies of brain regions across the lifespan. In fact, sex differences in expression of IDO2 was identified as also critical during puberty, with post-puberty being the emergence of sex differences in MDD and SCZ.

Other mechanisms that might account for opposing sex interaction effects, include balancing selection due to antagonistic pleiotropic effects (77), that could play a role in maintaining common susceptibility alleles in the population. Opposing effects suggest the potential presence of a ‘genetic switch’ for progression to either one of the diseases, in addition to shared genetic risk factors. Results in autism (78) and SCZ (79) support the idea that these disorders may be opposite extremes of a single gradient of mental disorders or due to diametric gene-dosage deviations caused by disrupted genomic imprinting (78) or copy number variants. Findings suggest that overall sex-specific and sex-dependent genetic correlations may obscure a more complex set of genetic relationships at the level of specific loci, brain regions, and pathways (80), and that timing of mechanisms implicated in sex effects is critical.

Our findings also identified genes associated with vascular development, interesting in light of the comorbidity of CVD with MDD (higher in women) (81) and SCZ. Results demonstrated genes involved in regulation of VEGF signaling were enriched among GxS loci for MDD. Sex differences were reported in VEGF levels (82), and brain expression of VEGF has been associated with cognitive aging and Alzheimer’s disease (83, 84). Further, the strongest GxS interaction was detected for SCZ in a locus in the *MOCOS* gene most highly expressed in endothelial cells lining blood vessels. Interestingly, our previous work on sex differences in neuronal migration due to impaired GABA-B signaling (62) was also significantly associated with sex differences in hypothalamic neurovascular development, being more severe in females and associated with depression-related behaviors (85). In fact, a recent meta-analysis of 22 available gene expression microarrays across multiple organs and tissues cited areas of the brain (i.e., anterior cingulate cortex, implicated in MDD, SCZ and BIP) with the most substantial sex differences in gene expression, followed by the heart (86).

Finally, sex-by-gene effects had implications for cognitive functions, not surprising given brain regions implicated by some of the significant loci in this study. For example, *ZNF385C* in MDD may play a role in cognition, since its paralogs *ZNF385B* and *ZNF385D*, have been associated with intelligence (87), general cognition, mathematical ability and educational attainment (88). It is possible that genes associated with cognitive abnormalities may be shared across disorders, given that the two strongest GxS interaction loci for BIP located near *TUSC1* and *FHL2* have been associated with educational attainment, other cognitive phenotypes, and depression (88, 89).

Although it seems intuitive that genes located on sex chromosomes would be involved in sex differences in disease risk, our analyses did not detect evidence for significant GxS interactions involving X chromosome SNPs. Lack of significance could be due to insensitive X chromosome modeling by sex, thus necessitating more refined models allowing for variability in X inactivation patterns and incorporation of the Y chromosome to clarify the role of sex chromosomes in disease risk. Recent data suggest tissue-specific patterns of X inactivation (90). Nevertheless, our results of GxS interactions for autosomal genes are consistent with transcriptomics data demonstrating sexually dimorphic expression in the brain of a substantial proportion of autosomal genes related to fundamental neural functions (61, 74, 91, 92) and data enriched for tissue-related diseases (92). These findings underscore the utility of studies like ours, with statistical power to test for interaction effects, that highlight genes worthy of deeper mechanistic investigations using transcriptomics and proteomics research and animal models.

A limitation of this study is the relatively low sex-stratified SNP heritability, in particular for MDD men (mean 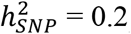). Nevertheless, all heritability estimates were greater than zero with very good precision (i.e., small standard errors), indicating the ability of this study to detect common variant effects. Genetic correlations between the sexes were high and only differed significantly for SCZ and BIP. In the latest PGC SCZ GWAS (93), the cross-sex *r*_*g*_ did not significantly differ from zero, which may, in part, be due to an increased SCZ sample size and different meta-analysis composition. While genetic correlations between the sexes within-disorder were high, most striking were the differences in genetic correlations by disorder by sex. High genetic correlations were observed between MDD (both sexes) and BIP women (0.42, 0.48), but much weaker with MDD (both sexes) and BIP men (0.13, 0.04). Although some have argued this may reflect study recruitment bias or misclassification (94), this is less likely for our study, given varying sample sizes across disorders (due to differing prevalences), and no genetic correlations by sex among SCZ compared with high correlations among MDD and BIP. Misclassification of cases is always a possibility, although clinical diagnoses were based on extensive DSM-IV or ICD-10 interviews, limiting the likelihood of this. Further, if there were bias, it would require similar and substantial bias across multiple international institutions.

The lack of detailed clinical data prevented examination of important questions related to symptom type, severity, age at onset, and cognitive deficits. These limitations emphasize the need for larger, deeply-phenotyped datasets to fully characterize sex differences in genetic and *clinical* characteristics of these disorders, as highlighted recently in (27). Further, alternative explanations for sex differences in incidence, presentation, and course, include genotype-by-environment interactions, e.g., implicating gonadal hormone regulation of genes, that we know from clinical and animal studies are sex-dependent. Finally, additional replication samples would significantly strengthen these findings.

### Conclusions

In the largest genome-wide GxS analysis of mood and psychotic disorders to date, we found substantial genetic overlap between men and women for SCZ, BIP, and MDD. However, we also found several loci with significant GxS interaction effects across and within disorder – *NKAIN2* at the variant level, *SLTM* at the gene level, and *VEGF* at pathway level. Functional genomics suggests that all genes were expressed in at least one brain region at some period across the lifespan, with most genes expressed in multiple brain regions associated with mood/anxiety and cognition.

Our results demonstrate that the risk for SCZ, MDD and BIP is impacted by interactions of genotype with sex, beyond the impact of gonadal steroid hormones. Though specific mechanisms remain unknown, our study underscores the importance of designing large-scale genetic studies that have the statistical power to test for interactions with sex. Dissecting the impact of sex, genes, and pathophysiology will identify potential targets for sex-dependent or sex-specific therapeutic interventions.

#### Resources

Summary statistics are available for download from https://www.med.unc.edu/pgc/ upon publication.

## Acknowledgments

These analyses were supported by a private donor, Ms. Gwill York, awarded to Dr. Jill Goldstein (JMG). JMG, ST, and RH’s time on these analyses was also supported, in part, by ORWH-NIMH U54 MH118919 (Goldstein and Handa, multi-PIs), and TP’s time by NIMH R01 MH092380 (Petryshen, PI). Funding sources for individual cohorts are provided in reference numbers (34–36).

## Consortia

### Schizophrenia Working Group of the Psychiatric Genomics Consortium

Stephan Ripke, Benjamin M. Neale, Aiden Corvin, James T. R. Walters, Kai-How Farh, Peter A. Holmans, Phil Lee, Brendan Bulik-Sullivan, David A. Collier, Hailiang Huang, Tune H. Pers, Ingrid Agartz, Esben Agerbo, Margot Albus, Madeline Alexander, Farooq Amin, Silviu A. Bacanu, Martin Begemann, Richard A. Belliveau Jr, Judit Bene, Sarah E. Bergen, Elizabeth Bevilacqua, Tim B. Bigdeli, Donald W. Black, Richard Bruggeman, Nancy G. Buccola, Randy L. Buckner, William Byerley, Wiepke Cahn, Guiqing Cai, Dominique Campion, Rita M. Cantor, Vaughan J. Carr, Noa Carrera, Stanley V. Catts, Kimberly D. Chambert, Raymond C. K. Chan, Ronald Y. L. Chen, Eric Y. H. Chen, Wei Cheng, Eric F. C. Cheung, Siow Ann Chong, C. Robert Cloninger, David Cohen, Nadine Cohen, Paul Cormican, Nick Craddock, James J. Crowley, David Curtis, Michael Davidson, Kenneth L. Davis, Franziska Degenhardt, Jurgen Del Favero, Ditte Demontis, Dimitris Dikeos, Timothy Dinan, Srdjan Djurovic, Gary Donohoe, Elodie Drapeau, Jubao Duan, Frank Dudbridge, Naser Durmishi, Peter Eichhammer, Johan Eriksson, Valentina Escott-Price, Laurent Essioux, Ayman H. Fanous, Martilias S. Farrell, Josef Frank, Lude Franke, Robert Freedman, Nelson B. Freimer, Marion Friedl, Joseph I. Friedman, Menachem Fromer, Giulio Genovese, Lyudmila Georgieva, Ina Giegling, Paola Giusti-Rodríguez, Stephanie Godard, Jacqueline I. Goldstein, Vera Golimbet, Srihari Gopal, Jacob Gratten, Lieuwe de Haan, Christian Hammer, Marian L. Hamshere, Mark Hansen, Thomas Hansen, Vahram Haroutunian, Annette M. Hartmann, Frans A. Henskens, Stefan Herms, Joel N. Hirschhorn, Per Hoffmann, Andrea Hofman, Mads V. Hollegaard, David M. Hougaard, Masashi Ikeda, Inge Joa, Antonio Julià, René S. Kahn, Luba Kalaydjieva, Sena Karachanak-Yankova, Juha Karjalainen, David Kavanagh, Matthew C. Keller, James L. Kennedy, Andrey Khrunin, Yunjung Kim, Janis Klovins, James A. Knowles, Bettina Konte, Vaidutis Kucinskas, Zita Ausrele Kucinskiene, Hana Kuzelova-Ptackova, Anna K. Kähler, Claudine Laurent, Jimmy Lee Chee Keong, S. Hong Lee, Sophie E. Legge, Bernard Lerer, Miaoxin Li, Tao Li, Kung-Yee Liang, Jeffrey Lieberman, Svetlana Limborska, Carmel M. Loughland, Jan Lubinski, Jouko Lönnqvist, Milan Macek Jr, Patrik K. E. Magnusson, Brion S. Maher, Wolfgang Maier, Jacques Mallet, Sara Marsal, Manuel Mattheisen, Morten Mattingsdal, Robert W. McCarley†, Colm McDonald, Andrew M. McIntosh, Sandra Meier, Carin J. Meijer, Bela Melegh, Ingrid Melle, Raquelle I. Mesholam-Gately, Andres Metspalu, Patricia T. Michie, Lili Milani, Vihra Milanova, Younes Mokrab, Derek W. Morris, Ole Mors, Kieran C. Murphy, Robin M. Murray, Inez Myin-Germeys, Bertram Müller-Myhsok, Mari Nelis, Igor Nenadic, Deborah A. Nertney, Gerald Nestadt, Kristin K. Nicodemus, Liene Nikitina-Zake, Laura Nisenbaum, Annelie Nordin, Eadbhard O’Callaghan, Colm O’Dushlaine, F. Anthony O’Neill, Sang-Yun Oh, Ann Olincy, Line Olsen, Jim Van Os, Psychosis Endophenotypes International Consortium, Christos Pantelis, George N. Papadimitriou, Sergi Papiol, Elena Parkhomenko, Michele T. Pato, Tiina Paunio, Milica Pejovic-Milovancevic, Diana O. Perkins, Olli Pietiläinen, Jonathan Pimm, Andrew J. Pocklington, John Powell, Alkes Price, Ann E. Pulver, Shaun M. Purcell, Digby Quested, Henrik B. Rasmussen, Abraham Reichenberg, Mark A. Reimers, Alexander L. Richards, Joshua L. Roffman, Panos Roussos, Douglas M. Ruderfer, Veikko Salomaa, Alan R. Sanders, Ulrich Schall, Christian R. Schubert, Thomas G. Schulze, Sibylle G. Schwab, Edward M. Scolnick, Rodney J. Scott, Larry J. Seidman†, Jianxin Shi, Engilbert Sigurdsson, Teimuraz Silagadze, Jeremy M. Silverman, Kang Sim, Petr Slominsky, Jordan W. Smoller, Hon-Cheong So, Chris C. A. Spencer, Eli A. Stahl, Hreinn Stefansson, Stacy Steinberg, Elisabeth Stogmann, Richard E. Straub, Eric Strengman, Jana Strohmaier, T. Scott Stroup, Mythily Subramaniam, Jaana Suvisaari, Dragan M. Svrakic, Jin P. Szatkiewicz, Erik Söderman, Srinivas Thirumalai, Draga Toncheva, Sarah Tosato, Juha Veijola, John Waddington, Dermot Walsh, Dai Wang, Qiang Wang, Bradley T. Webb, Mark Weiser, Dieter B. Wildenauer, Nigel M. Williams, Stephanie Williams, Stephanie H. Witt, Aaron R. Wolen, Emily H. M. Wong, Brandon K. Wormley, Hualin Simon Xi, Clement C. Zai, Xuebin Zheng, Fritz Zimprich, Naomi R. Wray, Kari Stefansson, Peter M. Visscher, Wellcome Trust Case-Control Consortium, Rolf Adolfsson, Ole A. Andreassen, Douglas H. R. Blackwood, Elvira Bramon, Joseph D. Buxbaum, Anders D. Børglum, Sven Cichon, Ariel Darvasi†, Enrico Domenici, Hannelore Ehrenreich, Tõnu Esko, Pablo V. Gejman, Michael Gill, Hugh Gurling, Christina M. Hultman, Nakao Iwata, Assen V. Jablensky, Erik G. Jönsson, Kenneth S. Kendler, George Kirov, Jo Knight, Todd Lencz, Douglas F. Levinson, Qingqin S. Li, Jianjun Liu, Anil K. Malhotra, Steven A. McCarroll, Andrew McQuillin, Jennifer L. Moran, Preben B. Mortensen, Sathish Periyasamy, Murray J. Cairns, Paul A. Tooney, Jing Qin Wu, Brian Kelly, Bryan J. Mowry, Markus M. Nöthen, Roel A. Ophoff, Michael J. Owen, Aarno Palotie, Carlos N. Pato, Tracey L. Petryshen, Danielle Posthuma, Marcella Rietschel, Brien P. Riley, Dan Rujescu, Pak C. Sham, Pamela Sklar, David St Clair, Daniel R. Weinberger, Jens R. Wendland, Thomas Werge, Mark J. Daly, Patrick F. Sullivan, Michael C. O’Donovan.

#### Wellcome Trust Case-Control Consortium

Management Committee: Peter Donnelly, Ines Barroso, Jenefer M. Blackwell, Elvira Bramon, Matthew A. Brown, Juan P. Casas, Aiden Corvin, Panos Deloukas, Audrey Duncanson, Janusz Jankowski, Hugh S. Markus, Christopher G. Mathew, Colin N. A. Palmer, Robert Plomin, Anna Rautanen, Stephen J. Sawcer, Richard C. Trembath, Ananth C. Viswanathan, Nicholas W. Wood. Data and Analysis Group: Chris C. A. Spencer, Gavin Band, Céline Bellenguez, Peter Donnelly, Colin Freeman, Eleni Giannoulatou, Garrett Hellenthal, Richard Pearson, Matti Pirinen, Amy Strange, Zhan Su, Damjan Vukcevic. DNA, Genotyping, Data QC, and Informatics: Cordelia Langford, Ines Barroso, Hannah Blackburn, Suzannah J. Bumpstead, Panos Deloukas, Serge Dronov, Sarah Edkins, Matthew Gillman, Emma Gray, Rhian Gwilliam, Naomi Hammond, Sarah E. Hunt, Alagurevathi Jayakumar, Jennifer Liddle, Owen T. McCann, Simon C. Potter, Radhi Ravindrarajah, Michelle Ricketts, Avazeh Tashakkori-Ghanbaria, Matthew Waller, Paul Weston, Pamela Whittaker, Sara Widaa. Publications Committee: Christopher G. Mathew, Jenefer M. Blackwell, Matthew A. Brown, Aiden Corvin, Mark I. McCarthy, Chris C. A. Spencer.

#### Psychosis Endophenotype International Consortium

Maria J. Arranz, Steven Bakker, Stephan Bender, Elvira Bramon, David A. Collier, Benedicto Crespo-Facorro, Jeremy Hall, Conrad Iyegbe, Assen V. Jablensky, René S. Kahn, Luba Kalaydjieva, Stephen Lawrie, Cathryn M. Lewis, Kuang Lin, Don H. Linszen, Ignacio Mata, Andrew M. McIntosh, Robin M. Murray, Roel A. Ophoff, Jim Van Os, John Powell, Dan Rujescu, Muriel Walshe, Matthias Weisbrod, Durk Wiersma.

### Bipolar Disorder Working Group of the Psychiatric Genomics Consortium

Eli A. Stahl, Gerome Breen, Andreas J. Forstner, Andrew McQuillin, Stephan Ripke, Vassily Trubetskoy, Manuel Mattheisen, Yunpeng Wang, Jonathan R. I. Coleman, Héléna A. Gaspar, Christiaan A. de Leeuw, Stacy Steinberg, Jennifer M. Whitehead Pavlides, Maciej Trzaskowski, Enda M. Byrne, Tune H. Pers, Peter A. Holmans, Alexander L. Richards, Liam Abbott, Esben Agerbo, Huda Akil, Diego Albani, Ney Alliey-Rodriguez, Thomas D. Als, Adebayo Anjorin, Verneri Antilla, Swapnil Awasthi, Judith A. Badner, Marie Bækvad-Hansen, Jack D. Barchas, Nicholas Bass, Michael Bauer, Richard Belliveau, Sarah E. Bergen, Carsten Bøcker Pedersen, Erlend Bøen, Marco P. Boks, James Boocock, Monika Budde, William Bunney, Margit Burmeister, Jonas Bybjerg-Grauholm, William Byerley, Miquel Casas, Felecia Cerrato, Pablo Cervantes, Kimberly Chambert, Alexander W. Charney, Danfeng Chen, Claire Churchhouse, Toni-Kim Clarke, William Coryell, David W. Craig, Cristiana Cruceanu, David Curtis, Piotr M. Czerski, Anders M. Dale, Simone de Jong, Franziska Degenhardt, Jurgen Del-Favero, J. Raymond DePaulo, Srdjan Djurovic, Amanda L. Dobbyn, Ashley Dumont, Torbjørn Elvsåshagen, Valentina Escott-Price, Chun Chieh Fan, Sascha B. Fischer, Matthew Flickinger, Tatiana M. Foroud, Liz Forty, Josef Frank, Christine Fraser, Nelson B. Freimer, Katrin Gade, Diane Gage, Julie Garnham, Claudia Giambartolomei, Marianne Giørtz Pedersen, Jaqueline Goldstein, Scott D. Gordon, Katherine Gordon-Smith, Elaine K. Green, Melissa J. Green, Tiffany A. Greenwood, Jakob Grove, Weihua Guan, José Guzman-Parra, Marian L. Hamshere, Martin Hautzinger, Urs Heilbronner, Stefan Herms, Maria Hipolito, Per Hoffmann, Dominic Holland, Laura Huckins, Stéphane Jamain, Jessica S. Johnson, Radhika Kandaswamy, Robert Karlsson, James L. Kennedy, Sarah Kittel-Schneider, James A. Knowles, Manolis Kogevinas, Anna C. Koller, Ralph Kupka, Catharina Lavebratt, Jacob Lawrence, William B. Lawson, Markus Leber, Phil H. Lee, Shawn E. Levy, Jun Z. Li, Chunyu Liu, Susanne Lucae, Anna Maaser, Donald J. MacIntyre, Pamela B. Mahon, Wolfgang Maier, Lina Martinsson, Steve McCarroll, Peter McGuffin, Melvin G. McInnis, James D. McKay, Helena Medeiros, Sarah E. Medland, Fan Meng, Lili Milani, Grant W. Montgomery, Derek W. Morris, Thomas W. Mühleisen, Niamh Mullins, Hoang Nguyen, Caroline M. Nievergelt, Annelie Nordin Adolfsson, Evaristus A. Nwulia, Claire O’Donovan, Loes M. Olde Loohuis, Anil P. S. Ori, Lilijana Oruc, Urban Ösby, Roy H. Perlis, Amy Perry, Andrea Pfennig, James B. Potash, Shaun M. Purcell, Eline J. Regeer, Andreas Reif, Céline S. Reinbold, John P. Rice, Fabio Rivas, Margarita Rivera, Panos Roussos, Douglas M. Ruderfer, Euijung Ryu, Cristina Sánchez-Mora, Alan F. Schatzberg, William A. Scheftner, Nicholas J. Schork, Cynthia Shannon Weickert, Tatyana Shehktman, Paul D. Shilling, Engilbert Sigurdsson, Claire Slaney, Olav B. Smeland, Janet L. Sobell, Christine Søholm Hansen, Anne T. Spijker, David St Clair, Michael Steffens, John S. Strauss, Fabian Streit, Jana Strohmaier, Szabolcs Szelinger, Robert C. Thompson, Thorgeir E. Thorgeirsson, Jens Treutlein, Helmut Vedder, Weiqing Wang, Stanley J. Watson, Thomas W. Weickert, Stephanie H. Witt, Simon Xi, Wei Xu, Allan H. Young, Peter Zandi, Peng Zhang, Sebastian Zöllner, eQTLGen Consortium, BIOS Consortium, Rolf Adolfsson, Ingrid Agartz, Martin Alda, Lena Backlund, Bernhard T. Baune, Frank Bellivier, Wade H. Berrettini, Joanna M. Biernacka, Douglas H. R. Blackwood, Michael Boehnke, Anders D. Børglum, Aiden Corvin, Nicholas Craddock, Mark J. Daly, Udo Dannlowski, Tõnu Esko, Bruno Etain, Mark Frye, Janice M. Fullerton, Elliot S. Gershon, Michael Gill, Fernando Goes, Maria Grigoroiu-Serbanescu, Joanna Hauser, David M. Hougaard, Christina M. Hultman, Ian Jones, Lisa A. Jones, René S. Kahn, George Kirov, Mikael Landén, Marion Leboyer, Cathryn M. Lewis, Qingqin S. Li, Jolanta Lissowska, Nicholas G. Martin, Fermin Mayoral, Susan L. McElroy, Andrew M. McIntosh, Francis J. McMahon, Ingrid Melle, Andres Metspalu, Philip B. Mitchell, Gunnar Morken, Ole Mors, Preben Bo Mortensen, Bertram Müller-Myhsok, Richard M. Myers, Benjamin M. Neale, Vishwajit Nimgaonkar, Merete Nordentoft, Markus M. Nöthen, Michael C. O’Donovan, Ketil J. Oedegaard, Michael J. Owen, Sara A. Paciga, Carlos Pato, Michele T. Pato, Danielle Posthuma, Josep Antoni Ramos-Quiroga, Marta Ribasés, Marcella Rietschel, Guy A. Rouleau, Martin Schalling, Peter R. Schofield, Thomas G. Schulze, Alessandro Serretti, Jordan W. Smoller, Hreinn Stefansson, Kari Stefansson, Eystein Stordal, Patrick F. Sullivan, Gustavo Turecki, Arne E. Vaaler, Eduard Vieta, John B. Vincent, Thomas Werge, John I. Nurnberger, Naomi R. Wray, Arianna Di Florio, Howard J. Edenberg, Sven Cichon, Roel A. Ophoff, Laura J. Scott, Ole A. Andreassen, John Kelsoe, Pamela Sklar†.

### Major Depressive Disorder Working Group of the Psychiatric Genomics Consortium

Naomi R. Wray, Stephan Ripke, Manuel Mattheisen, Maciej Trzaskowski, Enda M. Byrne, Abdel Abdellaoui, Mark J. Adams, Esben Agerbo, Tracy M. Air, Till F. M. Andlauer, Silviu-Alin Bacanu, Marie Bækvad-Hansen, Aartjan T. F. Beekman, Tim B. Bigdeli, Elisabeth B. Binder, Julien Bryois, Henriette N. Buttenschøn, Jonas Bybjerg-Grauholm, Na Cai, Enrique Castelao, Jane Hvarregaard Christensen, Toni-Kim Clarke, Jonathan R. I. Coleman, Lucía Colodro-Conde, Baptiste Couvy-Duchesne, Nick Craddock, Gregory E. Crawford, Gail Davies, Ian J. Deary, Franziska Degenhardt, Eske M. Derks, Nese Direk, Conor V. Dolan, Erin C. Dunn, Thalia C. Eley, Valentina Escott-Price, Farnush Farhadi Hassan Kiadeh, Hilary K. Finucane, Jerome C. Foo, Andreas J. Forstner, Josef Frank, Héléna A. Gaspar, Michael Gill, Fernando S. Goes, Scott D. Gordon, Jakob Grove, Lynsey S. Hall, Christine Søholm Hansen, Thomas F. Hansen, Stefan Herms, Ian B. Hickie, Per Hoffmann, Georg Homuth, Carsten Horn, Jouke-Jan Hottenga, David M. Hougaard, David M. Howard, Marcus Ising, Rick Jansen, Ian Jones, Lisa A. Jones, Eric Jorgenson, James A. Knowles, Isaac S. Kohane, Julia Kraft, Warren W. Kretzschmar, Zoltán Kutalik, Yihan Li, Penelope A. Lind, Donald J. MacIntyre, Dean F. MacKinnon, Robert M. Maier, Wolfgang Maier, Jonathan Marchini, Hamdi Mbarek, Patrick McGrath, Peter McGuffin, Sarah E. Medland, Divya Mehta, Christel M. Middeldorp, Evelin Mihailov, Yuri Milaneschi, Lili Milani, Francis M. Mondimore, Grant W. Montgomery, Sara Mostafavi, Niamh Mullins, Matthias Nauck, Bernard Ng, Michel G. Nivard, Dale R. Nyholt, Paul F. O’Reilly, Hogni Oskarsson, Michael J. Owen, Jodie N. Painter, Carsten Bøcker Pedersen, Marianne Giørtz Pedersen, Roseann E. Peterson, Wouter J. Peyrot, Giorgio Pistis, Danielle Posthuma, Jorge A. Quiroz, Per Qvist, John P. Rice, Brien P. Riley, Margarita Rivera, Saira Saeed Mirza, Robert Schoevers, Eva C. Schulte, Ling Shen, Jianxin Shi, Stanley I. Shyn, Engilbert Sigurdsson, Grant C. B. Sinnamon, Johannes H. Smit, Daniel J. Smith, Hreinn Stefansson, Stacy Steinberg, Fabian Streit, Jana Strohmaier, Katherine E. Tansey, Henning Teismann, Alexander Teumer, Wesley Thompson, Pippa A. Thomson, Thorgeir E. Thorgeirsson, Matthew Traylor, Jens Treutlein, Vassily Trubetskoy, André G. Uitterlinden, Daniel Umbricht, Sandra Van der Auwera, Albert M. van Hemert, Alexander Viktorin, Peter M. Visscher, Yunpeng Wang, Bradley T. Webb, Shantel Marie Weinsheimer, Jürgen Wellmann, Gonneke Willemsen, Stephanie H. Witt, Yang Wu, Hualin S. Xi, Jian Yang, Futao Zhang, Volker Arolt, Bernhard T. Baune, Klaus Berger, Dorret I. Boomsma, Sven Cichon, Udo Dannlowski, Eco JC de Geus, J. Raymond DePaulo, Enrico Domenici, Katharina Domschke, Tõnu Esko, Hans J. Grabe, Steven P. Hamilton, Caroline Hayward, Andrew C. Heath, Kenneth S. Kendler, Stefan Kloiber, Glyn Lewis, Qingqin S. Li, Susanne Lucae, Pamela AF Madden, Patrik K. Magnusson, Nicholas G. Martin, Andrew M. McIntosh, Andres Metspalu, Ole Mors, Preben Bo Mortensen, Bertram Müller-Myhsok, Merete Nordentoft, Markus M. Nöthen, Michael C. O’Donovan, Sara A. Paciga, Nancy L. Pedersen, Brenda WJH Penninx, Roy H. Perlis, David J. Porteous, James B. Potash, Martin Preisig, Marcella Rietschel, Catherine Schaefer, Thomas G. Schulze, Jordan W. Smoller, Kari Stefansson, Henning Tiemeier, Rudolf Uher, Henry Völzke, Myrna M. Weissman, Thomas Werge, Cathryn M. Lewis, Douglas F. Levinson, Gerome Breen, Anders D. Børglum, Patrick F. Sullivan.

† deceased

### Sex differences cross-disorder analysis group of the Psychiatric Genomics Consortium

Martin Alda, Gabriëlla A. M. Blokland, Anders D. Børglum, Marco Bortolato, Janita Bralten, Gerome Breen, Cynthia M. Bulik, Christie L. Burton, Enda M. Byrne, Caitlin E. Carey, Jonathan R. I. Coleman, Lea K. Davis, Ditte Demontis, Laramie E. Duncan, Howard J. Edenberg, Lauren Erdman, Stephen V. Faraone, Jill M. Goldstein, Slavina B. Goleva, Jakob Grove, Wei Guo, Christopher Hübel, Laura M. Huckins, Ekaterina A. Khramtsova, Phil H. Lee, Joanna Martin, Carol A. Mathews, Manuel Mattheisen, Benjamin M. Neale, Roseann E. Peterson, Tracey L. Petryshen, Elise Robinson, Jordan W. Smoller, Eli Stahl, Barbara E. Stranger, Michela Traglia, Raymond K. Walters, Lauren A. Weiss, Thomas Werge, Stacey J. Winham, Naomi R. Wray, Yin Yao.

### iPSYCH

#### Management Group

Anders D. Børglum, David M. Hougaard, Merete Nordentoft, Ole Mors, Preben Bo Mortensen, Thomas Werge, Kristjar Skajaa. *Advisory Board:* Markus Nöthen, Michael Owen, Robert H. Yolken, Niels Plath, Jonathan Mill, Daniel Geschwind. Affiliations for all consortium members and acknowledgements for specific cohorts are provided in the Supplement.

## Disclosures

All authors declare that they have no conflicts of interest. JG is on the scientific advisory board for and has equity in Cala Health, a neuromodulation company, although this is unrelated to the topic in this study; and TLP is an employee of Concert Pharmaceuticals, also unrelated to this study. JWS is an unpaid member of the Bipolar/Depression Research Community Advisory Panel of 23andMe.

## Notes

### Summary of Updates

Results now include secondary regression model, with additional covariates. Results, including tables and figures, reordered. Table 3 added. Discussion updated. Supplemental files updated.

## References

1. Salk RH, Hyde JS, Abramson LY (2017): Gender differences in depression in representative national samples: Meta-analyses of diagnoses and symptoms. Psychol Bull. 143:783–822.

2. Jongsma HE, Turner C, Kirkbride JB, Jones PB (2019): International incidence of psychotic disorders, 2002-17: A systematic review and meta-analysis. Lancet Public Health. 4:e229–e244.

3. Diflorio A, Jones I (2010): Is sex important? Gender differences in bipolar disorder. Int Rev Psychiatry. 22:437–452.

4. Erol A, Winham SJ, McElroy SL, Frye MA, Prieto ML, Cuellar-Barboza AB, et al. (2015): Sex differences in the risk of rapid cycling and other indicators of adverse illness course in patients with bipolar I and II disorder. Bipolar Disord. 17:670–676.

5. Falkenburg J, Tracy DK (2014): Sex and schizophrenia: A review of gender differences. Psychosis. 6:61–69.

6. Leung A, Chue P (2000): Sex differences in schizophrenia, a review of the literature. Acta Psychiatr Scand Suppl. 401:3–38.

7. Schuch JJ, Roest AM, Nolen WA, Penninx BW, de Jonge P (2014): Gender differences in major depressive disorder: Results from the Netherlands study of depression and anxiety. J Affect Disord. 156:156–163.

8. Mareckova K, Holsen L, Admon R, Whitfield-Gabrieli S, Seidman LJ, Buka SL, et al. (2017): Neural - hormonal responses to negative affective stimuli: Impact of dysphoric mood and sex. J Affect Disord. 222:88–97.

9. Mareckova K, Holsen LM, Admon R, Makris N, Seidman L, Buka S, et al. (2016): Brain activity and connectivity in response to negative affective stimuli: Impact of dysphoric mood and sex across diagnoses. Hum Brain Mapp. 37:3733–3744.

10. Polderman TJ, Benyamin B, de Leeuw CA, Sullivan PF, van Bochoven A, Visscher PM, et al. (2015): Meta-analysis of the heritability of human traits based on fifty years of twin studies. Nat Genet. 47:702–709.

11. Vink JM, Bartels M, van Beijsterveldt TC, van Dongen J, van Beek JH, Distel MA, et al. (2012): Sex differences in genetic architecture of complex phenotypes? PLoS One. 7:e47371.

12. Stringer S, Polderman TJC, Posthuma D (2017): Majority of human traits do not show evidence for sex-specific genetic and environmental effects. Sci Rep. 7:8688.

13. Weiss LA, Pan L, Abney M, Ober C (2006): The sex-specific genetic architecture of quantitative traits in humans. Nat Genet. 38:218–222.

14. Yang J, Bakshi A, Zhu Z, Hemani G, Vinkhuyzen AA, Nolte IM, et al. (2015): Genome-wide genetic homogeneity between sexes and populations for human height and body mass index. Hum Mol Genet. 24:7445–7449.

15. Goldstein JM, Faraone SV, Chen WJ, Tsuang MT (1995): Genetic heterogeneity may in part explain sex differences in the familial risk for schizophrenia. Biol Psychiatry. 38:808–813.

16. Goldstein JM, Cherkerzian S, Tsuang MT, Petryshen TL (2013): Sex differences in the genetic risk for schizophrenia: History of the evidence for sex-specific and sex-dependent effects. Am J Med Genet B Neuropsychiatr Genet. 162B:698–710.

17. Goldstein JM (1997): Sex differences in schizophrenia: Epidemiology, genetics and the brain. Int Rev Psychiatr. 9:399–408.

18. Goldstein JM, Seidman LJ, O’Brien LM, Horton NJ, Kennedy DN, Makris N, et al. (2002): Impact of normal sexual dimorphisms on sex differences in structural brain abnormalities in schizophrenia assessed by magnetic resonance imaging. Arch Gen Psychiatry. 59:154–164.

19. Sekar A, Bialas AR, de Rivera H, Davis A, Hammond TR, Kamitaki N, et al. (2016): Schizophrenia risk from complex variation of complement component 4. Nature. 530:177–183.

20. Kamitaki N, Sekar A, Handsaker RE, Rivera Hd, Tooley K, Morris DL, et al. (2020): Complement genes contribute sex-biased vulnerability in diverse disorders. Nature. 582:577–581.

21. van Loo HM, Aggen SH, Gardner CO, Kendler KS (2018): Sex similarities and differences in risk factors for recurrence of major depression. Psychol Med. 48:1685–1693.

22. Bertschy G, Velten M, Weibel S (2016): Major depression: Does gender influence the risk of recurrence? A systematic review. Eur J Psychiat. 30:7–27.

23. Smith DJ, Nicholl BI, Cullen B, Martin D, Ul-Haq Z, Evans J, et al. (2013): Prevalence and characteristics of probable major depression and bipolar disorder within UK biobank: Cross-sectional study of 172,751 participants. PLoS One. 8:e75362.

24. Duncan LE, Ratanatharathorn A, Aiello AE, Almli LM, Amstadter AB, Ashley-Koch AE, et al. (2018): Largest GWAS of PTSD (N=20 070) yields genetic overlap with schizophrenia and sex differences in heritability. Mol Psychiatry. 23:666–673.

25. Nievergelt CM, Maihofer AX, Klengel T, Atkinson EG, Chen CY, Choi KW, et al. (2019): International meta-analysis of PTSD genome-wide association studies identifies sex- and ancestry-specific genetic risk loci. Nat Commun. 10:4558.

26. Mitra I, Tsang K, Ladd-Acosta C, Croen LA, Aldinger KA, Hendren RL, et al. (2016): Pleiotropic mechanisms indicated for sex differences in autism. PLoS Genet. 12:e1006425.

27. Khramtsova EA, Heldman R, Derks EM, Yu D, Tourette Syndrome/Obsessive-Compulsive Disorder Working Group of the Psychiatric Genomics C, Davis LK, et al. (2018): Sex differences in the genetic architecture of obsessive-compulsive disorder. Am J Med Genet B Neuropsychiatr Genet. 180:351–364.

28. Hyde CL, Nagle MW, Tian C, Chen X, Paciga SA, Wendland JR, et al. (2016): Identification of 15 genetic loci associated with risk of major depression in individuals of European descent. Nat Genet. 48:1031–1036.

29. Walters R, Abbott L, Bryant S, Churchhouse C, Palmer D, Neale B (2018): Heritability of >2,000 traits and disorders in the UK Biobank. http://wwwnealelabis/uk-biobank/.

30. Hübel C, Gaspar HA, Coleman JRI, Finucane H, Purves KL, Hanscombe KB, et al. (2018): Genomics of body fat percentage may contribute to sex bias in anorexia nervosa. Am J Med Genet B Neuropsychiatr Genet. 180:428–438.

31. Trzaskowski M, Mehta D, Peyrot WJ, Hawkes D, Davies D, Howard DM, et al. (2019): Quantifying between-cohort and between-sex genetic heterogeneity in major depressive disorder. Am J Med Genet B Neuropsychiatr Genet. 180:439–447.

32. Martin J, Walters RK, Demontis D, Mattheisen M, Lee SH, Robinson E, et al. (2018): A genetic investigation of sex bias in the prevalence of attention-deficit/hyperactivity disorder. Biol Psychiatry. 83:1044–1053.

33. Rich-Edwards JW, Kaiser UB, Chen GL, Manson JE, Goldstein JM (2018): Sex and gender differences research design for basic, clinical, and population studies: Essentials for investigators. Endocr Rev. 39:424–439.

34. Major Depressive Disorder Working Group of the Psychiatric GWAS Consortium, Ripke S, Wray NR, Lewis CM, Hamilton SP, Weissman MM, et al. (2013): A mega-analysis of genome-wide association studies for major depressive disorder. Mol Psychiatry. 18:497–511.

35. Psychiatric Genomics Consortium Schizophrenia Working Group (2014): Biological insights from 108 schizophrenia-associated genetic loci. Nature. 511:421–427.

36. Psychiatric GWAS Consortium Bipolar Disorder Working Group (2011): Large-scale genome-wide association analysis of bipolar disorder identifies a new susceptibility locus near ODZ4. Nat Genet. 43:977–983.

37. Pedersen CB, Bybjerg-Grauholm J, Pedersen MG, Grove J, Agerbo E, Baekvad-Hansen M, et al. (2018): The iPSYCH2012 case-cohort sample: New directions for unravelling genetic and environmental architectures of severe mental disorders. Mol Psychiatry. 23:6–14.

38. Lam M, Awasthi S, Watson HJ, Goldstein J, Panagiotaropoulou G, Trubetskoy V, et al. (2020): RICOPILI: Rapid Imputation for COnsortias PIpeLIne. Bioinformatics. 36:930–933.

39. Chang CC, Chow CC, Tellier LC, Vattikuti S, Purcell SM, Lee JJ (2015): Second-generation PLINK: Rising to the challenge of larger and richer datasets. Gigascience. 4:7.

40. Willer CJ, Li Y, Abecasis GR (2010): METAL: Fast and efficient meta-analysis of genomewide association scans. Bioinformatics. 26:2190–2191.

41. Bulik-Sullivan BK, Loh PR, Finucane HK, Ripke S, Yang J, Schizophrenia Working Group of the Psychiatric Genomics Consortium, et al. (2015): LD Score regression distinguishes confounding from polygenicity in genome-wide association studies. Nat Genet. 47:291–295.

42. Zheng J, Erzurumluoglu AM, Elsworth BL, Kemp JP, Howe L, Haycock PC, et al. (2017): LD Hub: A centralized database and web interface to perform LD score regression that maximizes the potential of summary level GWAS data for SNP heritability and genetic correlation analysis. Bioinformatics. 33:272–279.

43. Keller MC (2014): Gene x environment interaction studies have not properly controlled for potential confounders: The problem and the (simple) solution. Biol Psychiatry. 75:18–24.

44. Bhattacharjee S, Rajaraman P, Jacobs KB, Wheeler WA, Melin BS, Hartge P, et al. (2012): A subset-based approach improves power and interpretation for the combined analysis of genetic association studies of heterogeneous traits. Am J Hum Genet. 90:821–835.

45. Benner C, Spencer CC, Havulinna AS, Salomaa V, Ripatti S, Pirinen M (2016): FINEMAP: Efficient variable selection using summary data from genome-wide association studies. Bioinformatics. 32:1493–1501.

46. Hormozdiari F, Kostem E, Kang EY, Pasaniuc B, Eskin E (2014): Identifying causal variants at loci with multiple signals of association. Genetics. 198:497–508.

47. de Leeuw CA, Mooij JM, Heskes T, Posthuma D (2015): MAGMA: Generalized gene-set analysis of GWAS data. PLoS computational biology. 11:e1004219.

48. Network & Pathway Analysis Subgroup of Psychiatric Genomics Consortium (2015): Psychiatric genome-wide association study analyses implicate neuronal, immune and histone pathways. Nat Neurosci. 18:199–209.

49. Pardiñas AF, Holmans P, Pocklington AJ, Escott-Price V, Ripke S, Carrera N, et al. (2018): Common schizophrenia alleles are enriched in mutation-intolerant genes and in regions under strong background selection. Nat Genet. 50:381–389.

50. Gorokhova S, Bibert S, Geering K, Heintz N (2007): A novel family of transmembrane proteins interacting with beta subunits of the Na,K-ATPase. Hum Mol Genet. 16:2394–2410.

51. Davies G, Lam M, Harris SE, Trampush JW, Luciano M, Hill WD, et al. (2018): Study of 300,486 individuals identifies 148 independent genetic loci influencing general cognitive function. Nat Commun. 9:2098.

52. Aberg KA, Liu Y, Bukszar J, McClay JL, Khachane AN, Andreassen OA, et al. (2013): A comprehensive family-based replication study of schizophrenia genes. JAMA Psychiatry. 70:573–581.

53. Edwards AC, Bigdeli TB, Docherty AR, Bacanu S, Lee D, de Candia TR, et al. (2016): Meta-analysis of positive and negative symptoms reveals schizophrenia modifier genes. Schizophr Bull. 42:279–287.

54. Peltola MA, Kuja-Panula J, Lauri SE, Taira T, Rauvala H (2011): AMIGO is an auxiliary subunit of the Kv2.1 potassium channel. EMBO Rep. 12:1293–1299.

55. Bishop HI, Cobb MM, Kirmiz M, Parajuli LK, Mandikian D, Philp AM, et al. (2018): Kv2 ion channels determine the expression and localization of the associated AMIGO-1 cell adhesion molecule in adult brain neurons. Front Mol Neurosci. 11:1.

56. He CW, Liao CP, Pan CL (2018): Wnt signalling in the development of axon, dendrites and synapses. Open Biol. 8:180116.

57. Bem J, Brozko N, Chakraborty C, Lipiec MA, Kozinski K, Nagalski A, et al. (2019): Wnt/beta-catenin signaling in brain development and mental disorders: Keeping TCF7L2 in mind. FEBS Lett. 593:1654–1674.

58. Hennig KM, Fass DM, Zhao WN, Sheridan SD, Fu T, Erdin S, et al. (2017): WNT/beta-Catenin pathway and epigenetic mechanisms regulate the Pitt-Hopkins syndrome and schizophrenia risk gene TCF4. Mol Neuropsychiatry. 3:53–71.

59. Hoseth EZ, Krull F, Dieset I, Morch RH, Hope S, Gardsjord ES, et al. (2018): Exploring the Wnt signaling pathway in schizophrenia and bipolar disorder. Transl Psychiatry. 8:55.

60. Yu Z, Lin D, Zhong Y, Luo B, Liu S, Fei E, et al. (2019): Transmembrane protein 108 involves in adult neurogenesis in the hippocampal dentate gyrus. Cell Biosci. 9:9.

61. O’Brien HE, Hannon E, Jeffries AR, Davies W, Hill MJ, Anney RJ, et al. (2018): Sex differences in gene expression in the human fetal brain. bioRxiv. doi: https://doi.org/10.1101/483636.

62. McClellan KM, Stratton MS, Tobet SA (2010): Roles for gamma-aminobutyric acid in the development of the paraventricular nucleus of the hypothalamus. J Comp Neurol. 518:2710–2728.

63. Sellgren CM, Gracias J, Jungholm O, Perlis RH, Engberg G, Schwieler L, et al. (2019): Peripheral and central levels of kynurenic acid in bipolar disorder subjects and healthy controls. Transl Psychiatry. 9:37.

64. Nilsson LK, Linderholm KR, Engberg G, Paulson L, Blennow K, Lindstrom LH, et al. (2005): Elevated levels of kynurenic acid in the cerebrospinal fluid of male patients with schizophrenia. Schizophr Res. 80:315–322.

65. Miller CL, Llenos IC, Dulay JR, Weis S (2006): Upregulation of the initiating step of the kynurenine pathway in postmortem anterior cingulate cortex from individuals with schizophrenia and bipolar disorder. Brain Res. 1073-1074:25–37.

66. Sathyasaikumar KV, Stachowski EK, Wonodi I, Roberts RC, Rassoulpour A, McMahon RP, et al. (2011): Impaired kynurenine pathway metabolism in the prefrontal cortex of individuals with schizophrenia. Schizophr Bull. 37:1147–1156.

67. Kindler J, Lim CK, Weickert CS, Boerrigter D, Galletly C, Liu D, et al. (2019): Dysregulation of kynurenine metabolism is related to proinflammatory cytokines, attention, and prefrontal cortex volume in schizophrenia. Mol Psychiatry.

68. Strasser B, Becker K, Fuchs D, Gostner JM (2017): Kynurenine pathway metabolism and immune activation: Peripheral measurements in psychiatric and co-morbid conditions. Neuropharmacology. 112:286–296.

69. Ortona E, Pierdominici M, Rider V (2019): Sex Hormones and Gender Differences in Immune Responses. Front Immunol, pp 186.

70. Hochgreb-Hägele T, Koo DE, Bronner ME (2015): Znf385C mediates a novel p53-dependent transcriptional switch to control timing of facial bone formation. Dev Biol. 400:23–32.

71. Buniello A, MacArthur JAL, Cerezo M, Harris LW, Hayhurst J, Malangone C, et al. (2019): The NHGRI-EBI GWAS Catalog of published genome-wide association studies, targeted arrays and summary statistics 2019. Nucleic Acids Res. 47:D1005–D1012.

72. Fishilevich S, Nudel R, Rappaport N, Hadar R, Plaschkes I, Iny Stein T, et al. (2017): GeneHancer: Genome-wide integration of enhancers and target genes in GeneCards. Database (Oxford). 2017.

73. Ziats MN, Rennert OM (2013): Sex-biased gene expression in the developing brain: Implications for autism spectrum disorders. Mol Autism. 4:10.

74. Seney ML, Huo Z, Cahill K, French L, Puralewski R, Zhang J, et al. (2018): Opposite molecular signatures of depression in men and women. Biol Psychiatry. 84:18–27.

75. Labonté B, Engmann O, Purushothaman I, Menard C, Wang J, Tan C, et al. (2017): Sex-specific transcriptional signatures in human depression. Nat Med. 23:1102–1111.

76. McCarthy MM (2019): Sex differences in neuroimmunity as an inherent risk factor. Neuropsychopharmacology. 44:38–44.

77. Wang K, Baldassano R, Zhang H, Qu HQ, Imielinski M, Kugathasan S, et al. (2010): Comparative genetic analysis of inflammatory bowel disease and type 1 diabetes implicates multiple loci with opposite effects. Hum Mol Genet. 19:2059–2067.

78. Byars SG, Stearns SC, Boomsma JJ (2014): Opposite risk patterns for autism and schizophrenia are associated with normal variation in birth size: Phenotypic support for hypothesized diametric gene-dosage effects. Proc Biol Sci. 281:20140604.

79. Crespi B, Badcock C (2008): Psychosis and autism as diametrical disorders of the social brain. Behav Brain Sci. 31:241–261; discussion 261-320.

80. Cross-Disorder Group of the Psychiatric Genomics Consortium (2019): Genomic relationships, novel loci, and pleiotropic mechanisms across eight psychiatric disorders. Cell. 179:1469–1482 e1411.

81. Goldstein JM, Hale T, Foster SL, Tobet SA, Handa RJ (2019): Sex differences in major depression and comorbidity of cardiometabolic disorders: impact of prenatal stress and immune exposures. Neuropsychopharmacology. 44:59–70.

82. Malamitsi-Puchner A, Tziotis J, Tsonou A, Protonotariou E, Sarandakou A, Creatsas G (2000): Changes in serum levels of vascular endothelial growth factor in males and females throughout life. J Soc Gynecol Investig. 7:309–312.

83. Mahoney ER, Dumitrescu L, Moore AM, Cambronero FE, De Jager PL, Koran MEI, et al. (2019): Brain expression of the vascular endothelial growth factor gene family in cognitive aging and alzheimer’s disease. Mol Psychiatry. doi: https://doi.org/10.1038/s41380-019-0458-5.

84. Frahm KA, Schow MJ, Tobet SA (2012): The vasculature within the paraventricular nucleus of the hypothalamus in mice varies as a function of development, subnuclear location, and GABA signaling. Horm Metab Res. 44:619–624.

85. Frahm KA, Handa RJ, Tobet SA (2018): Embryonic exposure to dexamethasone affects nonneuronal cells in the adult paraventricular nucleus of the hypothalamus. J Endocr Soc. 2:140–153.

86. Mayne BT, Bianco-Miotto T, Buckberry S, Breen J, Clifton V, Shoubridge C, et al. (2016): Large scale gene expression meta-analysis reveals tissue-specific, sex-biased gene expression in humans. Front Genet. 7:183.

87. Savage JE, Jansen PR, Stringer S, Watanabe K, Bryois J, de Leeuw CA, et al. (2018): Genome-wide association meta-analysis in 269,867 individuals identifies new genetic and functional links to intelligence. Nat Genet. 50:912–919.

88. Lee JJ, Wedow R, Okbay A, Kong E, Maghzian O, Zacher M, et al. (2018): Gene discovery and polygenic prediction from a genome-wide association study of educational attainment in 1.1 million individuals. Nat Genet. 50:1112–1121.

89. Howard DM, Adams MJ, Shirali M, Clarke TK, Marioni RE, Davies G, et al. (2018): Genome-wide association study of depression phenotypes in UK Biobank identifies variants in excitatory synaptic pathways. Nat Commun. 9:1470.

90. Tukiainen T, Villani AC, Yen A, Rivas MA, Marshall JL, Satija R, et al. (2017): Landscape of X chromosome inactivation across human tissues. Nature. 550:244–248.

91. Shi L, Zhang Z, Su B (2016): Sex biased gene expression profiling of human brains at major developmental stages. Sci Rep. 6:21181.

92. Lopes-Ramos CM, Chen CY, Kuijjer ML, Paulson JN, Sonawane AR, Fagny M, et al. (2020): Sex differences in gene expression and regulatory networks across 29 human tissues. Cell Rep. 31:107795.

93. Schizophrenia Working Group of the Psychiatric Genomics Consortium, Ripke S, Walters JTR, O’Donovan MC (2020): Mapping genomic loci prioritises genes and implicates synaptic biology in schizophrenia. medRxiv. doi: https://doi.org/10.1101/2020.09.12.20192922.

94. Pirastu N, Cordioli M, Nandakumar P, Mignogna G, Abdellaoui A, Hollis B, et al. (2020): Genetic analyses identify widespread sex-differential participation bias. bioRxiv. doi: https://doi.org/10.1101/2020.03.22.001453.

