## Supplementary Methods and Figures for "Sex-Dependent Shared and Non-Shared Genetic Architecture Across Mood and Psychotic Disorders"

SUPPLEMENTARY METHODS 4

POWER ANALYSES 4
IBD FILTERING 4
SEX INTERACTION VERSUS SEX STRATIFICATION 4
LINKAGE DISEQUILIBRIUM SCORE REGRESSION 4
SNP-BY-SEX INTERACTION ANALYSES 5
X CHROMOSOME 5
OMNIBUS TEST 6
IDENTIFICATION OF CREDIBLE SNPS 7
GENE-BASED TEST IN MAGMA 7
PATHWAY GENE SET ENRICHMENT ANALYSES 7
BRAIN EXPRESSION ANALYSIS 8
EXPRESSION QUANTITATIVE TRAIT LOCUS ANALYSES. 8
EVALUATION OF GXS INTERACTION FOR SEX-DEPENDENT AND CROSS-DISORDER SNPS FROM PRIOR STUDIES 9

SUPPLEMENTARY RESULTS 9

BRAIN EXPRESSION ANALYSIS 9
EVALUATION OF GXS INTERACTION FOR SEX-DEPENDENT AND CROSS-DISORDER SNPS FROM PRIOR STUDIES 9

SUPPLEMENTARY TABLES 11

SUPPLEMENTARY TABLE 1. PGC COHORT CHARACTERISTICS 11
SUPPLEMENTARY TABLE 2. IPSYCH COHORT CHARACTERISTICS 11
SUPPLEMENTARY TABLE 3. POWER ANALYSES 11
SUPPLEMENTARY TABLE 4. SNP-BASED HERITABILITY 11
SUPPLEMENTARY TABLE 5. SNP-BASED GENETIC CORRELATIONS 11
SUPPLEMENTARY TABLE 6. META-ANALYSIS AUTOSOMAL GXS INTERACTION LOCI IN PGC+IPSYCH 12
SUPPLEMENTARY TABLE 7. OMNIBUS TEST AUTOSOMAL GXS INTERACTION LOCI IN PGC+IPSYCH 12
SUPPLEMENTARY TABLE 8. META-ANALYSIS CHRX GXS INTERACTION LOCI IN PGC+IPSYCH 12
SUPPLEMENTARY TABLE 9. OMNIBUS TEST CHRX GXS INTERACTION LOCI IN PGC+IPSYCH 13
SUPPLEMENTARY TABLE 10. CREDIBLE SNPS FOR GXS LOCI IN PGC+IPSYCH 13
SUPPLEMENTARY TABLE 11. GENE-BASED TEST IN PGC+IPSYCH 13
SUPPLEMENTARY TABLE 12. MSIGDB PATHWAY GENE SET ENRICHMENT ANALYSES IN PGC+IPSYCH 14
SUPPLEMENTARY TABLE 13. SELECTED PATHWAY GENE SET ENRICHMENT ANALYSES IN PGC+IPSYCH 14

SUPPLEMENTARY TABLE 14. LOOKUP OF INTERACTION FOR SNPS SHOWING SEX-STRATIFICATION OR GXS INTERACTION IN 23ANDME, PGC, AND UK BIOBANK 14

SUPPLEMENTARY FIGURES 15

SUPPLEMENTARY FIGURE 1. EXPERIMENTAL DESIGN. 15
SUPPLEMENTARY FIGURE 2. POWER ANALYSES. 16
SUPPLEMENTARY FIGURE 3. MIAMI PLOTS FOR SEX-STRATIFIED ANALYSES IN PGC + IPSYCH 17
SUPPLEMENTARY FIGURE 4. SCATTER PLOTS OF FEMALE VS MALE ASSOCIATIONS IN PGC + IPSYCH 21
SUPPLEMENTARY FIGURE 5. QUANTILE-QUANTILE (Q-Q) PLOTS FOR GXS INTERACTION IN PGC + IPSYCH 24
SUPPLEMENTARY FIGURE 6. MANHATTAN PLOTS OF THE GXS INTERACTION GWAS IN PGC + IPSYCH 28
SUPPLEMENTARY FIGURE 7. LOCUSZOOM PLOTS FOR LOCI WITH GXS INTERACTION IN PGC + IPSYCH 33
SUPPLEMENTARY FIGURE 8. FOREST PLOTS FOR PGC + IPSYCH 46
SUPPLEMENTARY FIGURE 9. X CHROMOSOME MODEL COMPARISONS IN PGC + IPSYCH 63
SUPPLEMENTARY FIGURE 10. MANHATTAN PLOTS FOR GENE-BASED GXS TESTS IN PGC + IPSYCH 66
SUPPLEMENTARY FIGURE 11. GTEX MULTI-TISSUE EXPRESSION FOR GXS LOCI 69
SUPPLEMENTARY FIGURE 12. ALLEN BRAIN ATLAS EXPRESSION ACROSS DEVELOPMENT FOR GXS LOCI 71
SUPPLEMENTARY FIGURE 13. LIFE BRAIN EXPRESSION COURSE DERIVED FROM THE HUMAN BRAIN TRANSCRIPTOME (HBT) PROJECT FOR GXS LOCI 74
SUPPLEMENTARY FIGURE 14. GTEX SEX-SPECIFIC MULTI-TISSUE EXPRESSION FOR GXS LOCI 77
SUPPLEMENTARY FIGURE 15. CELL TYPE-SPECIFIC BRAIN EXPRESSION DERIVED FROM THE STANFORD RNA-SEQ DATABASE FOR GXS LOCI 80

SUPPLEMENTARY TABLES PGC ONLY 82

SUPPLEMENTARY TABLE 15. META-ANALYSIS AUTOSOMAL GXS INTERACTION LOCI IN PGC 82
SUPPLEMENTARY TABLE 16. OMNIBUS TEST AUTOSOMAL GXS INTERACTION LOCI IN PGC 82
SUPPLEMENTARY TABLE 17. META-ANALYSIS CHRX GXS INTERACTION LOCI IN PGC 82
SUPPLEMENTARY TABLE 18. OMNIBUS TEST CHRX GXS INTERACTION LOCI IN PGC 82
SUPPLEMENTARY TABLE 19. CREDIBLE SNPS FOR GXS LOCI IN PGC 83
SUPPLEMENTARY TABLE 20. GENE-BASED TEST IN PGC 83
SUPPLEMENTARY TABLE 21. MSIGDB PATHWAY GENE SET ENRICHMENT ANALYSES IN PGC 83
SUPPLEMENTARY TABLE 22. SELECTED PATHWAY GENE SET ENRICHMENT ANALYSES IN PGC 84

SUPPLEMENTARY FIGURES PGC ONLY 85

SUPPLEMENTARY FIGURE 16. LD SCORE REGRESSION ESTIMATES OF SNP-BASED (A) HERITABILITY AND (B) GENETIC CORRELATIONS (SE) IN PGC ONLY 85
SUPPLEMENTARY FIGURE 17. MIAMI PLOTS FOR SEX-STRATIFIED ANALYSES IN PGC 86
SUPPLEMENTARY FIGURE 18. SCATTER PLOTS OF FEMALE VS MALE ASSOCIATIONS IN PGC 90
SUPPLEMENTARY FIGURE 19. QUANTILE-QUANTILE (Q-Q) PLOTS FOR GXS INTERACTION IN PGC 93
SUPPLEMENTARY FIGURE 20. MANHATTAN PLOTS OF THE GXS INTERACTION GWAS IN PGC 97
SUPPLEMENTARY FIGURE 21. LOCUSZOOM PLOTS FOR LOCI WITH GXS INTERACTION IN PGC 102
SUPPLEMENTARY FIGURE 22. FOREST PLOTS FOR PGC 120
SUPPLEMENTARY FIGURE 23. X CHROMOSOME MODEL COMPARISONS IN PGC 136
SUPPLEMENTARY FIGURE 24. MANHATTAN PLOTS FOR GENE-BASED GXS TESTS IN PGC 139

**ACKNOWLEDGEMENTS 142**

**CONSORTIUM MEMBERSHIP 144**

**SCHIZOPHRENIA WORKING GROUP OF THE PSYCHIATRIC GENOMICS CONSORTIUM 144**
WELLCOME TRUST CASE-CONTROL CONSORTIUM 2 145
PSYCHOSIS ENDOPHENOTYPE INTERNATIONAL CONSORTIUM 146
**BIPOLAR DISORDER WORKING GROUP OF THE PSYCHIATRIC GENOMICS CONSORTIUM 154**
**MAJOR DEPRESSIVE DISORDER WORKING GROUP OF THE PSYCHIATRIC GENOMICS CONSORTIUM 162**
**SEX DIFFERENCES CROSS-DISORDER ANALYSIS GROUP OF THE PSYCHIATRIC GENOMICS CONSORTIUM 168**
**IPSYCH 171**

**SUPPLEMENTARY REFERENCES 172**

### Supplementary Methods

#### Power analyses

Power analyses were carried out using the 'GeneticsDesign' Bioconductor package in R. At the listed within-disorder and cross-disorder sample sizes, and a MAF of 0.05, this study had 83%-99% power to detect disease risk interaction effects within-disorder at an odds ratio of >= 1.2, and 88% power to detect effects cross-disorder at an odds ratio of >= 1.1. Power estimates for varying effect sizes and for the different data configurations are presented in **Supplementary Table 3** and **Supplementary Figure 2**.

#### IBD Filtering

IBD analyses were performed using PLINK (1) to identify duplicate samples and/or cryptic relatedness. For sample pairs in which PI_HAT > 0.1, one sample was randomly excluded if both samples were cases or both controls, or the control sample was excluded if one sample was a case and the other a control. IBD filtering was first applied within each study cohort and subsequently across study cohorts within a disorder. For cross-disorder analyses, IBD filtering was applied across all cohorts across all three disorders.

#### Sex interaction versus sex stratification

An interaction test is powered to detect a difference between the sexes in genetic risk and needed to determine whether differences in effect sizes are statistically different between the sexes (2). On the other hand, a stratified analysis is required in order to characterize the effect size itself, and the direction of effect within each sex. A sex-stratified analysis followed by a Z-score difference test (Eq. 1) is equivalent to a formal test for GxS interaction when there is no interaction between covariates and the strata, and the trait variance are equivalent in the two strata. Thus, different information can be gained from both types of analyses.

Eq. 1: $$[Z - score = \frac{Beta_{female} - Beta_{male}}{\sqrt{SE^2_{female} + SE^2_{male}}}]$$

Here, we focused on the interaction analysis. However, to characterize the quality of the suggestive GxS interaction signals ($p<1\times10^{-6}$), included in the tables with GxS interaction results (**Tables 1-2, Supplementary Tables 6-9**), we also report sex-specific association statistics. Miami plots show the genome-wide sex-specific associations (**Supplementary Figure 3**). None of these GxS SNPs showed a genome-wide significant sex-specific signal. Furthermore, a scatter plot of -log10(*p*-values) for sex-specific genome-wide associations indicated little overlap in the top signals across sexes (**Supplementary Figure 4**).

#### Linkage Disequilibrium Score Regression

Estimates of $h^2_{SNP}$ were transformed to the liability scale assuming lifetime prevalence of the disorder in the population ($K$) of $K$=0.01, $K$=0.01, and $K$=0.10 for SZ, BD, and MDD females, respectively, and $K$=0.01, $K$=0.01, and $K$=0.05 for SZ, BD, and MDD males, respectively, based on a Danish population study (3). Estimates of $h^2_{SNP}$ increased minimally across a range of MAF cutoffs (MAF>1%, 2%, 5%), indicating rarer variants contributed little to the heritability estimates (**Supplementary Table 7**).

For traits with non-zero $h^2_{SNP}$ estimates in both sexes (significantly greater than zero; z=$h^2_{SNP}$/SE), we tested whether the estimates were significantly different between the sexes by calculating Z-scores using the Equation above (replacing Beta with $h^2_{SNP}$), and obtaining corresponding p-values from the standard normal distribution, followed by Bonferroni-correction for multiple testing based on 3 independent tests/disorders (p-value threshold = 0.017).

We also used LDSC (4) to estimate bivariate genetic correlations ($r_g$) between the sexes and between the three disorders. LDSC genetic correlations attributable to genome-wide SNPs ($r_g$) were estimated within (males/females) and across disorders (sex-interaction; males/females). The intent of these comparisons was to evaluate the extent of shared common variant genetic architectures in order to suggest hypotheses about the fundamental genetic basis of sex differences in these three disorders. These $r_g$ are mostly based on studies of independent subjects and the estimates should be unbiased by confounding of genetic and non-genetic effects (except if there is genotype by environment correlation). When GWAS studies include overlapping samples, $r_g$ estimates remain unbiased but the intercept of the LDSC regression increases as it is an estimate of the correlation between association statistics attributable to sample overlap. Subject overlap in itself does not bias $r_g$ (4, 5). Therefore, we used the data with only within-cohort/within-disorder IBD filtering for these analyses.

For between-sex, within-disorder correlations, we used one-tailed tests comparing to a standard normal distribution, to determine whether $r_g$ was significantly greater than zero (z=$r_g$/SE) and significantly less than 1 (z=$(1 - r_g)$/SE). Bonferroni-correction was applied to adjust for multiple testing based on 3 tests/disorders. Next, we determined whether the between-trait, within-sex correlations were different for males and females (see Equation above). Given the non-independent genetic correlations across disorders, rendering Bonferroni-correction overly conservative, we applied false discovery rate (FDR) correction to adjust for multiple tests.

All estimates of $h^2_{SNP}$ and $r_g$ are based on the autosomal contributions only, as LDSC currently does not allow for estimation of $h^2_{SNP}$ and $r_g$ from the X chromosome, due to its more complex composition.

We provided the within-disorder meta-analysis sex-stratified summary statistics, calculated based on the PGC-only samples, to Martin et al (6), who evaluated sex differences in heritability estimates and genetic correlations of multiple psychiatric disorders and relevant quantitative phenotypes in an expanded set of analyses.

**SNP-by-sex interaction analyses**

PLINK (1) was used to perform a genome-wide genotype–by–sex (GxS) interaction analysis of each study cohort, followed by standard-error weighted (i.e., inverse variance) meta-analysis of the GxS interaction results using METAL (7). GxS interaction analyses were performed using logistic regression with a main effect for each SNP, a main effect for sex, and SNP-by-sex interaction terms, using an additive model for each SNP. The first 10 ancestry principal components (PC) were included as covariates to adjust for population stratification. A secondary regression model included additional statistical controls in the form of 10 SNP-by-PC interaction terms and 10 sex-by-PC interaction terms in addition to the terms above (8). The "dosage" information score for imputed genotype was used to account for uncertainty of imputation. For all analyses, SNPs with poor imputation quality (IMPUTE2 INFO score < 0.6) or low minor allele frequency (MAF <0.01 for SNP-by-sex interaction analysis) were excluded.

**X chromosome**

The X chromosome is usually excluded from GWAS because the data has a different, sex-specific structure and, therefore, requires special analytical tools.(9) While there are two copies of each autosomal chromosome, males carry only one copy of the X chromosome whereas females, again, carry two copies. Therefore, at each SNP, females can carry one of three possible genotypes; that is, they can have 0, 1 or 2 copies of a specific allele. In contrast to this, there are only two possible genotypes for males, corresponding to 0 or 1 copies of a specific allele. Only for the so-called pseudo-autosomal regions, there exist homologous loci on the Y chromosome, and males can have up to 2 copies of a specific allele. In addition, one of the two female X chromosomes might be inactivated. In each cell, one of the two female X chromosomes is randomly selected to be silenced (10). This means that the expression levels of this chromosome are much lower than for the second chromosome in the cell. This mechanism of dose compensation should result in comparable expression levels for males and females despite the different number of chromosome copies. However, this inactivation is incomplete: while some genes or regions will be completely inactivated, some genes might show expression levels that are reduced only slightly or not at all. Therefore, to analyze X-chromosomal data, special quality control (separately for males and females) and test statistics are required (11). The choice of the best statistical test depends on the underlying genetic model and the inactivation patterns at a specific locus (12).

**Omnibus test**

As opposite risk effects of SNPs in cross-disorder analyses (i.e., a particular allele is associated with increased risk of one disorder and decreased risk of another disorder) might cancel each other out, we also performed a three-degrees-of-freedom (df) omnibus test (13-15) as a second analytical approach. This test was performed by summing the $\chi^2$ values for each individual disease meta-analysis, which enables detection of opposing allelic effects across disorders.

Association analysis based on SubSETs (ASSET) (16) is designed to be powerful for pooling association signals across multiple studies when true effects may exist only in a subset of the studies and could be in opposite directions across studies. The method explores all possible subsets of studies and evaluates fixed-effect meta-analysis test statistics for each subset. The final test statistic is obtained by maximizing the subset-specific test statistics over all possible subsets and then evaluating its significance after efficient adjustment for multiple testing, taking into account the correlation between test statistics across different subsets due to possible subject overlap (although here we removed this overlap using the IBD filtering described above). The method not only returns a *p*-value for significance for the overall evidence of association of a SNP across studies, but also outputs the "best subset" containing the studies that contributed to the overall association signal. For detection of association signals with effects in opposite directions, ASSET allows subset search separately for positively- and negatively- associated studies and then combines association signals from two directions using a chi-square test statistic.

Inclusion of East Asian ancestry SCZ cohorts, which represent a relatively small component of the SCZ dataset (7.56% of PGC; 7.03% of PGC+iPSYCH), did not substantially improve SNP-by-sex interaction results. For this reason, and given that the gene- and pathway-based analyses reported below required the application of an ancestry-specific reference panel, all subsequent analyses utilized only European ancestry cohorts.

**Identification of credible SNPs**

Linkage disequilibrium (LD)-independent SNPs with genome-wide significance ($p < 5\times10^{-8}$) and suggestive GxS signals ($p < 1\times10^{-6}$) were used as index SNPs to obtain credible SNPs (i.e., potentially causal in disease risk). All SNPs associated with $p < 1\times10^{-6}$ and SNPs in LD ($r^2 >$ 0.6) with the index SNP were selected. Correlations (LD structure) among this set of SNPs were calculated based on the 1000 Genomes Phase 1 European (CEU) reference panel. FINEMAP v1.4 (17) and CAVIAR v2.2 (18) (-r 0.95, posterior probability; -c 2, maximum number of causal SNPs) were applied to summary association statistics and LD structure for each index SNP locus (plink --bfile 1kgp1_ref_file --clump metal_output_file --clump-p1 1e-4 --clump-p2 0.05 --clump-r2 0.1 --clump-kb 250), and credible SNPs for each index SNP were identified. We summarize the posterior probabilities of all SNPs in the fine-mapping loci (**Supplementary Table 10**) and highlight the SNPs that are most likely to have a causal effect on mood and psychotic disorders. It is noteworthy that the SNPs with the highest posterior probability of causality are not necessarily the most statistically significant SNPs in the original GxS analysis.

**Gene-based test in MAGMA**

Briefly, the gene-based test evaluates whether the number of associated SNPs in/around a particular gene is greater than would be expected given the size and structure of that gene, as opposed to a SNP-based test, which does not take into account gene size and structure. The gene-based test in MAGMA (19) is based on a multiple linear principal components regression model, using an F-test to compute the gene *p*-value. This model first projects the SNP matrix for a gene onto its principal components (PC), pruning away PCs with very small eigenvalues, and then uses those PCs as predictors for the phenotype in the linear regression model. This improves power by removing redundant parameters, and guarantees that the model is identifiable in the presence of highly collinear SNPs. Although application of the linear regression model to a binary phenotype violates some assumptions of the F-test, comparison of the F-test *p*-values with *p*-values based on permutation of the F-statistic has shown that the F-test remains accurate.

We applied an adjusted genome-wide significant *p*-value threshold of $p < 2.6\times10^{-6}$, which accounts for 19,427 autosome and sex chromosome genes evaluated in the test (**Supplementary Table 11**).

**Pathway gene set enrichment analyses**

Using MAGMA (19), two sets of pathway/gene set enrichment analyses were carried out. Hypothesis-free analyses were performed for Gene Ontology (GO) pathways (20, 21) plus curated gene sets (including gene sets from BioCarta, KEGG, and Reactome) from the Molecular Signatures Database v6.2 (MSigDB; http://software.broadinstitute.org/gsea/msigdb/genesets.jsp) (22). Pathways analyzed contained a minimum of 10 genes because statistics for smaller gene sets tend to be over-dispersed (23), reducing down the number of MSigDB gene sets from 5917 GO + 4762 curated = 10,679 pathways to 10,353. Data-driven analyses included an additional 9 gene sets/pathways compiled from prior studies: three gene sets reported to be enriched across the PGC SZ, BD, and MDD cohorts (23), immune/neurotrophic, synaptic, and histone methylation, and six central nervous system (CNS) pathways that were enriched in the largest SZ GWAS to date (CLOZUK+PGC) (24): mouse phenome (MP) abnormal behavior, MP abnormal long-term potentiation, MP abnormal CNS electrophysiology, 5HT2C receptor complex, FMRP targets, and Voltage-gated $Ca^{2+}$ channels. The number of genes in each (top) pathway are listed in **Supplementary Tables 12-13**.

Ensembl gene definitions were used as the reference gene annotation and map. The different pathway sets were combined into one database and identical pathways merged. SNPs were assigned to genes based on human genome build 37 positions if they lay within 10 kb upstream or 10 kb downstream of the gene, to capture transcriptional regulatory elements. SNPs that mapped to more than one gene, were assigned to all such genes. Analyses were run according to the standard protocols for MAGMA, both with and without the MHC region (chromosome 6, between base pair position 25,000,000 and 35,000,000). MAGMA (19) is a "best SNP per gene" method that counts the number of genes in a pathway where a number of independent SNPs exceed a predefined significance, and adjusts for LD and genomic structure with corrected statistics derived by Monte Carlo simulation. This gene-set analysis also uses a regression structure to allow generalization to analysis of continuous properties of genes and simultaneous analysis of multiple gene sets and other gene properties. To determine whether any pathway gene sets annotate the top GWAS genes at a frequency greater than that would be expected by chance, a *p*-value was calculated using the hypergeometric distribution (25). Correction for multiple testing was applied by estimating the FDR (26).

**Brain expression analysis**

Brain expression (RNA Sequencing, RNA-Seq) data from the Genotype-Tissue Expression project (GTEx; 44 tissues, N>70; http://www.gtexportal.org (27, 28)), the Human Brain Transcriptome project (HBT; http://hbatlas.org (29, 30)), the Allen Brain Atlas (http://human.brain-map.org (31)), and the Stanford Brain RNA-Seq database (http://web.stanford.edu/group/barres_lab/brain_rnaseq.html (32, 33) were evaluated to validate and interpret the GxS interaction results (variants with a GxS interaction $p$-value $< 1\times10^{-6}$).

The expression levels from the Allen Brain Atlas were averaged across the 6 brain tissue samples and up to 6 probes per gene. As the experiments contained in the Stanford Brain RNA-Seq database (32, 33), i.e. expression of genes in neurons, astrocytes, and oligodendrocytes specifically, were done in mice, genes were mapped to human orthologous genes using Ensembl.

**Expression quantitative trait locus analyses.**

All variants with a GxS interaction $p$-value $< 1\times10^{-6}$ were analyzed further to test whether their genotype was associated with RNA(-Seq) expression. The most significant SNP from each locus having $p < 1\times10^{-6}$ in GxS interaction analyses was assessed for the possibility of genotype-specific gene expression patterns (or expression quantitative trait loci, eQTLs). To assess variants for their influence on expression of their closest genes in brain tissue, we conducted eQTL look-ups of the most associated SNPs in each locus ($p < 1\times10^{-6}$) and report GWAS SNPs in LD ($r^2 > 0.8$) with the top eQTLs in the following data sets: GTEx (27, 28), PsychENCODE (PEC; PFC, N=427; https://www.synapse.org/pec ; http://resource.psychencode.org) (34, 35), CommonMind Consortium (CMC; dorsolateral prefrontal cortex [DLPFC], Sage Synapse accession syn5650509, N=467; https://www.synapse.org/cmc) (36), the Lieber Institute for Brain Development (LIBD; DLPFC), accessed via the eQTL Browser (http://eqtl.brainseq.org/) (37, 38). For SNPs showing significant eQTLs in the GTEx dataset, we looked for replication in the other datasets. Expression QTLs that reached a threshold of $\alpha = 0.05$ in the GTEx dataset and replicated (defined as a threshold of $\alpha = 0.05$ in the same direction) in PEC, CMC, and/or LIBD are reported.

**Evaluation of GxS interaction for sex-dependent and cross-disorder SNPs from prior studies**

To assess overlap of GxS signals between the current study and prior published studies, GxS interaction results were compared to previously reported sex-dependent or sex-specific effects on psychiatric illness risk ($p<5\times10^{-8}$) from sex-stratified analyses by the PGC (2, 39, 40), ASD collection (41), 23andMe (42), and UK Biobank (43) (see Supplementary Methods). Additionally, GxS interaction effects were evaluated for SNPs with genome-wide significant main additive effects across sexes in the recent PGC cross-disorder group (CDG) study of eight disorders (includes the PGC SCZ, BIP, and MDD datasets analyzed in this study) (44).

For the UK Biobank GWAS, genome-wide sex-stratified summary statistics are available for download for a range of mental illness diagnoses. Lookups were performed for SNPs with a significant Z difference score ($p < 5\times10^{-8}$) between the sexes only. The Z difference score was calculated as described above. Additionally, GxS interaction effects were evaluated for genome-wide significant SNPs (main additive effect across sexes) from the recent PGC cross-disorder group (CDG) study of eight disorders (this study includes the PGC data analyzed here) (44).

### Supplementary Results

**Brain expression analysis**

Tissue and brain expression data were examined for genes located adjacent to SNPs with significant or suggestive evidence for GxS interactions ($p<1\times10^{-6}$; i.e. the SNPs listed in **Tables 1-2)**. As shown in **Supplementary Figures 11-13**, the NKAIN2 gene containing the omnibus genome-wide significant SNP (rs117780815) is specifically expressed in brain in the adult, being highest in spinal cord followed by hippocampus and substantia nigra, while expression during neurodevelopment is highest in prenatal cortex and neocortex. MOCOS expression is highest in tibial nerve in adulthood, and prenatally in cortex, hippocampus, and amygdala. IDO2 expression in adulthood is highest in cortex, and in childhood frontal cortex and amygdala. SLTM expression is highest in adulthood in hypothalamus, and in prenatal and childhood cortex. TUSC1 is fairly consistently expressed across the brain and across development from prenatal development through childhood to adulthood. FHL2 brain expression is relatively low prenatally, highest in mediodorsal nucleus of the thalamus in early childhood, and in neocortex in adulthood. SPAG17 expression is highest prenatally in hippocampus and amygdala, through childhood in hippocampus, and in the adult hypothalamus. ZNF385C expression is highest in the cerebellum (including cortex), throughout prenatal development, childhood, and adulthood. Among seven brain cell types, NKAIN2 expression is highest in oligodendrocytes, MOCOS in endothelial cells and microglia, IDO2 and FHL2 in oligodendrocyte precursor cells, SLTM, SPAG17 and ZNF385C in astrocytes, and TUSC1 in neuron (**Supplementary Figure 15**).

Evaluation of sex-specific expression detected different expression levels between males and females of several of the genes in some brain regions (**Supplementary Figure 21**).

**Evaluation of GxS interaction for sex-dependent and cross-disorder SNPs from prior studies**

Of four SNPs with nominally significant SNP-by-sex interactions ($p < 0.05$) identified in a 23andMe study of MDD (42), two SNPs exhibited nominally significant GxS interactions in our analyses (**Supplementary Table 14)** of MDD (rs2042772; $p$=0.037) and BIP (rs4543289; $p$=0.034). SNPs with significant sex-dependent effects ($p<5\times10^{-8}$) in prior within-disorder studies of ADHD, OCD, PTSD, and ASD (2, 39-41) or UK Biobank psychiatric phenotypes (43) had non-significant ($p_{FDR} > 0.05$) GxS interaction $p$-values in this study. Among the genome-wide significant results in a PGC cross-disorder (non-sex-stratified) analysis of 8 psychiatric disorders (44), rs7521492 had a *p*-value of $4.2\times10^{-4}$ in our GxS omnibus test of SCZ, BIP and rMDD ($p_{FDR}$ = 0.034); rs11688767 had a *p*-value of $3.2\times10^{-4}$ in our meta-analysis of rMDD ($p_{FDR}$ = 0.034). Of note, *CSMD1*, identified in the PGC-CDG cross-disorder analysis (44), was among our top cross-disorder GxS results (regular meta-analysis). However, the most significant SNP in each analysis differed.

### Supplementary Tables

#### Supplementary Table 1. PGC cohort characteristics

See SupplTable1_PGC_cohort_characteristics.xlsx

The cohorts have been previously described in references (45-47).
Note: Due to the nature of the sample composition, 3 SCZ trio cohorts and 2 BIP trio cohorts were excluded from analyses (and from this table).
*No X chromosome data; #Recurrent MDD data available.

#### Supplementary Table 2. iPSYCH cohort characteristics

See SupplTable2_iPSYCH_cohort_characteristics.xlsx

The cohort has been previously described in Pedersen et al (2018) (48). For this study, the large control dataset was semi-randomly split into subsets to match with the patients for each disorder. The number of controls in each set was decided upon based on the percentage of patients with that disorder.

#### Supplementary Table 3. Power analyses

See SupplTable3_Power.xlsx

Power analyses for varying effect sizes and different data configurations were carried out using the 'GeneticsDesign' Bioconductor package in R. At the listed within-disorder and cross-disorder sample sizes, and a MAF of 0.05, this study had 83%-99% power to detect disease risk interaction effects within-disorder at an odds ratio of >= 1.2, and 88% power to detect effects cross-disorder at an odds ratio of >= 1.1.

#### Supplementary Table 4. SNP-based heritability

See SupplTable4_SNP-based_heritability_LDSC.xlsx

Estimates of SNP-based heritability, $h^2$ (standard error, SE), were obtained for three minor allele frequency (MAF) cutoffs using LD Score Regression (LDSC) with population prevalences of $K$=0.0124, $K$=0.0107, $K$=0.1018, and $K$=0.0563 for SCZ, BIP, MDD, and rMDD females, respectively, and $K$=0.0173, $K$=0.0076, $K$=0.0563, and $K$=0.0256 for SCZ, BIP, MDD, and rMDD males, respectively (3), to transform from the observed heritability scale to the liability scale. Primary model refers to the regression model without additional interaction covariates; secondary model refers to the extended model with additional interaction covariates.
Abbreviations: BIP = bipolar disorder; (r)MDD = (recurrent) major depressive disorder; SCZ = schizophrenia.

#### Supplementary Table 5. SNP-based genetic correlations

See SupplTable5_SNP-based_rg_LDSC.xlsx

Estimates of SNP-based genetic correlations, $r_g$ (standard error, SE), were obtained using LD Score Regression (LDSC) with MAF threshold 0.01 and population prevalences of $K$=0.0124, $K$=0.0107, $K$=0.1018, and $K$=0.0563 for SCZ, BIP, MDD, and rMDD females, respectively, and $K$=0.0173, $K$=0.0076, $K$=0.0563, and $K$=0.0256 for SCZ, BIP, MDD, and rMDD males, respectively (3). Primary model refers to the regression model without additional interaction covariates; secondary model refers to the extended model with additional interaction covariates.

Abbreviations: BIP = bipolar disorder; (r)MDD = (recurrent) major depressive disorder; SCZ = schizophrenia.

**Supplementary Table 6. Meta-analysis Autosomal GxS interaction loci in PGC+iPSYCH**

See SupplTable6_MetaAnalysisSTDERR_auto_PGC+iPSYCH.xlsx

Cross-disorder and within-disorder meta-analyses were carried out using METAL, incorporating cohort-level summary statistics from PLINK. Listed are LD-independent SNPs with interaction $p$-values < $1{\times}10^{-6}$ in SCZ, BIP, (r)MDD, and cross-disorder. Loci were clumped using '*plink --bfile 1kgp_ref_file --clump metal_output --clump-p1 1e-4 --clump-p2 1e-4 --clump-r2 0.6 --clump-kb 3000*'. Primary model refers to the regression model without additional interaction covariates; secondary model refers to the extended model with additional interaction covariates.

Abbreviations: BIP = bipolar disorder; (r)MDD = (recurrent) major depressive disorder; SCZ = schizophrenia.

**Supplementary Table 7. Omnibus test Autosomal GxS interaction loci in PGC+iPSYCH**

See SupplTable7_OmnibusTestASSET_auto_PGC+iPSYCH.xlsx

Omnibus tests were carried out using ASSET, incorporating the within-disorder meta-analysis summary statistics from METAL. Listed are LD-independent SNPs with cross-disorder interaction $p$-values < $1{\times}10^{-6}$. Loci were clumped using '*plink --bfile 1kgp_ref_file --clump asset_output --clump-p1 1e-4 --clump-p2 1e-4 --clump-r2 0.6 --clump-kb 3000*'. Primary model refers to the regression model without additional interaction covariates; secondary model refers to the extended model with additional interaction covariates.

Abbreviations: BIP = bipolar disorder; (r)MDD = (recurrent) major depressive disorder; SCZ = schizophrenia.

**Supplementary Table 8. Meta-analysis chrX GxS interaction loci in PGC+iPSYCH**

See SupplTable8_MetaAnalysisSTDERR_xchr_PGC+iPSYCH.xlsx

Cross-disorder and within-disorder meta-analyses were carried out using METAL, incorporating cohort-level summary statistics from PLINK. Listed are LD-independent SNPs with interaction $p$-values < $1{\times}10^{-6}$ in SCZ, BIP, (r)MDD, and cross-disorder. Model A **(a)** effectively assumes complete and uniform X-inactivation in females and a similar effect size between males and females. Females are considered to have 0, 1, or 2 copies of an allele; males are considered to have 0 or 2 copies of the same allele. Model B **(b)** considers the allelic dosages for females to be 0,1, or 2 copies, and males to be 0 or 1 copy as in an autosomal analysis. Loci were clumped using '*plink --bfile 1kgp_ref_file --clump metal_output --clump-p1 1e-4 --clump-p2 1e-4 --clump-r2 0.6 --clump-kb 3000*'. Primary model refers to the regression model without additional interaction covariates; secondary model refers to the extended model with additional interaction covariates.

Abbreviations: BIP = bipolar disorder; (r)MDD = (recurrent) major depressive disorder; SCZ = schizophrenia.

**Supplementary Table 9. Omnibus test chrX GxS interaction loci in PGC+iPSYCH**

See SupplTable9_OmnibusTestASSET_xchr_PGC+iPSYCH.xlsx

Omnibus tests were carried out using ASSET, incorporating the within-disorder meta-analysis summary statistics from METAL. Listed are LD-independent SNPs with cross-disorder interaction *p*-values < $1\times10^{-6}$. Loci were clumped using '*plink --bfile 1kgp_ref_file --clump asset_output --clump-p1 1e-4 --clump-p2 1e-4 --clump-r2 0.6 --clump-kb 3000*'. Omnibus tests were carried out using ASSET, incorporating the within-disorder meta-analysis summary statistics from METAL. Listed are LD-independent SNPs with cross-disorder interaction *p*-values < $1\times10^{-6}$.

Abbreviations: BIP = bipolar disorder; (r)MDD = (recurrent) major depressive disorder; SCZ = schizophrenia.

**Supplementary Table 10. Credible SNPs for GxS loci in PGC+iPSYCH**

See SupplTable10_CredibleSNPs_FineMapping_PGC+iPSYCH.xlsx

Fine mapping was carried out using both FINEMAP and CAVIAR. Fine mapping using FINEMAP was carried out with settings: *--sss --corr-config 0.95 --n-causal-snps 5 --n-configs-top 50000 --prior-k0 0 --prior-std 0.05*. If there were less than 5 SNPs in the locus, *--n-causal-snps* was set to the number of SNPs in the locus according to LD. The most likely causal SNPs per locus are highlighted in bold font. The shotgun stochastic search (--sss) conducts a pre-defined number of iterations within the space of causal configurations. In each iteration, the neighborhood of the current causal configuration is defined by configurations that result from deleting, changing or adding a causal SNP from the current configuration. The next iteration starts by sampling a new causal configuration from the neighborhood based on the scores normalized within the neighborhood. Fine mapping using CAVIAR was carried out with settings: *-r 0.95 -c 5 -f 1*. If there were less than 5 SNPs in the locus, *-c* was set to the number of SNPs in the locus according to LD. Analyses used European ancestry only summary statistics. Loci with $p < 1\times10^{-6}$ were analyzed (index SNPs determined based on clumping using LD threshold 0.1). The most likely causal SNPs per locus are highlighted in bold font. Primary model refers to the regression model without additional interaction covariates; secondary model refers to the extended model with additional interaction covariates.

Abbreviations: PP_group = posterior probability that there is at least one causal signal among SNPs in the same group with this SNP; PP_causal = posterior probability that the SNP is causal; BP = base pair position; BIP = bipolar disorder; CHR = chromosome; (r)MDD = (recurrent) major depressive disorder; SCZ = schizophrenia; SNP = Single Nucleotide Polymorphism rs ID.

**Supplementary Table 11. Gene-based test in PGC+iPSYCH**

See SupplTable11_Gene-BasedTest_PGC+iPSYCH.xlsx

Gene-based analyses were carried out in MAGMA on the genomic control output with INFO score > 0.6, European ancestry only, and autosomal SNPs only, with the MHC region included. Genes with *p*-values < $1\times10^{-4}$ are shown. There was no difference in the *p*-values when the MHC region was excluded. There were minor differences in *p*-values when using INFO score > 0.8, but with the same top 10 genes. *Significant at genome-wide threshold for gene-based test of 0.05 / 19,427 genes = $2.6\times10^{-6}$. Primary model refers to the regression model without additional interaction covariates; secondary model refers to the extended model with additional interaction covariates.

Abbreviations: BP = base pair position; Chr = chromosome; N SNPs = number of SNPs in gene; N Param = number of parameters; N = sample size; Z = Z-statistic; BIP = bipolar disorder; MDD = major depressive disorder; rMDD = recurrent major depressive disorder; SCZ = schizophrenia.

**Supplementary Table 12. MSigDB pathway gene set enrichment analyses in PGC+iPSYCH**

See SupplTable12_MSigDB pathway GSEA_PGC+iPSYCH.xlsx

Enrichment analyses were carried out in MAGMA on the genomic control output with INFO score > 0.6, European ancestry only, and autosomal SNPs only. Analyses were run both with (top subtable) and without (bottom subtable) inclusion of the Chromosome 6 Major Histocompatibility Complex (MHC) region. Each (sub)table displays the top 10 gene sets based on the uncorrected *p*-value. Hyperlinks link to the GSEA/MSigDB website with a description of the pathway. Primary model refers to the regression model without additional interaction covariates; secondary model refers to the extended model with additional interaction covariates.

Abbreviations: BIP = bipolar disorder; MDD = major depressive disorder; $P_{BONF}$ = Bonferroni-corrected *p*-value; $P_{FDR}$ = False Discovery Rate-corrected *p*-value; rMDD = recurrent major depressive disorder; SCZ = schizophrenia; SE = Standard Error.

**Supplementary Table 13. Selected pathway gene set enrichment analyses in PGC+iPSYCH**

See SupplTable13_ Selected pathway GSEA_PGC+iPSYCH.xlsx

Analyses were run with (top) and without (bottom) inclusion of the Chromosome 6 MHC region in MAGMA. These analyses were carried out on the genomic control output with INFO score > 0.6, European ancestry only, and autosomal SNPs only. * Significant after adjusting *p*-values for multiple testing. Primary model refers to the regression model without additional interaction covariates; secondary model refers to the extended model with additional interaction covariates.

Abbreviations: BIP = bipolar disorder; CNS = central nervous system; MDD = major depressive disorder; MP = Mouse Phenome; $P_{FDR}$ = False Discovery Rate-corrected *p*-value; PGC-NPA = Psychiatric Genomics Consortium – Network and Pathway Analysis Working Group; rMDD = recurrent major depressive disorder; SCZ = schizophrenia; SE = Standard Error.

**Supplementary Table 14. Lookup of interaction for SNPs showing sex-stratification or GxS interaction in 23andme, PGC, and UK Biobank**

See SupplTable14_Prior_GWAS_SNP_Lookups.xlsx

Interaction results for the SNPs identified in sex-stratified analyses of other disorders and phenotypes, as well as SNPs identified in the recent PGC Crosss-Disorder GWAS. Reported are replications/validations with nominal *p*-values < 0.01 in the interaction study.

Abbreviations: BP = base pair position; CHR = Chromosome; SE = Standard Error; SNP = Single Nucleotide Polymorphism rs ID.

### Supplementary Figures

**Supplementary Figure 1. Experimental Design.**

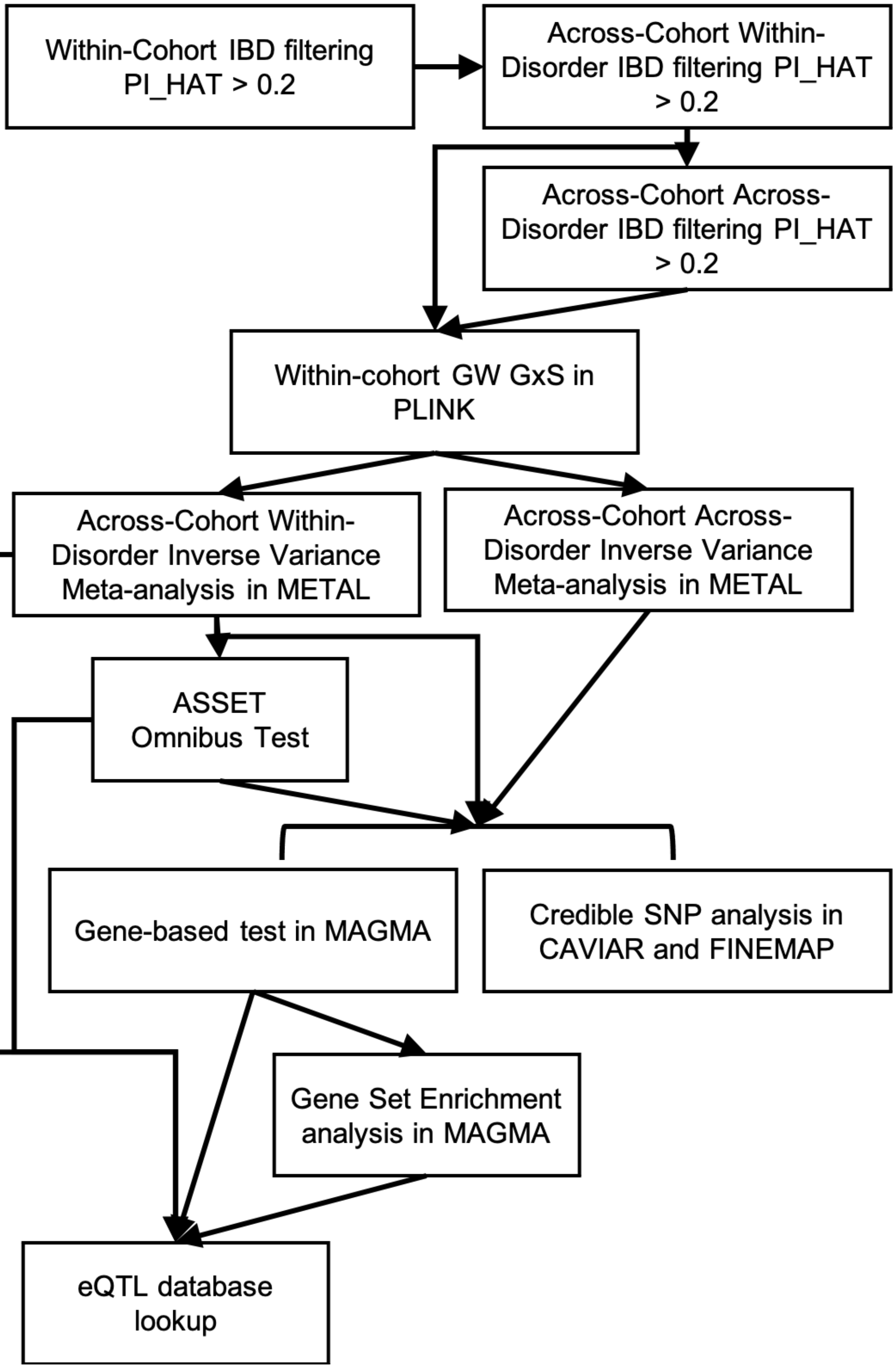

**Supplementary Figure 2. Power analyses.**
Abbreviations: MAF = Minor Allele Frequency; OR = Odds Ratio.

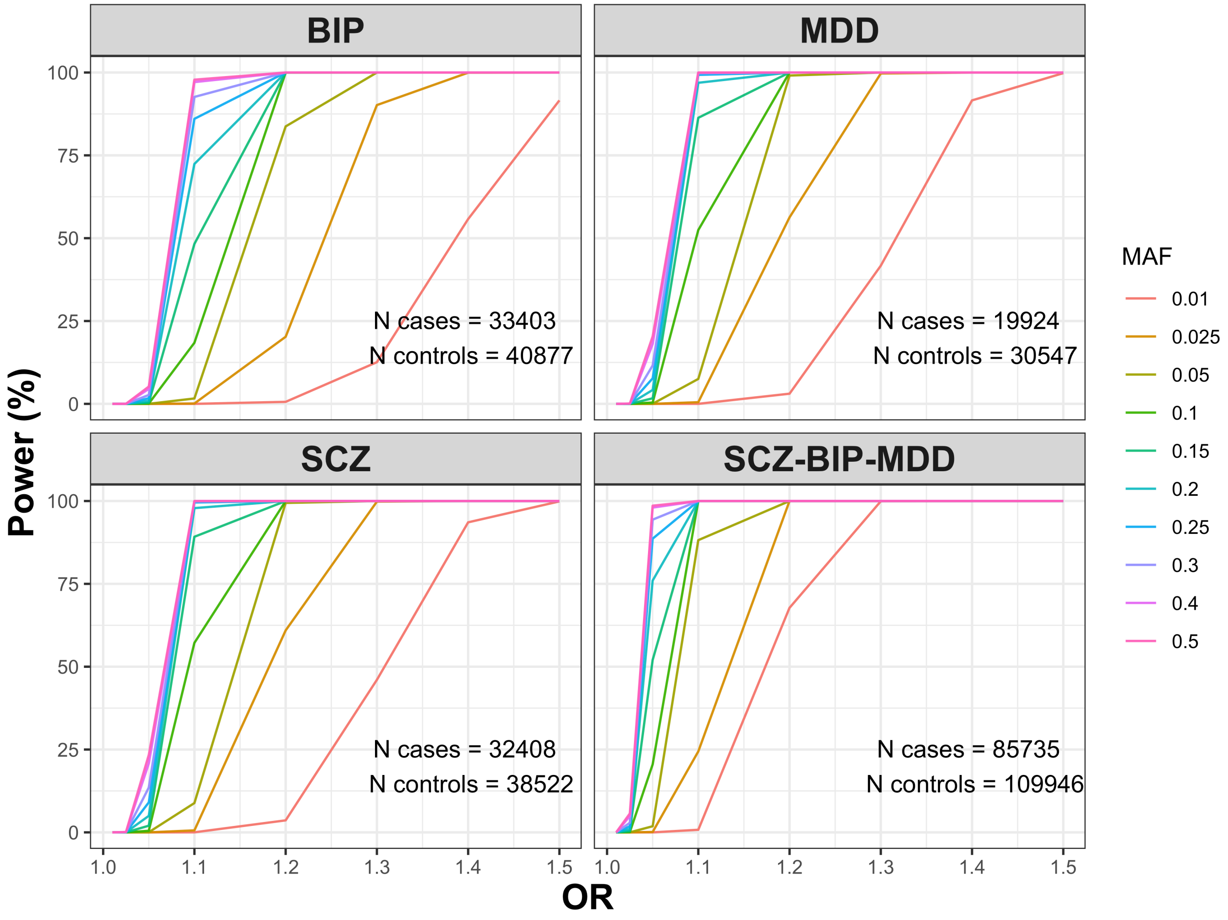

**Supplementary Figure 3. Miami plots for sex-stratified analyses in PGC + iPSYCH**

GWAS SNP main effects for men (blue) are plotted downward, and are plotted upward for women (green). Negative log10-transformed p-values for each variant (each dot) (y-axis) are plotted by chromosomal position (x-axis). The solid red and dotted blue horizontal lines represent the thresholds for genome-wide significant association ($p = 5\times10^{-8}$) and suggestive association ($p = 1\times10^{-5}$), respectively. Plotted are the regular meta-analysis results within and across disorders only; omnibus tests were not carried out for sex-stratified analyses. Plots were generated using the plot package in R.

Abbreviations: BIP = Bipolar Disorder; (r)MDD = (recurrent) Major Depressive Disorder; SCZ = Schizophrenia

### a) Schizophrenia – European ancestry only

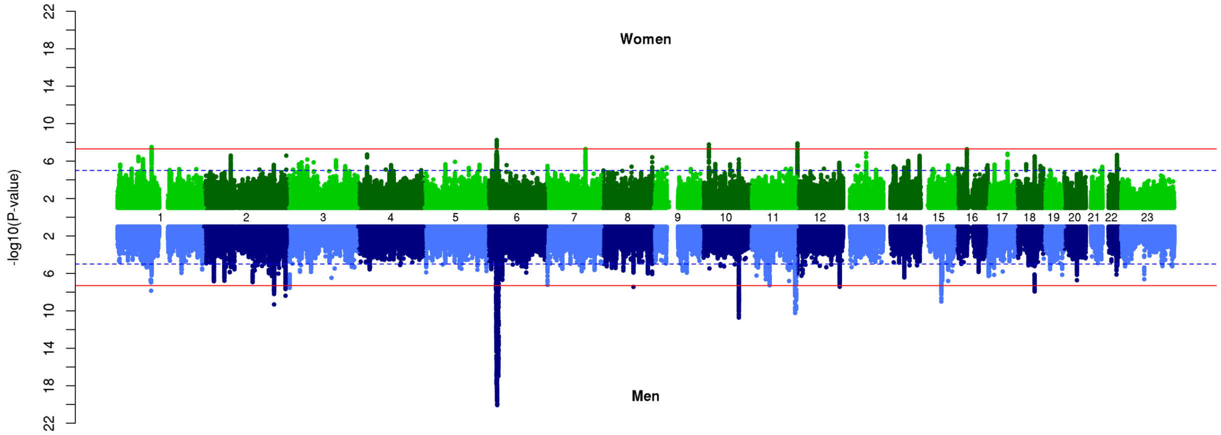


### b) Schizophrenia – European + East Asian ancestry

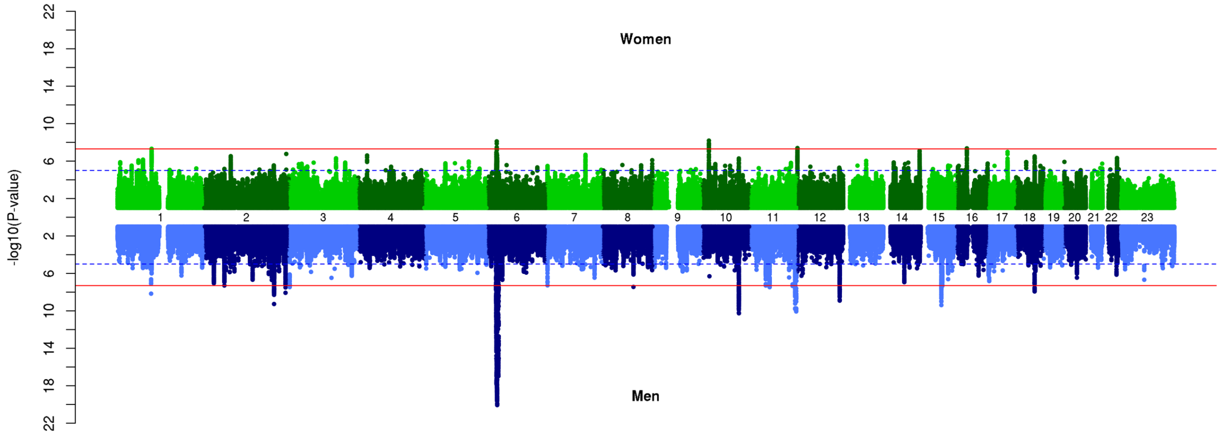

### c) Bipolar Disorder

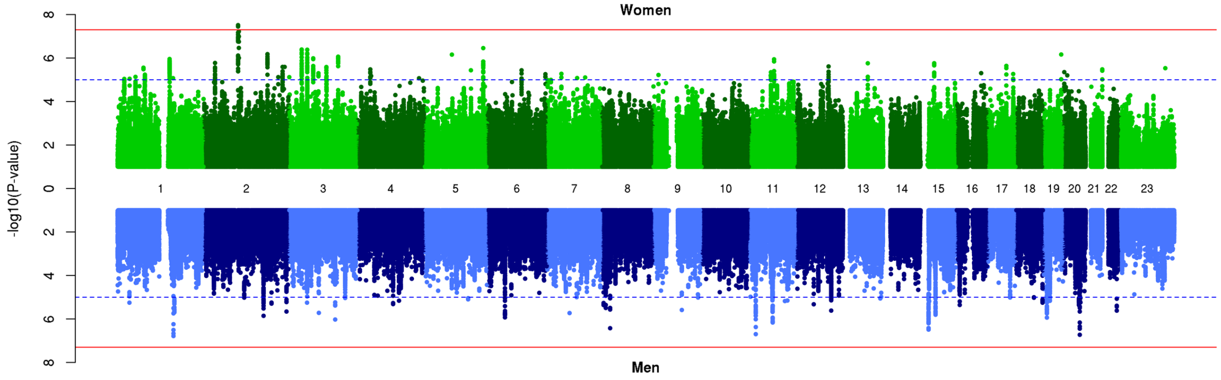


### d) Major Depressive Disorder

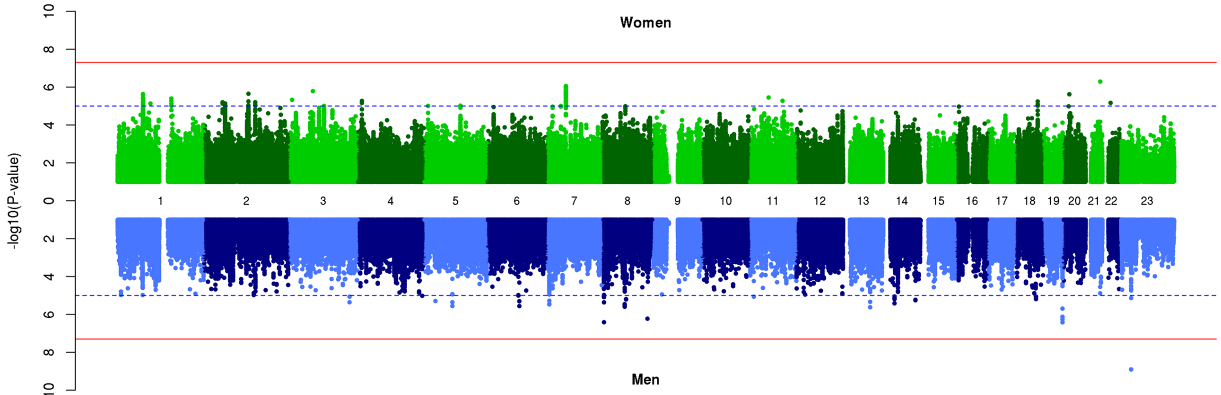


### e) Cross-Disorder SCZ-BIP-MDD – European ancestry only

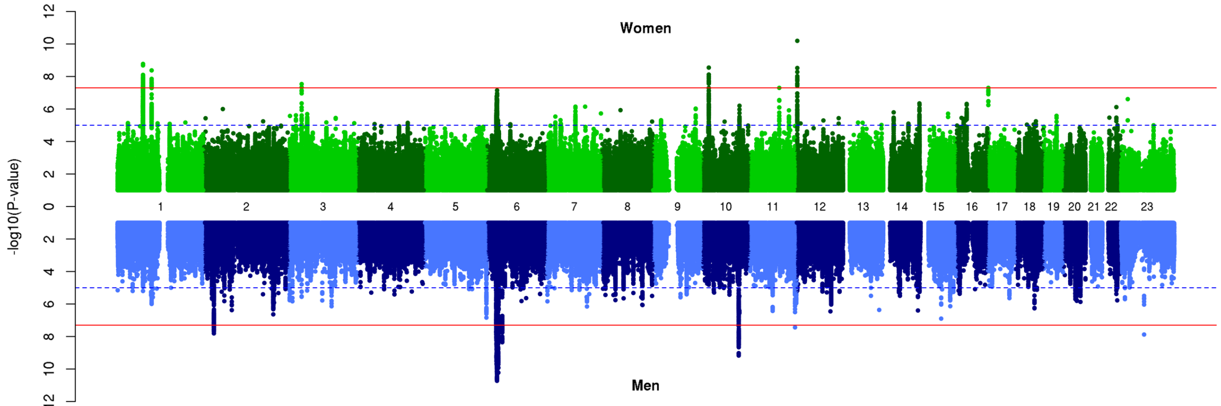

### f) Cross-Disorder SCZ-BIP-MDD – European + East Asian ancestry

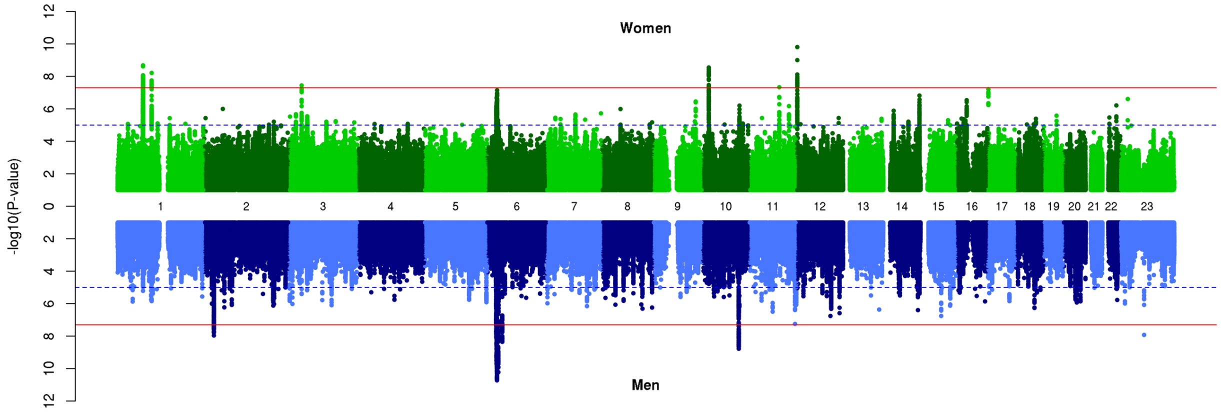


### g) Cross-Disorder SCZ-BIP-rMDD – European ancestry only

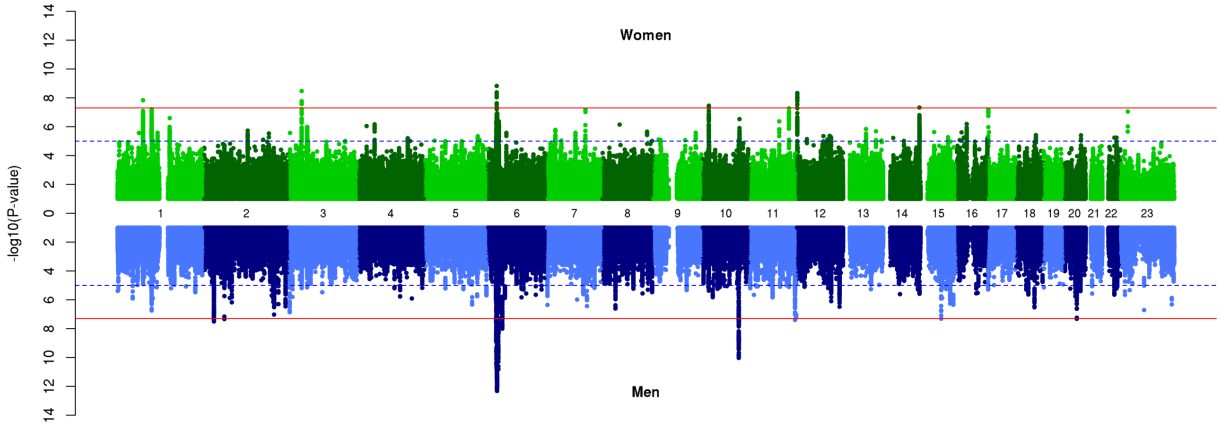


### h) Cross-Disorder SCZ-BIP-rMDD – European + East Asian ancestry

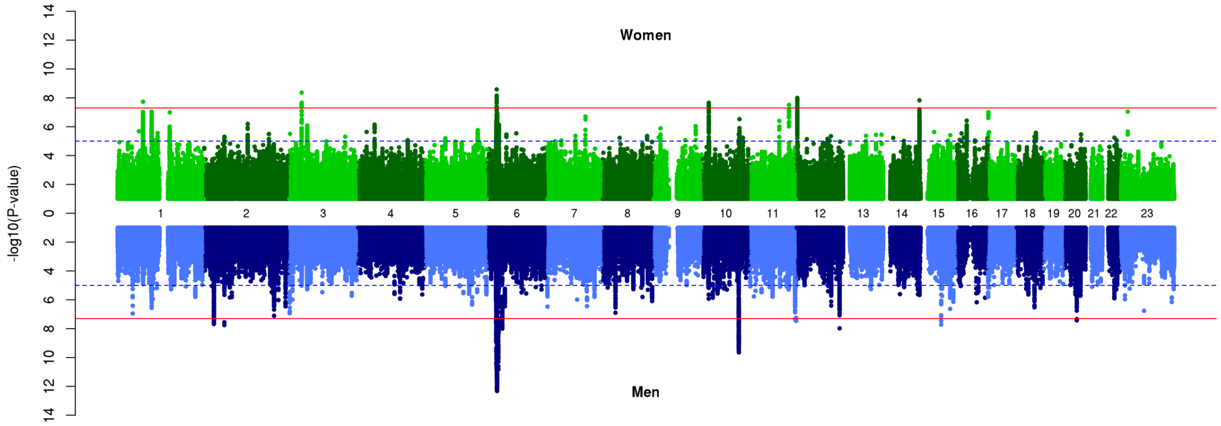

### i) Recurrent Major Depressive Disorder

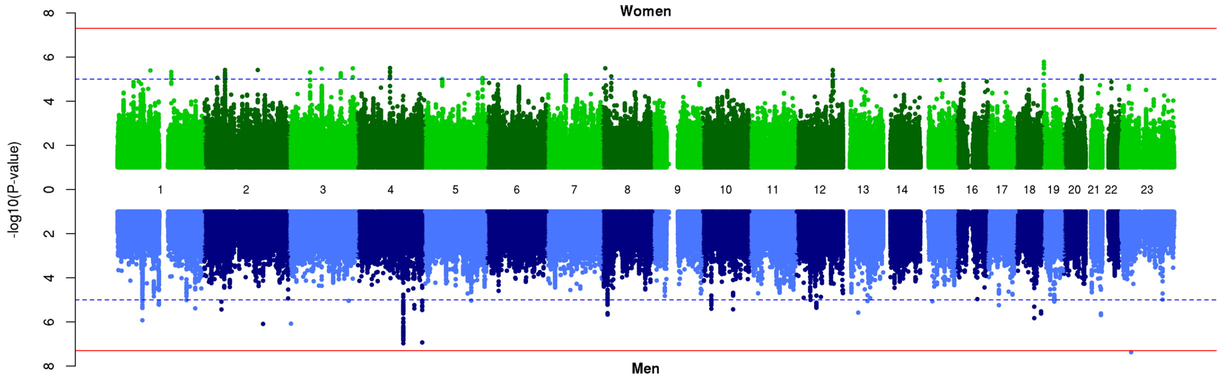

**Supplementary Figure 4. Scatter plots of female vs male associations in PGC + iPSYCH**

The scatter plots show little correlation (R) between GWAS SNP main effect *p*-values from the two sexes, indicating the strength of association differed substantially between the two sexes. Plots were generated using the plot package in R.

Abbreviations: BIP = Bipolar Disorder; (r)MDD = (recurrent) Major Depressive Disorder; SCZ = Schizophrenia; $R^2$ = proportion variance explained.

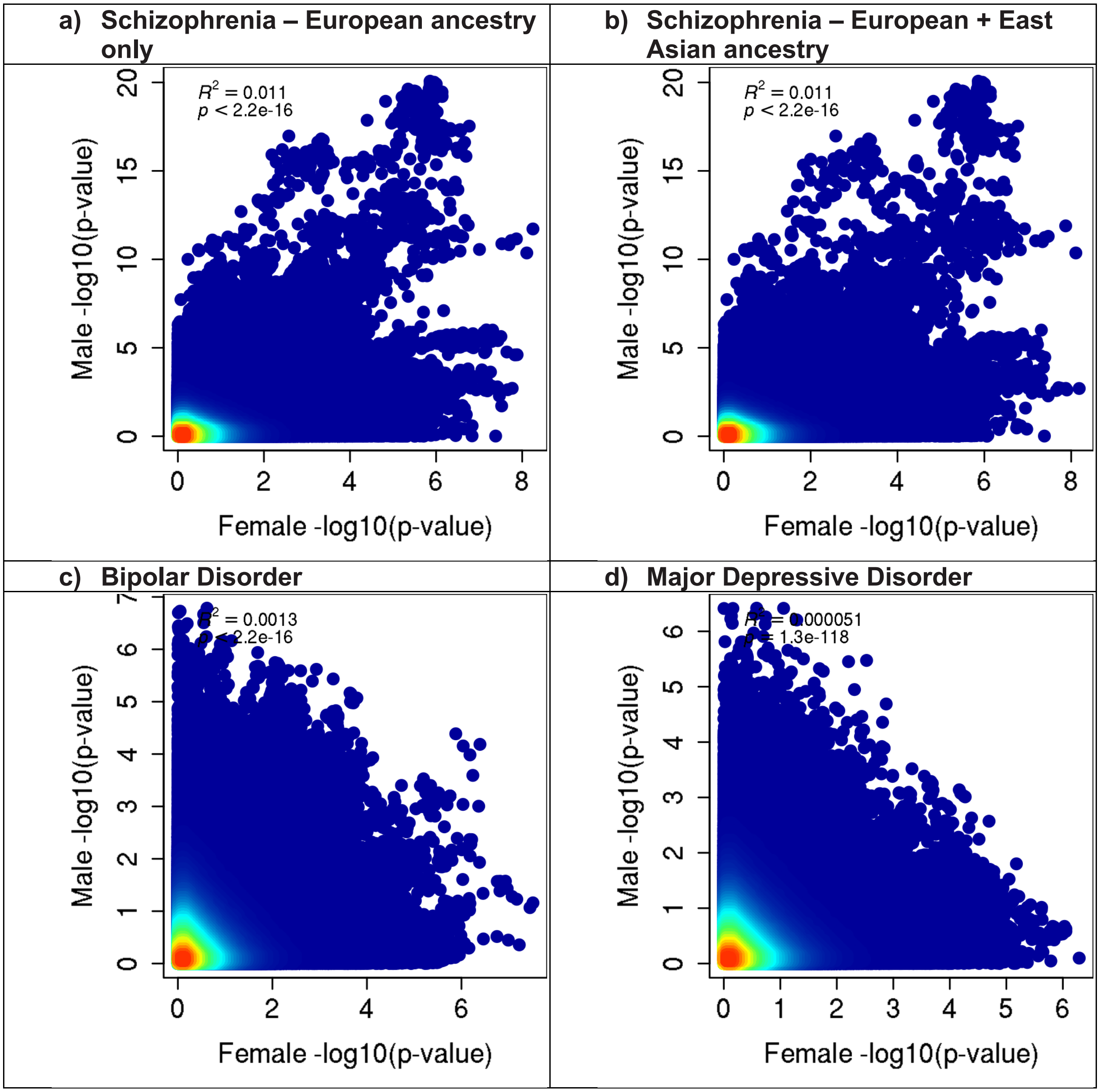

**e) Recurrent Major Depressive Disorder**

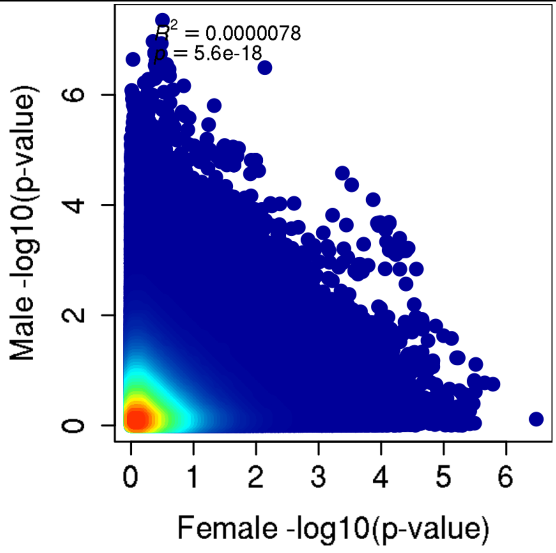


**f) Cross-Disorder SCZ-BIP-MDD – European ancestry only**

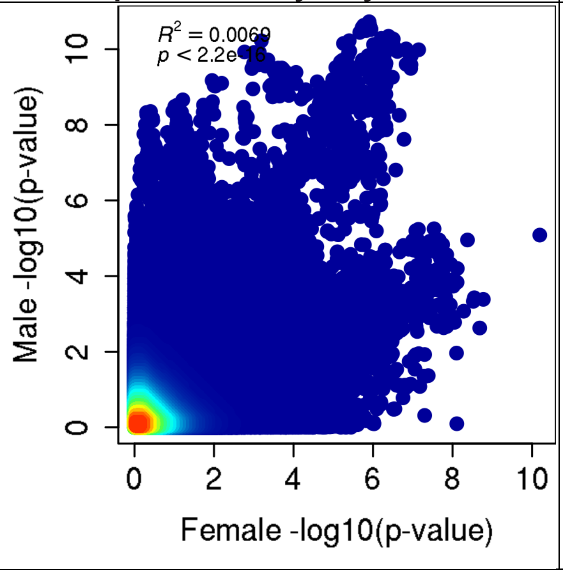


**g) Cross-Disorder SCZ-BIP-MDD – European + East Asian ancestry**

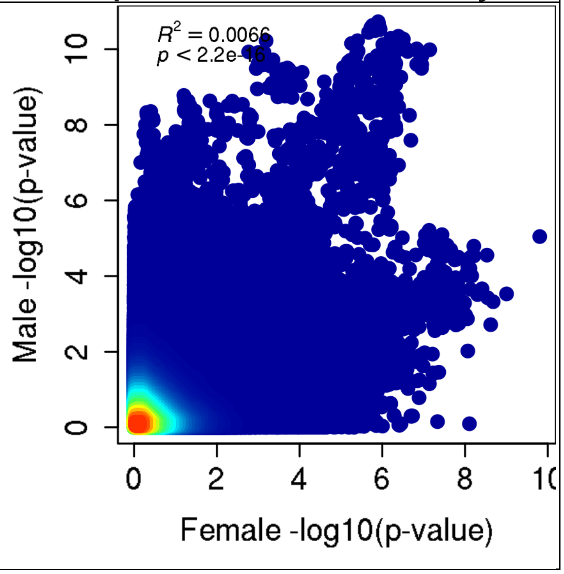

| h) Cross-Disorder SCZ-BIP-rMDD – European ancestry only | i) Cross-Disorder SCZ-BIP-rMDD – European + East Asian ancestry |
| --- | --- |
| 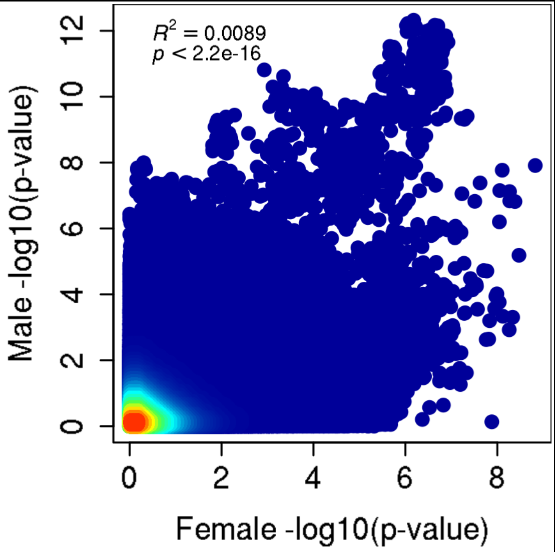 | 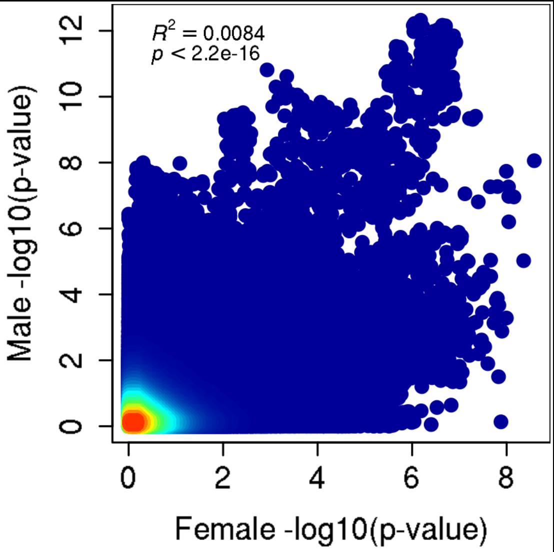 |

**Supplementary Figure 5. Quantile-Quantile (Q-Q) plots for GxS interaction in PGC + iPSYCH**

The Q-Q plot is used to assess the number and magnitude of observed associations compared with the expectations under no association. The nature of deviations from the identity line provide clues whether the observed associations are true associations or may be due to for example population stratification or cryptic relatedness.

Abbreviations: BIP = Bipolar Disorder; (r)MDD = (recurrent) Major Depressive Disorder; SCZ = Schizophrenia

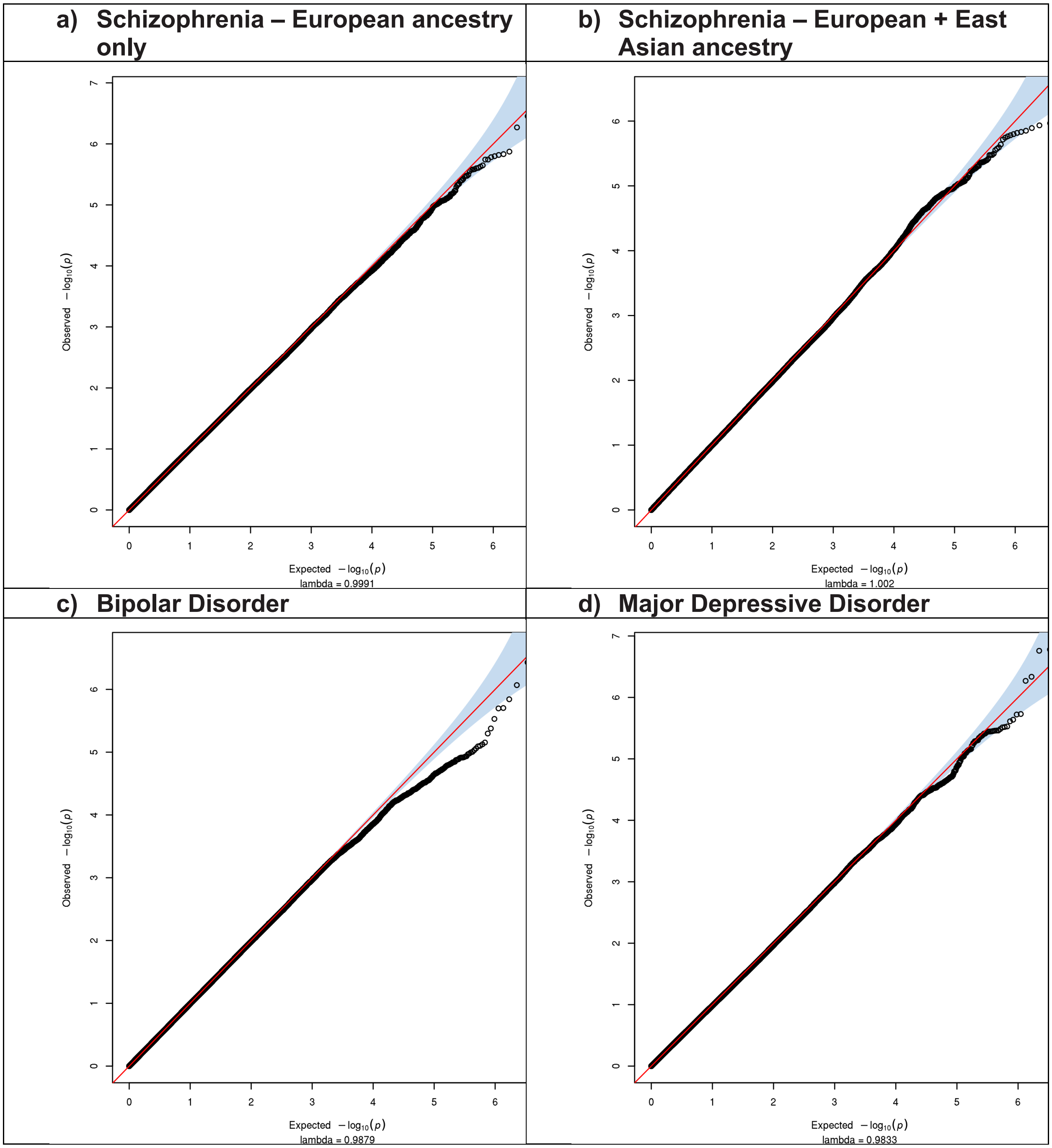

**e) Recurrent Major Depressive Disorder**

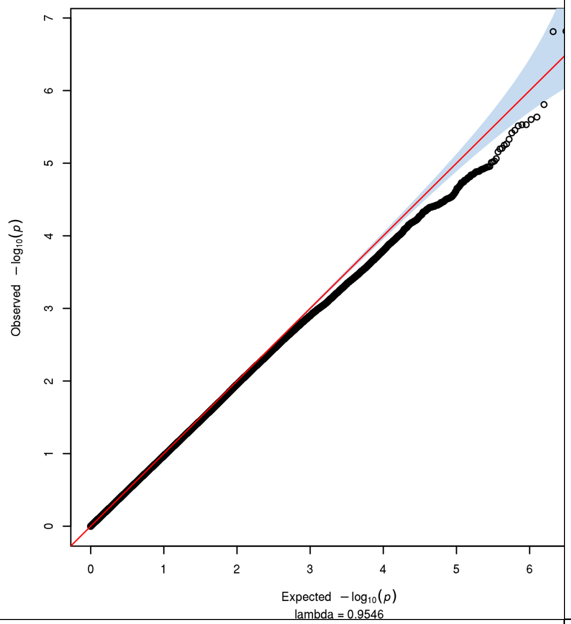


**f) Cross-Disorder SCZ-BIP-MDD – European ancestry only**

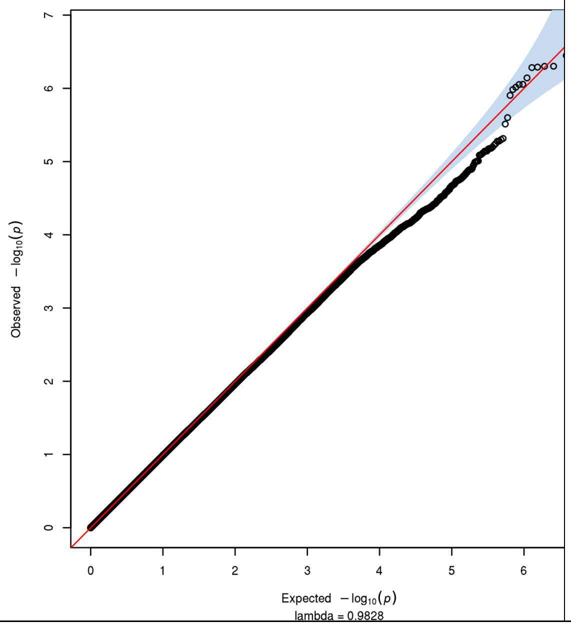


**g) Cross-Disorder SCZ-BIP-MDD – European + East Asian ancestry**

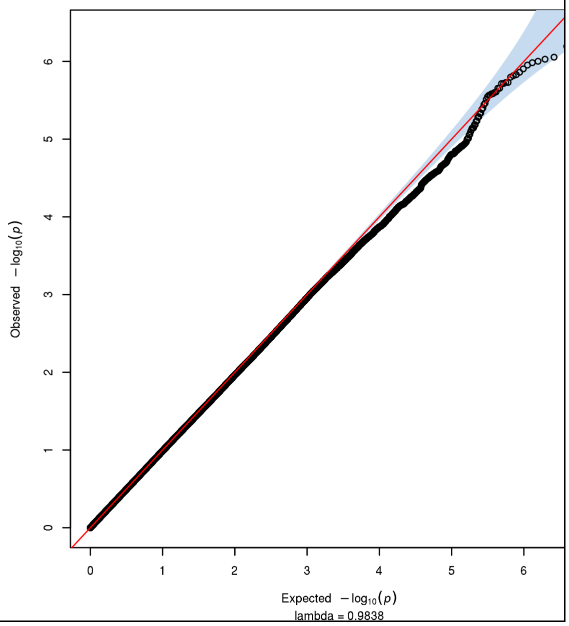

**h) Cross-Disorder SCZ-BIP-rMDD – European ancestry only**

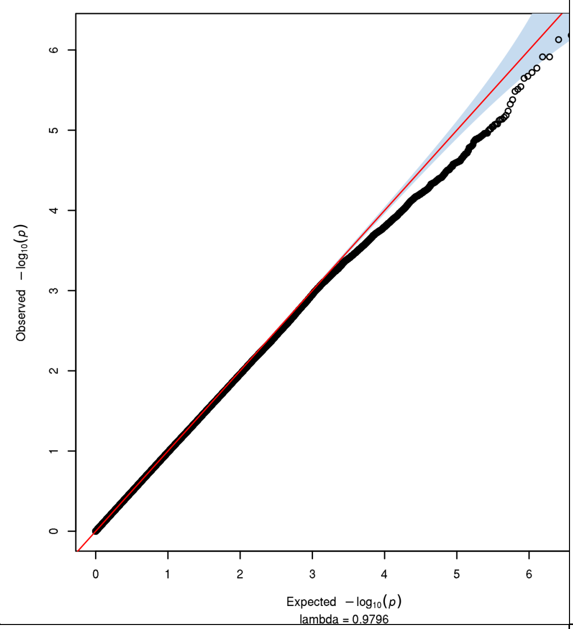


**i) Cross-Disorder SCZ-BIP-rMDD – European + East Asian ancestry**

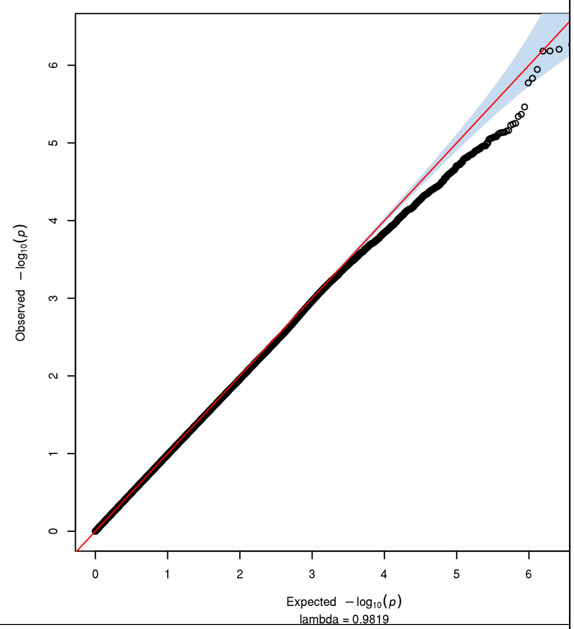


**j) Omnibus Test SCZ-BIP-MDD – European ancestry only**

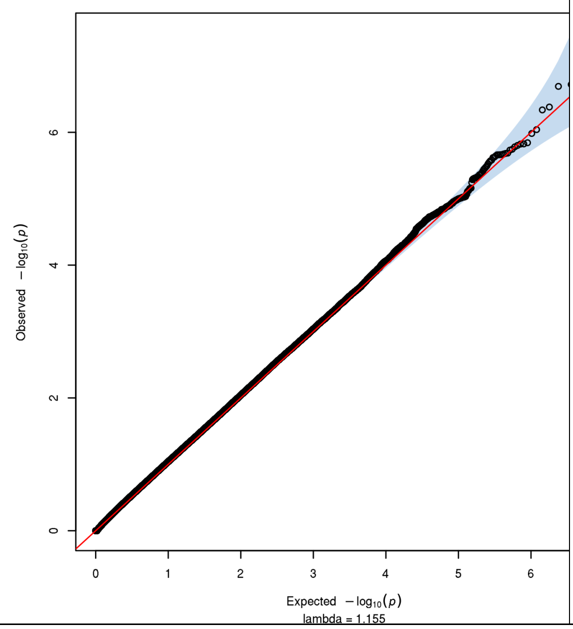


**k) Omnibus Test SCZ-BIP-MDD – European + East Asian ancestry**

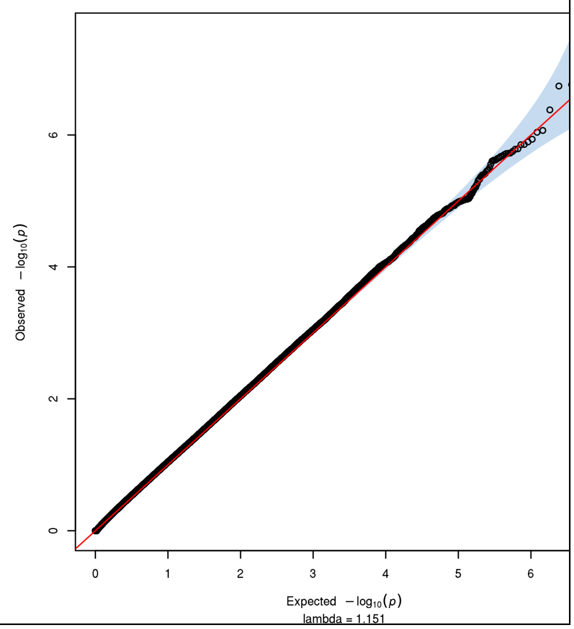

| l) Omnibus Test SCZ-BIP-rMDD – European ancestry only | m) Omnibus Test SCZ-BIP-rMDD – European + East Asian ancestry |
|---|---|

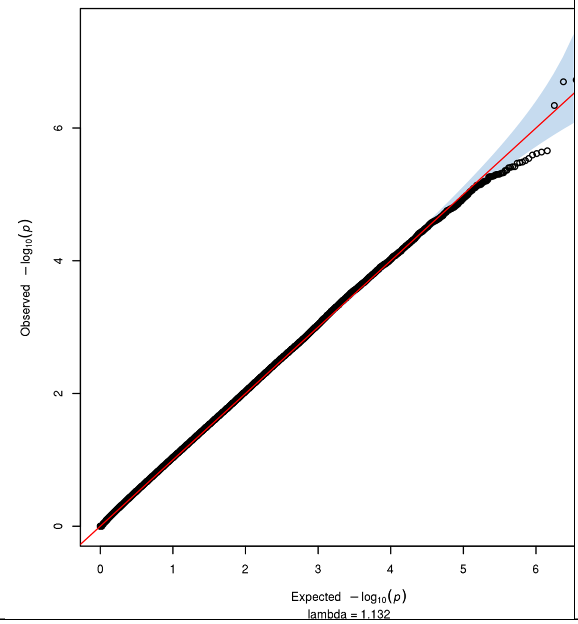


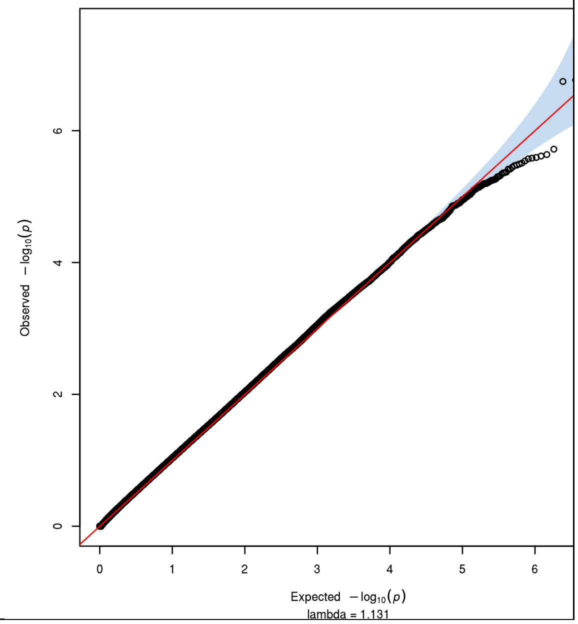

**Supplementary Figure 6. Manhattan plots of the GxS interaction GWAS in PGC + iPSYCH**

Negative log10-transformed p-values for each variant (each dot) (y-axis) are plotted by chromosomal position (x-axis). The red and blue lines represent the thresholds for genome-wide significant association ($p = 5{\times}10^{-8}$) and suggestive association ($p = 1{\times}10^{-5}$), respectively. P-values for X chromosome (23) model B (alleles: females 0, 1, or 2; males 0 or 1) are included.

Abbreviations: BIP = Bipolar Disorder; (r)MDD = (recurrent) Major Depressive Disorder; SCZ = Schizophrenia

**a) Schizophrenia – European ancestry only**

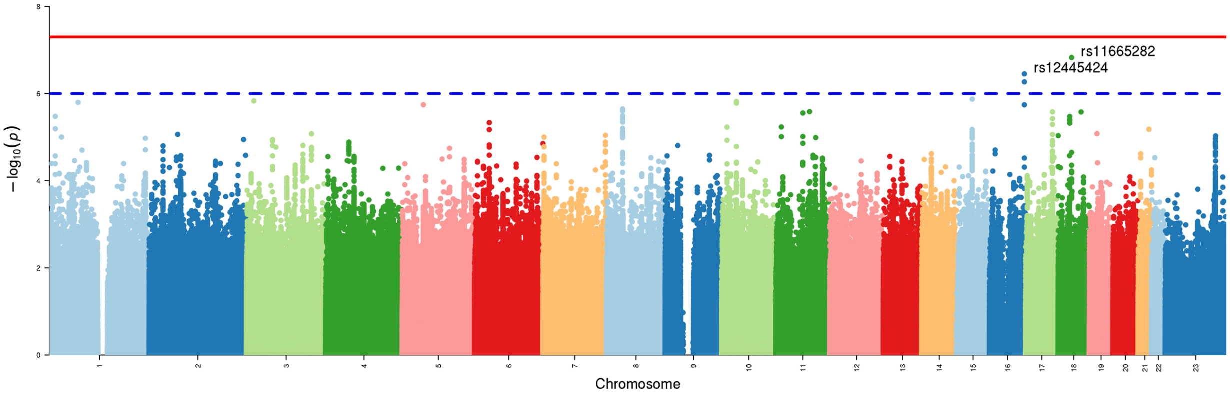


**b) Schizophrenia – European + East Asian ancestry**

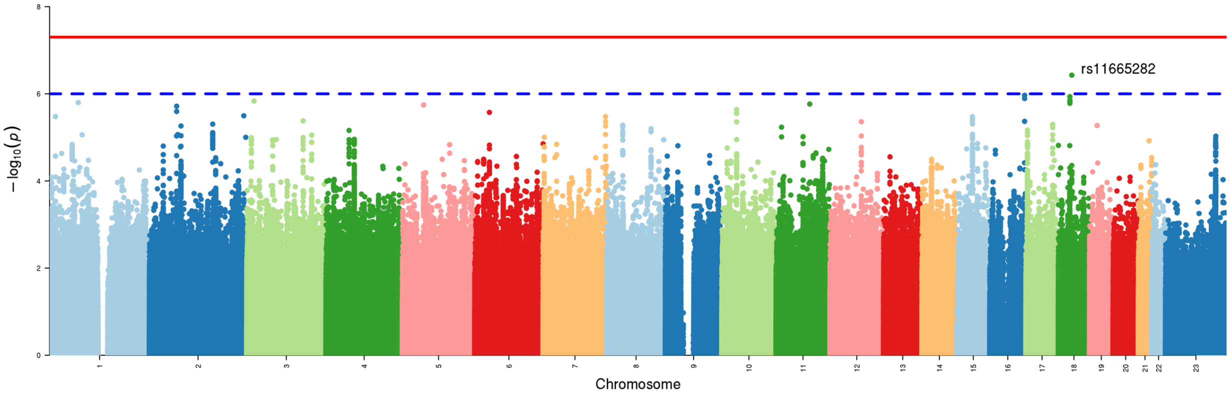


**c) Bipolar Disorder**

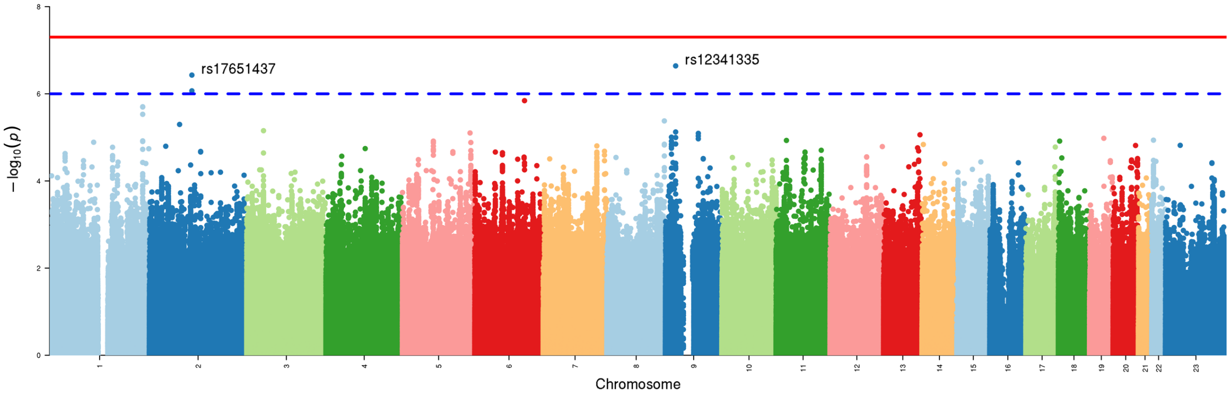

### d) Major Depressive Disorder

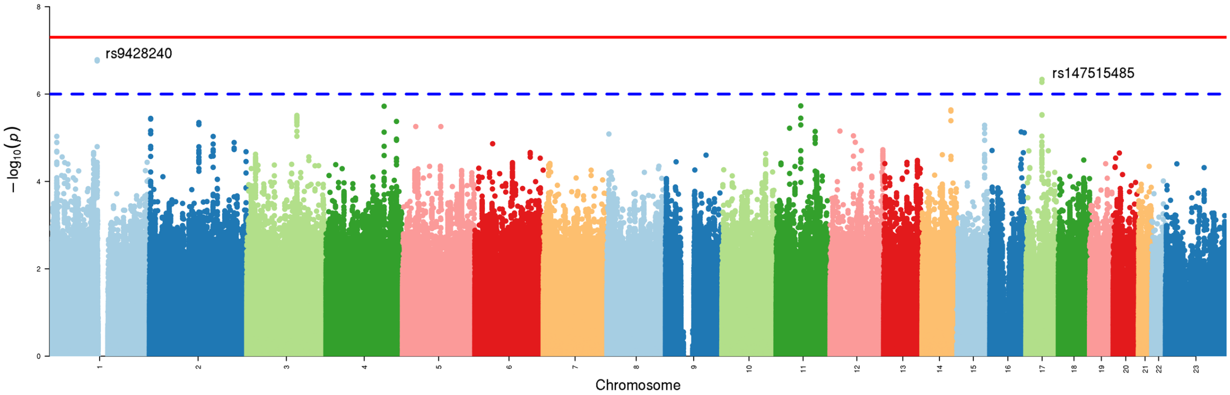


### e) Recurrent Major Depressive Disorders

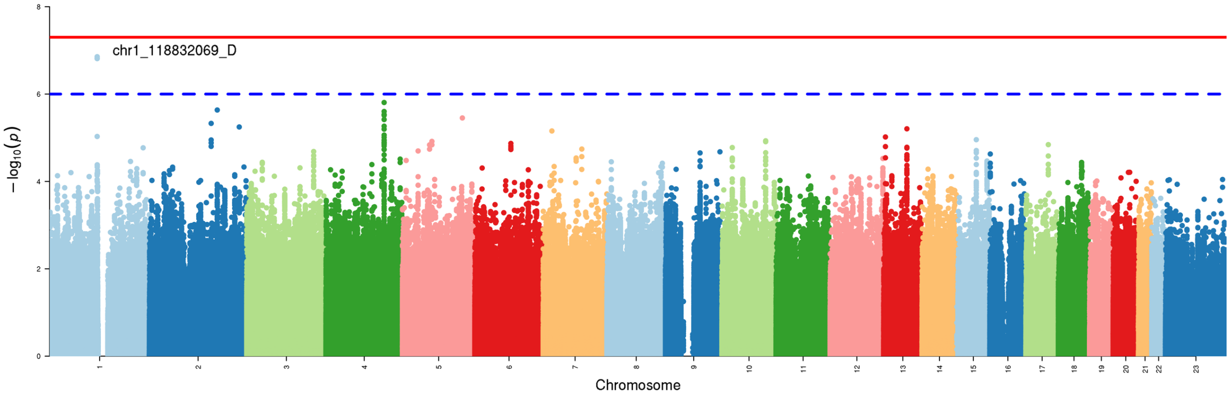


### f) Cross-Disorder SCZ-BIP-MDD – European ancestry only

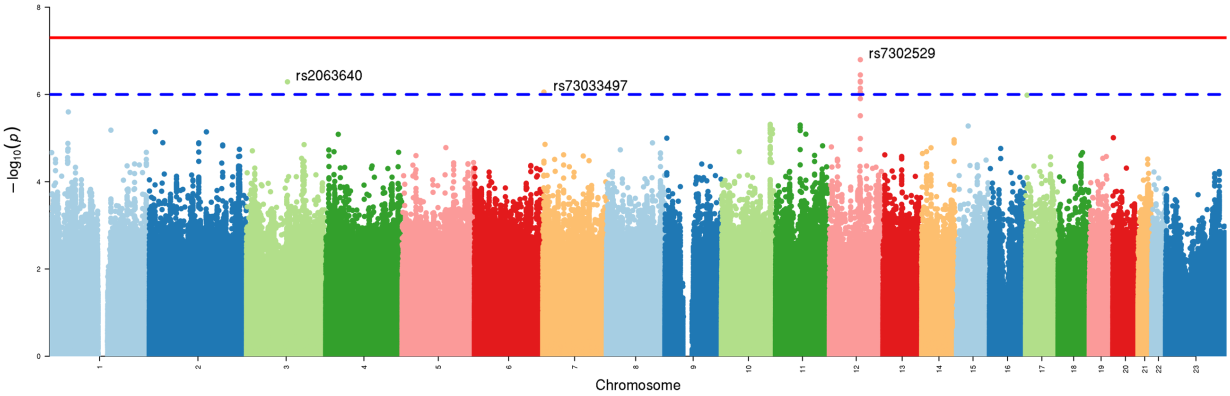

### g) Cross-Disorder SCZ-BIP-MDD – European + East Asian ancestry

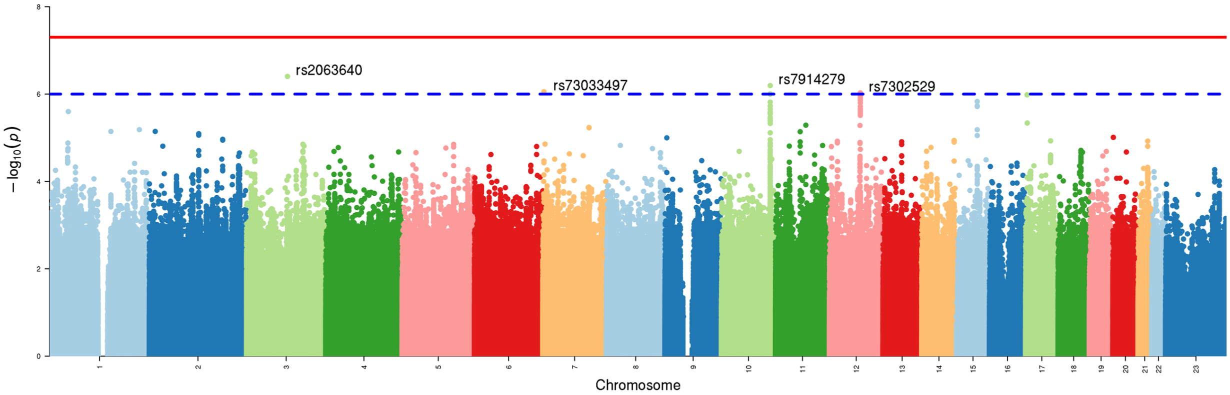


### h) Cross-Disorder SCZ-BIP-rMDD – European ancestry only

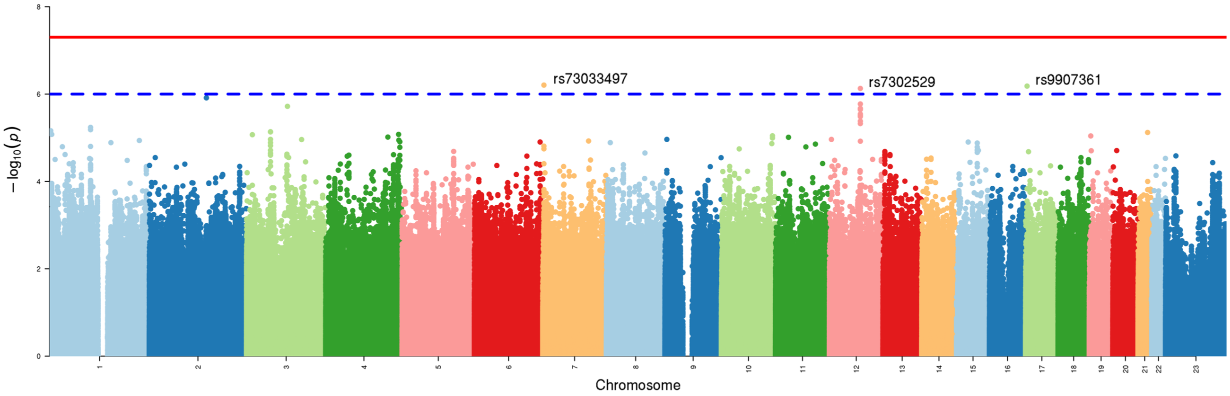


### i) Cross-Disorder SCZ-BIP-rMDD – European + East Asian ancestry

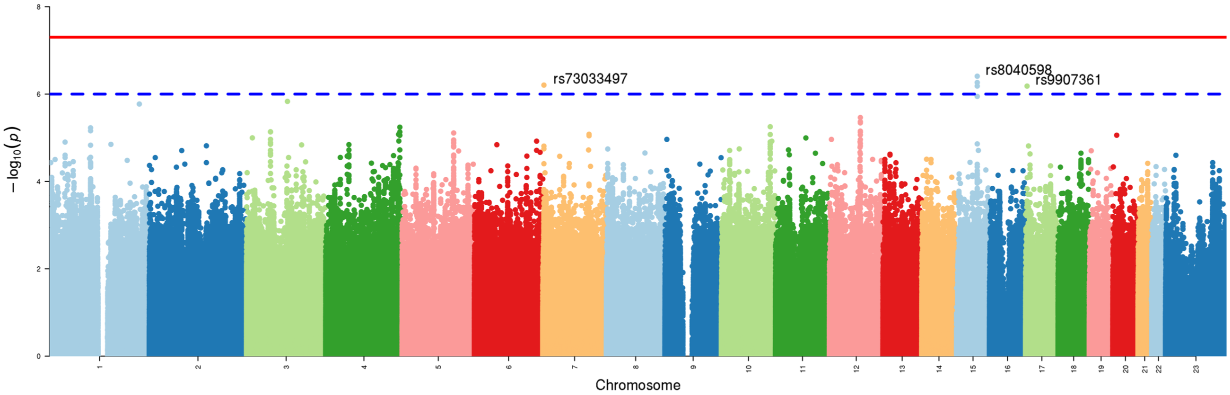

### j) Omnibus Test SCZ-BIP-MDD – European ancestry only

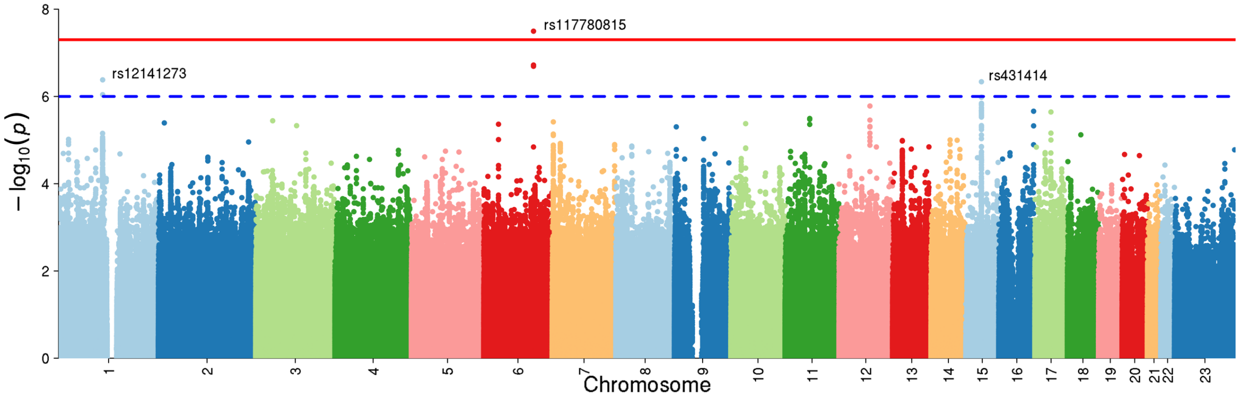


### k) Omnibus Test SCZ-BIP-MDD – European + East Asian ancestry

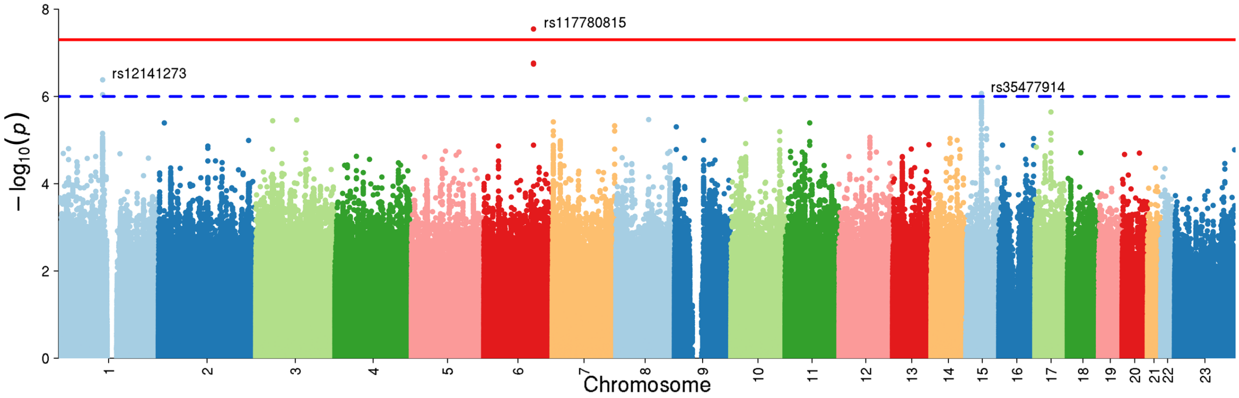


### l) Omnibus Test SCZ-BIP-rMDD – European ancestry only

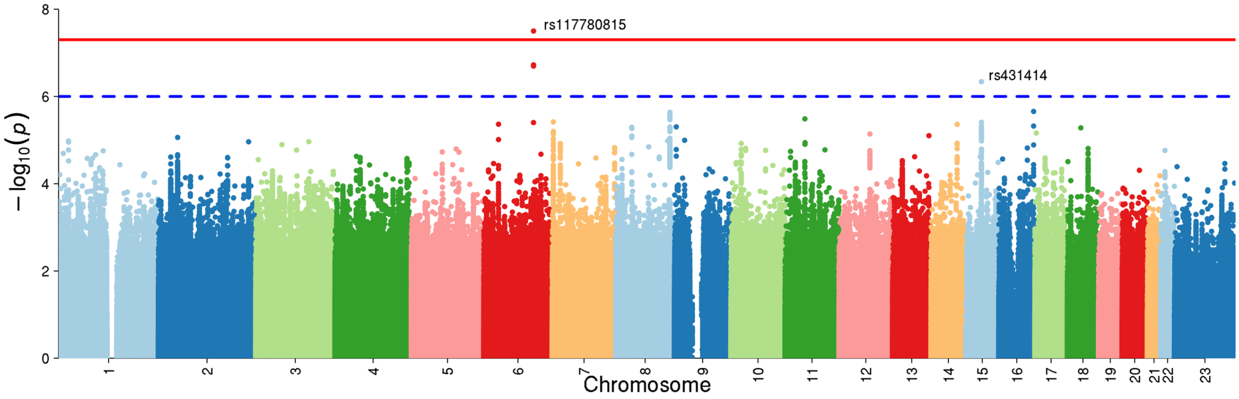

### m) Omnibus Test SCZ-BIP-rMDD – European + East Asian ancestry

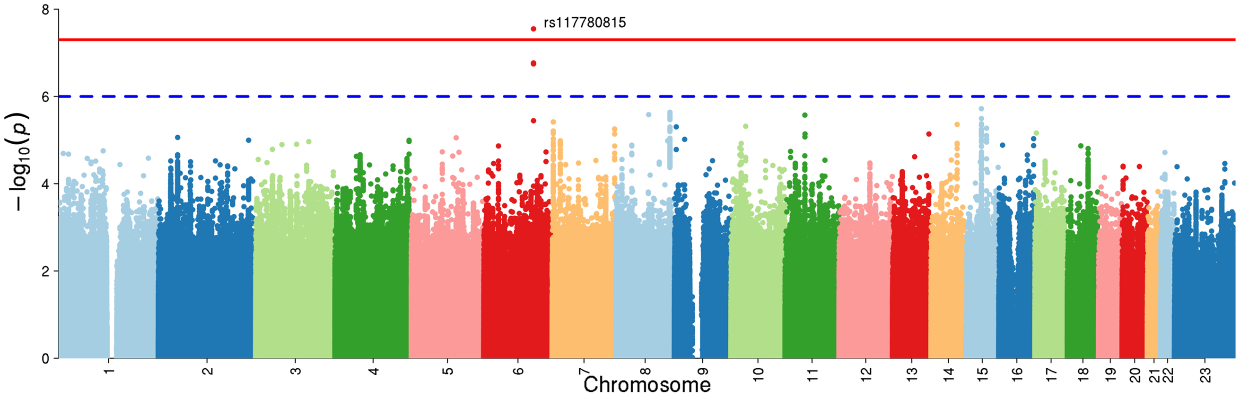

**Supplementary Figure 7. LocusZoom plots for loci with GxS interaction in PGC + iPSYCH**

Plots were generated using the LocusZoom 1.4 Standalone application (49) for loci with GxS interaction p < 1×10$^{-6}$.

Abbreviations: chr = chromosome; cM = centimorgans; Mb = megabases; $r^2$ = linkage disequilibrium level; BIP = Bipolar Disorder; (r)MDD = (recurrent) Major Depressive Disorder; SCZ = Schizophrenia

### a) Schizophrenia – European ancestry only

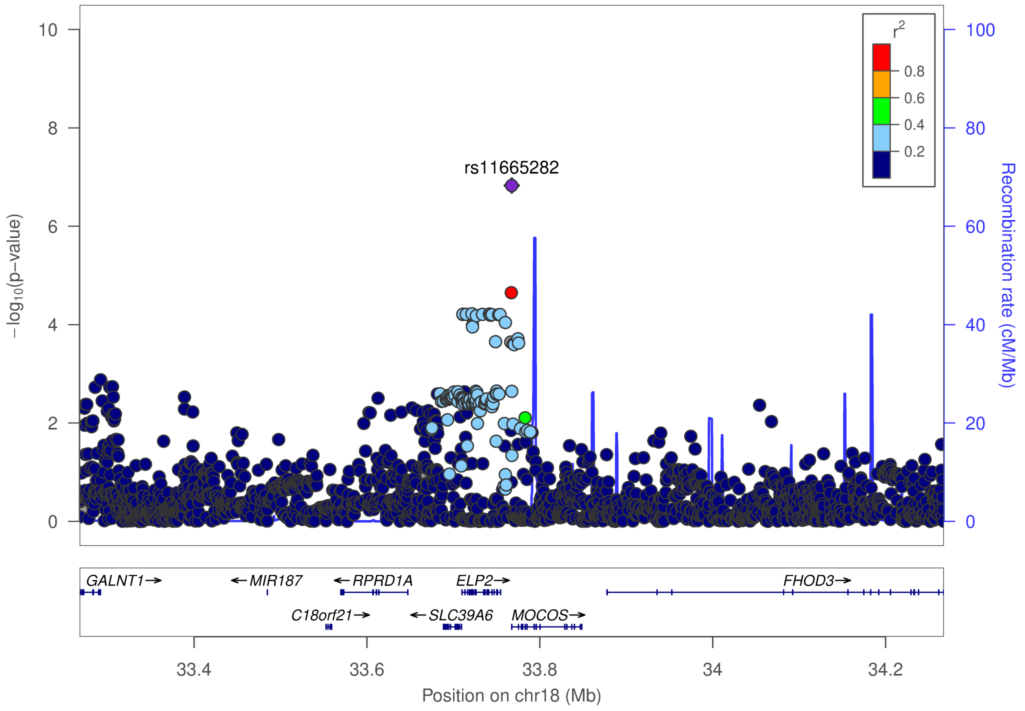

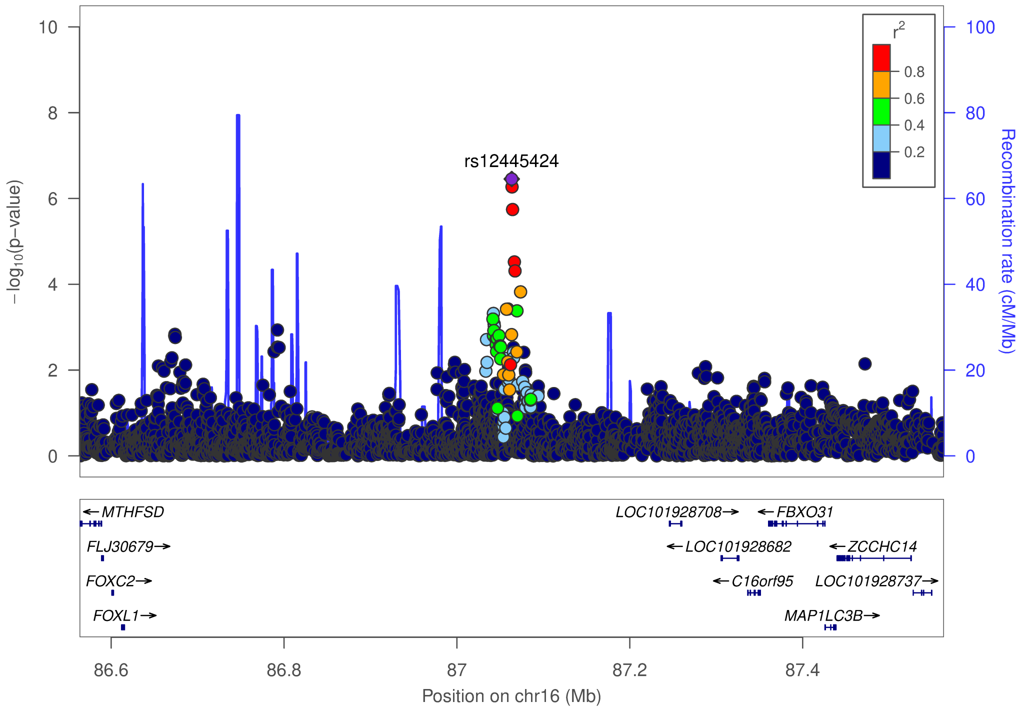


### b) Schizophrenia – European + East Asian ancestry

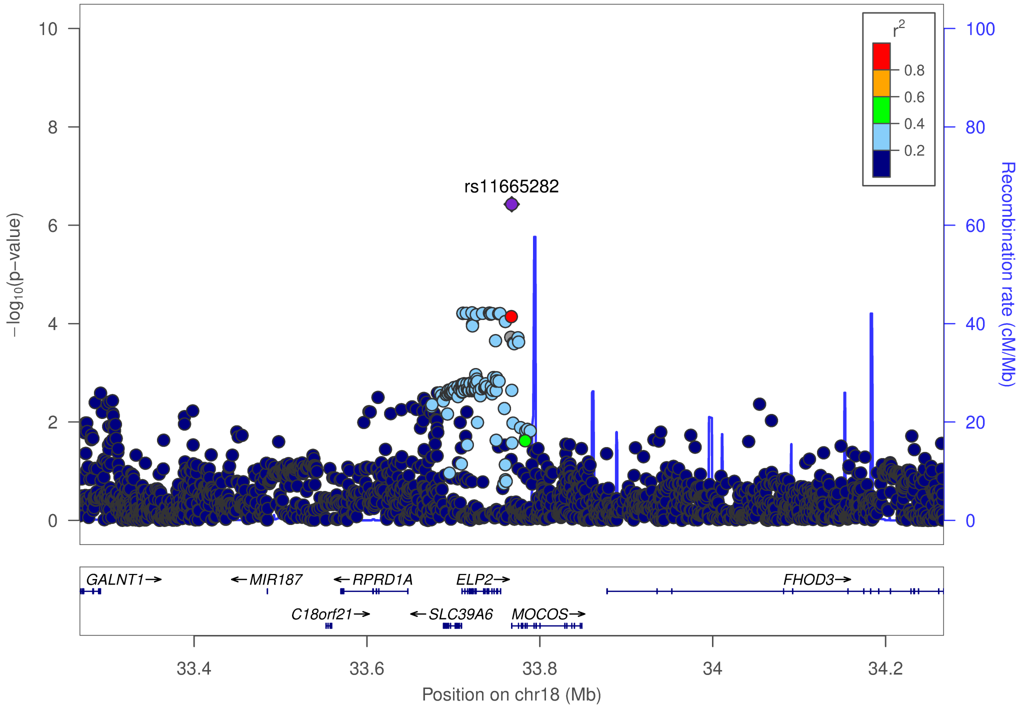

### c) Bipolar Disorder

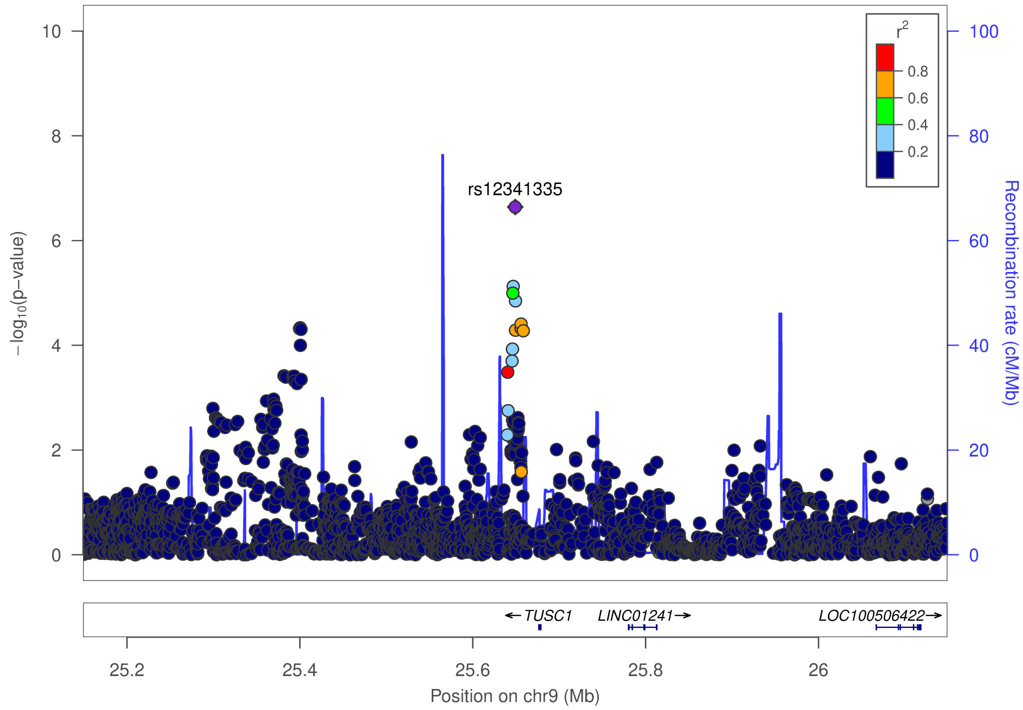


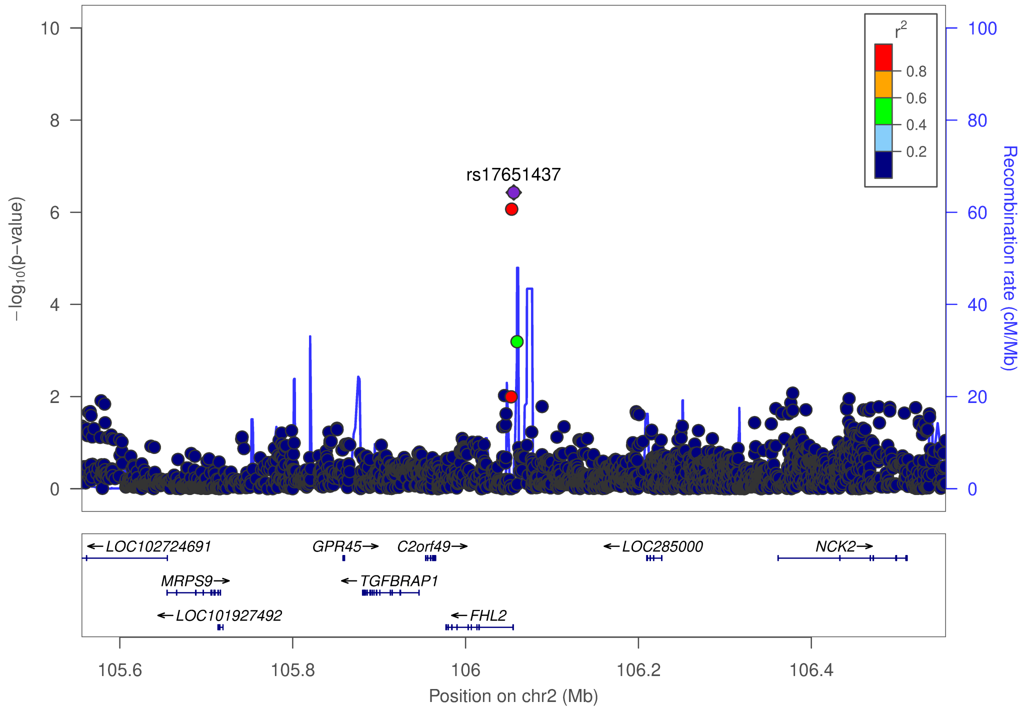

### d) Major Depressive Disorder

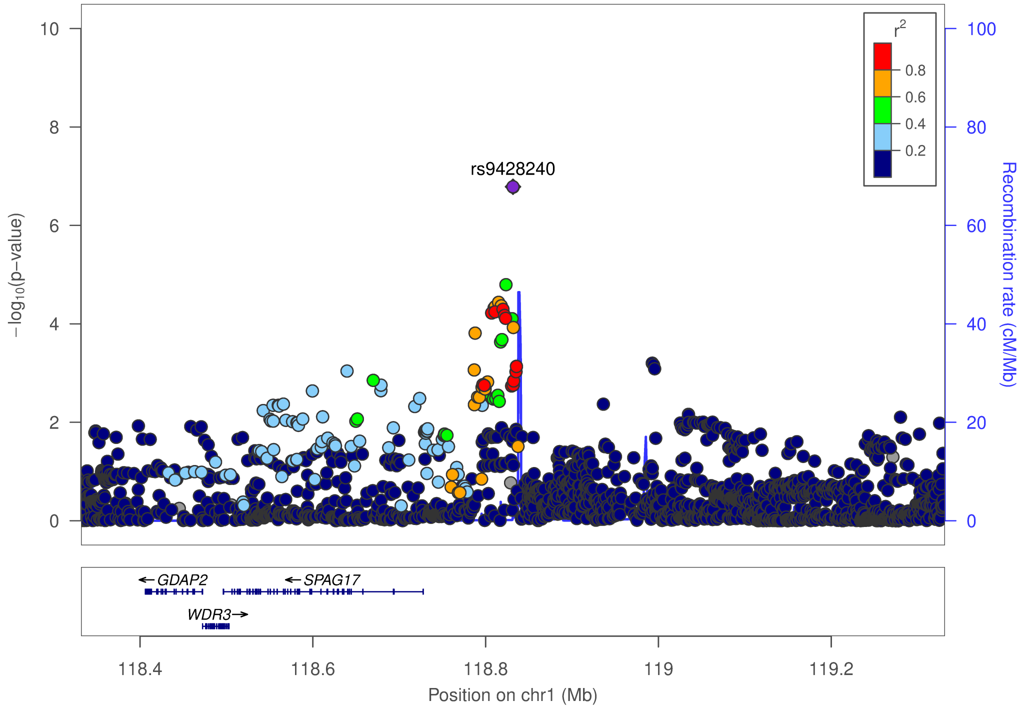


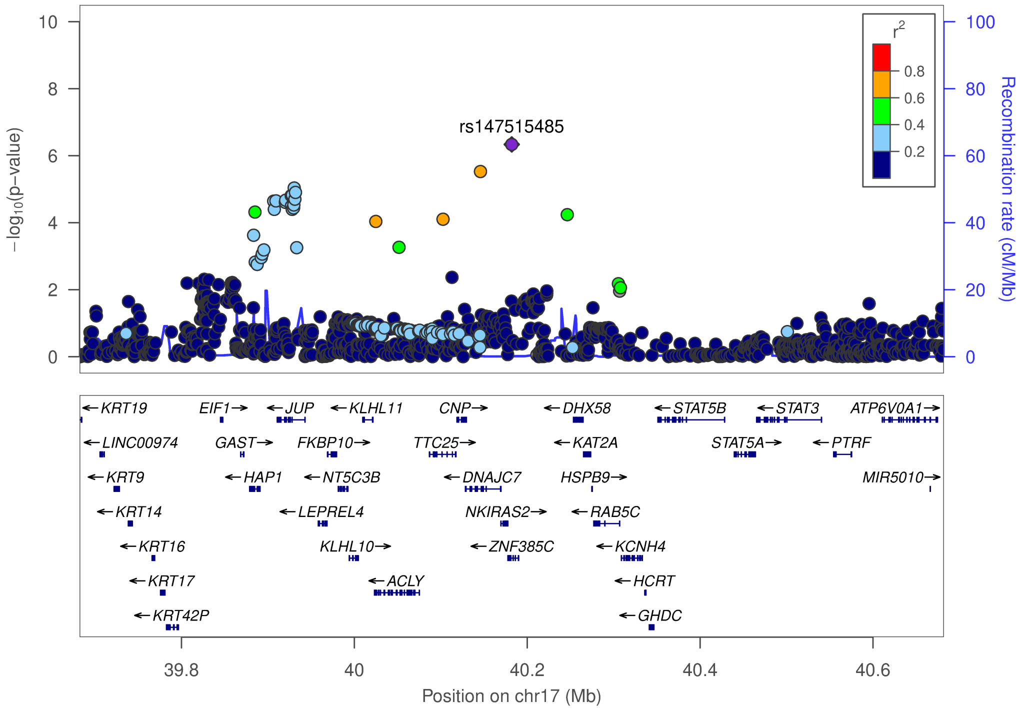

### e) Recurrent Major Depressive Disorder

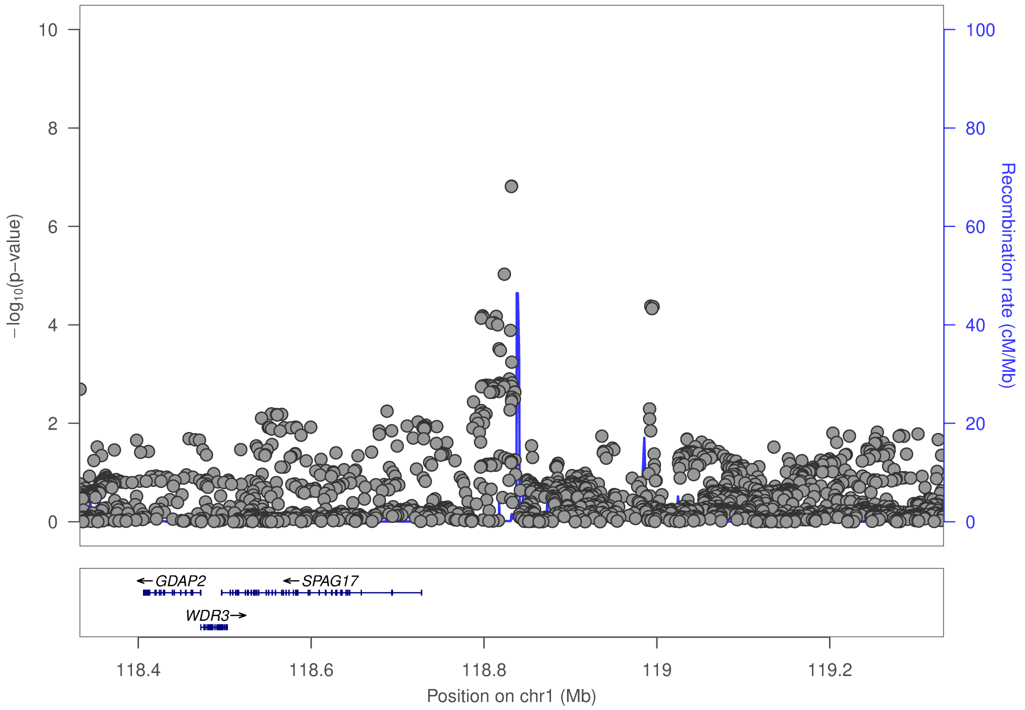


### f) Cross-Disorder SCZ-BIP-MDD – European ancestry only

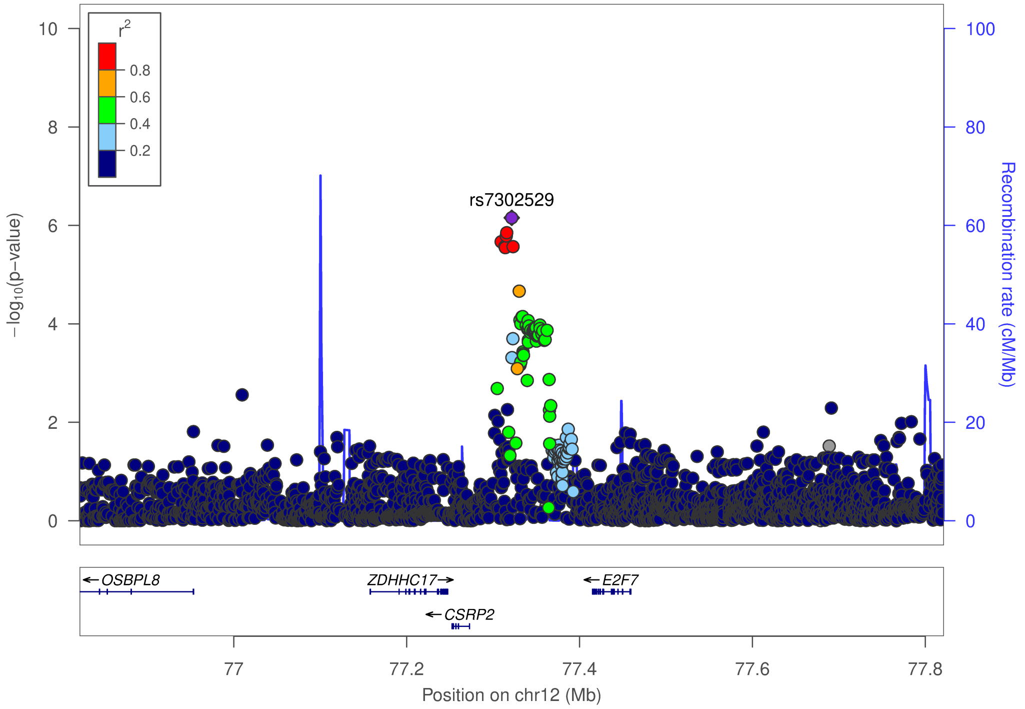

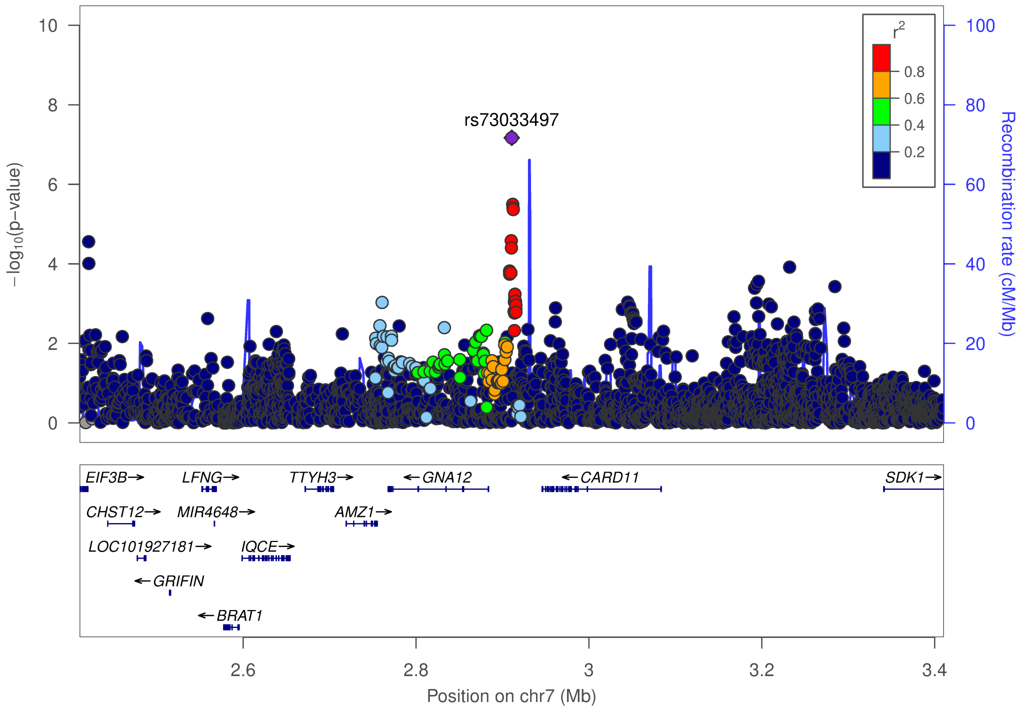


### g) Cross-Disorder SCZ-BIP-MDD – European + East Asian ancestry

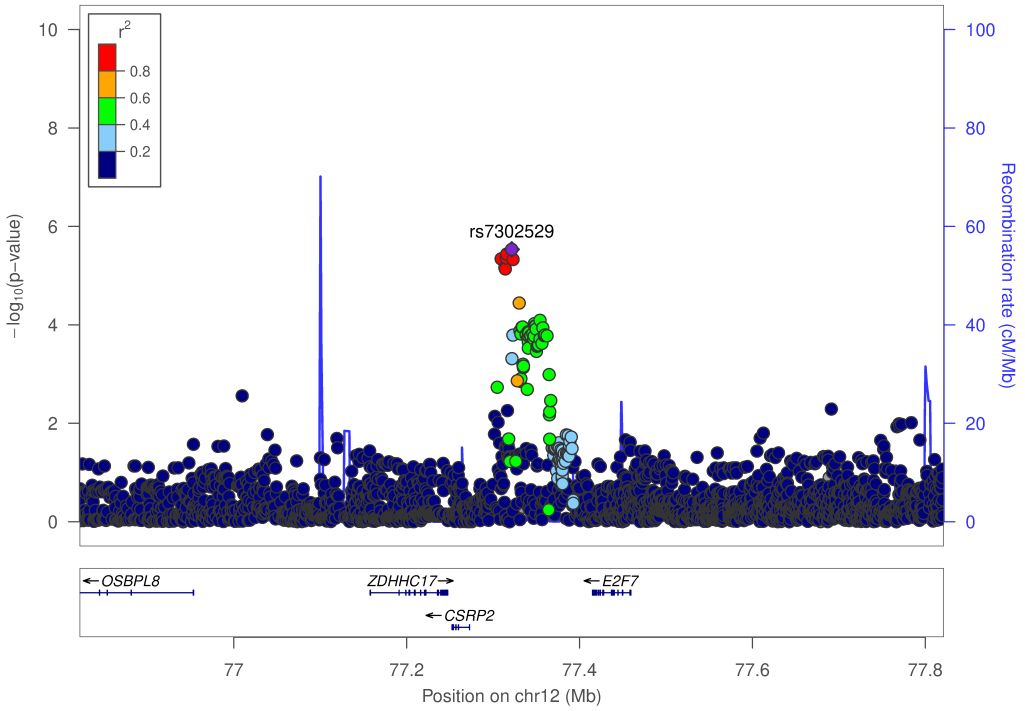

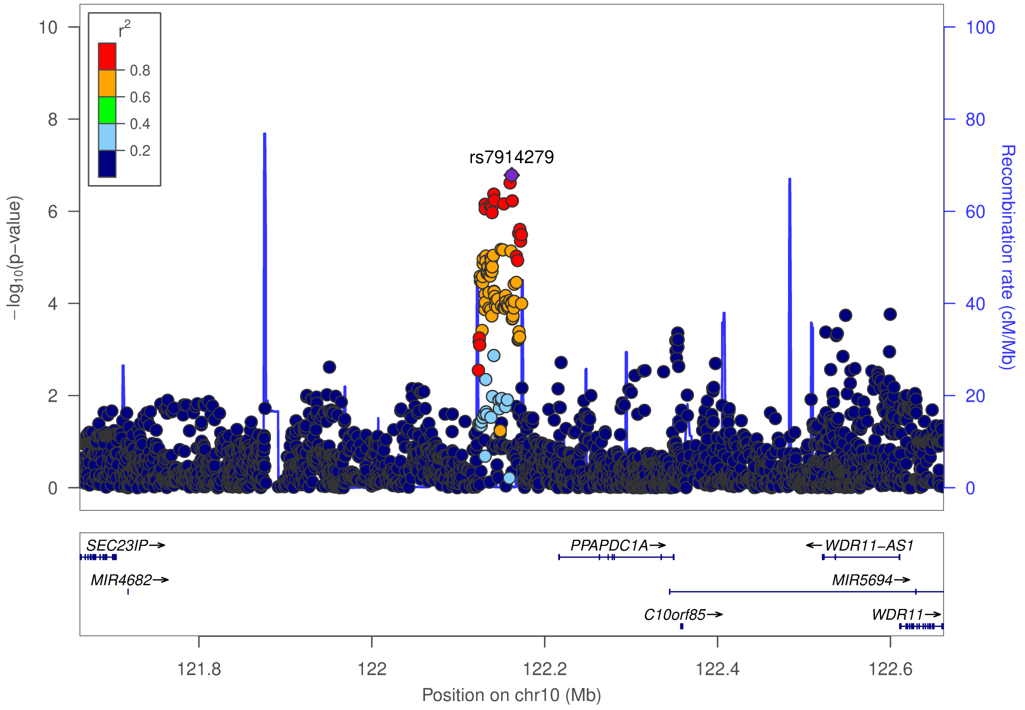


### h) Cross-Disorder SCZ-BIP-rMDD – European ancestry only

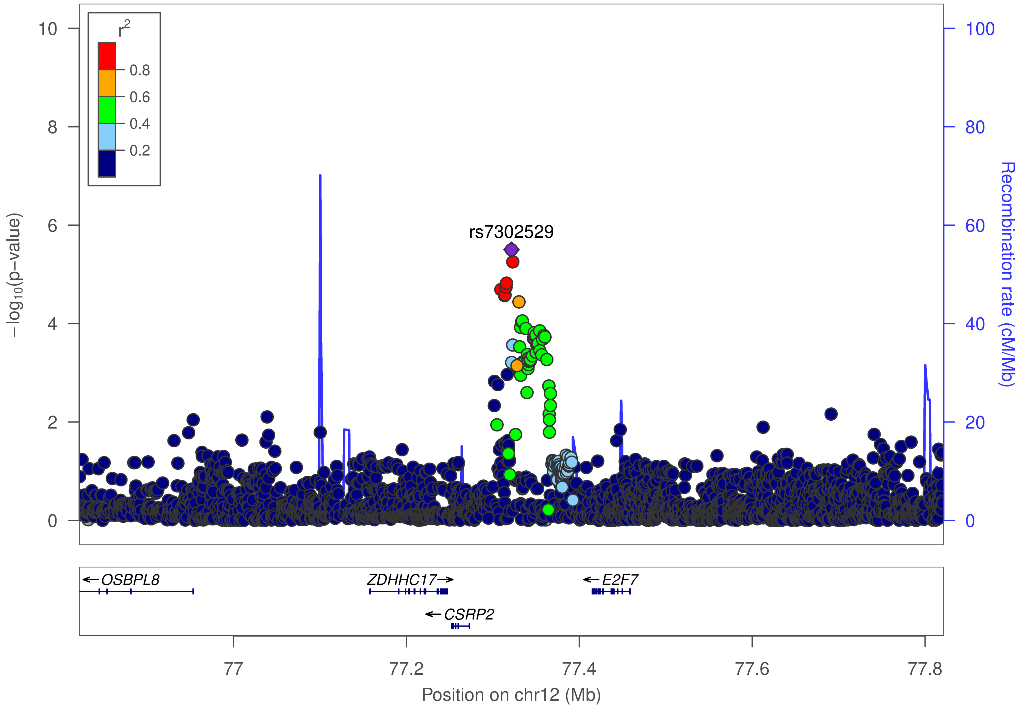

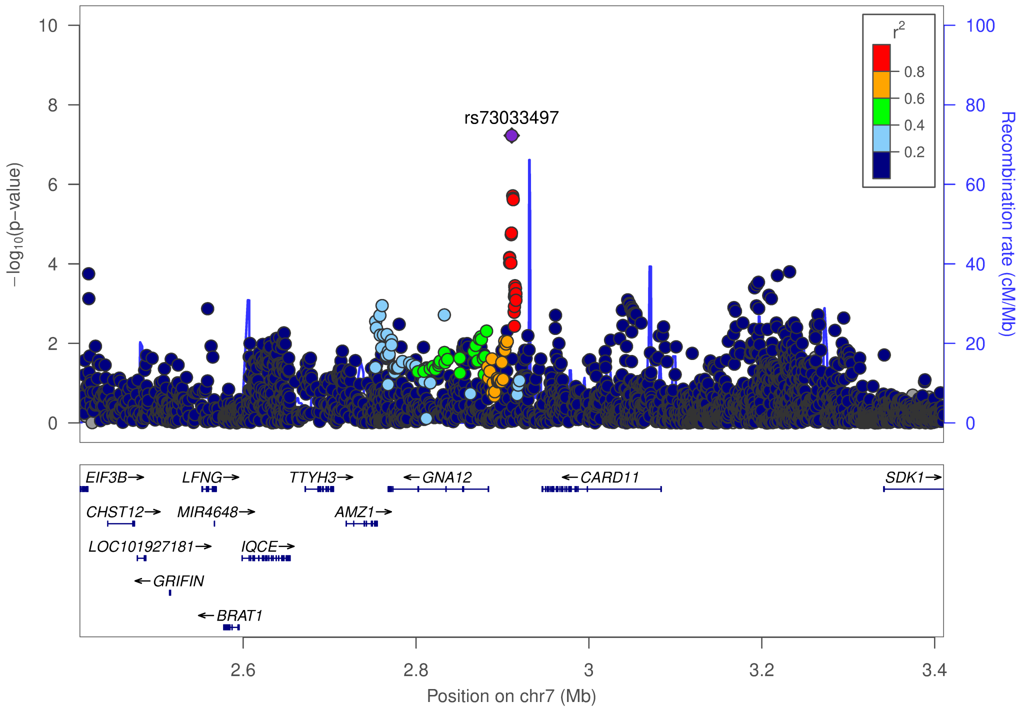


### i) Cross-Disorder SCZ-BIP-rMDD – European + East Asian ancestry

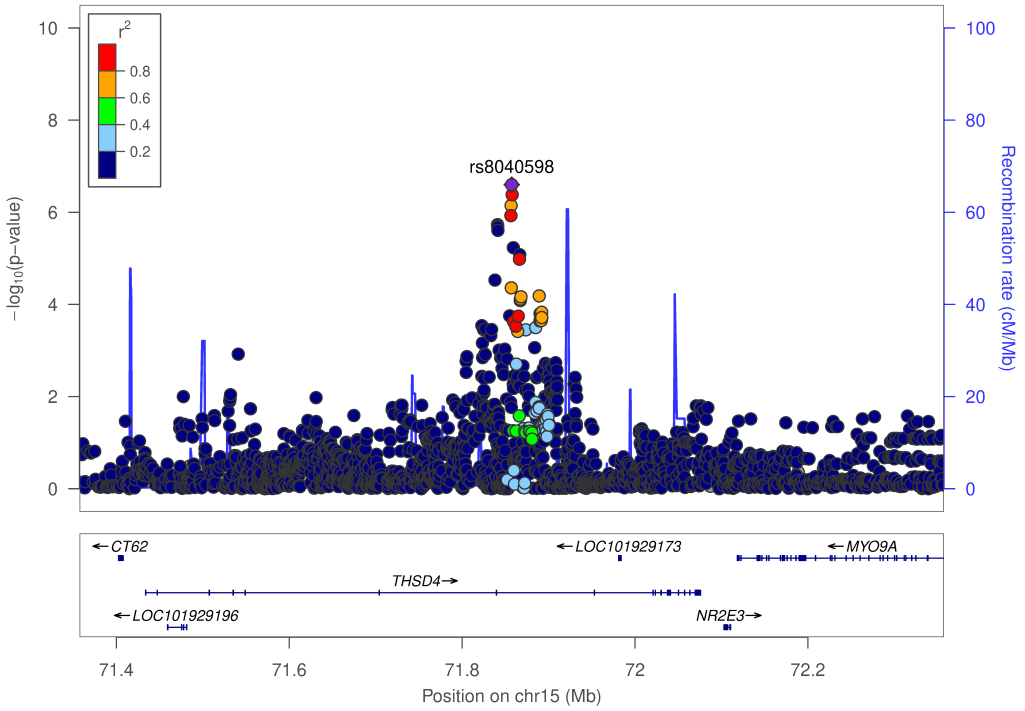

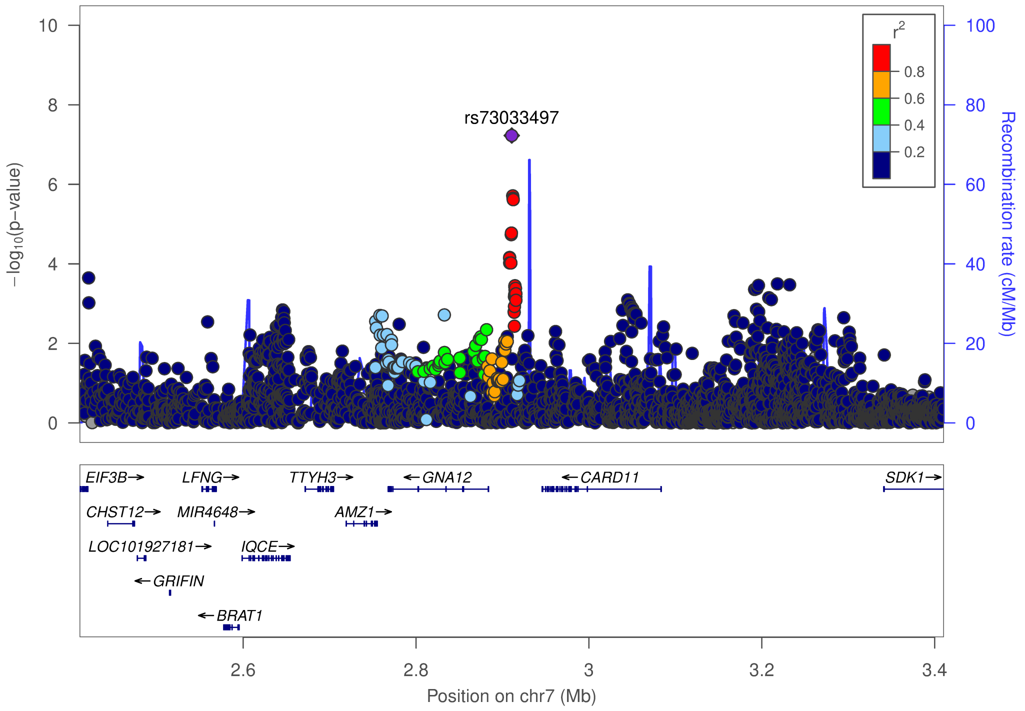


### j) Omnibus Test SCZ-BIP-MDD – European ancestry

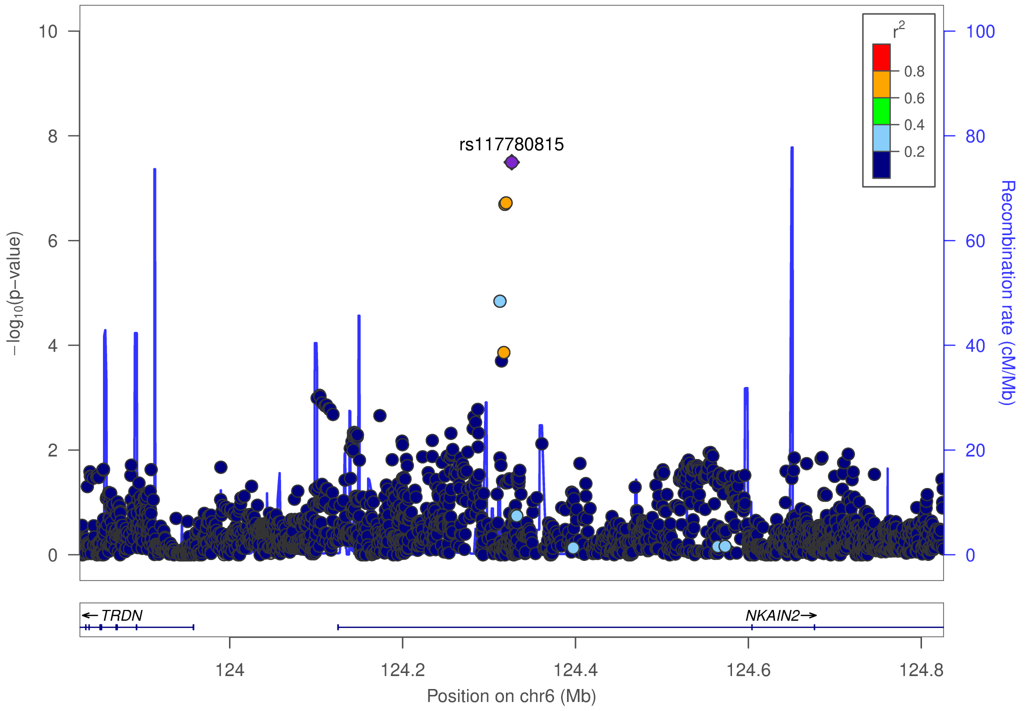

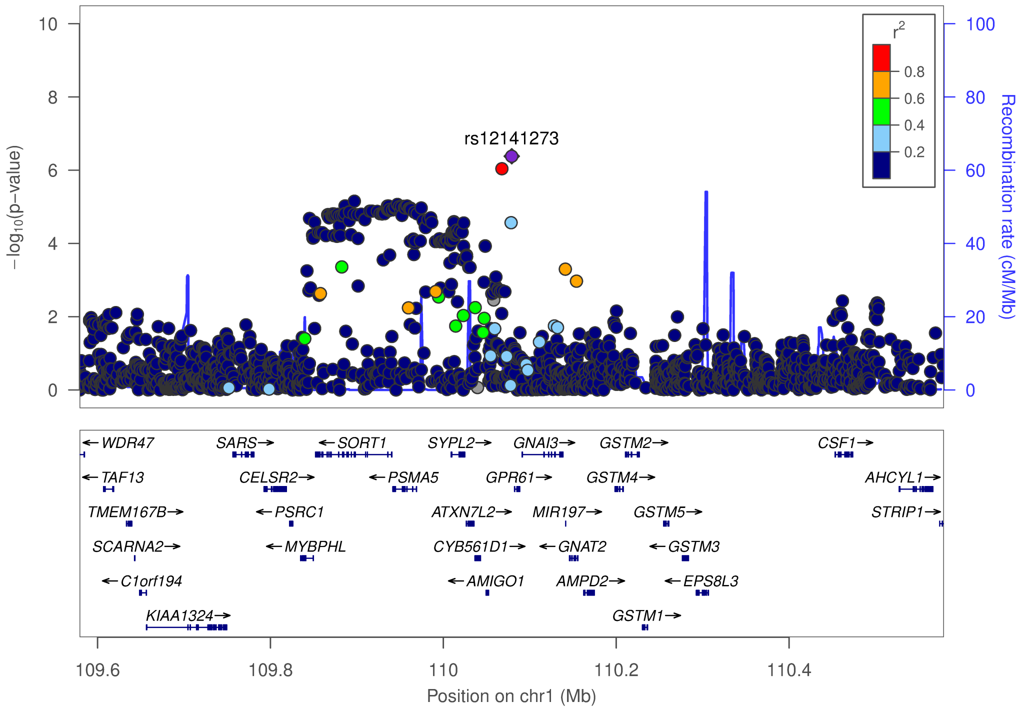


### k) Omnibus Test SCZ-BIP-MDD – European + East Asian ancestry

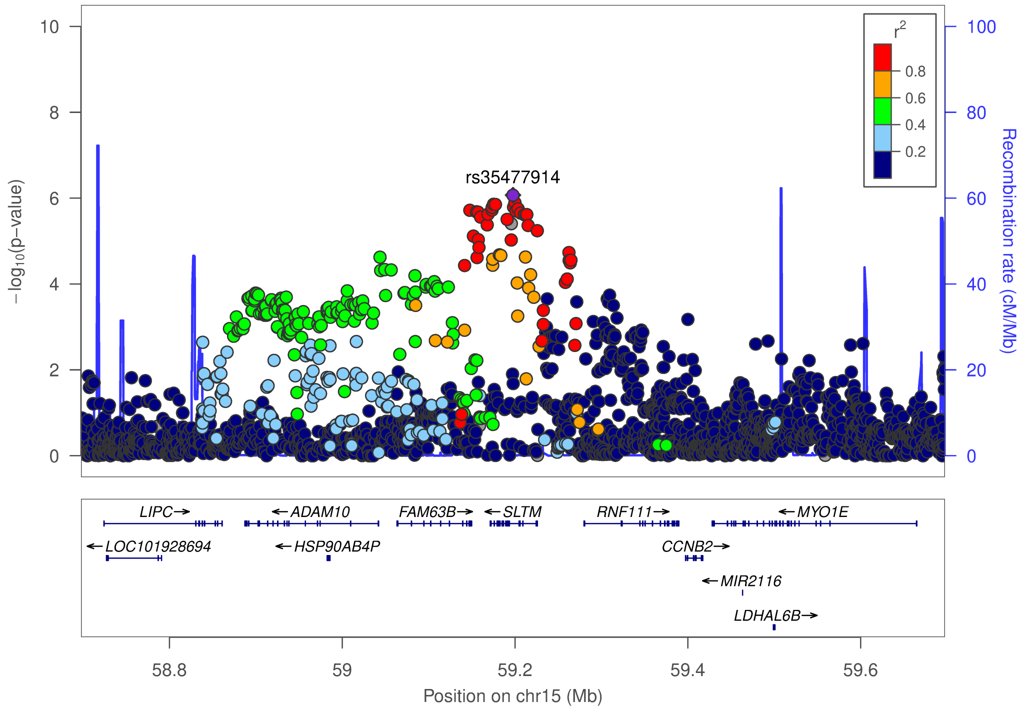

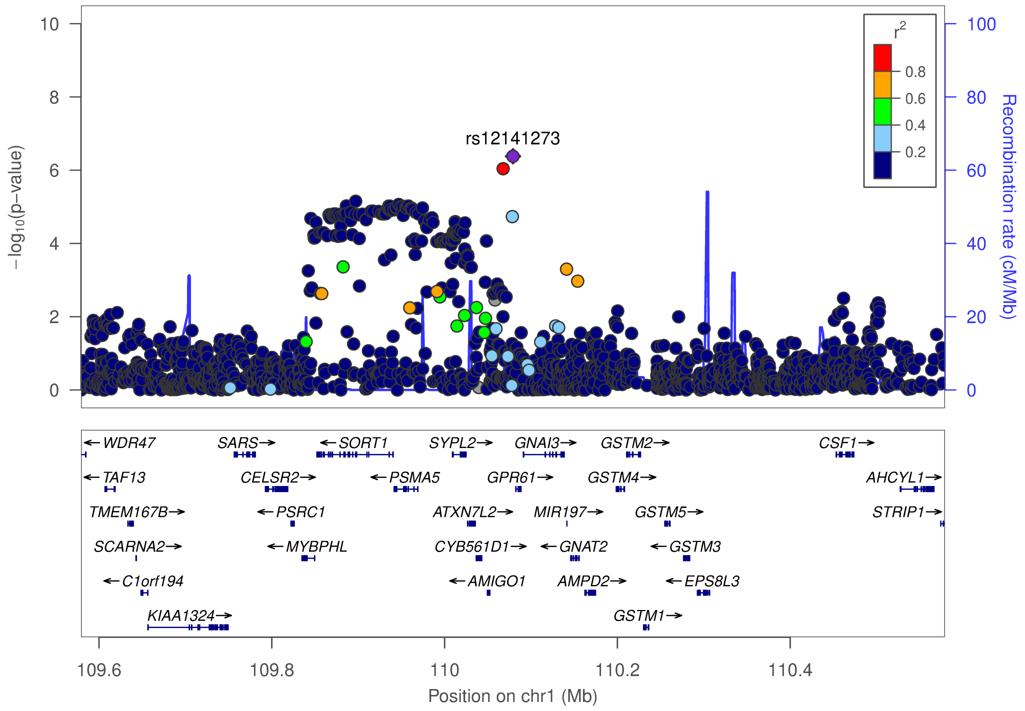
rs12141273
r2
0.8
0.6
0.4
0.2
−log10(p−value)
Recombination rate (cM/Mb)
WDR47
SARS
SORT1
SYPL2
GNAI3
GSTM2
CSF1
TAF13
CELSR2
PSMA5
GPR61
GSTM4
AHCYL1
TMEM167B
PSRC1
ATXN7L2
MIR197
GSTM5
STRIP1
SCARNA2
MYBPHL
CYB561D1
GNAT2
GSTM3
C1orf194
AMIGO1
AMPD2
EPS8L3
KIAA1324
GSTM1
109.6
109.8
110
110.2
110.4
Position on chr1 (Mb)

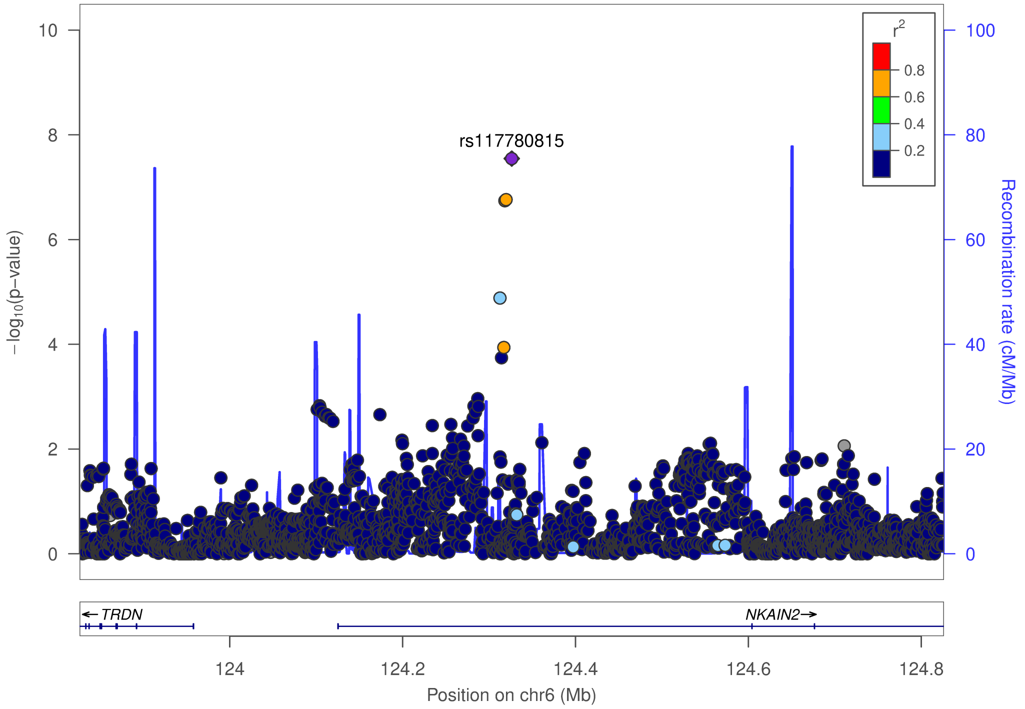
rs117780815
r2
0.8
0.6
0.4
0.2
−log10(p−value)
Recombination rate (cM/Mb)
TRDN
NKAIN2
124
124.2
124.4
124.6
124.8
Position on chr6 (Mb)

### I) Omnibus Test SCZ-BIP-rMDD – European ancestry

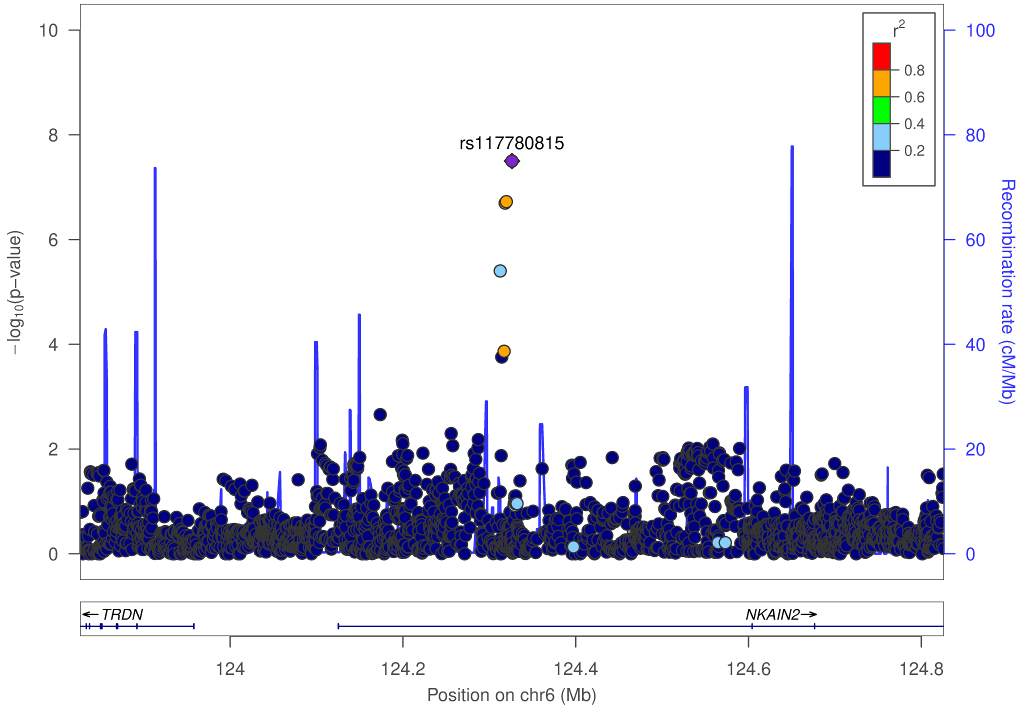


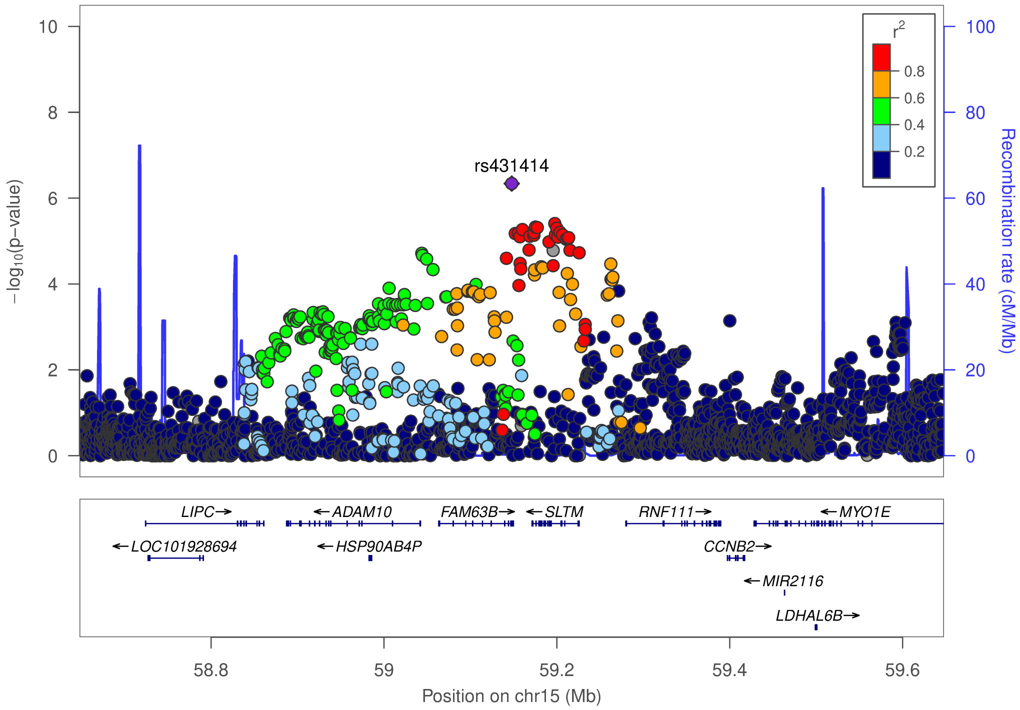

**m) Omnibus Test SCZ-BIP-rMDD – European + East Asian ancestry**

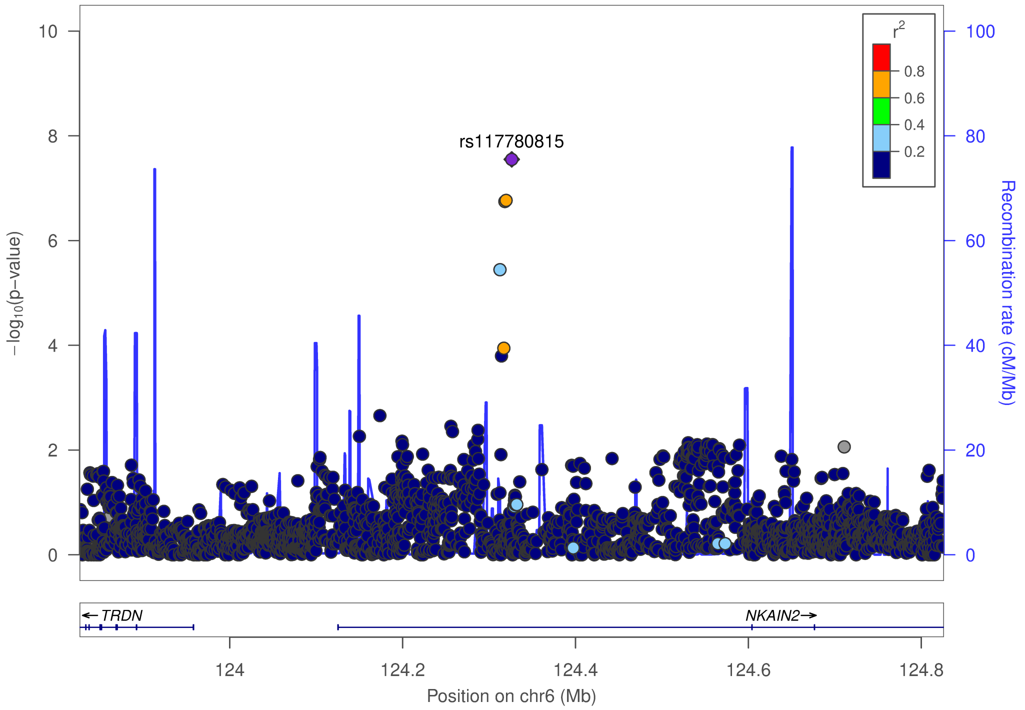

**Supplementary Figure 8. Forest plots for PGC + iPSYCH**

Plots were generated using the rmeta package in R for loci (index SNPs) with GxS interaction $p < 1 \times 10^{-6}$.

Abbreviations: BIP = Bipolar Disorder; (r)MDD = (recurrent) Major Depressive Disorder; SCZ = Schizophrenia; HC F N = number of female healthy controls; HC M N = number of male healthy controls; PT F N = number of female patients; PT M N = number of male patients; Study = cohort abbreviation used by PGC; Meta = meta-analysis results

### a) Schizophrenia – European ancestry only

rs11665282 (A/G)
Schizophrenia

| Study | HC F N | HC M N | PT F N | PT M N | p-value |
|---|---|---|---|---|---|
| aarh | 388 | 458 | 384 | 458 | 1.56E-04 |
| ajsz | 414 | 1170 | 327 | 563 | 7.00E-02 |
| asrb | 153 | 121 | 147 | 279 | 4.02E-01 |
| boco | 621 | 644 | 756 | 952 | 1.20E-01 |
| cims | 22 | 40 | 10 | 54 | 7.70E-01 |
| clm2 | 1965 | 1950 | 914 | 2431 | 1.16E-01 |
| clo3 | 880 | 1013 | 627 | 1433 | 5.74E-02 |
| cou3 | 341 | 329 | 206 | 314 | 9.69E-01 |
| denm | 187 | 258 | 190 | 261 | 3.28E-02 |
| edin | 131 | 144 | 95 | 263 | 1.13E-01 |
| egcu | 821 | 297 | 167 | 62 | 6.00E-01 |
| ersw | 121 | 182 | 92 | 141 | 4.17E-01 |
| fii6 | 592 | 437 | 152 | 204 | 3.81E-01 |
| gras | 410 | 698 | 335 | 688 | 4.15E-02 |
| irwt | 433 | 537 | 422 | 844 | 9.19E-01 |
| lacw | 59 | 60 | 51 | 104 | 3.78E-01 |
| lie2 | 143 | 125 | 37 | 95 | 1.71E-01 |
| lie5 | 200 | 183 | 124 | 364 | 3.36E-01 |
| mgs2 | 1241 | 1058 | 777 | 1760 | 3.46E-02 |
| munc | 153 | 136 | 150 | 267 | 9.14E-01 |
| pews | 95 | 134 | 62 | 87 | 7.68E-02 |
| s234 | 1063 | 1087 | 746 | 1123 | 6.72E-01 |
| swe5 | 1190 | 1306 | 694 | 1033 | 7.87E-02 |
| swe6 | 533 | 549 | 372 | 515 | 7.44E-01 |
| ucla | 297 | 291 | 171 | 519 | 7.40E-02 |
| umeb | 288 | 236 | 136 | 174 | 8.26E-01 |
| **Meta** | **12719** | **13443** | **8134** | **14988** | **1.49E-07** |

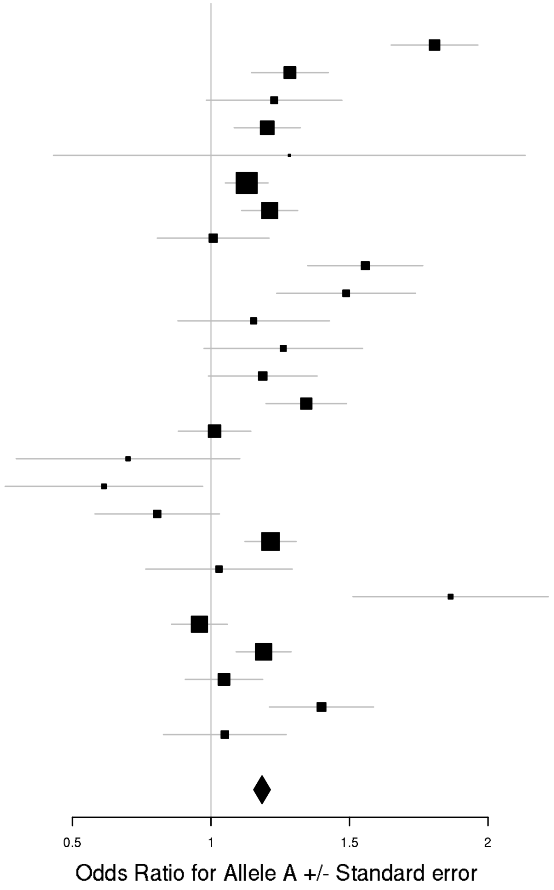

rs12445424 (A/G)
Schizophrenia

| Study | HC F N | HC M N | PT F N | PT M N | p-value |
|---|---|---|---|---|---|
| aarh | 393 | 463 | 387 | 465 | 2.02E-01 |
| aber | 244 | 440 | 182 | 530 | 4.59E-01 |
| ajsz | 414 | 1173 | 327 | 567 | 1.26E-01 |
| asrb | 154 | 122 | 148 | 280 | 2.99E-03 |
| boco | 628 | 650 | 764 | 960 | 6.64E-01 |
| buls | 317 | 288 | 104 | 91 | 2.16E-01 |
| cati | 21 | 90 | 90 | 304 | 8.06E-01 |
| caws | 145 | 125 | 131 | 260 | 4.37E-01 |
| cims | 22 | 40 | 11 | 54 | 8.30E-01 |
| clm2 | 1981 | 1964 | 924 | 2455 | 3.75E-02 |
| clo3 | 886 | 1019 | 631 | 1444 | 5.46E-01 |
| cou3 | 342 | 333 | 208 | 319 | 6.02E-01 |
| denm | 189 | 261 | 193 | 266 | 5.10E-02 |
| dubl | 579 | 237 | 79 | 176 | 8.93E-01 |
| edin | 135 | 146 | 99 | 265 | 6.23E-02 |
| egcu | 828 | 299 | 169 | 62 | 4.70E-01 |
| ersw | 123 | 182 | 93 | 143 | 3.11E-01 |
| fi3m | 466 | 452 | 85 | 101 | 1.24E-01 |
| fii6 | 596 | 438 | 153 | 207 | 9.14E-01 |
| gras | 415 | 708 | 342 | 688 | 4.45E-02 |
| ipsy | 791 | 1645 | 1325 | 1470 | 6.22E-02 |
| irwt | 443 | 549 | 428 | 859 | 1.60E-02 |
| lacw | 59 | 60 | 51 | 105 | 6.12E-01 |
| lie2 | 144 | 125 | 37 | 95 | 7.55E-03 |
| lie5 | 204 | 183 | 126 | 365 | 5.62E-01 |
| mgs2 | 1273 | 1074 | 790 | 1792 | 2.63E-02 |
| msaf | 58 | 80 | 121 | 200 | 8.17E-01 |
| munc | 155 | 136 | 149 | 261 | 3.09E-01 |
| pewb | 907 | 867 | 142 | 398 | 1.34E-01 |
| pews | 96 | 139 | 63 | 87 | 8.13E-01 |
| port | 131 | 78 | 133 | 213 | 2.94E-01 |
| s234 | 1081 | 1108 | 761 | 1141 | 3.36E-01 |
| swe1 | 102 | 101 | 95 | 116 | 4.90E-01 |
| swe5 | 1200 | 1313 | 697 | 1041 | 9.90E-01 |
| swe6 | 537 | 550 | 372 | 518 | 6.40E-01 |
| top8 | 202 | 201 | 162 | 215 | 2.45E-01 |
| ucla | 300 | 295 | 171 | 522 | 3.97E-01 |
| uclo | 281 | 203 | 144 | 358 | 4.94E-01 |
| umeb | 293 | 241 | 139 | 177 | 6.66E-01 |
| umes | 337 | 287 | 80 | 103 | 1.18E-01 |
| zhh1 | 98 | 92 | 63 | 127 | 2.30E-01 |
| **Meta** | **17548** | **18757** | **11158** | **19800** | **3.52E-07** |

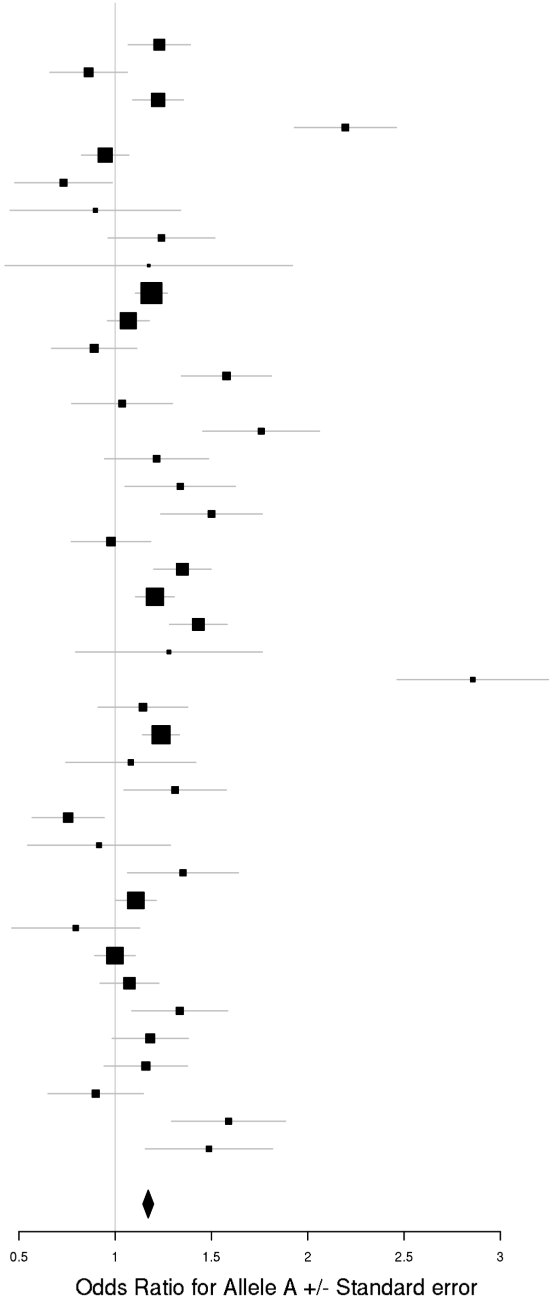

### b) Schizophrenia – European + East Asian ancestry

rs11665282 (A/G)
Schizophrenia

| Study | HC F N | HC M N | PT F N | PT M N | p-value |
|---|---|---|---|---|---|
| aarh | 388 | 458 | 384 | 458 | 1.56E-04 |
| ajsz | 414 | 1170 | 327 | 563 | 7.00E-02 |
| asrb | 153 | 121 | 147 | 279 | 4.02E-01 |
| boco | 621 | 644 | 756 | 952 | 1.20E-01 |
| cims | 22 | 40 | 10 | 54 | 7.70E-01 |
| clm2 | 1965 | 1950 | 914 | 2431 | 1.16E-01 |
| clo3 | 880 | 1013 | 627 | 1433 | 5.74E-02 |
| cou3 | 341 | 329 | 206 | 314 | 9.69E-01 |
| denm | 187 | 258 | 190 | 261 | 3.28E-02 |
| edin | 131 | 144 | 95 | 263 | 1.13E-01 |
| egcu | 821 | 297 | 167 | 62 | 6.00E-01 |
| ersw | 121 | 182 | 92 | 141 | 4.17E-01 |
| fii6 | 592 | 437 | 152 | 204 | 3.81E-01 |
| gras | 410 | 698 | 335 | 688 | 4.15E-02 |
| irwt | 433 | 537 | 422 | 844 | 9.19E-01 |
| lacw | 59 | 60 | 51 | 104 | 3.78E-01 |
| lie2 | 143 | 125 | 37 | 95 | 1.71E-01 |
| lie5 | 200 | 183 | 124 | 364 | 3.36E-01 |
| mgs2 | 1241 | 1058 | 777 | 1760 | 3.46E-02 |
| munc | 153 | 136 | 150 | 267 | 9.14E-01 |
| pews | 95 | 134 | 62 | 87 | 7.68E-02 |
| s234 | 1063 | 1087 | 746 | 1123 | 6.72E-01 |
| swe5 | 1190 | 1306 | 694 | 1033 | 7.87E-02 |
| swe6 | 533 | 549 | 372 | 515 | 7.44E-01 |
| ucla | 297 | 291 | 171 | 519 | 7.40E-02 |
| umeb | 288 | 236 | 136 | 174 | 8.26E-01 |
| jpn1 | 212 | 212 | 231 | 248 | 1.30E-01 |
| tcr1 | 327 | 560 | 285 | 556 | 8.20E-01 |
| **Meta** | **13258** | **14215** | **8650** | **15792** | **3.74E-07** |

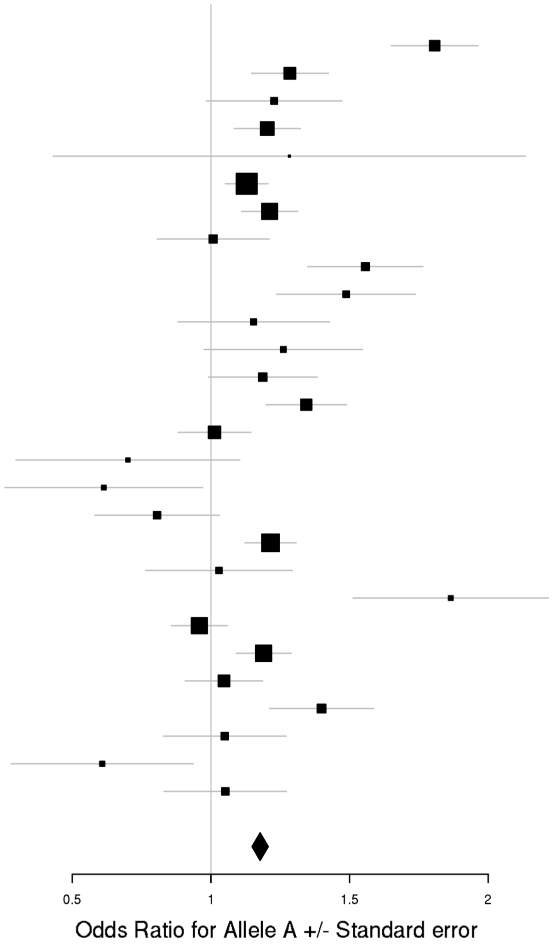

### c) Bipolar Disorder

rs12341335 (T/C)
Bipolar Disorder

| Study | HC F N | HC M N | PT F N | PT M N | p-value |
|---|---|---|---|---|---|
| bmau | 891 | 901 | 189 | 140 | 6.52E-02 |
| bmg2 | 211 | 232 | 117 | 64 | 8.20E-01 |
| bmg3 | 584 | 279 | 286 | 193 | 2.83E-01 |
| bonn | 593 | 607 | 343 | 321 | 1.04E-02 |
| gsk1 | 524 | 369 | 413 | 237 | 5.74E-01 |
| hal2 | 234 | 243 | 234 | 170 | 4.76E-01 |
| may1 | 377 | 377 | 560 | 374 | 3.92E-01 |
| mich | 153 | 135 | 9 | 3 | 9.11E-01 |
| rom3 | 107 | 86 | 134 | 96 | 2.23E-03 |
| swei | 1726 | 1846 | 786 | 470 | 9.50E-03 |
| top8 | 130 | 161 | 81 | 56 | 7.60E-02 |
| ucl2 | 244 | 444 | 397 | 324 | 1.50E-01 |
| ume4 | 291 | 257 | 350 | 211 | 4.39E-02 |
| usc2 | 680 | 472 | 633 | 661 | 8.94E-04 |
| **Meta** | **6745** | **6409** | **4532** | **3320** | **2.29E-07** |

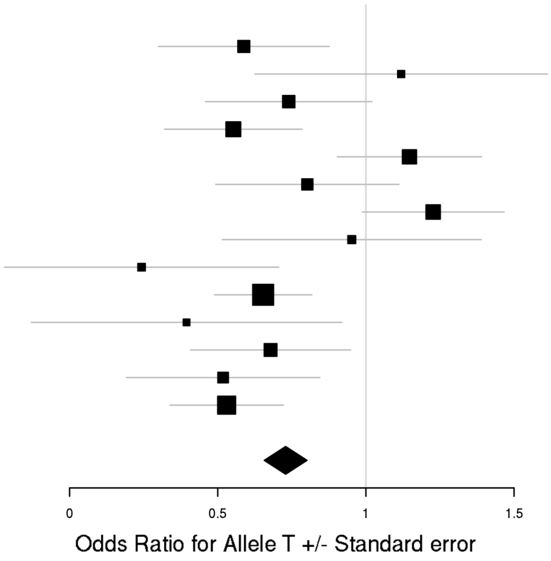

rs17651437 (T/C)
Bipolar Disorder

| Study | HC F N | HC M N | PT F N | PT M N | p-value |
|---|---|---|---|---|---|
| bmau | 889 | 898 | 185 | 140 | 1.48E-01 |
| bmg2 | 210 | 231 | 116 | 63 | 1.20E-01 |
| bmg3 | 587 | 281 | 282 | 194 | 2.78E-01 |
| bmpo | 207 | 471 | 249 | 159 | 5.41E-01 |
| bmsp | 143 | 143 | 137 | 111 | 4.24E-01 |
| bonn | 596 | 605 | 340 | 316 | 7.82E-01 |
| dub1 | 560 | 236 | 78 | 71 | 2.46E-02 |
| edi1 | 133 | 141 | 159 | 119 | 2.22E-01 |
| fat2 | 577 | 480 | 417 | 302 | 7.69E-01 |
| fran | 916 | 684 | 261 | 189 | 3.31E-01 |
| gain | 191 | 206 | 314 | 295 | 9.92E-02 |
| gsk1 | 527 | 369 | 412 | 239 | 7.32E-01 |
| hal2 | 233 | 242 | 231 | 170 | 7.61E-01 |
| icuk | 1661 | 1723 | 1710 | 757 | 5.31E-02 |
| may1 | 375 | 376 | 556 | 374 | 3.74E-01 |
| mich | 156 | 135 | 9 | 3 | 1.26E-01 |
| rom3 | 103 | 87 | 134 | 94 | 5.90E-01 |
| st2c | 551 | 632 | 361 | 272 | 9.17E-01 |
| stp1 | 378 | 406 | 502 | 417 | 1.56E-01 |
| swa2 | 1103 | 1122 | 516 | 361 | 1.53E-02 |
| swei | 1693 | 1820 | 779 | 460 | 1.58E-01 |
| top7 | 188 | 184 | 251 | 185 | 1.61E-01 |
| top8 | 130 | 159 | 78 | 55 | 5.23E-01 |
| ucl2 | 242 | 445 | 395 | 319 | 3.15E-01 |
| uclo | 283 | 211 | 242 | 172 | 3.51E-01 |
| wtcc | 499 | 456 | 977 | 576 | 5.24E-03 |
| **Meta** | **13131** | **12743** | **9691** | **6413** | **3.72E-07** |

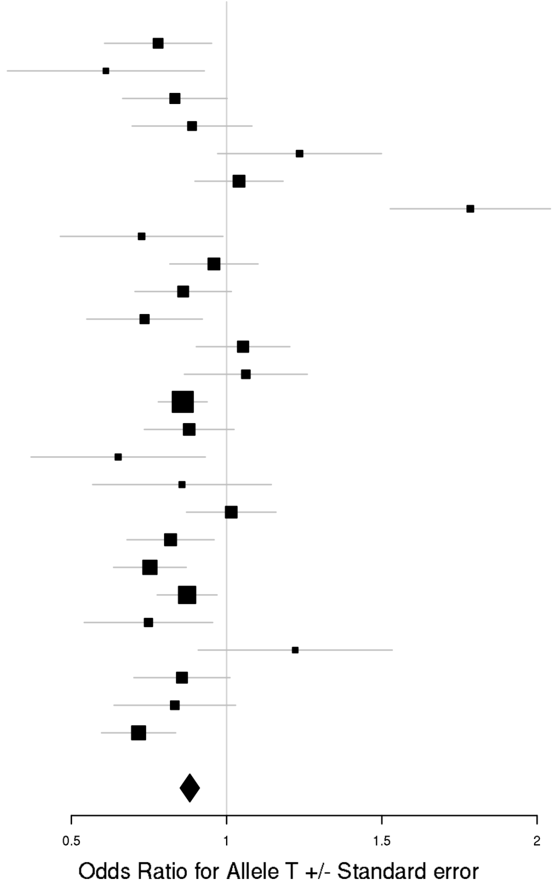

### d) Major Depressive Disorder

rs9428240 (T/C)
Major Depressive Disorder

| Study | HC F N | HC M N | PT F N | PT M N | p-value |
|---|---|---|---|---|---|
| boma | 530 | 536 | 377 | 210 | 4.96E-01 |
| col3 | 605 | 818 | 358 | 148 | 1.14E-02 |
| edi2 | 138 | 146 | 221 | 151 | 1.81E-01 |
| gens | 535 | 784 | 713 | 291 | 2.53E-01 |
| gep3 | 1346 | 1351 | 329 | 141 | 7.98E-01 |
| grdg | 292 | 176 | 365 | 106 | 6.33E-01 |
| grnd | 232 | 241 | 683 | 143 | 8.15E-01 |
| gsk2 | 577 | 276 | 563 | 284 | 1.93E-02 |
| i2b3 | 531 | 534 | 532 | 266 | 2.21E-01 |
| mmi2 | 277 | 227 | 314 | 267 | 6.01E-01 |
| mmo4 | 185 | 193 | 136 | 130 | 9.15E-01 |
| nes1 | 976 | 627 | 1021 | 471 | 1.06E-03 |
| qi6c | 364 | 223 | 346 | 151 | 8.67E-01 |
| rad3 | 820 | 557 | 1315 | 550 | 1.17E-03 |
| rage | 103 | 113 | 213 | 111 | 4.36E-01 |
| rai2 | 178 | 161 | 90 | 19 | 4.71E-01 |
| rau2 | 195 | 182 | 174 | 48 | 8.03E-01 |
| rde4 | 208 | 306 | 93 | 40 | 6.58E-01 |
| rot4 | 570 | 422 | 181 | 63 | 3.10E-01 |
| shp0 | 493 | 638 | 254 | 119 | 3.62E-03 |
| shpt | 223 | 276 | 119 | 46 | 9.23E-01 |
| stm2 | 422 | 503 | 551 | 368 | 1.06E-01 |
| twg2 | 1182 | 1417 | 773 | 321 | 2.33E-01 |
| **Meta** | **10982** | **10707** | **9721** | **4444** | **1.64E-07** |

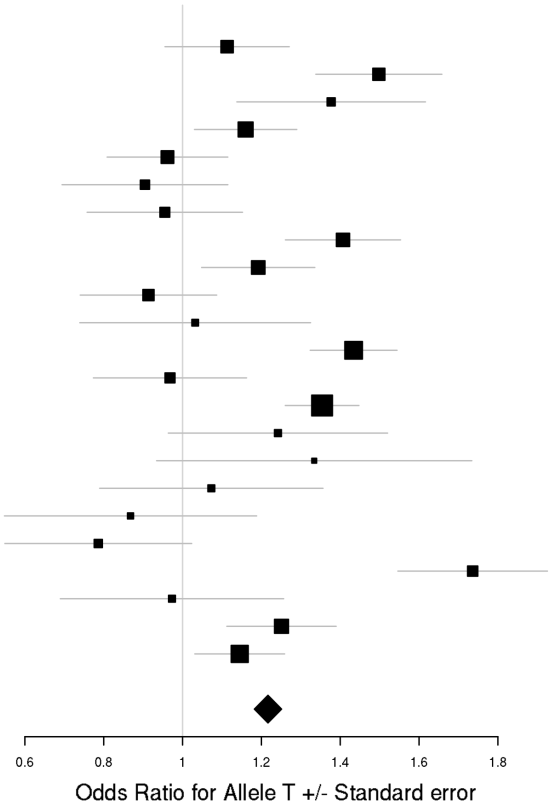

rs147515485 (T/C)
Major Depressive Disorder

| Study | HC F N | HC M N | PT F N | PT M N | p-value |
|---|---|---|---|---|---|
| boma | 531 | 531 | 376 | 210 | 1.15E-01 |
| col3 | 603 | 807 | 355 | 148 | 5.30E-01 |
| edi2 | 137 | 146 | 220 | 151 | 1.63E-01 |
| gens | 530 | 777 | 706 | 291 | 9.76E-01 |
| gep3 | 1337 | 1341 | 324 | 141 | 1.69E-01 |
| grdg | 292 | 173 | 361 | 104 | 7.74E-02 |
| grnd | 230 | 238 | 681 | 142 | 9.00E-01 |
| gsk2 | 574 | 277 | 559 | 285 | 1.92E-01 |
| i2b3 | 527 | 529 | 532 | 264 | 6.83E-01 |
| ipsy | 6512 | 6512 | 5048 | 5048 | 1.99E-05 |
| mmo4 | 186 | 192 | 136 | 130 | 5.87E-01 |
| nes1 | 964 | 619 | 1009 | 468 | 2.49E-02 |
| qi3c | 358 | 207 | 494 | 357 | 5.71E-01 |
| qi6c | 363 | 221 | 346 | 152 | 2.07E-01 |
| qio2 | 303 | 222 | 399 | 160 | 2.35E-01 |
| rad3 | 811 | 557 | 1306 | 546 | 7.41E-01 |
| rage | 102 | 114 | 213 | 110 | 4.12E-01 |
| rau2 | 192 | 181 | 174 | 48 | 8.49E-01 |
| rde4 | 204 | 305 | 90 | 40 | 7.72E-01 |
| rot4 | 569 | 418 | 181 | 62 | 2.61E-02 |
| shp0 | 486 | 632 | 253 | 118 | 2.78E-01 |
| shpt | 224 | 275 | 119 | 46 | 1.11E-01 |
| stm2 | 421 | 496 | 543 | 363 | 7.78E-01 |
| twg2 | 1179 | 1417 | 773 | 319 | 5.65E-03 |
| **Meta** | **17813** | **17187** | **15288** | **9703** | **4.61E-07** |

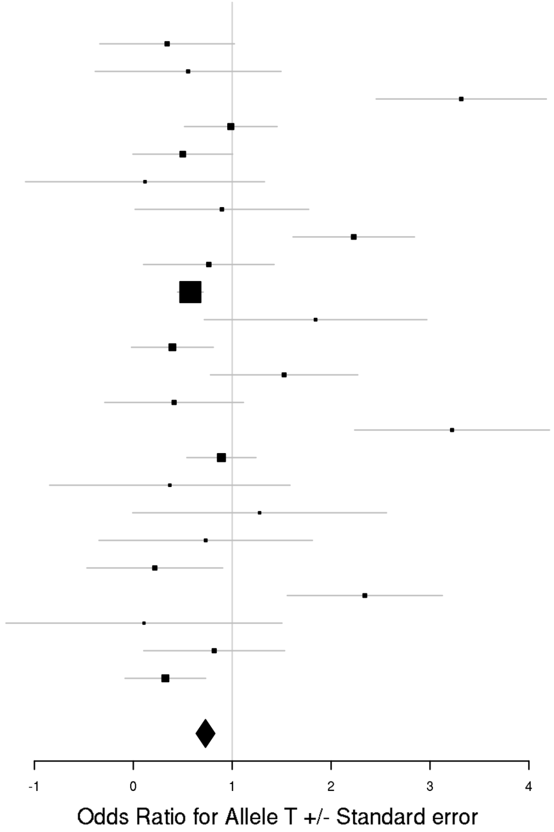

### e) Recurrent Major Depressive Disorder

chr1_118832069_D (D/I2)
Recurrent Major Depressive Disorder

| Study | HC F N | HC M N | PT F N | PT M N | p-value |
|---|---|---|---|---|---|
| boma | 530 | 536 | 297 | 149 | 4.47E-01 |
| col3 | 604 | 818 | 355 | 146 | 1.09E-02 |
| gens | 535 | 782 | 711 | 291 | 2.22E-01 |
| gep3 | 1345 | 1349 | 260 | 85 | 7.27E-01 |
| grnd | 231 | 238 | 673 | 140 | 6.86E-01 |
| gsk2 | 577 | 275 | 440 | 226 | 1.02E-02 |
| mmi2 | 276 | 227 | 229 | 174 | 8.85E-01 |
| mmo4 | 185 | 193 | 104 | 89 | 6.97E-01 |
| nes1 | 976 | 627 | 482 | 220 | 4.93E-04 |
| qi6c | 364 | 222 | 156 | 84 | 9.28E-01 |
| rad3 | 817 | 556 | 1016 | 419 | 1.19E-03 |
| rage | 103 | 113 | 182 | 75 | 3.21E-01 |
| rai2 | 178 | 161 | 89 | 19 | 5.07E-01 |
| rau2 | 195 | 182 | 175 | 47 | 8.59E-01 |
| rde4 | 208 | 306 | 41 | 18 | 1.34E-01 |
| rot4 | 570 | 422 | 58 | 14 | 8.87E-01 |
| shp0 | 493 | 638 | 128 | 52 | 5.52E-03 |
| **Meta** | **8187** | **7645** | **5396** | **2248** | **1.39E-07** |

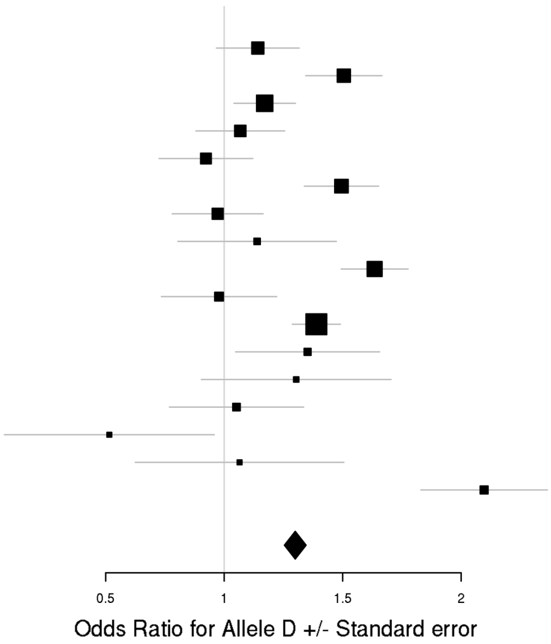

### f) Cross-Disorder SCZ-BIP-MDD – European ancestry only

rs7302529 (T/C)
Cross-Disorder SCZ-BIP-MDD

| Dx | Study | HC F N | HC M N | PT F N | PT M N | p-value |
|---|---|---|---|---|---|---|
| SCZ | aarh | 391 | 463 | 387 | 465 | 9.74E-03 |
| SCZ | ajsz | 414 | 1171 | 327 | 567 | 3.63E-01 |
| SCZ | asrb | 153 | 121 | 148 | 280 | 4.17E-02 |
| SCZ | boco | 160 | 160 | 761 | 952 | 3.97E-01 |
| SCZ | cims | 20 | 33 | 11 | 52 | 5.10E-01 |
| SCZ | clm2 | 671 | 657 | 908 | 2393 | 1.28E-01 |
| SCZ | clo3 | 684 | 818 | 627 | 1435 | 8.16E-02 |
| SCZ | denm | 188 | 259 | 193 | 266 | 2.61E-02 |
| SCZ | egcu | 828 | 299 | 169 | 62 | 1.20E-01 |
| SCZ | ersw | 120 | 175 | 92 | 141 | 9.94E-01 |
| SCZ | fi3m | 469 | 457 | 85 | 101 | 6.67E-01 |
| SCZ | fii6 | 585 | 427 | 152 | 207 | 1.85E-01 |
| SCZ | lacw | 23 | 21 | 51 | 104 | 9.77E-01 |
| SCZ | lie2 | 144 | 124 | 37 | 94 | 7.65E-01 |
| SCZ | lie5 | 204 | 182 | 126 | 363 | 3.83E-01 |
| SCZ | munc | 156 | 136 | 150 | 266 | 4.51E-01 |
| SCZ | swe5 | 585 | 654 | 690 | 1034 | 3.19E-01 |
| SCZ | swe6 | 259 | 283 | 360 | 511 | 9.70E-01 |
| SCZ | ucla | 278 | 282 | 170 | 522 | 9.44E-01 |
| SCZ | umeb | 183 | 146 | 109 | 154 | 2.63E-01 |
| SCZ | umes | 266 | 232 | 72 | 93 | 2.68E-01 |
| BIP | bmau | 895 | 895 | 184 | 140 | 8.66E-01 |
| BIP | bmg2 | 210 | 226 | 117 | 64 | 4.03E-01 |
| BIP | bmg3 | 158 | 54 | 288 | 197 | 7.26E-01 |
| BIP | bmpo | 213 | 476 | 250 | 160 | 9.67E-01 |
| BIP | bmsp | 144 | 147 | 138 | 114 | 1.62E-01 |
| BIP | bonn | 151 | 142 | 345 | 318 | 5.85E-01 |
| BIP | fran | 934 | 693 | 262 | 189 | 8.29E-01 |
| BIP | gsk1 | 408 | 299 | 415 | 239 | 9.25E-01 |
| BIP | icuk | 616 | 671 | 1691 | 741 | 8.58E-01 |
| BIP | may1 | 372 | 369 | 553 | 371 | 6.07E-02 |
| BIP | mich | 69 | 60 | 9 | 3 | 3.86E-01 |
| BIP | swei | 873 | 898 | 765 | 459 | 2.69E-03 |
| BIP | top8 | 126 | 152 | 80 | 55 | 8.16E-01 |
| BIP | ucl2 | 240 | 440 | 393 | 319 | 7.70E-01 |
| BIP | ume4 | 136 | 121 | 294 | 183 | 4.19E-01 |
| BIP | usc2 | 650 | 433 | 632 | 654 | 5.01E-02 |
| MDD | boma | 131 | 149 | 377 | 209 | 5.57E-02 |
| MDD | gep3 | 382 | 361 | 328 | 142 | 8.69E-01 |
| MDD | grdg | 290 | 176 | 365 | 106 | 1.11E-01 |
| MDD | grnd | 199 | 212 | 682 | 143 | 3.73E-01 |
| MDD | gsk2 | 134 | 65 | 561 | 285 | 7.44E-01 |
| MDD | mmi2 | 278 | 228 | 315 | 265 | 5.93E-02 |
| MDD | mmo4 | 181 | 189 | 135 | 129 | 7.39E-01 |
| MDD | rad3 | 694 | 485 | 1303 | 548 | 2.59E-01 |
| MDD | rage | 29 | 34 | 211 | 109 | 1.30E-01 |
| MDD | rai2 | 173 | 157 | 89 | 19 | 1.17E-02 |
| MDD | rau2 | 196 | 182 | 175 | 48 | 4.59E-01 |
| MDD | rot4 | 571 | 422 | 181 | 63 | 6.89E-01 |
| MDD | shpt | 224 | 276 | 119 | 46 | 9.51E-01 |
| MDD | twg2 | 1112 | 1336 | 763 | 317 | 1.71E-01 |
| | **Meta** | **17550** | **17448** | **17634** | **16697** | **1.60E-07** |

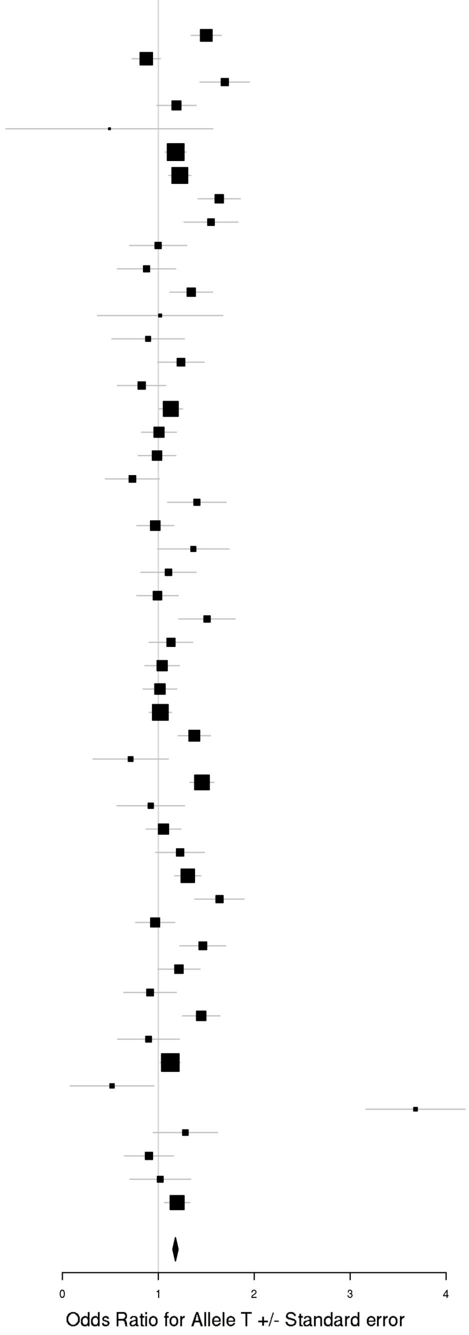

## rs73033497 (A/T)
### Cross-Disorder SCZ-BIP-MDD

| Dx | Study | HC F N | HC M N | PT F N | PT M N | p-value |
|---|---|---|---|---|---|---|
| SCZ | aarh | 383 | 455 | 380 | 454 | 3.47E-01 |
| SCZ | ajsz | 410 | 1155 | 320 | 561 | 6.30E-01 |
| SCZ | buls | 312 | 281 | 101 | 91 | 5.29E-01 |
| SCZ | cims | 20 | 33 | 11 | 52 | 1.69E-01 |
| SCZ | cou3 | 89 | 73 | 200 | 310 | 4.88E-01 |
| SCZ | dubl | 315 | 129 | 78 | 175 | 6.91E-02 |
| SCZ | edin | 37 | 37 | 98 | 262 | 4.76E-01 |
| SCZ | egcu | 815 | 294 | 165 | 61 | 3.88E-01 |
| SCZ | ersw | 120 | 178 | 91 | 140 | 6.08E-01 |
| SCZ | fi3m | 464 | 452 | 83 | 99 | 3.87E-01 |
| SCZ | irwt | 432 | 544 | 424 | 847 | 5.10E-02 |
| SCZ | s234 | 508 | 543 | 732 | 1109 | 1.46E-01 |
| SCZ | swe5 | 580 | 651 | 676 | 1015 | 5.72E-04 |
| SCZ | swe6 | 258 | 280 | 359 | 502 | 2.69E-01 |
| SCZ | top8 | 101 | 101 | 158 | 212 | 1.21E-01 |
| BIP | bmg3 | 154 | 53 | 282 | 194 | 4.10E-01 |
| BIP | dub1 | 280 | 112 | 76 | 68 | 3.99E-01 |
| BIP | gain | 84 | 64 | 311 | 288 | 1.09E-01 |
| BIP | may1 | 373 | 368 | 552 | 366 | 1.53E-01 |
| BIP | swei | 854 | 891 | 757 | 453 | 9.57E-01 |
| BIP | top8 | 127 | 150 | 79 | 55 | 5.81E-01 |
| BIP | ucl2 | 241 | 436 | 387 | 319 | 2.61E-01 |
| MDD | gens | 129 | 193 | 702 | 291 | 3.21E-01 |
| MDD | shp0 | 485 | 623 | 250 | 116 | 2.09E-01 |
| MDD | shpt | 223 | 273 | 118 | 46 | 4.03E-01 |
| MDD | twg2 | 1104 | 1321 | 756 | 309 | 3.76E-01 |
| | **Meta** | **8878** | **9690** | **8135** | **8395** | **8.82E-07** |

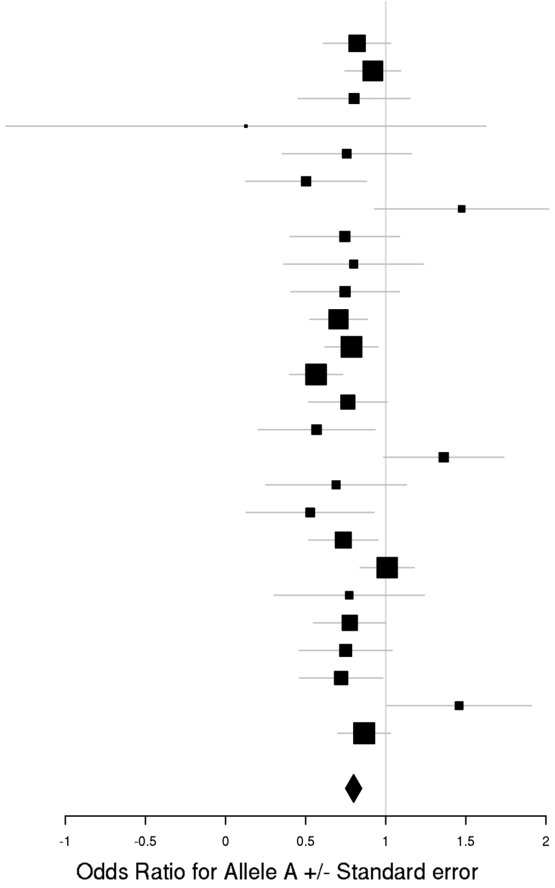

### g) Cross-Disorder SCZ-BIP-MDD – European + East Asian ancestry

rs7914279 (T/G)
Cross-Disorder SCZ-BIP-MDD

| Dx | Study | HC F N | HC M N | PT F N | PT M N | p-value |
|---|---|---|---|---|---|---|
| SCZ | aarh | 389 | 463 | 387 | 464 | 1.07E-01 |
| SCZ | aber | 243 | 438 | 182 | 528 | 1.86E-01 |
| SCZ | ajsz | 414 | 1171 | 327 | 567 | 4.16E-01 |
| SCZ | asrb | 153 | 121 | 147 | 280 | 9.31E-01 |
| SCZ | boco | 160 | 160 | 755 | 951 | 8.98E-01 |
| SCZ | buls | 317 | 288 | 104 | 91 | 3.10E-01 |
| SCZ | cati | 8 | 44 | 90 | 304 | 4.22E-02 |
| SCZ | caws | 65 | 63 | 129 | 256 | 4.80E-01 |
| SCZ | cims | 20 | 33 | 11 | 52 | 5.13E-01 |
| SCZ | clm2 | 679 | 667 | 919 | 2444 | 9.05E-01 |
| SCZ | clo3 | 689 | 820 | 628 | 1433 | 1.23E-01 |
| SCZ | cou3 | 92 | 73 | 202 | 316 | 9.51E-01 |
| SCZ | denm | 188 | 259 | 193 | 266 | 2.74E-01 |
| SCZ | dubl | 319 | 129 | 79 | 176 | 2.74E-01 |
| SCZ | edin | 37 | 37 | 98 | 264 | 7.76E-02 |
| SCZ | egcu | 828 | 299 | 169 | 62 | 1.22E-01 |
| SCZ | ersw | 121 | 178 | 92 | 142 | 4.66E-01 |
| SCZ | fi3m | 458 | 450 | 82 | 101 | 5.11E-01 |
| SCZ | fii6 | 594 | 437 | 153 | 207 | 6.34E-01 |
| SCZ | gras | 411 | 705 | 339 | 688 | 3.85E-01 |
| SCZ | ipsy | 789 | 1639 | 1320 | 1464 | 4.15E-01 |
| SCZ | irwt | 435 | 547 | 425 | 857 | 4.00E-01 |
| SCZ | lacw | 23 | 21 | 51 | 104 | 7.68E-01 |
| SCZ | lie2 | 143 | 124 | 37 | 94 | 1.07E-01 |
| SCZ | lie5 | 203 | 182 | 126 | 361 | 5.38E-01 |
| SCZ | mgs2 | 527 | 381 | 785 | 1783 | 2.52E-02 |
| SCZ | msaf | 57 | 80 | 121 | 200 | 2.76E-02 |
| SCZ | pewb | 886 | 857 | 140 | 396 | 9.65E-01 |
| SCZ | pews | 96 | 138 | 63 | 87 | 7.71E-02 |
| SCZ | port | 131 | 78 | 133 | 212 | 5.05E-01 |
| SCZ | s234 | 518 | 549 | 744 | 1131 | 8.05E-02 |
| SCZ | swe1 | 99 | 97 | 94 | 116 | 8.54E-01 |
| SCZ | swe5 | 583 | 652 | 689 | 1033 | 7.35E-01 |
| SCZ | swe6 | 258 | 283 | 360 | 511 | 4.27E-01 |
| SCZ | top8 | 102 | 102 | 160 | 214 | 1.84E-01 |
| SCZ | ucla | 278 | 281 | 169 | 522 | 4.49E-01 |
| SCZ | uclo | 143 | 95 | 144 | 358 | 6.40E-01 |
| SCZ | umeb | 185 | 150 | 109 | 155 | 8.72E-02 |
| SCZ | umes | 267 | 235 | 72 | 93 | 4.23E-01 |
| SCZ | zhh1 | 97 | 91 | 63 | 127 | 2.30E-01 |
| SCZ | hok2 | 1342 | 666 | 153 | 322 | 1.25E-01 |
| SCZ | jpn1 | 215 | 212 | 238 | 253 | 1.42E-01 |
| SCZ | tcr1 | 336 | 570 | 289 | 562 | 4.47E-01 |
| BIP | bmau | 895 | 894 | 184 | 140 | 4.58E-01 |
| BIP | bmg2 | 210 | 226 | 117 | 64 | 4.57E-01 |
| BIP | bmg3 | 158 | 54 | 288 | 197 | 2.39E-01 |
| BIP | bmsp | 144 | 145 | 137 | 113 | 5.60E-01 |
| BIP | bonn | 150 | 141 | 345 | 318 | 9.62E-01 |
| BIP | dub1 | 283 | 115 | 77 | 70 | 8.89E-01 |
| BIP | edi1 | 43 | 38 | 154 | 118 | 5.90E-01 |
| BIP | fat2 | 254 | 170 | 411 | 301 | 1.59E-01 |
| BIP | gain | 85 | 65 | 314 | 294 | 5.25E-01 |
| BIP | gsk1 | 406 | 299 | 415 | 239 | 3.75E-01 |
| BIP | hal2 | 65 | 73 | 234 | 170 | 3.30E-01 |
| BIP | icuk | 628 | 678 | 1720 | 754 | 6.06E-01 |
| BIP | ipsy | 255 | 295 | 606 | 357 | 7.28E-01 |
| BIP | may1 | 375 | 374 | 558 | 373 | 9.35E-02 |
| BIP | mich | 69 | 60 | 9 | 3 | 6.19E-01 |
| BIP | rom3 | 107 | 87 | 135 | 96 | 4.91E-01 |
| BIP | st2c | 400 | 499 | 361 | 271 | 3.61E-01 |
| BIP | stp1 | 287 | 313 | 502 | 415 | 3.88E-01 |
| BIP | swa2 | 581 | 573 | 499 | 350 | 2.97E-01 |
| BIP | swei | 871 | 897 | 763 | 459 | 3.49E-01 |
| BIP | top7 | 104 | 104 | 250 | 182 | 3.26E-01 |
| BIP | top8 | 128 | 154 | 81 | 56 | 8.19E-01 |
| BIP | ucl2 | 242 | 442 | 394 | 324 | 3.68E-01 |
| BIP | uclo | 139 | 110 | 239 | 170 | 8.52E-01 |
| BIP | ume4 | 139 | 121 | 298 | 184 | 1.40E-01 |
| BIP | usc2 | 657 | 435 | 632 | 655 | 7.30E-01 |
| BIP | wtcc | 204 | 186 | 972 | 573 | 8.75E-01 |
| MDD | boma | 131 | 149 | 376 | 207 | 5.89E-02 |
| MDD | cof3 | 61 | 65 | 74 | 46 | 7.77E-02 |
| MDD | col3 | 608 | 819 | 358 | 148 | 3.93E-01 |
| MDD | edi2 | 36 | 37 | 220 | 151 | 9.82E-01 |
| MDD | gens | 130 | 195 | 713 | 291 | 1.65E-01 |
| MDD | gep3 | 382 | 361 | 328 | 141 | 8.26E-01 |
| MDD | grdg | 290 | 176 | 365 | 106 | 5.63E-01 |
| MDD | grnd | 199 | 211 | 682 | 143 | 3.93E-01 |
| MDD | gsk2 | 134 | 65 | 561 | 284 | 9.23E-01 |
| MDD | i2b3 | 238 | 253 | 530 | 266 | 4.25E-03 |
| MDD | ipsy | 6544 | 6544 | 5075 | 5075 | 9.26E-03 |
| MDD | mmo4 | 185 | 193 | 137 | 130 | 7.15E-01 |
| MDD | nes1 | 951 | 618 | 1020 | 472 | 3.86E-01 |
| MDD | qi6c | 362 | 221 | 346 | 152 | 1.43E-01 |
| MDD | qio2 | 297 | 222 | 403 | 159 | 8.57E-01 |
| MDD | rad3 | 694 | 485 | 1297 | 548 | 1.13E-01 |
| MDD | rage | 29 | 34 | 211 | 108 | 8.33E-01 |
| MDD | rai2 | 172 | 156 | 89 | 19 | 6.07E-01 |
| MDD | rau2 | 196 | 181 | 175 | 48 | 4.86E-01 |
| MDD | rde4 | 208 | 303 | 92 | 40 | 7.90E-02 |
| MDD | rot4 | 570 | 422 | 180 | 63 | 4.59E-01 |
| MDD | shp0 | 494 | 637 | 254 | 119 | 7.96E-01 |
| MDD | shpt | 224 | 276 | 119 | 46 | 1.32E-01 |
| MDD | stm2 | 280 | 357 | 551 | 368 | 8.19E-02 |
| MDD | twg2 | 1111 | 1334 | 757 | 316 | 8.23E-01 |
| | **Meta** | **36283** | **36727** | **37168** | **37239** | **6.39E-07** |

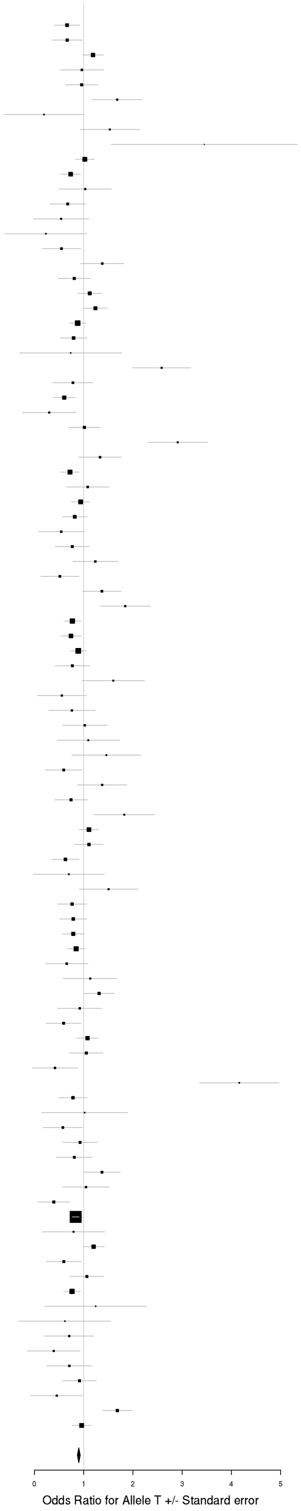

## rs73033497 (A/T)
### Cross-Disorder SCZ-BIP-MDD

| Dx | Study | HC F N | HC M N | PT F N | PT M N | p-value |
|---|---|---|---|---|---|---|
| SCZ | aarh | 383 | 455 | 380 | 454 | 3.47E-01 |
| SCZ | ajsz | 410 | 1155 | 320 | 561 | 6.30E-01 |
| SCZ | buls | 312 | 281 | 101 | 91 | 5.29E-01 |
| SCZ | cims | 20 | 33 | 11 | 52 | 1.69E-01 |
| SCZ | cou3 | 89 | 73 | 200 | 310 | 4.88E-01 |
| SCZ | dubl | 315 | 129 | 78 | 175 | 6.91E-02 |
| SCZ | edin | 37 | 37 | 98 | 262 | 4.76E-01 |
| SCZ | egcu | 815 | 294 | 165 | 61 | 3.88E-01 |
| SCZ | ersw | 120 | 178 | 91 | 140 | 6.08E-01 |
| SCZ | fi3m | 464 | 452 | 83 | 99 | 3.87E-01 |
| SCZ | irwt | 432 | 544 | 424 | 847 | 5.10E-02 |
| SCZ | s234 | 508 | 543 | 732 | 1109 | 1.46E-01 |
| SCZ | swe5 | 580 | 651 | 676 | 1015 | 5.72E-04 |
| SCZ | swe6 | 258 | 280 | 359 | 502 | 2.69E-01 |
| SCZ | top8 | 101 | 101 | 158 | 212 | 1.21E-01 |
| BIP | bmg3 | 154 | 53 | 282 | 194 | 4.10E-01 |
| BIP | dub1 | 280 | 112 | 76 | 68 | 3.99E-01 |
| BIP | gain | 84 | 64 | 311 | 288 | 1.09E-01 |
| BIP | may1 | 373 | 368 | 552 | 366 | 1.53E-01 |
| BIP | swei | 854 | 891 | 757 | 453 | 9.57E-01 |
| BIP | top8 | 127 | 150 | 79 | 55 | 5.81E-01 |
| BIP | ucl2 | 241 | 436 | 387 | 319 | 2.61E-01 |
| MDD | gens | 129 | 193 | 702 | 291 | 3.21E-01 |
| MDD | shp0 | 485 | 623 | 250 | 116 | 2.09E-01 |
| MDD | shpt | 223 | 273 | 118 | 46 | 4.03E-01 |
| MDD | twg2 | 1104 | 1321 | 756 | 309 | 3.76E-01 |
| | **Meta** | **8878** | **9690** | **8135** | **8395** | **8.82E-07** |

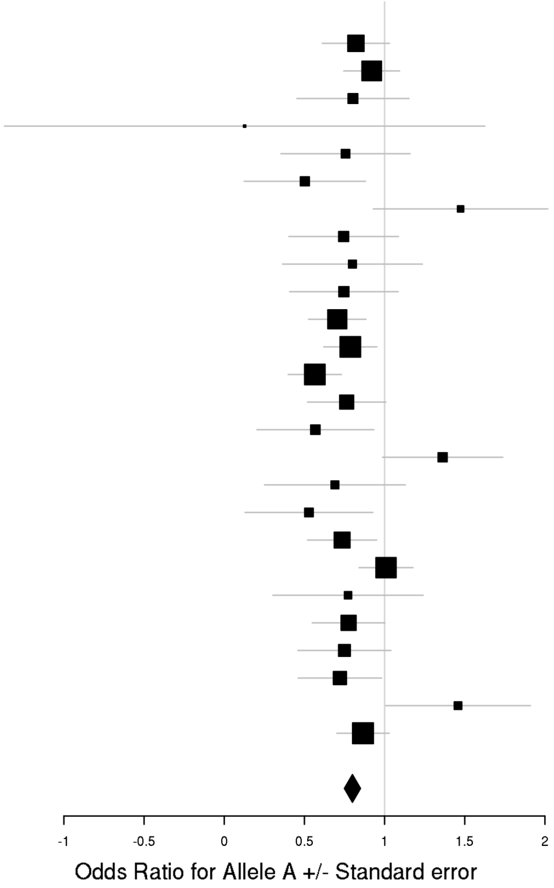

rs7302529 (T/C)
Cross-Disorder SCZ-BIP-MDD

| Dx | Study | HC F N | HC M N | PT F N | PT M N | p-value |
|---|---|---|---|---|---|---|
| SCZ | aarh | 391 | 463 | 387 | 465 | 9.74E-03 |
| SCZ | ajsz | 414 | 1171 | 327 | 567 | 3.63E-01 |
| SCZ | asrb | 153 | 121 | 148 | 280 | 4.17E-02 |
| SCZ | boco | 160 | 160 | 761 | 952 | 3.97E-01 |
| SCZ | cims | 20 | 33 | 11 | 52 | 5.10E-01 |
| SCZ | clm2 | 671 | 657 | 908 | 2393 | 1.28E-01 |
| SCZ | clo3 | 684 | 818 | 627 | 1435 | 8.16E-02 |
| SCZ | denm | 188 | 259 | 193 | 266 | 2.61E-02 |
| SCZ | egcu | 828 | 299 | 169 | 62 | 1.20E-01 |
| SCZ | ersw | 120 | 175 | 92 | 141 | 9.94E-01 |
| SCZ | fi3m | 469 | 457 | 85 | 101 | 6.67E-01 |
| SCZ | fii6 | 585 | 427 | 152 | 207 | 1.85E-01 |
| SCZ | lacw | 23 | 21 | 51 | 104 | 9.77E-01 |
| SCZ | lie2 | 144 | 124 | 37 | 94 | 7.65E-01 |
| SCZ | lie5 | 204 | 182 | 126 | 363 | 3.83E-01 |
| SCZ | munc | 156 | 136 | 150 | 266 | 4.51E-01 |
| SCZ | swe5 | 585 | 654 | 690 | 1034 | 3.19E-01 |
| SCZ | swe6 | 259 | 283 | 360 | 511 | 9.70E-01 |
| SCZ | ucla | 278 | 282 | 170 | 522 | 9.44E-01 |
| SCZ | umeb | 183 | 146 | 109 | 154 | 2.63E-01 |
| SCZ | umes | 266 | 232 | 72 | 93 | 2.68E-01 |
| SCZ | hok2 | 1344 | 668 | 154 | 322 | 2.33E-01 |
| SCZ | tcr1 | 336 | 570 | 289 | 563 | 6.13E-01 |
| BIP | bmau | 895 | 895 | 184 | 140 | 8.66E-01 |
| BIP | bmg2 | 210 | 226 | 117 | 64 | 4.03E-01 |
| BIP | bmg3 | 158 | 54 | 288 | 197 | 7.26E-01 |
| BIP | bmpo | 213 | 476 | 250 | 160 | 9.67E-01 |
| BIP | bmsp | 144 | 147 | 138 | 114 | 1.62E-01 |
| BIP | bonn | 151 | 142 | 345 | 318 | 5.85E-01 |
| BIP | fran | 934 | 693 | 262 | 189 | 8.29E-01 |
| BIP | gsk1 | 408 | 299 | 415 | 239 | 9.25E-01 |
| BIP | icuk | 616 | 671 | 1691 | 741 | 8.58E-01 |
| BIP | may1 | 372 | 369 | 553 | 371 | 6.07E-02 |
| BIP | mich | 69 | 60 | 9 | 3 | 3.86E-01 |
| BIP | swei | 873 | 898 | 765 | 459 | 2.69E-03 |
| BIP | top8 | 126 | 152 | 80 | 55 | 8.16E-01 |
| BIP | ucl2 | 240 | 440 | 393 | 319 | 7.70E-01 |
| BIP | ume4 | 136 | 121 | 294 | 183 | 4.19E-01 |
| BIP | usc2 | 650 | 433 | 632 | 654 | 5.01E-02 |
| MDD | boma | 131 | 149 | 377 | 209 | 5.57E-02 |
| MDD | gep3 | 382 | 361 | 328 | 142 | 8.69E-01 |
| MDD | grdg | 290 | 176 | 365 | 106 | 1.11E-01 |
| MDD | grnd | 199 | 212 | 682 | 143 | 3.73E-01 |
| MDD | gsk2 | 134 | 65 | 561 | 285 | 7.44E-01 |
| MDD | mmi2 | 278 | 228 | 315 | 265 | 5.93E-02 |
| MDD | mmo4 | 181 | 189 | 135 | 129 | 7.39E-01 |
| MDD | rad3 | 694 | 485 | 1303 | 548 | 2.59E-01 |
| MDD | rage | 29 | 34 | 211 | 109 | 1.30E-01 |
| MDD | rai2 | 173 | 157 | 89 | 19 | 1.17E-02 |
| MDD | rau2 | 196 | 182 | 175 | 48 | 4.59E-01 |
| MDD | rot4 | 571 | 422 | 181 | 63 | 6.89E-01 |
| MDD | shpt | 224 | 276 | 119 | 46 | 9.51E-01 |
| MDD | twg2 | 1112 | 1336 | 763 | 317 | 1.71E-01 |
| | **Meta** | **19230** | **18686** | **18077** | **17582** | **9.37E-07** |

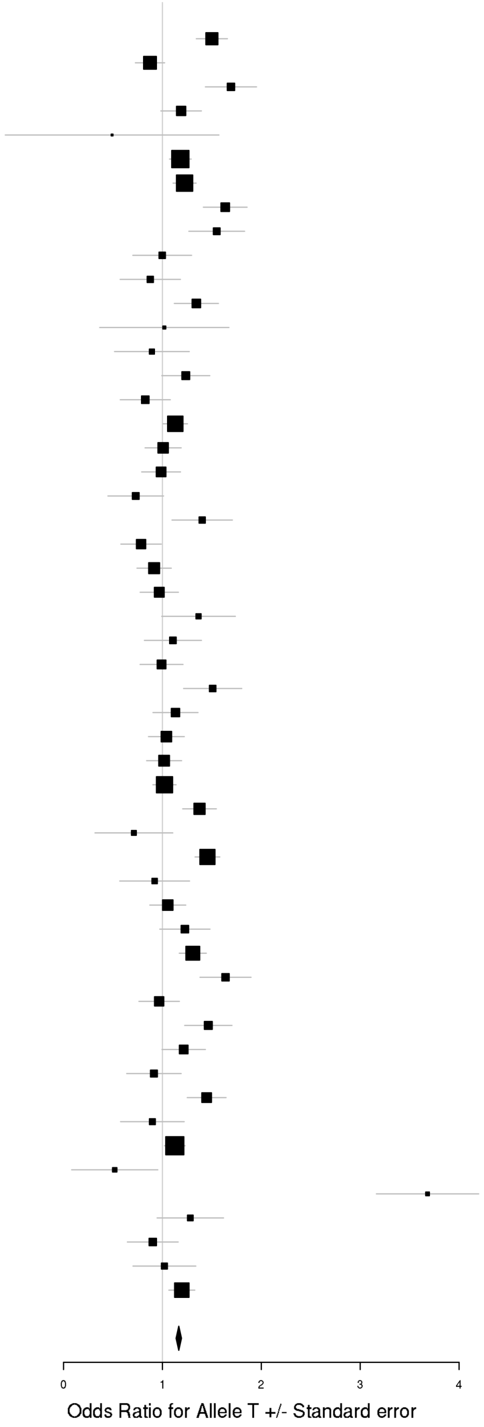

### h) Cross-Disorder SCZ-BIP-rMDD – European ancestry only

rs73033497 (A/T)
Cross-Disorder SCZ-BIP-RMDD

| Dx | Study | HC F N | HC M N | PT F N | PT M N | p-value |
|---|---|---|---|---|---|---|
| SCZ | aarh | 383 | 455 | 380 | 454 | 3.47E-01 |
| SCZ | ajsz | 410 | 1155 | 320 | 561 | 6.30E-01 |
| SCZ | buls | 312 | 281 | 101 | 91 | 5.29E-01 |
| SCZ | cims | 20 | 33 | 11 | 52 | 1.69E-01 |
| SCZ | cou3 | 89 | 73 | 200 | 310 | 4.88E-01 |
| SCZ | dubl | 315 | 129 | 78 | 175 | 6.91E-02 |
| SCZ | edin | 37 | 37 | 98 | 262 | 4.76E-01 |
| SCZ | egcu | 815 | 294 | 165 | 61 | 3.88E-01 |
| SCZ | ersw | 120 | 178 | 91 | 140 | 6.08E-01 |
| SCZ | fi3m | 464 | 452 | 83 | 99 | 3.87E-01 |
| SCZ | irwt | 432 | 544 | 424 | 847 | 5.10E-02 |
| SCZ | s234 | 508 | 543 | 732 | 1109 | 1.46E-01 |
| SCZ | swe5 | 580 | 651 | 676 | 1015 | 5.72E-04 |
| SCZ | swe6 | 258 | 280 | 359 | 502 | 2.69E-01 |
| SCZ | top8 | 101 | 101 | 158 | 212 | 1.21E-01 |
| BIP | bmg3 | 154 | 53 | 282 | 194 | 4.10E-01 |
| BIP | dub1 | 280 | 112 | 76 | 68 | 3.99E-01 |
| BIP | gain | 84 | 64 | 311 | 288 | 1.09E-01 |
| BIP | may1 | 373 | 368 | 552 | 366 | 1.53E-01 |
| BIP | swei | 854 | 891 | 757 | 453 | 9.57E-01 |
| BIP | top8 | 127 | 150 | 79 | 55 | 5.81E-01 |
| BIP | ucl2 | 241 | 436 | 387 | 319 | 2.61E-01 |
| rMDD | gens | 129 | 193 | 702 | 291 | 3.21E-01 |
| rMDD | shp0 | 485 | 623 | 125 | 51 | 2.25E-01 |
| | **Meta** | **7551** | **8096** | **7136** | **7975** | **6.22E-07** |

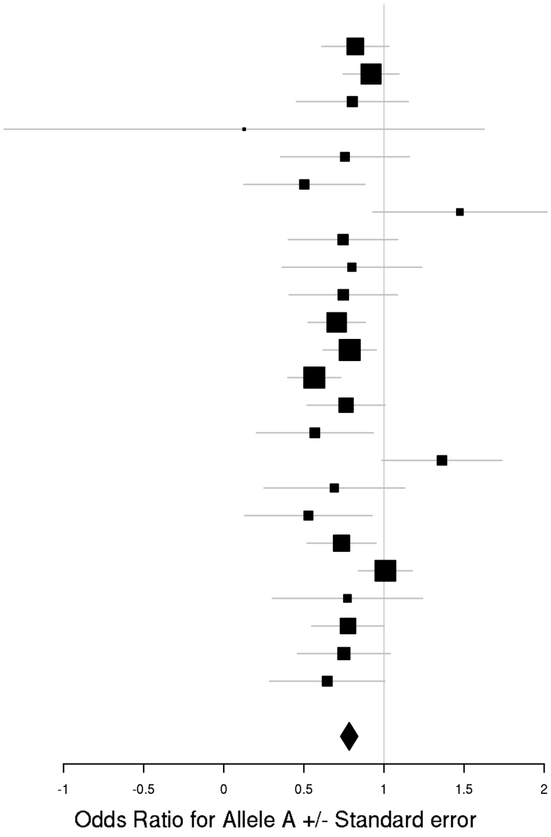

## rs7302529 (T/C)
### Cross-Disorder SCZ-BIP-RMDD

| Dx | Study | HC F N | HC M N | PT F N | PT M N | p-value |
|---|---|---|---|---|---|---|
| SCZ | aarh | 391 | 463 | 387 | 465 | 9.74E-03 |
| SCZ | ajsz | 414 | 1171 | 327 | 567 | 3.63E-01 |
| SCZ | asrb | 153 | 121 | 148 | 280 | 4.17E-02 |
| SCZ | boco | 160 | 160 | 761 | 952 | 3.97E-01 |
| SCZ | cims | 20 | 33 | 11 | 52 | 5.10E-01 |
| SCZ | clm2 | 671 | 657 | 908 | 2393 | 1.28E-01 |
| SCZ | clo3 | 684 | 818 | 627 | 1435 | 8.16E-02 |
| SCZ | denm | 188 | 259 | 193 | 266 | 2.61E-02 |
| SCZ | egcu | 828 | 299 | 169 | 62 | 1.20E-01 |
| SCZ | ersw | 120 | 175 | 92 | 141 | 9.94E-01 |
| SCZ | fi3m | 469 | 457 | 85 | 101 | 6.67E-01 |
| SCZ | fii6 | 585 | 427 | 152 | 207 | 1.85E-01 |
| SCZ | lacw | 23 | 21 | 51 | 104 | 9.77E-01 |
| SCZ | lie2 | 144 | 124 | 37 | 94 | 7.65E-01 |
| SCZ | lie5 | 204 | 182 | 126 | 363 | 3.83E-01 |
| SCZ | munc | 156 | 136 | 150 | 266 | 4.51E-01 |
| SCZ | swe5 | 585 | 654 | 690 | 1034 | 3.19E-01 |
| SCZ | swe6 | 259 | 283 | 360 | 511 | 9.70E-01 |
| SCZ | ucla | 278 | 282 | 170 | 522 | 9.44E-01 |
| SCZ | umeb | 183 | 146 | 109 | 154 | 2.63E-01 |
| SCZ | umes | 266 | 232 | 72 | 93 | 2.68E-01 |
| BIP | bmau | 895 | 895 | 184 | 140 | 8.66E-01 |
| BIP | bmg2 | 210 | 226 | 117 | 64 | 4.03E-01 |
| BIP | bmg3 | 158 | 54 | 288 | 197 | 7.26E-01 |
| BIP | bmpo | 213 | 476 | 250 | 160 | 9.67E-01 |
| BIP | bmsp | 144 | 147 | 138 | 114 | 1.62E-01 |
| BIP | bonn | 151 | 142 | 345 | 318 | 5.85E-01 |
| BIP | fran | 934 | 693 | 262 | 189 | 8.29E-01 |
| BIP | gsk1 | 408 | 299 | 415 | 239 | 9.25E-01 |
| BIP | icuk | 616 | 671 | 1691 | 741 | 8.58E-01 |
| BIP | may1 | 372 | 369 | 553 | 371 | 6.07E-02 |
| BIP | mich | 69 | 60 | 9 | 3 | 3.86E-01 |
| BIP | swei | 873 | 898 | 765 | 459 | 2.69E-03 |
| BIP | top8 | 126 | 152 | 80 | 55 | 8.16E-01 |
| BIP | ucl2 | 240 | 440 | 393 | 319 | 7.70E-01 |
| BIP | ume4 | 136 | 121 | 294 | 183 | 4.19E-01 |
| BIP | usc2 | 650 | 433 | 632 | 654 | 5.01E-02 |
| rMDD | boma | 131 | 149 | 299 | 148 | 1.20E-01 |
| rMDD | gep3 | 382 | 361 | 260 | 85 | 8.82E-01 |
| rMDD | grnd | 199 | 212 | 677 | 140 | 3.29E-01 |
| rMDD | gsk2 | 134 | 65 | 438 | 227 | 7.43E-01 |
| rMDD | mmi2 | 278 | 228 | 230 | 173 | 2.70E-02 |
| rMDD | mmo4 | 181 | 189 | 104 | 89 | 4.70E-01 |
| rMDD | rad3 | 694 | 485 | 1005 | 417 | 2.52E-01 |
| rMDD | rage | 29 | 34 | 180 | 73 | 2.86E-01 |
| rMDD | rai2 | 173 | 157 | 88 | 19 | 1.05E-02 |
| rMDD | rau2 | 196 | 182 | 175 | 47 | 5.17E-01 |
| rMDD | rot4 | 571 | 422 | 57 | 14 | 2.25E-01 |
| | **Meta** | **15924** | **15660** | **15543** | **15700** | **7.43E-07** |

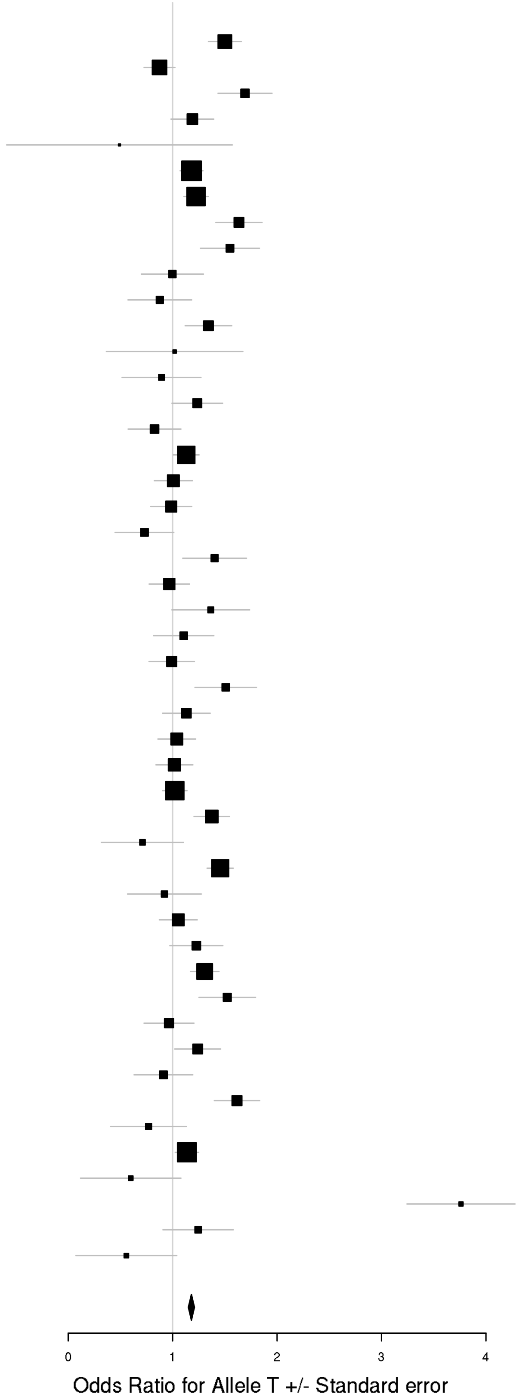

### i) Cross-Disorder SCZ-BIP-rMDD – European + East Asian ancestry

rs8040598 (A/G)
Cross-Disorder SCZ-BIP-RMDD

| Dx | Study | HC F N | HC M N | PT F N | PT M N | p-value |
|---|---|---|---|---|---|---|
| SCZ | aarh | 391 | 463 | 387 | 465 | 1.79E-02 |
| SCZ | ajsz | 414 | 1171 | 327 | 567 | 9.91E-01 |
| SCZ | asrb | 153 | 121 | 148 | 280 | 5.70E-01 |
| SCZ | boco | 160 | 160 | 761 | 952 | 8.52E-01 |
| SCZ | buls | 315 | 288 | 104 | 91 | 2.83E-01 |
| SCZ | cims | 20 | 33 | 11 | 52 | 3.20E-01 |
| SCZ | clm2 | 681 | 668 | 923 | 2450 | 7.24E-01 |
| SCZ | clo3 | 692 | 824 | 629 | 1442 | 7.56E-01 |
| SCZ | denm | 188 | 259 | 193 | 266 | 2.63E-01 |
| SCZ | dubl | 318 | 129 | 79 | 176 | 4.90E-01 |
| SCZ | edin | 37 | 37 | 98 | 264 | 8.31E-01 |
| SCZ | egcu | 828 | 299 | 169 | 62 | 6.33E-01 |
| SCZ | ersw | 121 | 178 | 92 | 142 | 4.48E-02 |
| SCZ | fi3m | 469 | 457 | 85 | 101 | 7.04E-01 |
| SCZ | fii6 | 594 | 437 | 153 | 207 | 9.44E-02 |
| SCZ | gras | 415 | 707 | 342 | 693 | 2.76E-01 |
| SCZ | irwt | 434 | 546 | 424 | 855 | 3.93E-01 |
| SCZ | lacw | 23 | 21 | 51 | 104 | 8.93E-01 |
| SCZ | lie2 | 144 | 124 | 37 | 94 | 8.01E-02 |
| SCZ | lie5 | 204 | 182 | 126 | 363 | 1.15E-01 |
| SCZ | mgs2 | 527 | 381 | 783 | 1777 | 6.40E-01 |
| SCZ | msaf | 57 | 79 | 121 | 198 | 9.62E-01 |
| SCZ | munc | 156 | 136 | 150 | 266 | 7.86E-01 |
| SCZ | pewb | 885 | 856 | 140 | 396 | 2.76E-01 |
| SCZ | pews | 96 | 138 | 63 | 87 | 9.01E-01 |
| SCZ | s234 | 517 | 548 | 744 | 1131 | 3.86E-02 |
| SCZ | swe5 | 585 | 654 | 690 | 1034 | 7.45E-01 |
| SCZ | swe6 | 259 | 283 | 360 | 511 | 1.69E-01 |
| SCZ | top8 | 102 | 101 | 160 | 214 | 5.05E-01 |
| SCZ | ucla | 278 | 282 | 170 | 522 | 1.12E-01 |
| SCZ | umeb | 185 | 150 | 110 | 156 | 7.40E-01 |
| SCZ | umes | 269 | 235 | 72 | 93 | 1.49E-02 |
| SCZ | hok2 | 1344 | 668 | 154 | 322 | 1.30E-02 |
| SCZ | tcr1 | 336 | 570 | 289 | 563 | 7.50E-02 |
| BIP | bmau | 895 | 895 | 184 | 140 | 3.38E-02 |
| BIP | bmg2 | 210 | 226 | 117 | 64 | 1.41E-01 |
| BIP | bmg3 | 158 | 54 | 288 | 197 | 8.90E-01 |
| BIP | bmpo | 213 | 476 | 250 | 160 | 5.69E-01 |
| BIP | bmsp | 144 | 147 | 138 | 114 | 8.53E-01 |
| BIP | bonn | 151 | 142 | 345 | 318 | 1.58E-01 |
| BIP | dub1 | 281 | 115 | 77 | 70 | 6.79E-01 |
| BIP | fran | 934 | 693 | 262 | 189 | 4.42E-01 |
| BIP | gain | 85 | 65 | 314 | 294 | 9.12E-01 |
| BIP | gsk1 | 408 | 299 | 415 | 239 | 3.77E-01 |
| BIP | may1 | 375 | 374 | 560 | 374 | 8.14E-01 |
| BIP | mich | 69 | 60 | 9 | 3 | 4.69E-01 |
| BIP | rom3 | 107 | 87 | 135 | 96 | 8.33E-01 |
| BIP | swa2 | 581 | 572 | 496 | 350 | 2.06E-01 |
| BIP | swei | 873 | 898 | 765 | 459 | 8.96E-02 |
| BIP | top8 | 128 | 155 | 81 | 56 | 2.13E-01 |
| BIP | ucl2 | 243 | 442 | 397 | 324 | 5.34E-01 |
| BIP | ume4 | 139 | 121 | 299 | 184 | 5.56E-01 |
| BIP | usc2 | 659 | 438 | 635 | 659 | 2.29E-01 |
| rMDD | boma | 131 | 149 | 299 | 148 | 6.11E-01 |
| rMDD | cof3 | 60 | 64 | 53 | 37 | 4.09E-01 |
| rMDD | gens | 130 | 195 | 711 | 290 | 2.29E-01 |
| rMDD | gep3 | 382 | 361 | 260 | 85 | 1.00E+ |
| rMDD | grnd | 199 | 212 | 677 | 140 | 3.53E-01 |
| rMDD | gsk2 | 134 | 65 | 438 | 227 | 7.99E-01 |
| rMDD | mmi2 | 278 | 228 | 230 | 173 | 3.05E-01 |
| rMDD | mmo4 | 185 | 193 | 105 | 89 | 5.09E-01 |
| rMDD | qi3c | 369 | 209 | 326 | 253 | 8.20E-01 |
| rMDD | qi6c | 363 | 221 | 156 | 85 | 3.74E-01 |
| rMDD | rad3 | 694 | 485 | 1005 | 417 | 5.90E-01 |
| rMDD | rage | 29 | 34 | 180 | 73 | 9.60E-01 |
| rMDD | rai2 | 172 | 154 | 87 | 19 | 3.01E-01 |
| rMDD | rau2 | 196 | 182 | 175 | 47 | 3.80E-01 |
| rMDD | rde4 | 208 | 303 | 41 | 18 | 4.18E-01 |
| rMDD | rot4 | 571 | 422 | 57 | 14 | 4.12E-01 |
| rMDD | shp0 | 494 | 637 | 128 | 52 | 8.48E-01 |
| | **Meta** | **23425** | **22558** | **19829** | **23353** | **3.90E-07** |

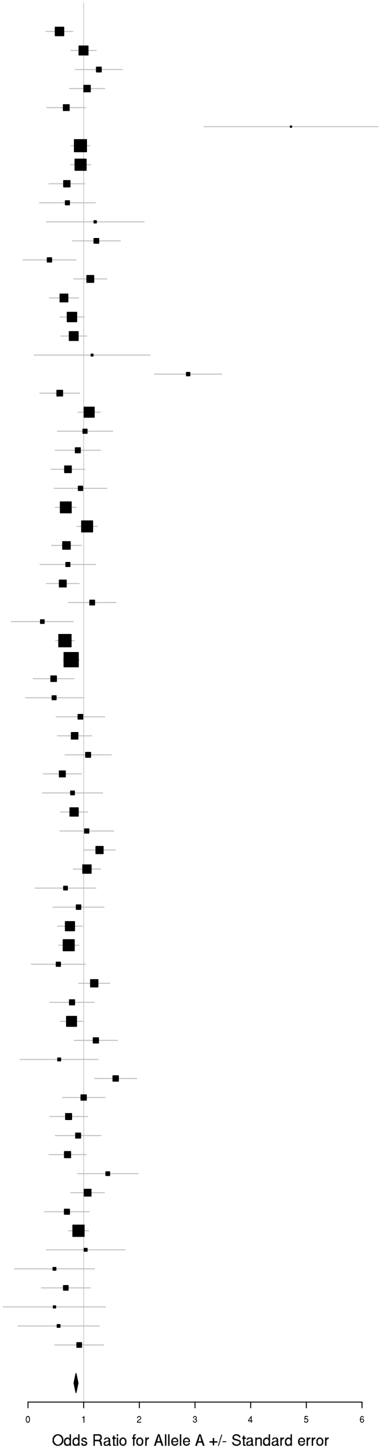

rs73033497 (A/T)
Cross-Disorder SCZ-BIP-RMDD

| Dx | Study | HC F N | HC M N | PT F N | PT M N | p-value |
|---|---|---|---|---|---|---|
| SCZ | aarh | 383 | 455 | 380 | 454 | 3.47E-01 |
| SCZ | ajsz | 410 | 1155 | 320 | 561 | 6.30E-01 |
| SCZ | buls | 312 | 281 | 101 | 91 | 5.29E-01 |
| SCZ | cims | 20 | 33 | 11 | 52 | 1.69E-01 |
| SCZ | cou3 | 89 | 73 | 200 | 310 | 4.88E-01 |
| SCZ | dubl | 315 | 129 | 78 | 175 | 6.91E-02 |
| SCZ | edin | 37 | 37 | 98 | 262 | 4.76E-01 |
| SCZ | egcu | 815 | 294 | 165 | 61 | 3.88E-01 |
| SCZ | ersw | 120 | 178 | 91 | 140 | 6.08E-01 |
| SCZ | fi3m | 464 | 452 | 83 | 99 | 3.87E-01 |
| SCZ | irwt | 432 | 544 | 424 | 847 | 5.10E-02 |
| SCZ | s234 | 508 | 543 | 732 | 1109 | 1.46E-01 |
| SCZ | swe5 | 580 | 651 | 676 | 1015 | 5.72E-04 |
| SCZ | swe6 | 258 | 280 | 359 | 502 | 2.69E-01 |
| SCZ | top8 | 101 | 101 | 158 | 212 | 1.21E-01 |
| BIP | bmg3 | 154 | 53 | 282 | 194 | 4.10E-01 |
| BIP | dub1 | 280 | 112 | 76 | 68 | 3.99E-01 |
| BIP | gain | 84 | 64 | 311 | 288 | 1.09E-01 |
| BIP | may1 | 373 | 368 | 552 | 366 | 1.53E-01 |
| BIP | swei | 854 | 891 | 757 | 453 | 9.57E-01 |
| BIP | top8 | 127 | 150 | 79 | 55 | 5.81E-01 |
| BIP | ucl2 | 241 | 436 | 387 | 319 | 2.61E-01 |
| rMDD | gens | 129 | 193 | 702 | 291 | 3.21E-01 |
| rMDD | shp0 | 485 | 623 | 125 | 51 | 2.25E-01 |
| | **Meta** | **7551** | **8096** | **7136** | **7975** | **6.22E-07** |

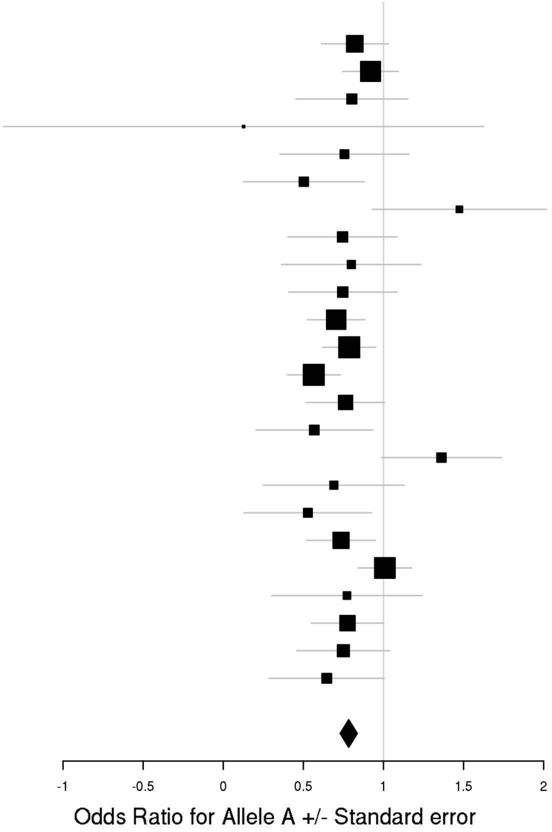

**Supplementary Figure 9. X chromosome model comparisons in PGC + iPSYCH**

GxS interactions with X-linked SNPs were tested using two different models. Model A assumed complete and uniform X-inactivation in females and similar effect size between males and females by assigning 0, 1, or 2 copies of an allele to females and 0 or 2 copies to males. As these assumptions often do not hold, Model B assigned 0 or 1 copy to males.

The scatter plots show substantial correlation (R) between *p*-values from the two X chromosome models, indicating the results from the two models did not differ substantially.

Plots were generated using the plot package in R.

Abbreviations: BIP = Bipolar Disorder; (r)MDD = (recurrent) Major Depressive Disorder; SCZ = Schizophrenia; R2 = proportion variance explained.

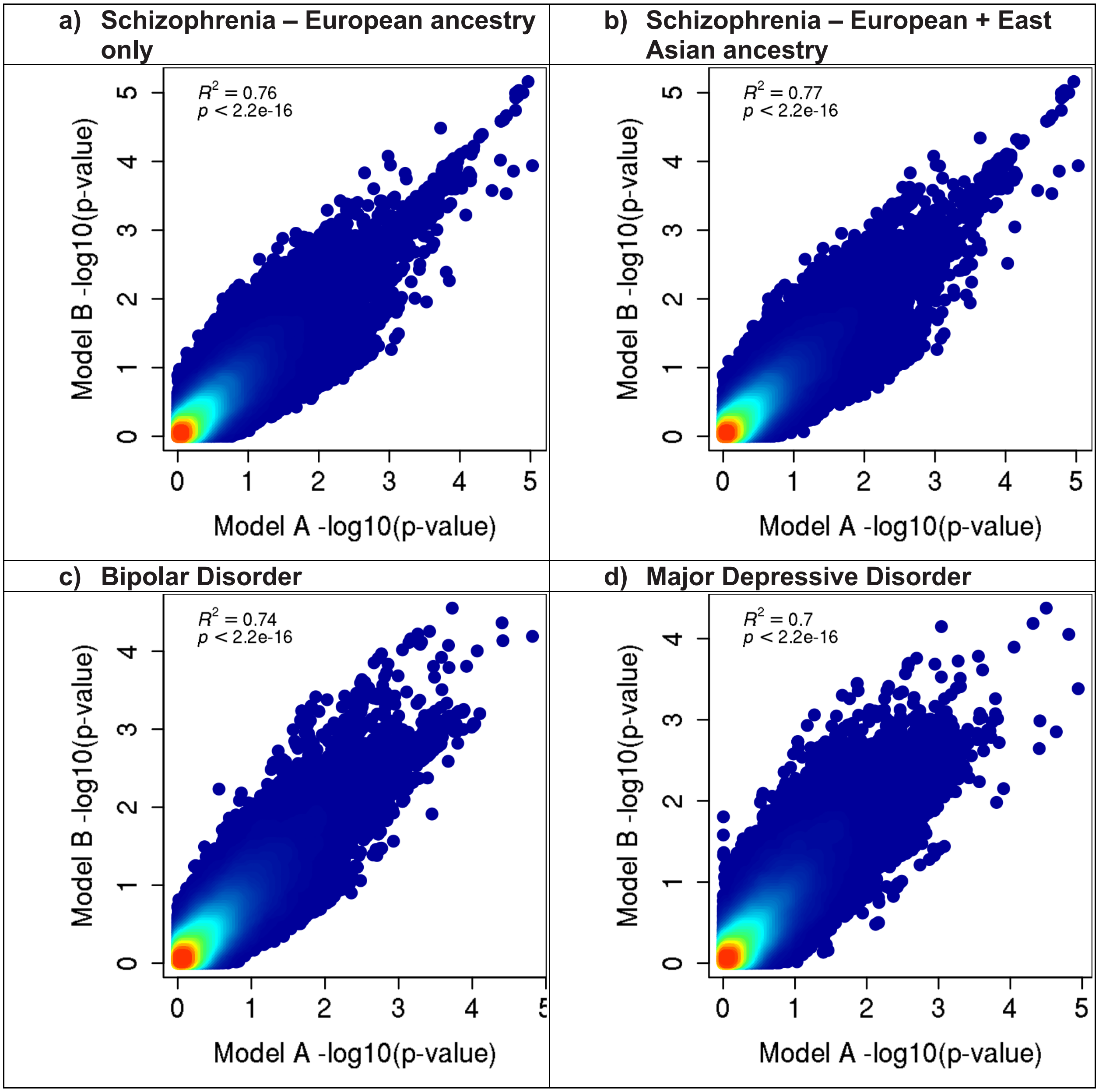

**e) Recurrent Major Depressive Disorder**

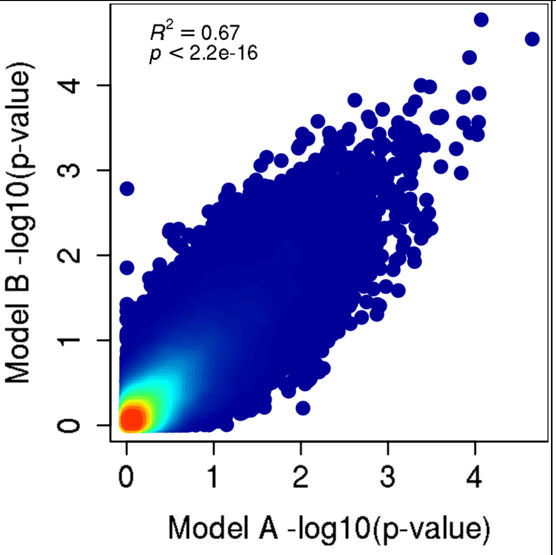


**f) Cross-Disorder SCZ-BIP-MDD – European ancestry only**

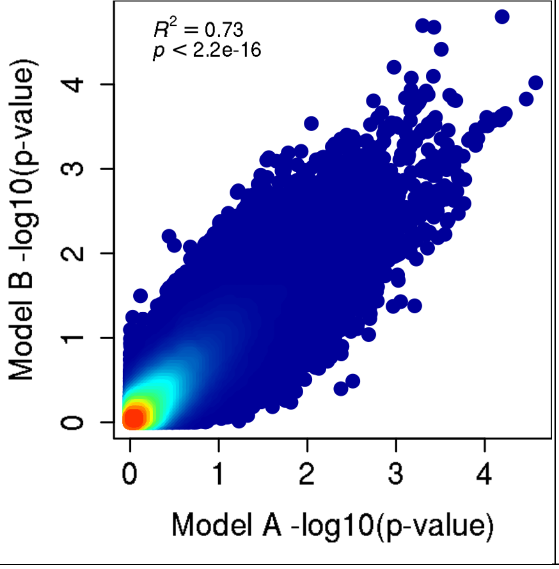


**g) Cross-Disorder SCZ-BIP-MDD – European + East Asian ancestry**

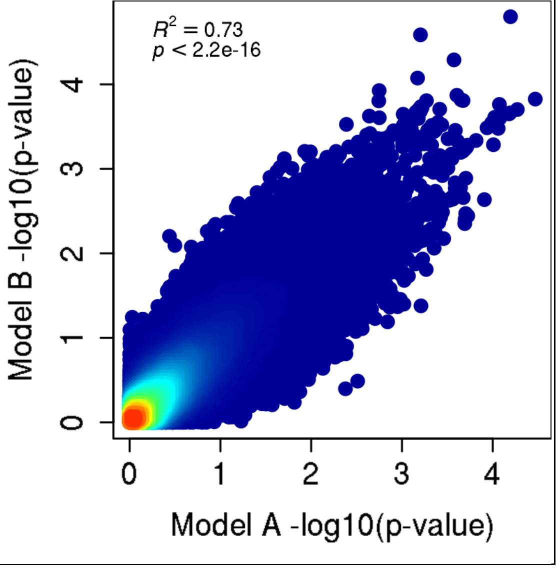

| **h) Cross-Disorder SCZ-BIP-rMDD – European ancestry only** | **i) Cross-Disorder SCZ-BIP-rMDD – European + East Asian ancestry** |
|---|---|
| 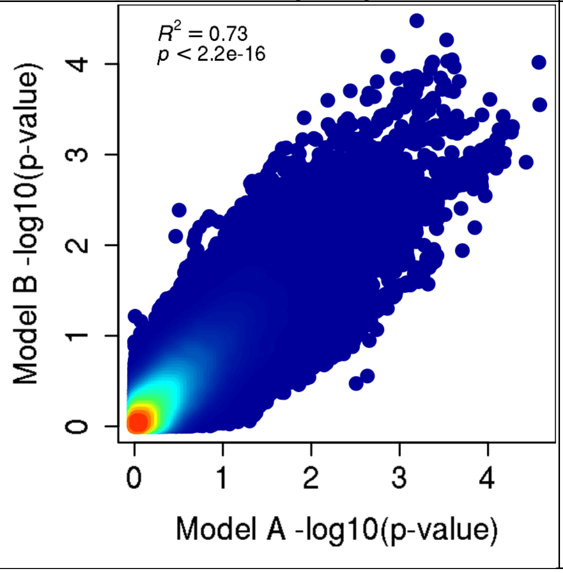 | 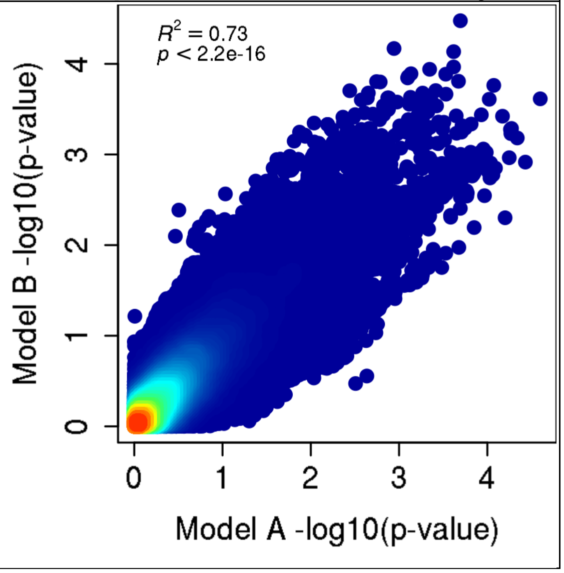 |

**Supplementary Figure 10. Manhattan plots for gene-based GxS tests in PGC + iPSYCH**

These analyses were carried out in MAGMA on the genomic control output with INFO score > 0.6, *European ancestry only*, and autosomal SNPs only, with the MHC region included.

Negative log10-transformed p-values for each gene (y-axis) are plotted by chromosomal position (x-axis). Each dot represents a gene, and the solid red and dotted blue horizontal lines represent the thresholds for genome-wide significant association ($p = 2.57{\times}10^{-6}$) and suggestive association ($p = 1{\times}10^{-5}$), respectively.

Plots were generated using the plot package in R.

Abbreviations: BIP = Bipolar Disorder; (r)MDD = (recurrent) Major Depressive Disorder; SCZ = Schizophrenia; SLTM = SAFB Like Transcription Modulator

**a) Schizophrenia**

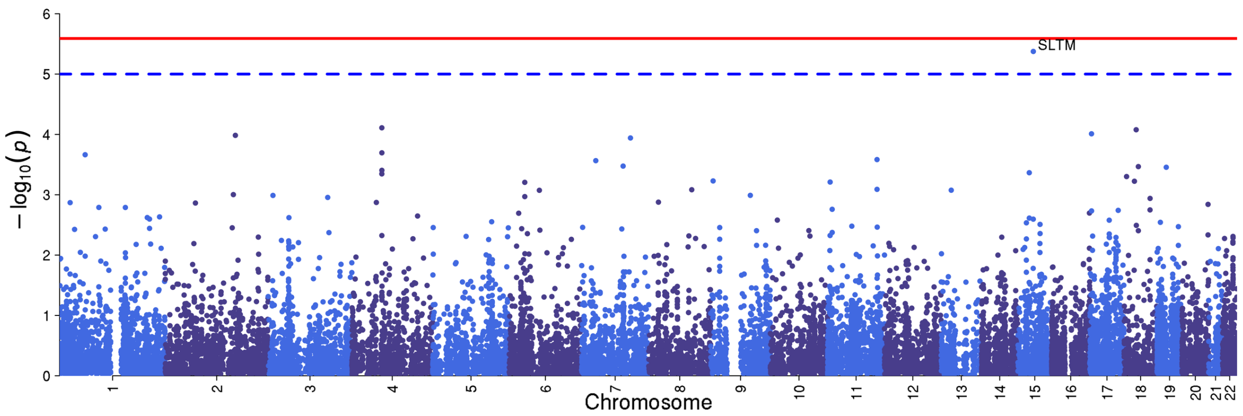


**b) Bipolar Disorder**

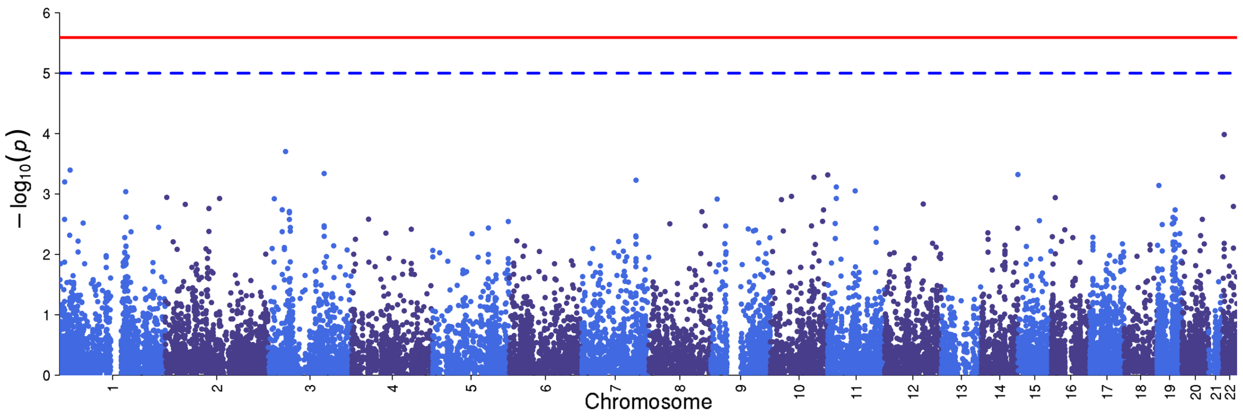

### c) Major Depressive Disorder

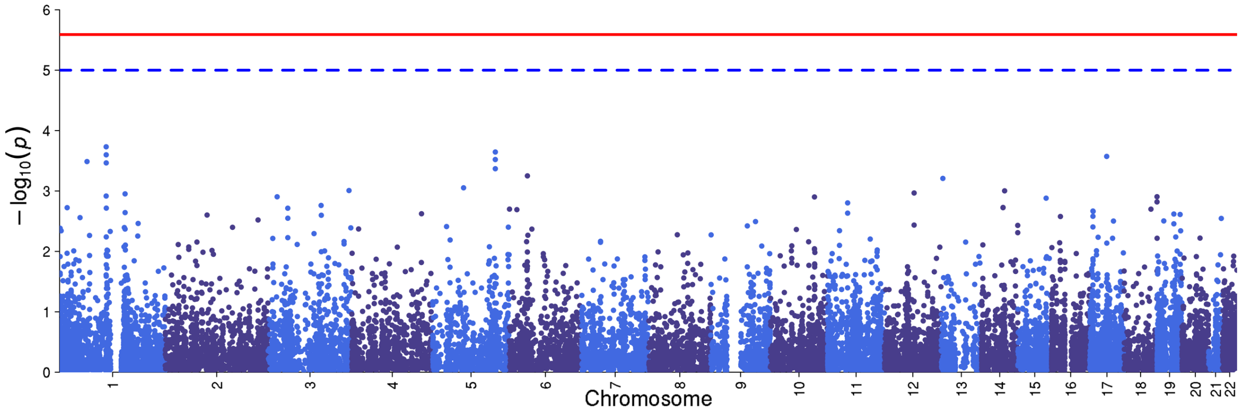


### d) Recurrent Major Depressive Disorder

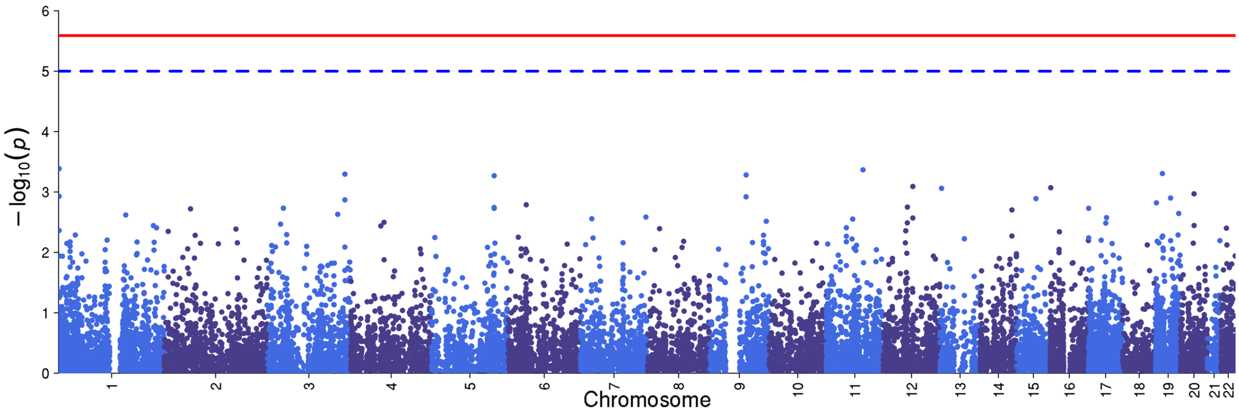


### e) Cross-Disorder SCZ-BIP-MDD

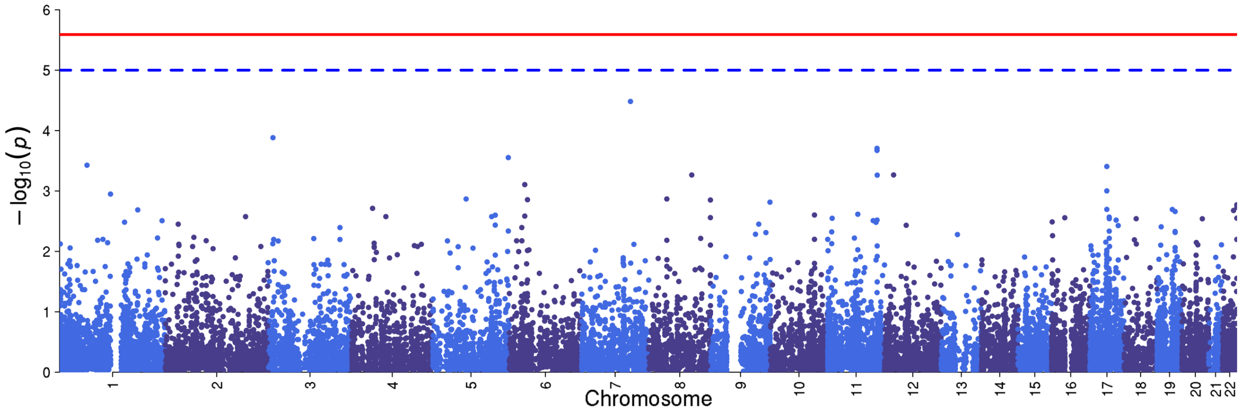

**f) Cross-Disorder SCZ-BIP-rMDD**

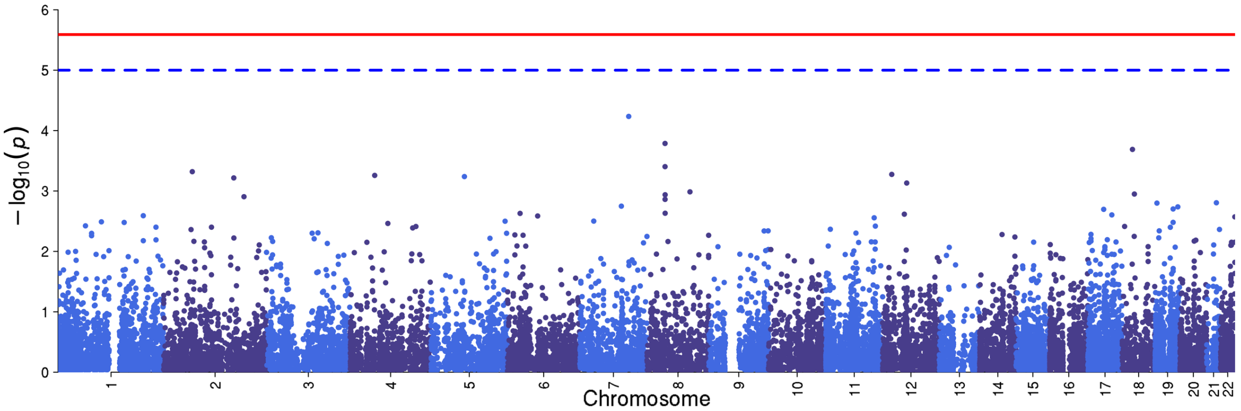


**g) Omnibus Test SCZ-BIP-MDD**

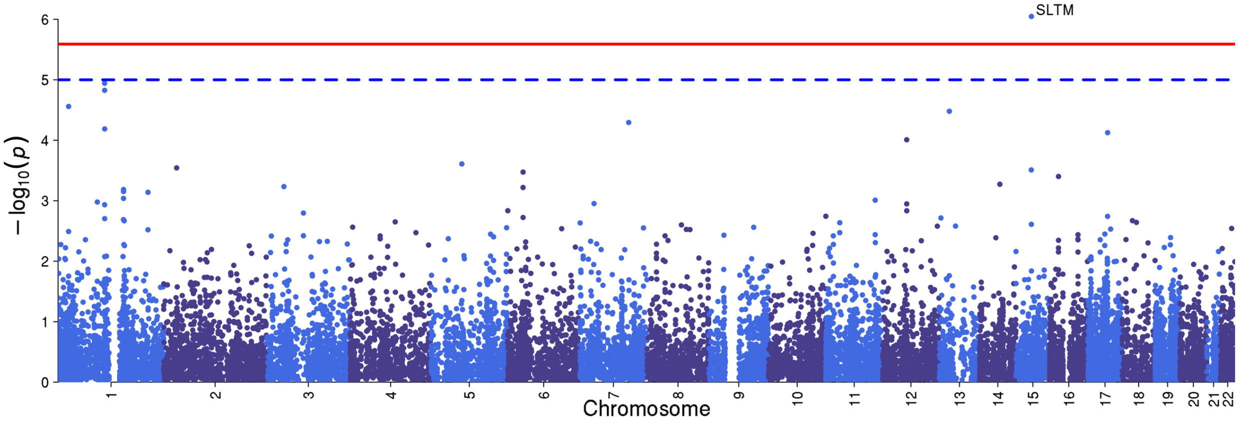


**h) Omnibus Test SCZ-BIP-rMDD**

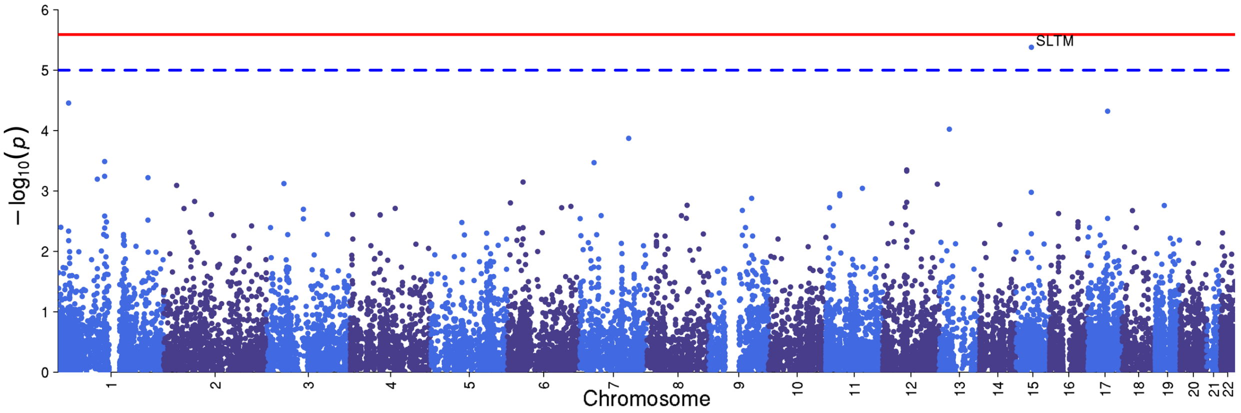

**Supplementary Figure 11. GTEx multi-tissue expression for GxS loci**

This plot was generated via website gtexportal.org.

Genes were included based on the following thresholds: SNP-based GxS interaction $p < 1\times10^{-6}$ and genes with gene-based test $p$-values $< 2.7\times10^{-6}$.

The "Brain - Frontal Cortex" and "Brain – Cortex", and the "Brain – Cerebellum" and "Brain - Cerebellar Hemisphere" samples should be considered as sample duplicates. One set of each pair (the "Brain – Cortex" and "Brain - Cerebellum)" were sampled at the same time as the remaining donor non-brain tissue samples, and were preserved in PAXgene tissue fixative solution. The remaining whole brain was then shipped to the University of Miami Brain Endowment Bank, where 8-11 brain sub-regions were sampled. The "Brain - Frontal Cortex" and "Brain - Cerebellar Hemisphere" were re-sampled at this time, as close as possible to the original sampling sites. All brain sub-regions sampled at the Miami Brain bank were preserved by snap freezing. Hence the paired brain regions differ in the time of sampling (those re-sampled at the Brain Bank, have a longer ischemic time) and in the manner in which the sample was preserved.

Abbreviations: ACC = Anterior Cingulate Cortex; BA = Brodmann Area; BG = basal ganglia; C1 = cervical-1; NAcc = Nucleus Accumbens; PFC = prefrontal cortex; TPM = Transcripts Per Kilobase Million mapped reads.

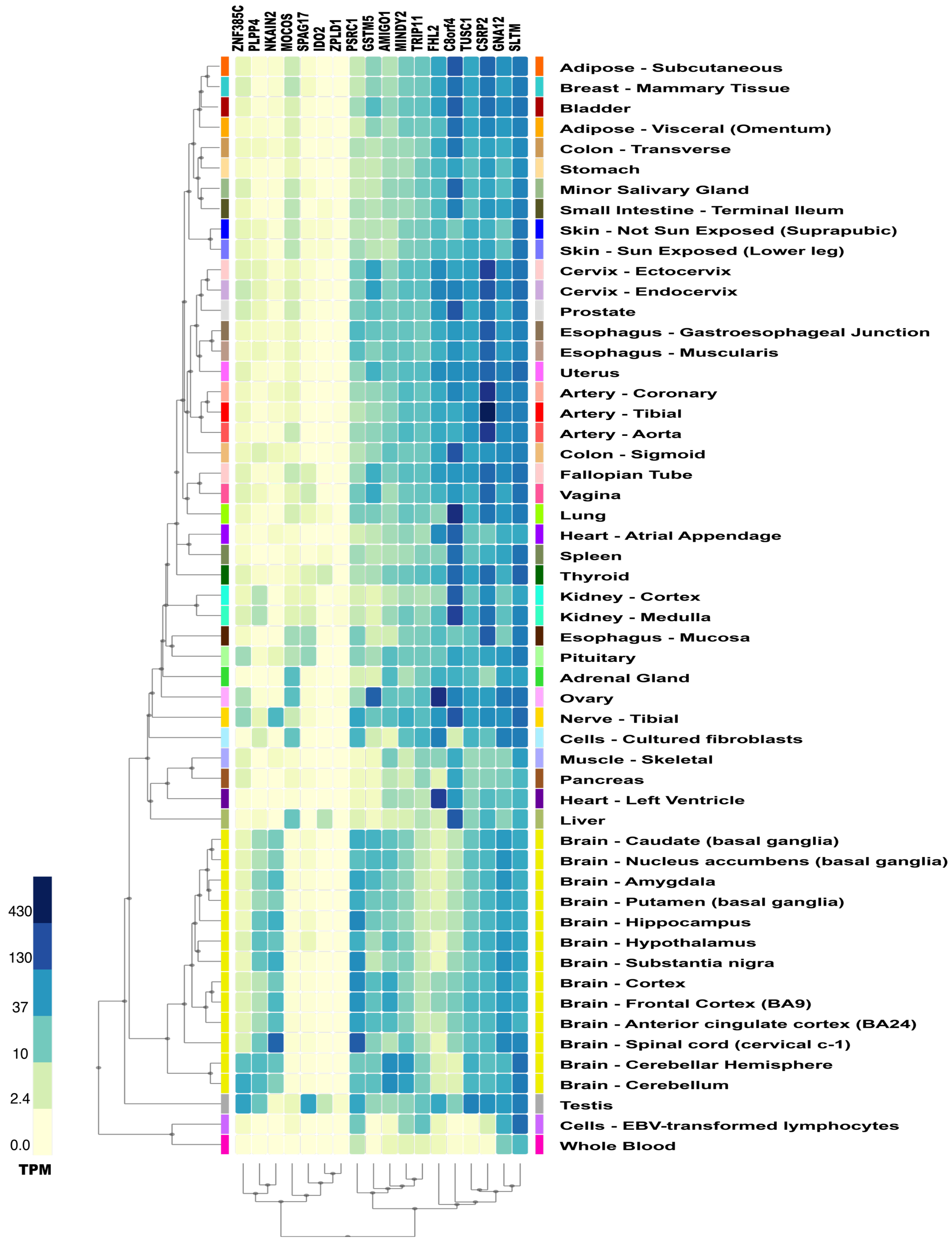
ZNF385C
PLPP4
NKAIN2
MOCOS
SPAG17
IDO2
ZPLD1
PSRC1
GSTM5
AMIGO1
MINDY2
TRIP11
FHL2
C8orf4
TUSC1
CSRP2
GNA12
SLTM
Adipose - Subcutaneous
Breast - Mammary Tissue
Bladder
Adipose - Visceral (Omentum)
Colon - Transverse
Stomach
Minor Salivary Gland
Small Intestine - Terminal Ileum
Skin - Not Sun Exposed (Suprapubic)
Skin - Sun Exposed (Lower leg)
Cervix - Ectocervix
Cervix - Endocervix
Prostate
Esophagus - Gastroesophageal Junction
Esophagus - Muscularis
Uterus
Artery - Coronary
Artery - Tibial
Artery - Aorta
Colon - Sigmoid
Fallopian Tube
Vagina
Lung
Heart - Atrial Appendage
Spleen
Thyroid
Kidney - Cortex
Kidney - Medulla
Esophagus - Mucosa
Pituitary
Adrenal Gland
Ovary
Nerve - Tibial
Cells - Cultured fibroblasts
Muscle - Skeletal
Pancreas
Heart - Left Ventricle
Liver
Brain - Caudate (basal ganglia)
Brain - Nucleus accumbens (basal ganglia)
Brain - Amygdala
Brain - Putamen (basal ganglia)
Brain - Hippocampus
Brain - Hypothalamus
Brain - Substantia nigra
Brain - Cortex
Brain - Frontal Cortex (BA9)
Brain - Anterior cingulate cortex (BA24)
Brain - Spinal cord (cervical c-1)
Brain - Cerebellar Hemisphere
Brain - Cerebellum
Testis
Cells - EBV-transformed lymphocytes
Whole Blood
430
130
37
10
2.4
0.0
TPM

**Supplementary Figure 12. Allen Brain Atlas expression across development for GxS loci**

Plots were generated using the ggplot package in R. Data were downloaded from https://human.brain-map.org/. Genes were included, in alphabetical order, based on the following thresholds: SNP-based GxS interaction $p < 1\times10^{-6}$ and genes with gene-based test $p$-values $< 2.7\times10^{-6}$.

Most of the genes examined were expressed in multiple brain regions at several stages from prenatal neurodevelopment through adulthood. However, some of the genes are predominantly expressed prenatally in one or more regions (*CRSP2*, *MOCOS, C8orf4* [= *TCIM*], *SPAG17*) or, in the case of *IDO2,* at the beginning of puberty (8-12 years) in prefrontal and orbitofrontal cortex.

Abbreviations: RPKM = Reads Per Kilobase of transcript per Million mapped reads; A1C = primary auditory cortex (core); AMY = amygdaloid complex; CB = cerebellum; CBC = cerebellar cortex; CGE = caudal ganglionic eminence; DFC = dorsolateral prefrontal cortex; DTH = dorsal thalamus; HIP = hippocampus (hippocampal formation); IPC = posteroventral (inferior) parietal cortex; ITC = inferolateral temporal cortex (area TEv, area 20); LGE = lateral ganglionic eminence; M1C = primary motor cortex (area M1, area 4); M1C-S1C = primary motor-sensory cortex (samples); MD = mediodorsal nucleus of thalamus; MFC = anterior (rostral) cingulate (medial prefrontal) cortex; MGE = medial ganglionic eminence; Ocx = occipital neocortex; OFC = orbital frontal cortex; PCx = parietal neocortex; S1C = primary somatosensory cortex (area S1, areas 3,1,2); STC = posterior (caudal) superior temporal cortex (area 22c); STR = striatum; TCx = temporal neocortex; URL = upper (rostral) rhombic lip; V1C = primary visual cortex (striate cortex, area V1/17); VFC = ventrolateral prefrontal cortex

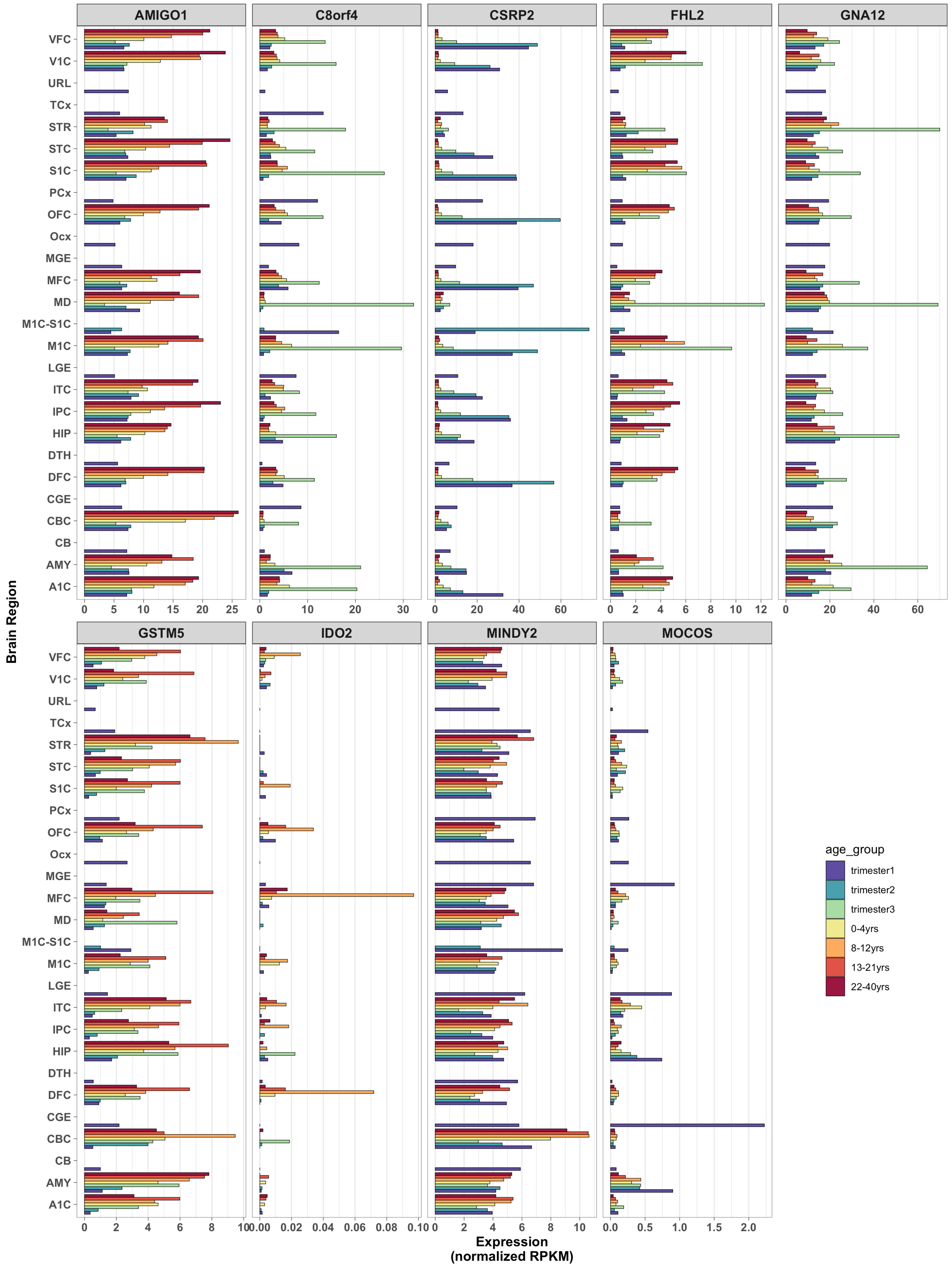
AMIGO1
C8orf4
CSRP2
FHL2
GNA12
GSTM5
IDO2
MINDY2
MOCOS
Brain Region
VFC
V1C
URL
TCx
STR
STC
S1C
PCx
OFC
Ocx
MGE
MFC
MD
M1C-S1C
M1C
LGE
ITC
IPC
HIP
DTH
DFC
CGE
CBC
CB
AMY
A1C
Expression
(normalized RPKM)
age_group
trimester1
trimester2
trimester3
0-4yrs
8-12yrs
13-21yrs
22-40yrs

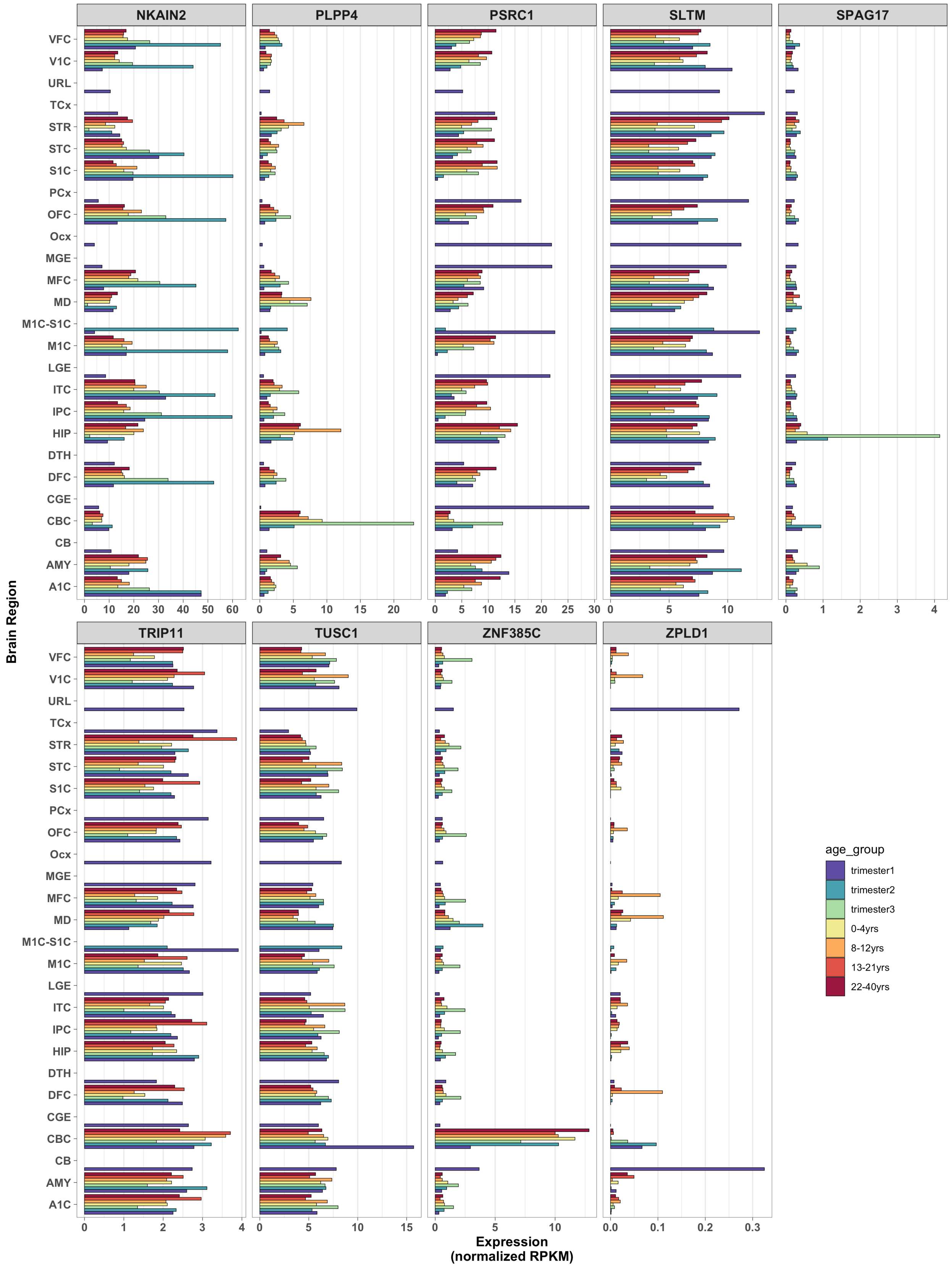
NKAIN2
PLPP4
PSRC1
SLTM
SPAG17
TRIP11
TUSC1
ZNF385C
ZPLD1
VFC
V1C
URL
TCx
STR
STC
S1C
PCx
OFC
Ocx
MGE
MFC
MD
M1C-S1C
M1C
LGE
ITC
IPC
HIP
DTH
DFC
CGE
CBC
CB
AMY
A1C
Brain Region
Expression
(normalized RPKM)
age_group
trimester1
trimester2
trimester3
0-4yrs
8-12yrs
13-21yrs
22-40yrs

**Supplementary Figure 13. Life brain expression course derived from the Human Brain Transcriptome (HBT) Project for GxS loci**

Plots were generated via website hbatlas.org. Genes were included, in alphabetical order, based on the following thresholds: SNP-based GxS interaction $p < 1\times10^{-6}$ and genes with gene-based test $p$-values $< 2.7\times10^{-6}$. Periods 1 through 7 are prenatal; Periods 8 and 9 are infant and toddler, respectively; Periods 10 and 11 are childhood; Periods 12 and 13 correspond to age ranges 12-20 years and 20-40 years, respectively; Periods 14 and 15 are middle age and 65+, respectively. Most of the genes examined were expressed in multiple brain regions at several stages from prenatal neurodevelopment through adulthood. However, some of the genes are predominantly expressed prenatally in one or more regions (*CRSP2*, *MOCOS, C8orf4* [= *TCIM*], *SPAG17*) or, in the case of *IDO2,* at the beginning of puberty (8-12 years) in prefrontal and orbitofrontal cortex.

Abbreviations: CBC = cerebellar cortex; MD = mediodorsal nucleus of the thalamus; STR = striatum; AMY = amygdala; HIP = hippocampus; NCX = 11 areas of neocortex.

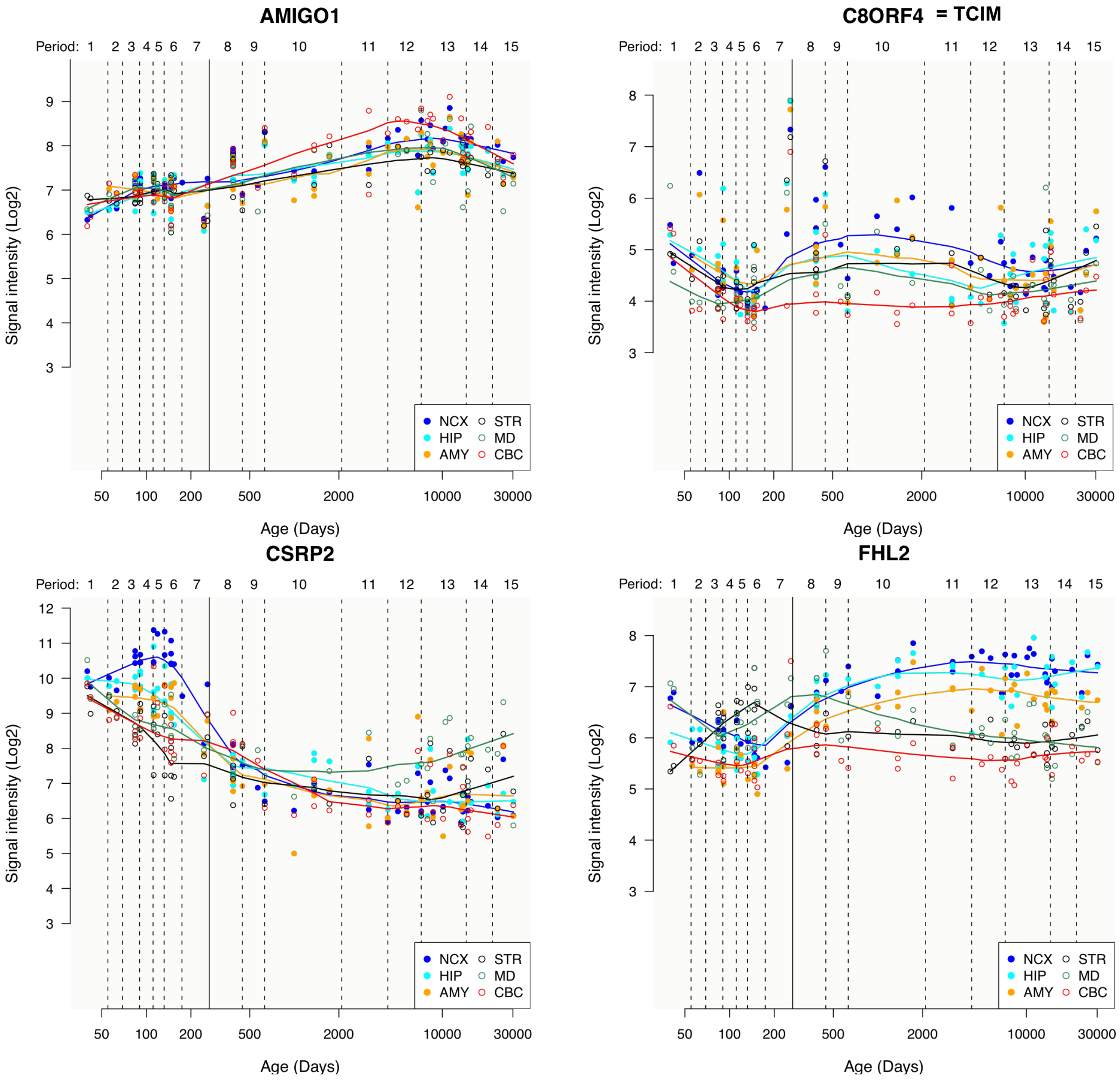

**GNA12**

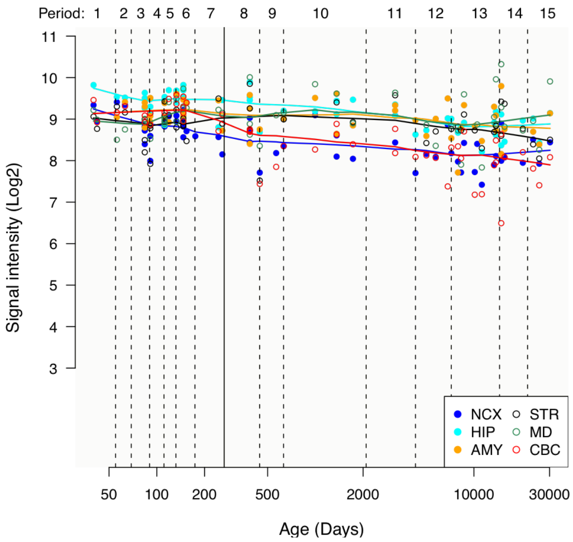


**GSTM5**

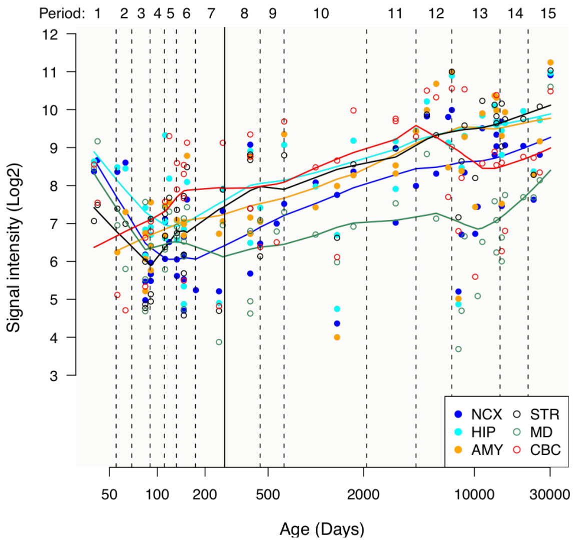


**IDO2**

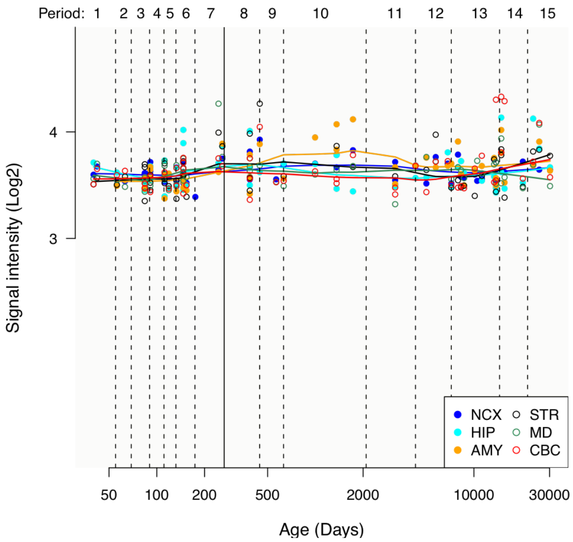


**MINDY2 = FAM63B**

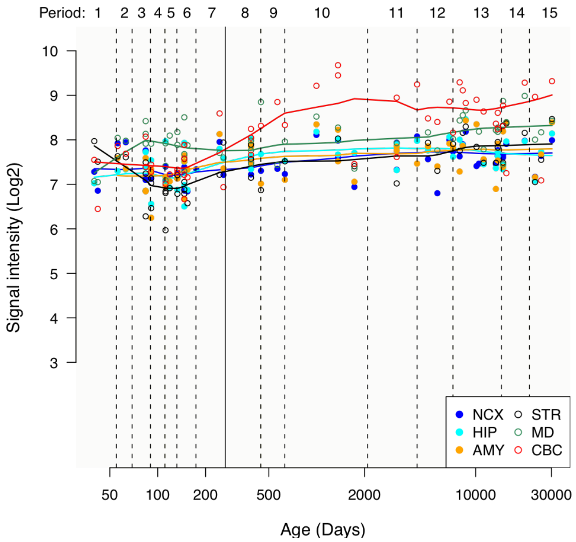


**MOCOS**

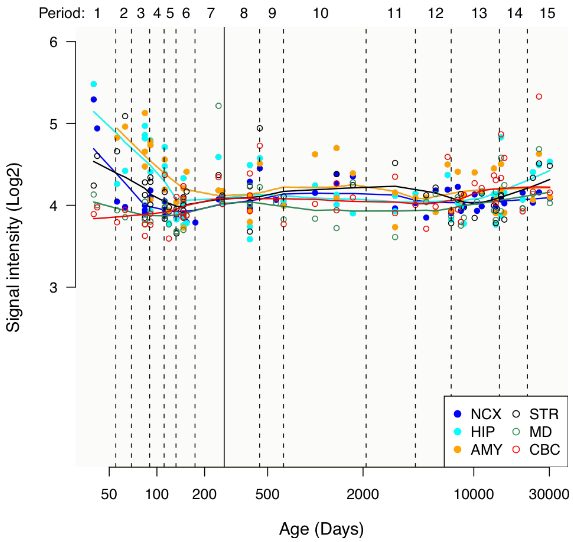


**NKAIN2**

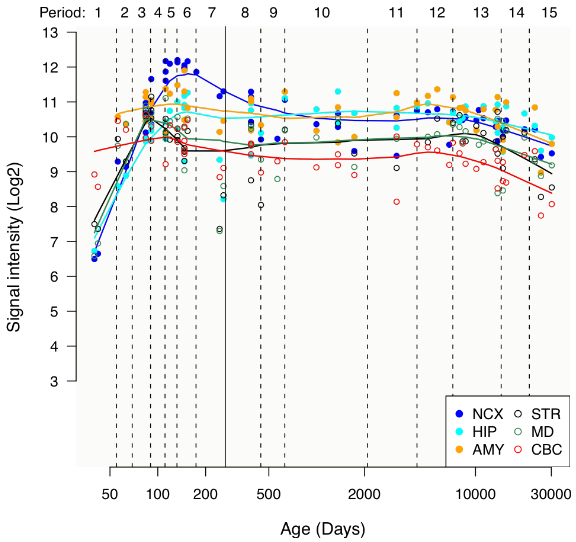

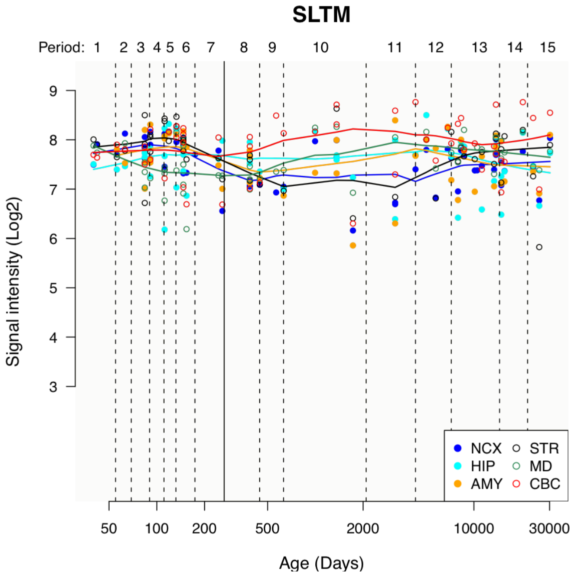
SLTM
Period: 1 2 3 4 5 6 7 8 9 10 11 12 13 14 15
Signal intensity (Log2)
Age (Days)
NCX STR
HIP MD
AMY CBC

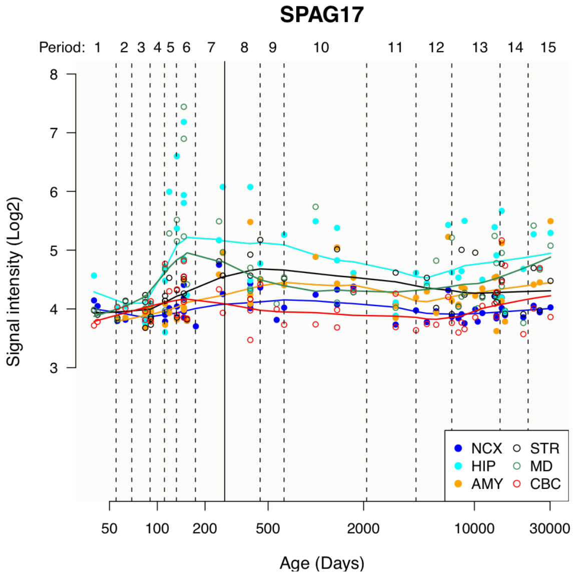
SPAG17
Period: 1 2 3 4 5 6 7 8 9 10 11 12 13 14 15
Signal intensity (Log2)
Age (Days)
NCX STR
HIP MD
AMY CBC

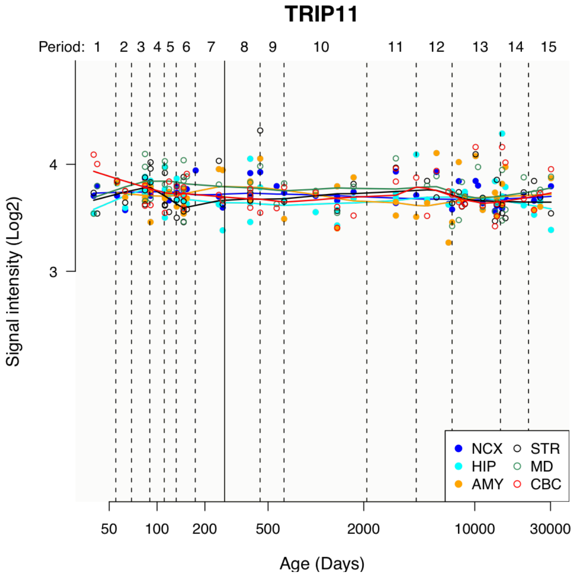
TRIP11
Period: 1 2 3 4 5 6 7 8 9 10 11 12 13 14 15
Signal intensity (Log2)
Age (Days)
NCX STR
HIP MD
AMY CBC

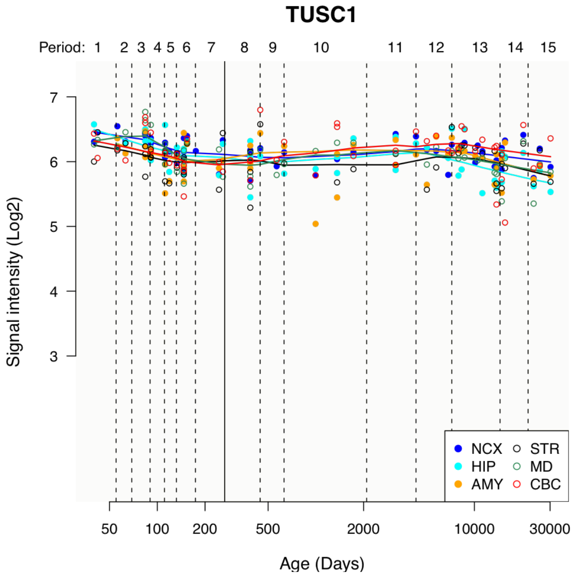
TUSC1
Period: 1 2 3 4 5 6 7 8 9 10 11 12 13 14 15
Signal intensity (Log2)
Age (Days)
NCX STR
HIP MD
AMY CBC

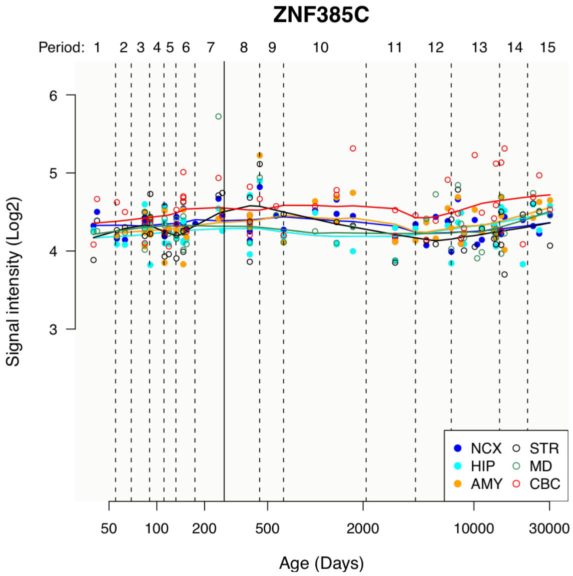
ZNF385C
Period: 1 2 3 4 5 6 7 8 9 10 11 12 13 14 15
Signal intensity (Log2)
Age (Days)
NCX STR
HIP MD
AMY CBC

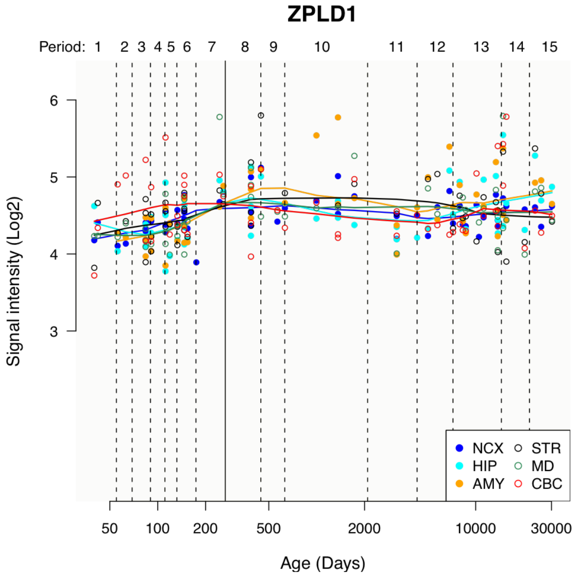
ZPLD1
Period: 1 2 3 4 5 6 7 8 9 10 11 12 13 14 15
Signal intensity (Log2)
Age (Days)
NCX STR
HIP MD
AMY CBC

**Supplementary Figure 14. GTEx sex-specific multi-tissue expression for GxS loci**

Expression data (v.8) were obtained from the Genotype-Tissue Expression (GTEx) project and downloaded from gtexportal.org. Tissue expression per gene was filtered for outliers with values > $9^{th}$ decile prior to t-test comparisons. Plots were generated using the ggplot package in R. Genes were included, in alphabetical order, based on the following thresholds: SNP-based GxS interaction $p < 1\times10^{-6}$ and genes with gene-based test $p$-values $< 2.7\times10^{-6}$. Evaluation of sex-specific expression detected significantly different expression levels between males and females of several of the genes, particularly in PFC, ACC, pituitary, and hypothalamus. * p < 0.05; ** p < 0.01 (Bonferroni-corrected for 14 tissues compared). The “Brain - Frontal Cortex” and “Brain – Cortex”, and the “Brain – Cerebellum” and “Brain - Cerebellar Hemisphere” samples should be considered as sample duplicates. One set of each pair (the “Brain – Cortex” and “Brain – Cerebellum”) were sampled at the same time as the remaining donor non-brain tissue samples, and were preserved in PAXgene tissue fixative solution. The remaining whole brain was then shipped to the University of Miami Brain Endowment Bank, where 8-11 brain sub-regions were sampled. The “Brain - Frontal Cortex” and “Brain - Cerebellar Hemisphere” were re-sampled at this time, as close as possible to the original sampling sites. All brain sub-regions sampled at the Miami Brain bank were preserved by snap freezing. Hence the paired brain regions differ in the time of sampling (those re-sampled at the Brain Bank, have a longer ischemic time) and in the mannesr in which the sample was preserved.

Abbreviations: ACC = Anterior Cingulate Cortex; BA = Brodmann Area; BG = basal ganglia; C1 = cervical-1; F = Females; M = Males; NAcc = Nucleus Accumbens; PFC = prefrontal cortex; TPM = Transcripts Per Kilobase Million mapped reads.

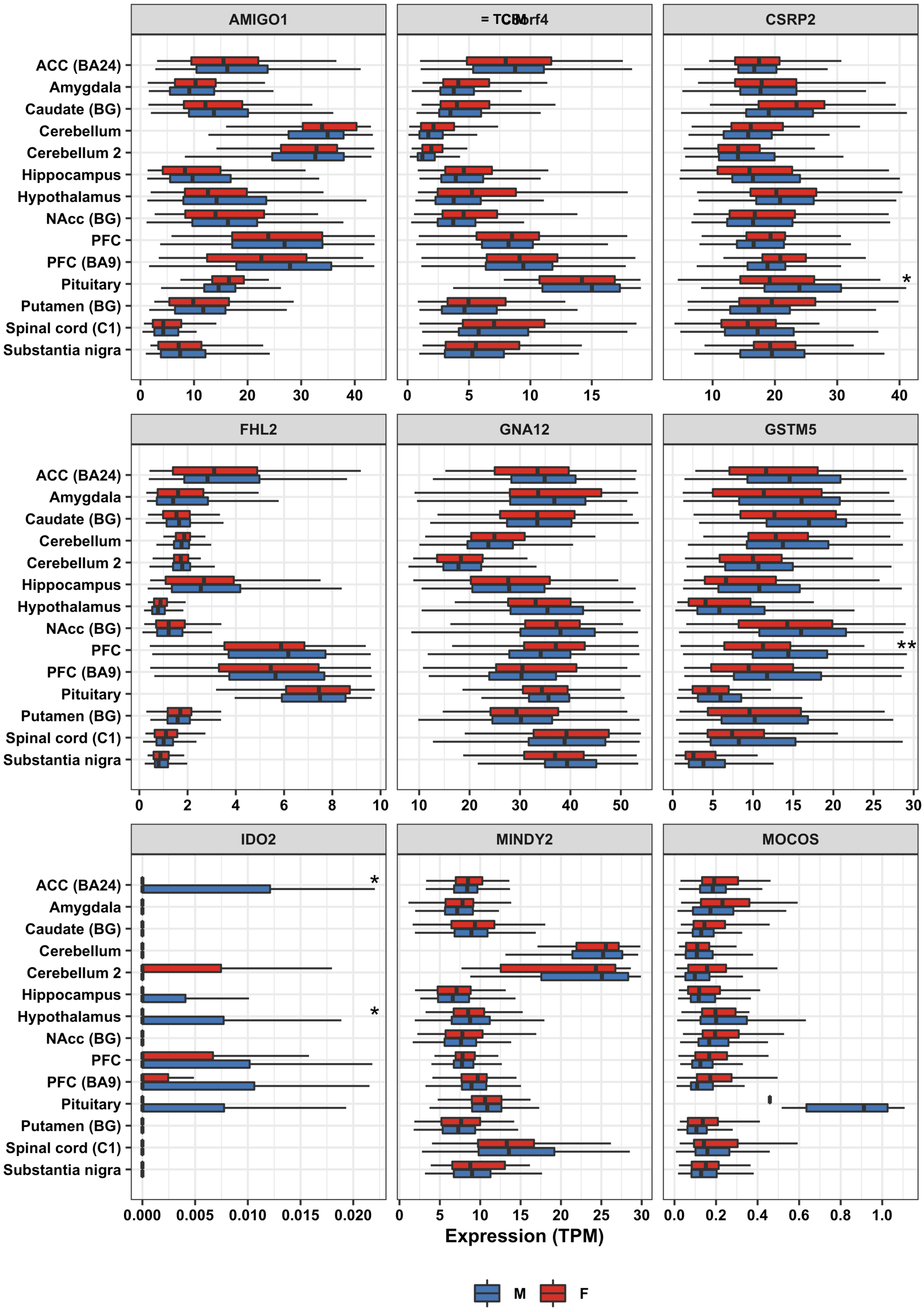
AMIGO1
= TCIM
C8orf4
CSRP2
FHL2
GNA12
GSTM5
IDO2
MINDY2
MOCOS
ACC (BA24)
Amygdala
Caudate (BG)
Cerebellum
Cerebellum 2
Hippocampus
Hypothalamus
NAcc (BG)
PFC
PFC (BA9)
Pituitary
Putamen (BG)
Spinal cord (C1)
Substantia nigra
*
**
Expression (TPM)
M
F

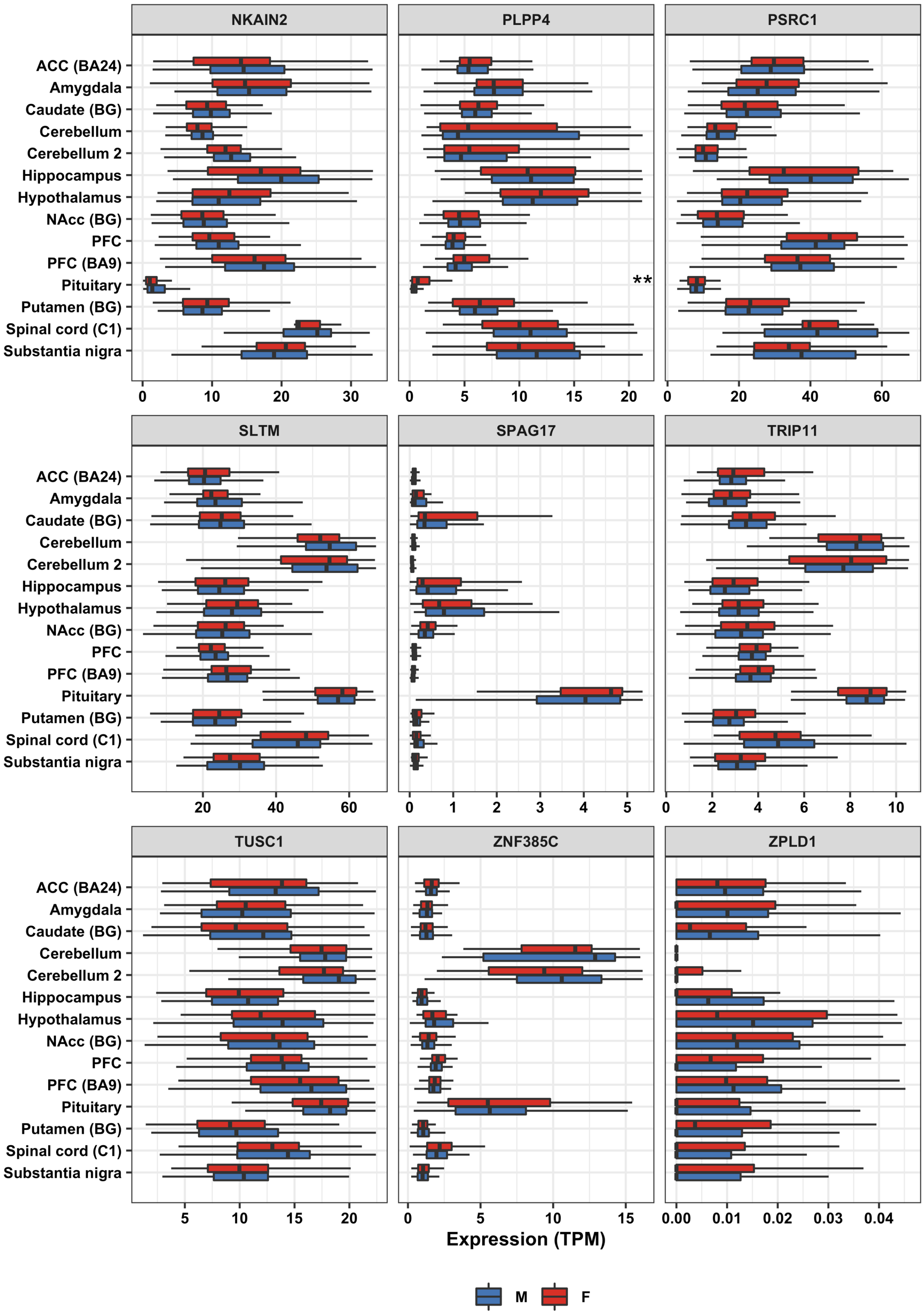
NKAIN2
PLPP4
PSRC1
SLTM
SPAG17
TRIP11
TUSC1
ZNF385C
ZPLD1
ACC (BA24)
Amygdala
Caudate (BG)
Cerebellum
Cerebellum 2
Hippocampus
Hypothalamus
NAcc (BG)
PFC
PFC (BA9)
Pituitary
Putamen (BG)
Spinal cord (C1)
Substantia nigra
**
Expression (TPM)
M
F

**Supplementary Figure 15. Cell type-specific brain expression derived from the Stanford RNA-Seq database for GxS loci**

Mouse brain expression data were downloaded from https://www.brainrnaseq.org/. Genes were mapped to human orthologous genes using Ensembl. Genes were included, in alphabetical order, based on the following thresholds: SNP-based GxS interaction $p < 1\times10^{-6}$ and genes with gene-based test p-values $< 2.7\times10^{-6}$. Plots were generated using the ggplot package in R. Among seven brain cell types, the genes examined are expressed in various cell types, with no preponderance of expression in a particular type.

Abbreviations: FPKM = Fragments Per Kilobase of transcript per Million mapped reads.

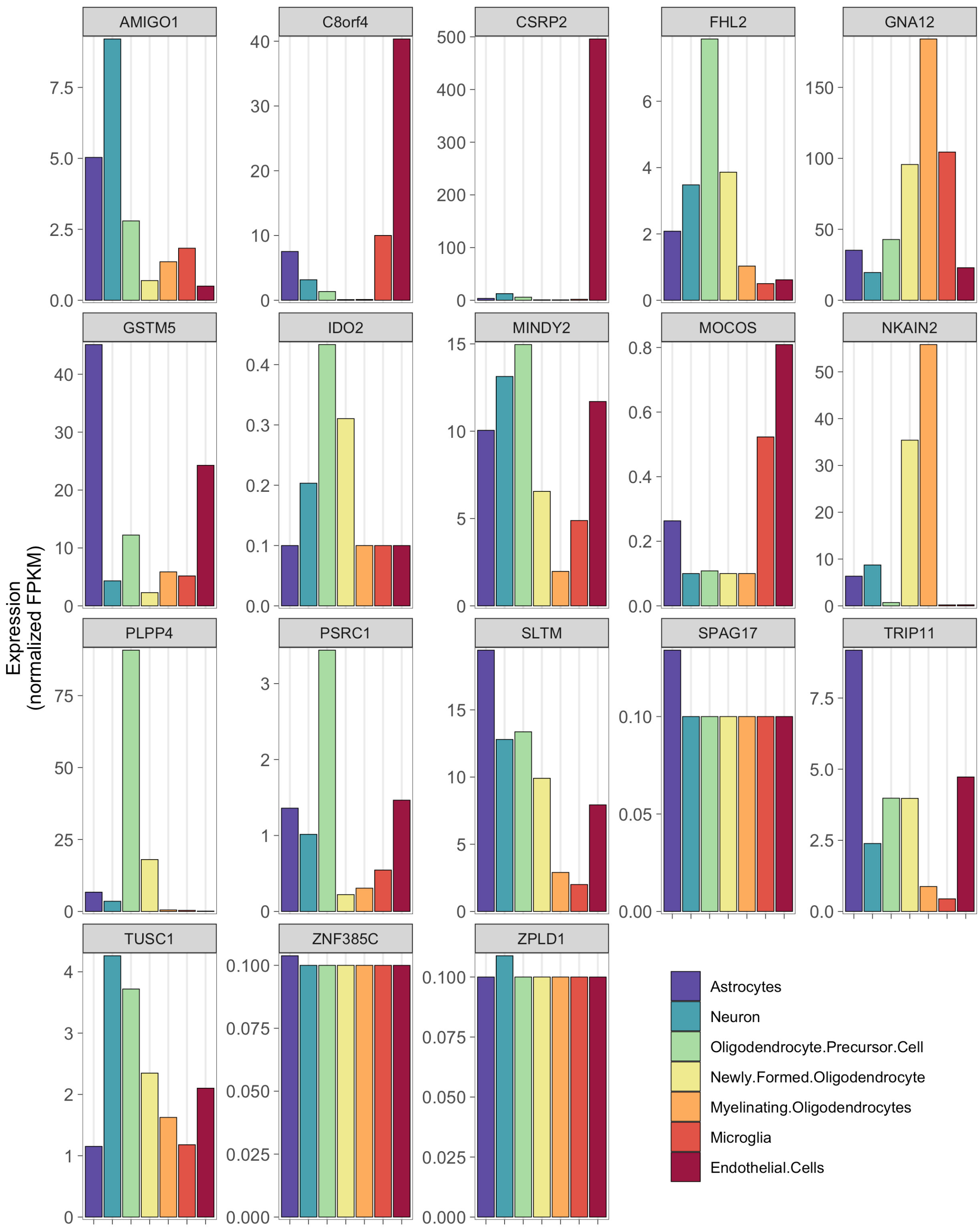
AMIGO1
C8orf4
CSRP2
FHL2
GNA12
GSTM5
IDO2
MINDY2
MOCOS
NKAIN2
PLPP4
PSRC1
SLTM
SPAG17
TRIP11
TUSC1
ZNF385C
ZPLD1
Expression
(normalized FPKM)
Astrocytes
Neuron
Oligodendrocyte.Precursor.Cell
Newly.Formed.Oligodendrocyte
Myelinating.Oligodendrocytes
Microglia
Endothelial.Cells

### Supplementary Tables PGC only

#### Supplementary Table 15. Meta-analysis Autosomal GxS interaction loci in PGC

See SupplTable15_MetaAnalysisSTDERR_auto_PGC.xlsx

Cross-disorder and within-disorder meta-analyses were carried out using METAL, incorporating cohort-level summary statistics from PLINK. Listed are LD-independent SNPs with interaction *p*-values < $1{\times}10^{-6}$ in SCZ, BIP, (r)MDD, and cross-disorder. Loci were clumped using '*plink --bfile 1kgp_ref_file --clump metal_output --clump-p1 1e-4 --clump-p2 1e-4 --clump-r2 0.6 --clump-kb 3000*'

Abbreviations: BIP = bipolar disorder; (r)MDD = (recurrent) major depressive disorder; SCZ = schizophrenia.

#### Supplementary Table 16. Omnibus test Autosomal GxS interaction loci in PGC

See SupplTable16_OmnibusTestASSET_auto_PGC.xlsx

Omnibus tests were carried out using ASSET, incorporating the within-disorder meta-analysis summary statistics from METAL. Listed are LD-independent SNPs with cross-disorder interaction *p*-values < $1{\times}10^{-6}$. Loci were clumped using '*plink --bfile 1kgp_ref_file --clump asset_output --clump-p1 1e-4 --clump-p2 1e-4 --clump-r2 0.6 --clump-kb 3000*'

Abbreviations: BIP = bipolar disorder; (r)MDD = (recurrent) major depressive disorder; SCZ = schizophrenia.

#### Supplementary Table 17. Meta-analysis chrX GxS interaction loci in PGC

See SupplTable17_MetaAnalysisSTDERR_xchr_PGC.xlsx

Cross-disorder and within-disorder meta-analyses were carried out using METAL, incorporating cohort-level summary statistics from PLINK. Listed are LD-independent SNPs with interaction *p*-values < $1{\times}10^{-6}$ in SCZ, BIP, (r)MDD, and cross-disorder. Model A **(a)** effectively assumes complete and uniform X-inactivation in females and a similar effect size between males and females. Females are considered to have 0, 1, or 2 copies of an allele; males are considered to have 0 or 2 copies of the same allele. Model B **(b)** considers the allelic dosages for females to be 0,1, or 2 copies, and males to be 0 or 1 copy as in an autosomal analysis. Loci were clumped using '*plink --bfile 1kgp_ref_file --clump metal_output --clump-p1 1e-4 --clump-p2 1e-4 --clump-r2 0.6 --clump-kb 3000*'

Abbreviations: BIP = bipolar disorder; (r)MDD = (recurrent) major depressive disorder; SCZ = schizophrenia.

#### Supplementary Table 18. Omnibus test chrX GxS interaction loci in PGC

See SupplTable18_OmnibusTestASSET_xchr_PGC.xlsx

Omnibus tests were carried out using ASSET, incorporating the within-disorder meta-analysis summary statistics from METAL. Listed are LD-independent SNPs with cross-disorder interaction *p*-values < $1{\times}10^{-6}$. Loci were clumped using '*plink --bfile 1kgp_ref_file --clump asset_output --clump-p1 1e-4 --clump-p2 1e-4 --clump-r2 0.6 --clump-kb 3000*'

Abbreviations: BIP = bipolar disorder; (r)MDD = (recurrent) major depressive disorder; SCZ = schizophrenia.

**Supplementary Table 19. Credible SNPs for GxS loci in PGC**

See SupplTable19_CredibleSNPs_FineMapping_PGC.xlsx

Fine mapping was carried out using both FINEMAP and CAVIAR. Fine mapping using FINEMAP was carried out with settings: *--sss --corr-config 0.95 --n-causal-snps 5 --n-configs-top 50000 --prior-k0 0 --prior-std 0.05*. If there were less than 5 SNPs in the locus, *--n-causal-snps* was set to the number of SNPs in the locus according to LD. The most likely causal SNPs per locus are highlighted in bold font. The shotgun stochastic search (--sss) conducts a pre-defined number of iterations within the space of causal configurations. In each iteration, the neighborhood of the current causal configuration is defined by configurations that result from deleting, changing or adding a causal SNP from the current configuration. The next iteration starts by sampling a new causal configuration from the neighborhood based on the scores normalized within the neighborhood. Fine mapping using CAVIAR was carried out with settings: *-r 0.95 -c 5 -f 1*. If there were less than 5 SNPs in the locus, *-c* was set to the number of SNPs in the locus according to LD. Analyses used European ancestry only summary statistics. Loci with $p < 1\times10^{-6}$ were analyzed (index SNPs determined based on clumping using LD threshold 0.1). The most likely causal SNPs per locus are highlighted in bold font.

Abbreviations: PP_group = posterior probability that there is at least one causal signal among SNPs in the same group with this SNP; PP_causal = posterior probability that the SNP is causal; BP = base pair position; BIP = bipolar disorder; CHR = chromosome; (r)MDD = (recurrent) major depressive disorder; SCZ = schizophrenia; SNP = Single Nucleotide Polymorphism rs ID.

**Supplementary Table 20. Gene-based test in PGC**

See SupplTable20_Gene-BasedTest_PGC.xlsx

Gene-based analyses were carried out in MAGMA on the genomic control output with INFO score > 0.6, European ancestry only, and autosomal SNPs only, with the MHC region included. Genes with *p*-values $< 1\times10^{-4}$ are shown. There was no difference in the *p*-values when the MHC region was excluded. There were minor differences in *p*-values when using INFO score > 0.8, but with the same top 10 genes. *Significant at genome-wide threshold for gene-based test of 0.05 / 19,427 genes = $2.6\times10^{-6}$.

Abbreviations: BP = base pair position; Chr = chromosome; N SNPs = number of SNPs in gene; N Param = number of parameters; N = sample size; Z = Z-statistic; BIP = bipolar disorder; MDD = major depressive disorder; rMDD = recurrent major depressive disorder; SCZ = schizophrenia.

**Supplementary Table 21. MSigDB pathway gene set enrichment analyses in PGC**

See SupplTable21_MSigDB pathway GSEA_PGC.xlsx

Enrichment analyses were carried out in MAGMA on the genomic control output with INFO score > 0.6, European ancestry only, and autosomal SNPs only. Analyses were run both with (top subtable) and without (bottom subtable) inclusion of the Chromosome 6 MHC region. Each (sub)table displays the top 10 gene sets based on the uncorrected *p*-value. Hyperlinks link to the GSEA/MSigDB website with a description of the pathway.

Abbreviations: BIP = bipolar disorder; MDD = major depressive disorder; $P_{BONF}$ = Bonferroni-corrected *p*-value; $P_{FDR}$ = False Discovery Rate-corrected *p*-value; rMDD = recurrent major depressive disorder; SCZ = schizophrenia; SE = Standard Error.

**Supplementary Table 22. Selected pathway gene set enrichment analyses in PGC**

See SupplTable22_Selected pathway GSEA_PGC.xlsx

Analyses were run with (top) and without (bottom) inclusion of the Chromosome 6 MHC region in MAGMA. These analyses were carried out on the genomic control output with INFO score > 0.6, European ancestry only, and autosomal SNPs only. * Significant after adjusting *p*-values for multiple testing.

Abbreviations: BIP = bipolar disorder; CNS = central nervous system; MDD = major depressive disorder; MP = Mouse Phenome; $P_{FDR}$ = False Discovery Rate-corrected *p*-value; PGC-NPA = Psychiatric Genomics Consortium – Network and Pathway Analysis Working Group; rMDD = recurrent major depressive disorder; SCZ = schizophrenia; SE = Standard Error.

### Supplementary Figures PGC only

**Supplementary Figure 16. LD Score Regression estimates of SNP-based (a) heritability and (b) genetic correlations (SE) in PGC only**

This graph shows $h^2$ and $r_g$ estimates for MAF > 0.01.

a) Heritability estimates were substantially different between the sexes for SCZ ($p_{FDR}$ = 0.019) and MDD ($p_{FDR}$ = 0.005), but not BIP ($p_{FDR}$ = 0.381).

b) SNP-based genetic correlations ($r_g$) between males and females within each disorder ranged between 0.86 and 1 and were significantly different from 1 for SCZ ($p_{FDR}$ = 0.039) and BIP ($p_{FDR}$ = 0.039), but not MDD ($p_{FDR}$ = 0.397). No significant differences in the cross-disorder genetic correlations between males and females, with the exception of $r_g$ between BIP and MDD ($r_{gF}$ = 0.42; $r_{gM}$ = 0.04; $p_{FDR}$ = 0.044).

Abbreviations: BIP = Bipolar Disorder; MDD = Major Depressive Disorder; SCZ = Schizophrenia; F = Females; M = Males; SE = standard error.

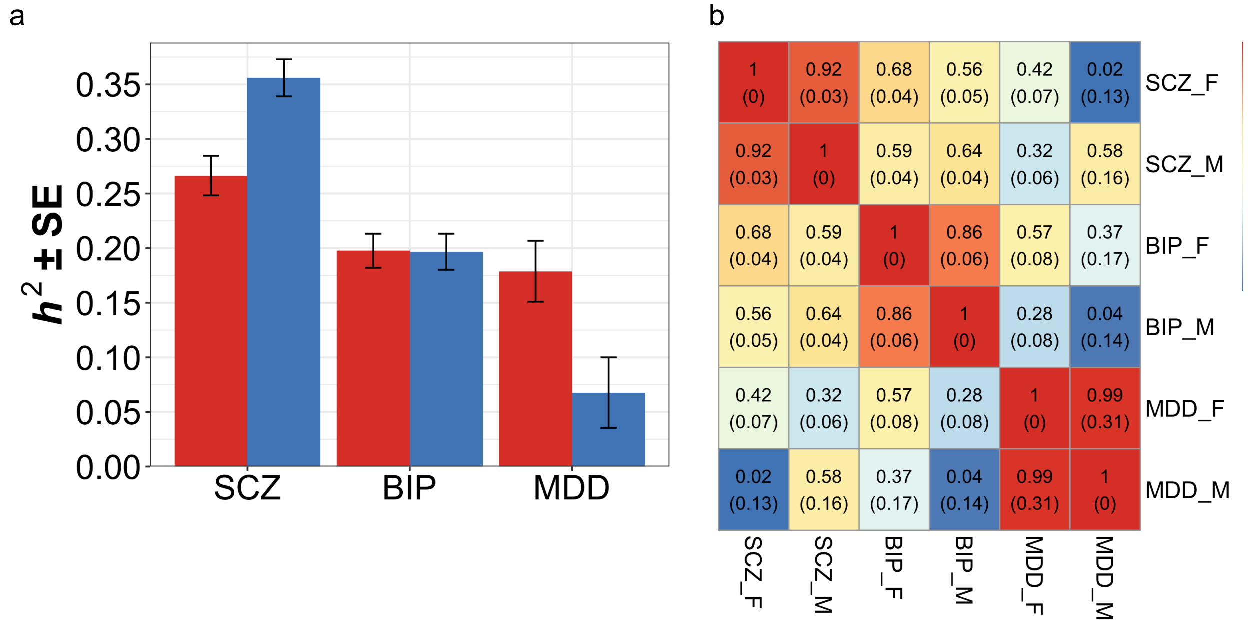

**Supplementary Figure 17. Miami plots for sex-stratified analyses in PGC**

GWAS SNP main effects for men (blue) are plotted downward, and are plotted upward for women (green). Negative log10-transformed p-values for each variant (each dot) (y-axis) are plotted by chromosomal position (x-axis). The solid red and dotted blue horizontal lines represent the thresholds for genome-wide significant association ($p = 5{\times}10^{-8}$) and suggestive association ($p = 1{\times}10^{-5}$), respectively. Plotted are the regular meta-analysis results within and across disorders only; omnibus tests were not carried out for sex-stratified analyses.

Plots were generated using the plot package in R.

Abbreviations: BIP = Bipolar Disorder; (r)MDD = (recurrent) Major Depressive Disorder; SCZ = Schizophrenia

### a) Schizophrenia – European ancestry only

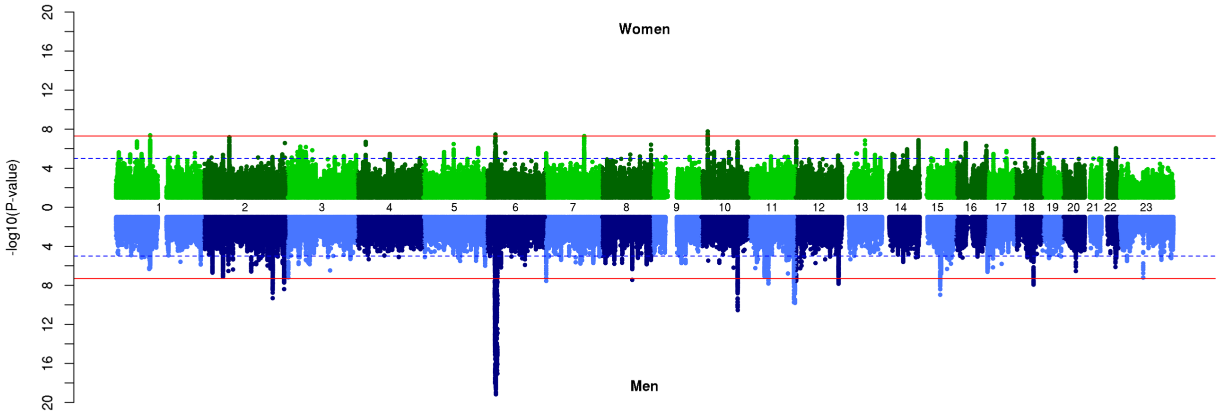


### b) Schizophrenia – European + East Asian ancestry

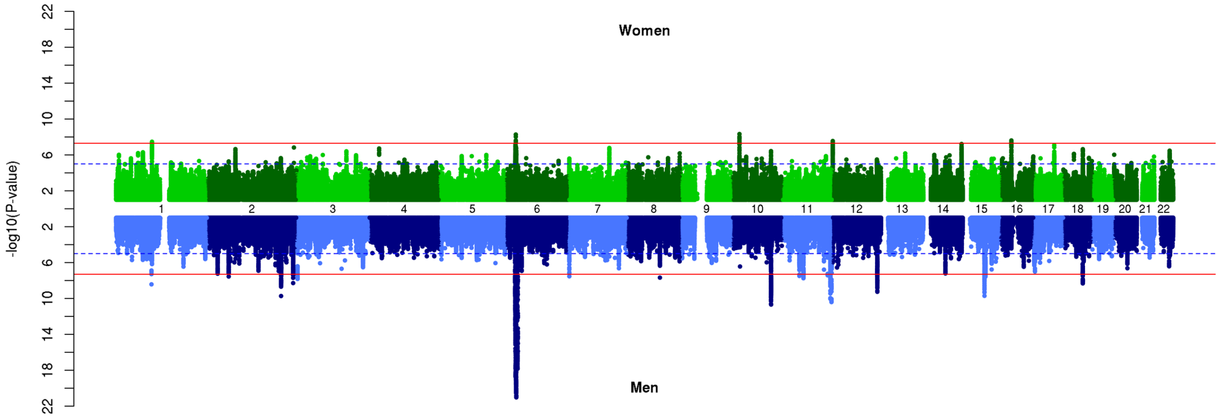

### c) Bipolar Disorder

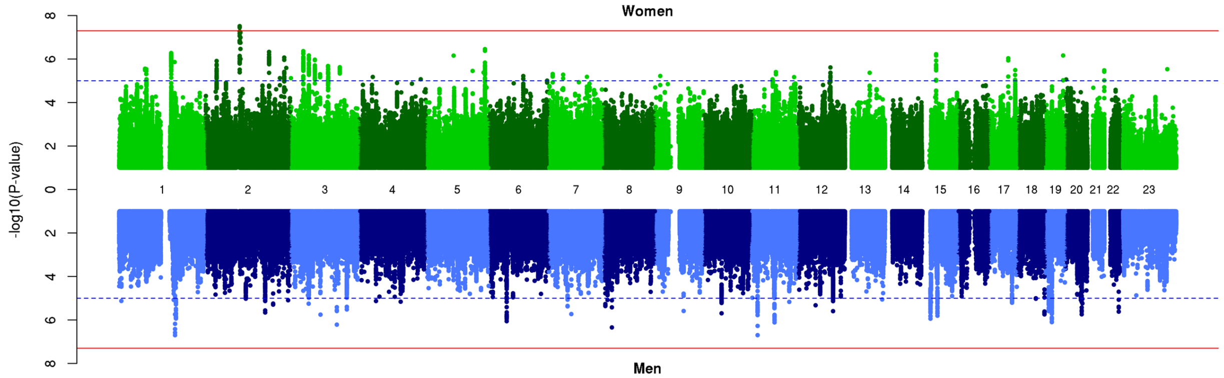


### d) Major Depressive Disorder

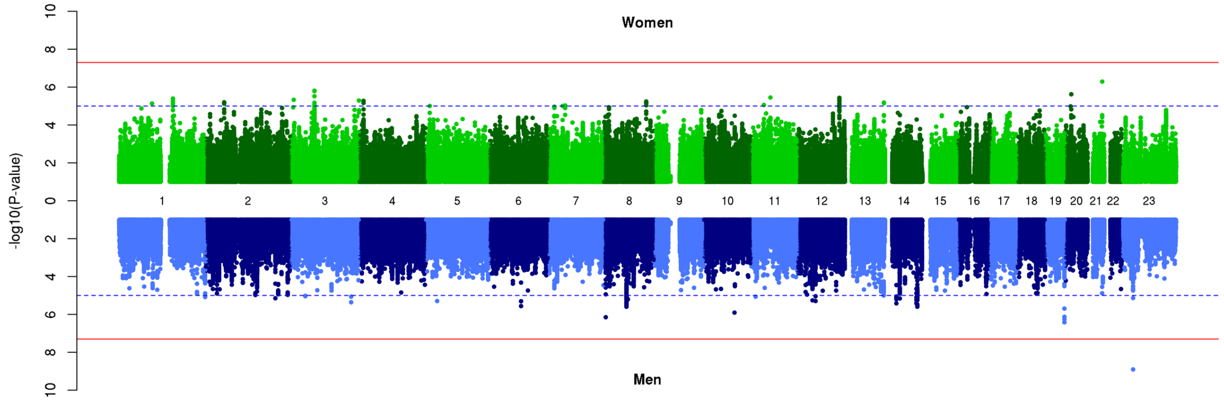


### e) Recurrent Major Depressive Disorder

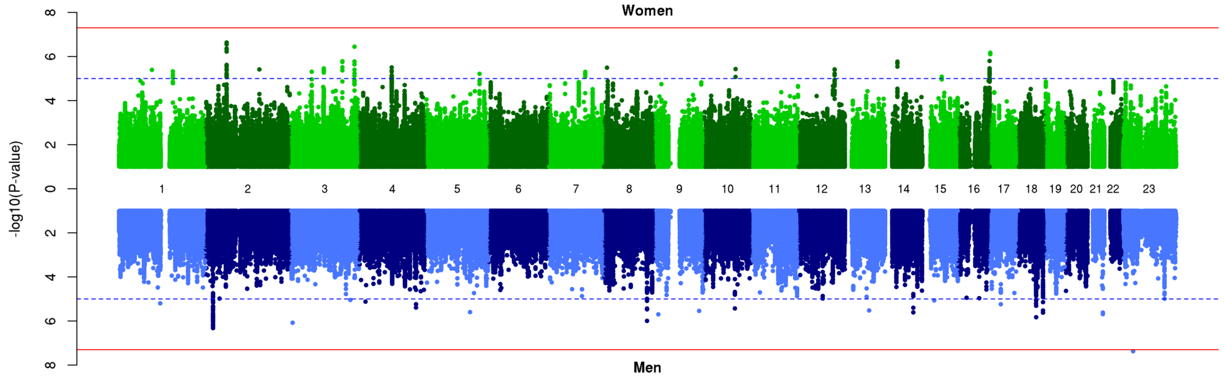

### f) Cross-Disorder SCZ-BIP-MDD – European ancestry only

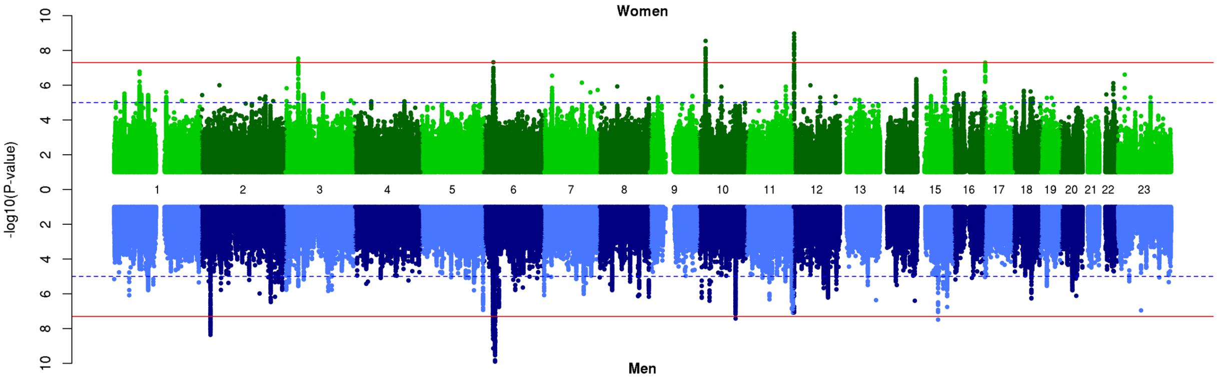


### g) Cross-Disorder SCZ-BIP-MDD – European + East Asian ancestry

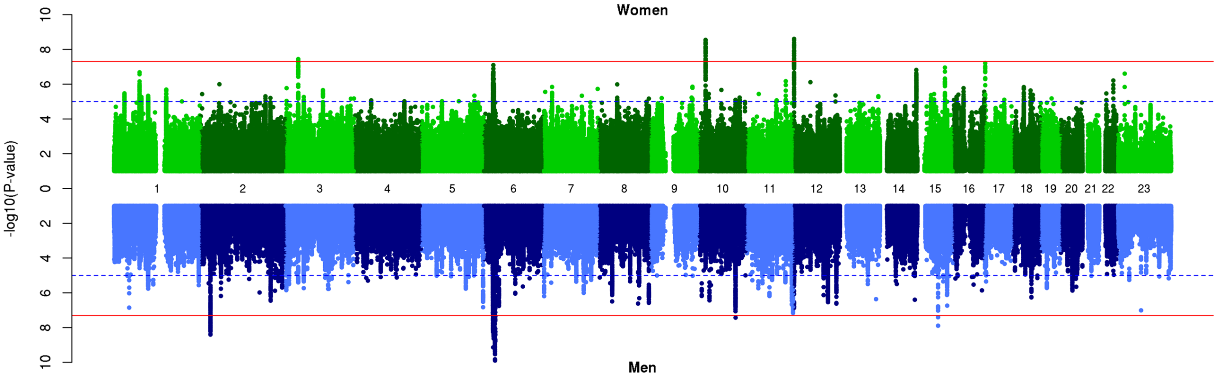


### h) Cross-Disorder SCZ-BIP-rMDD – European ancestry only

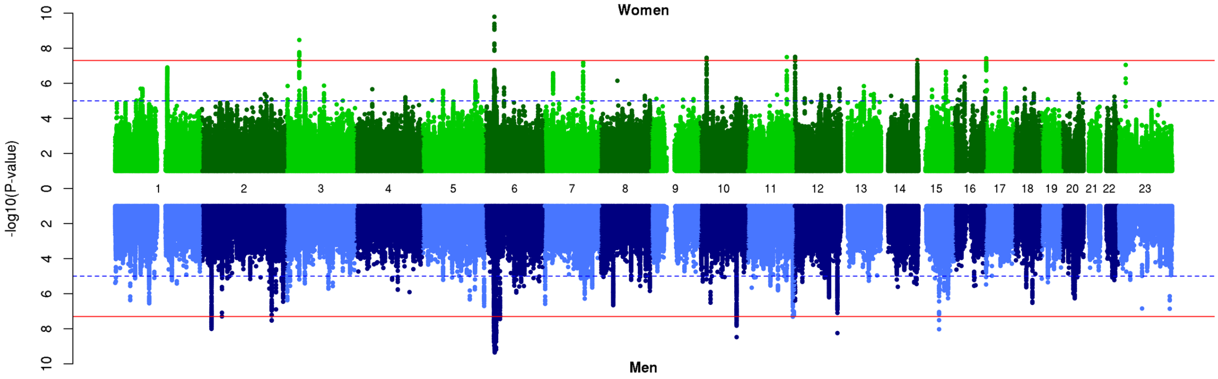

### i) Cross-Disorder SCZ-BIP-rMDD – European + East Asian ancestry

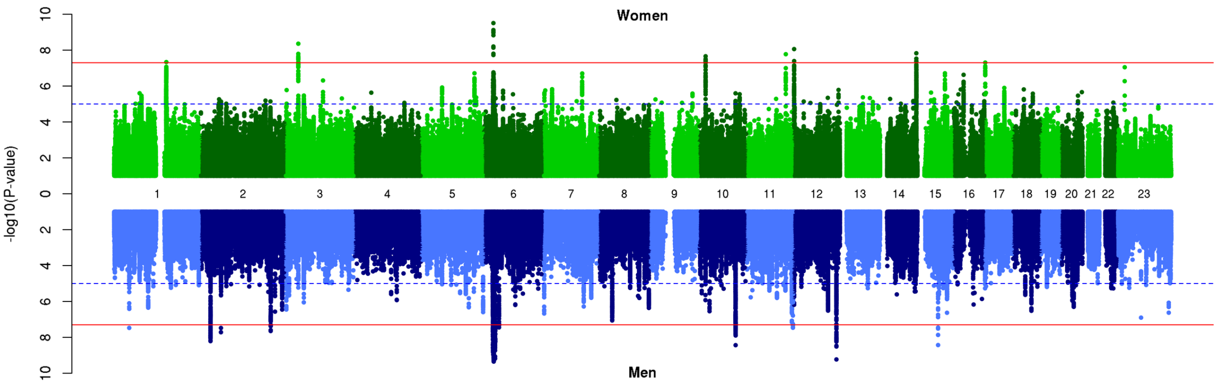

**Supplementary Figure 18. Scatter plots of female vs male associations in PGC**

The scatter plots show little correlation (R) between GWAS SNP main effect *p*-values from the two sexes, indicating the strength of association differed substantially between the two sexes.

Plots were generated using the plot package in R.

Abbreviations: BIP = Bipolar Disorder; (r)MDD = (recurrent) Major Depressive Disorder; SCZ = Schizophrenia; $R^2$ = proportion variance explained.

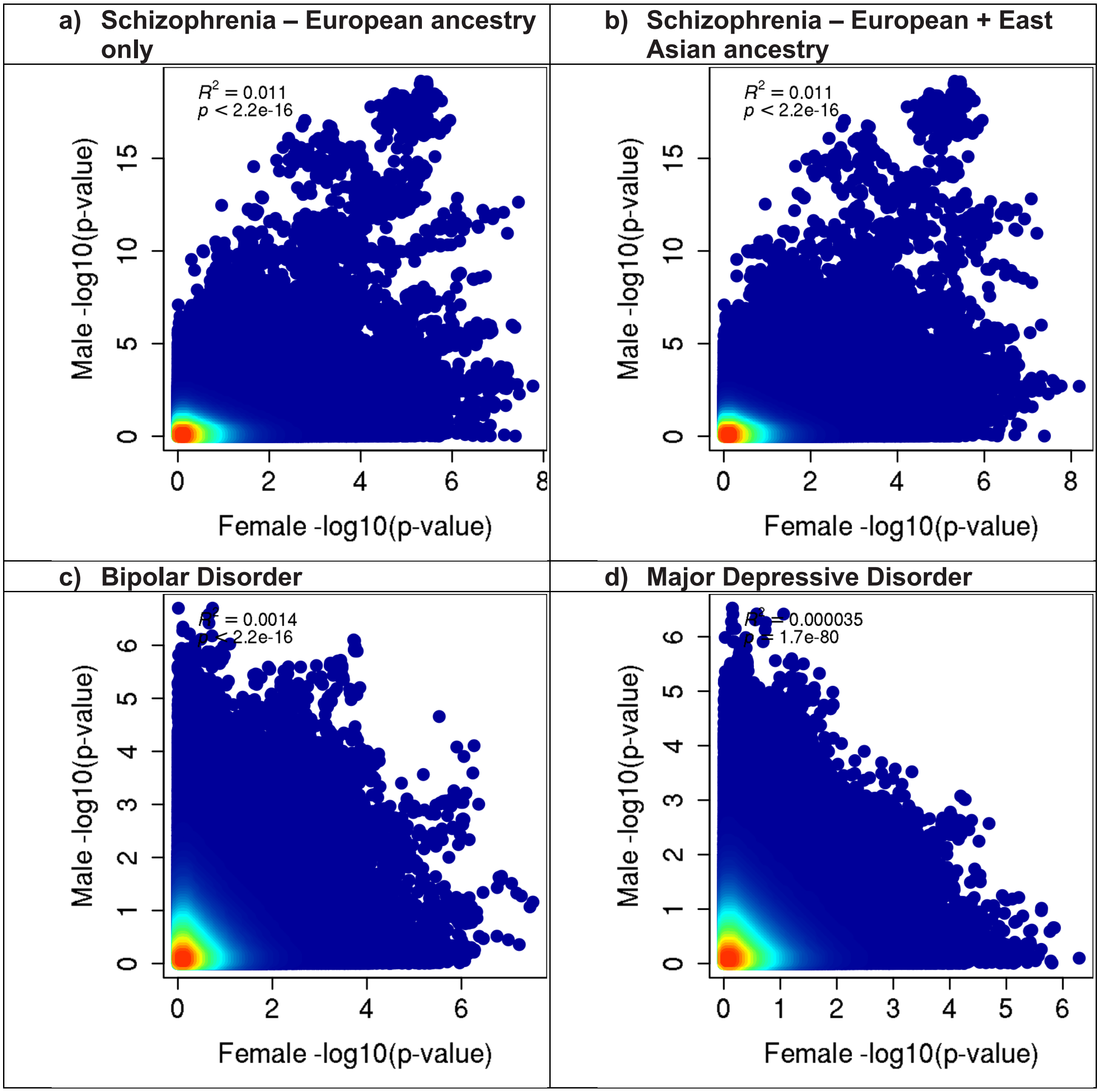

**e) Recurrent Major Depressive Disorder**

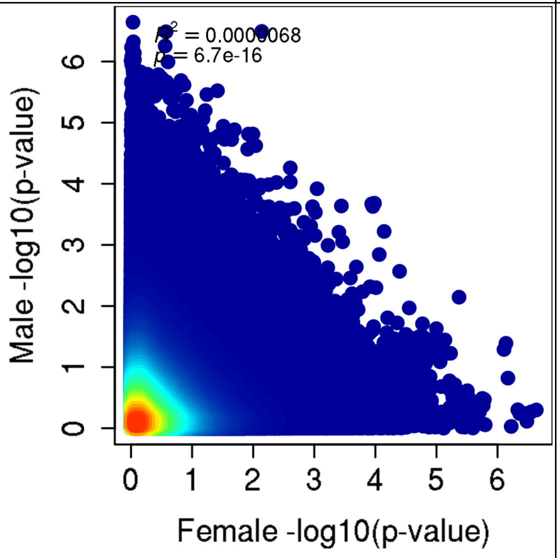


**f) Cross-Disorder SCZ-BIP-MDD – European ancestry only**

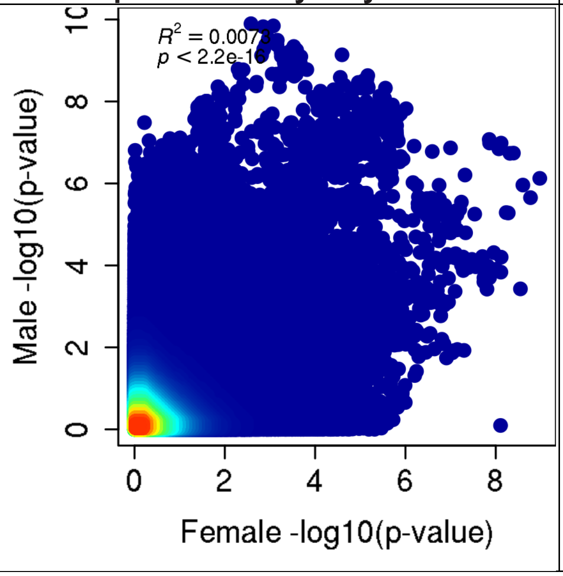


**g) Cross-Disorder SCZ-BIP-MDD – European + East Asian ancestry**

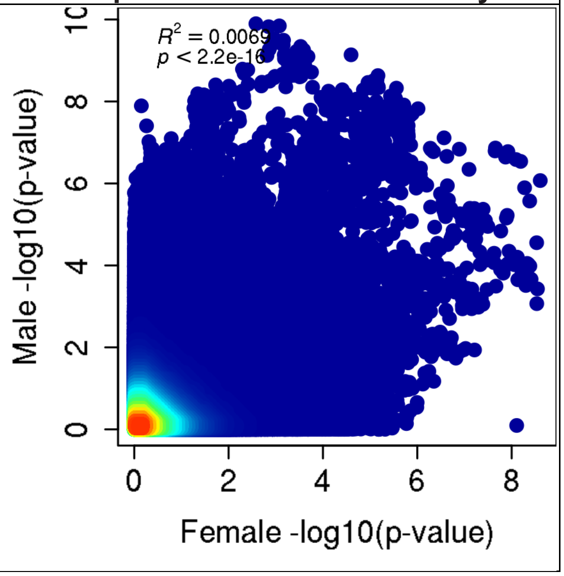

| h) Cross-Disorder SCZ-BIP-rMDD – European ancestry only | i) Cross-Disorder SCZ-BIP-rMDD – European + East Asian ancestry |
|---|---|
| 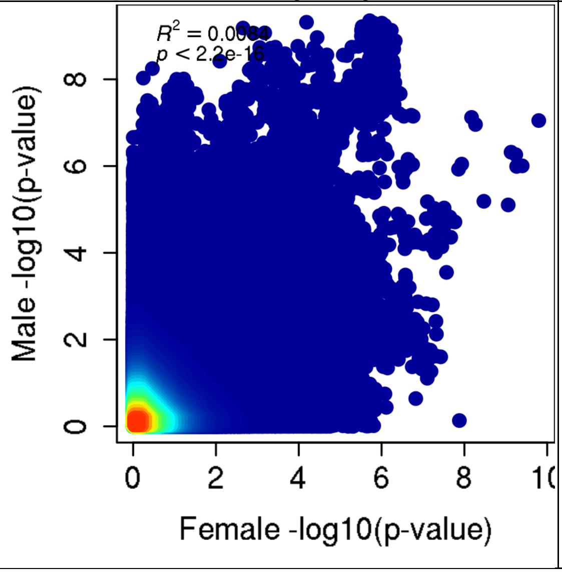 | 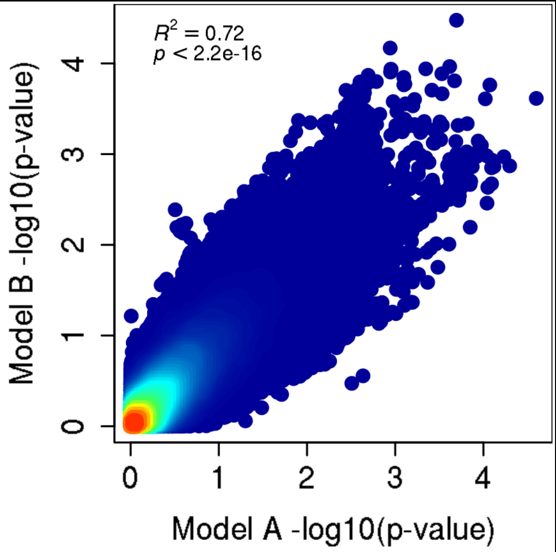 |

**Supplementary Figure 19. Quantile-Quantile (Q-Q) plots for GxS interaction in PGC**

The Q-Q plot is used to assess the number and magnitude of observed associations compared with the expectations under no association. The nature of deviations from the identity line provide clues whether the observed associations are true associations or may be due to for example population stratification or cryptic relatedness.

Abbreviations: BIP = Bipolar Disorder; (r)MDD = (recurrent) Major Depressive Disorder; SCZ = Schizophrenia

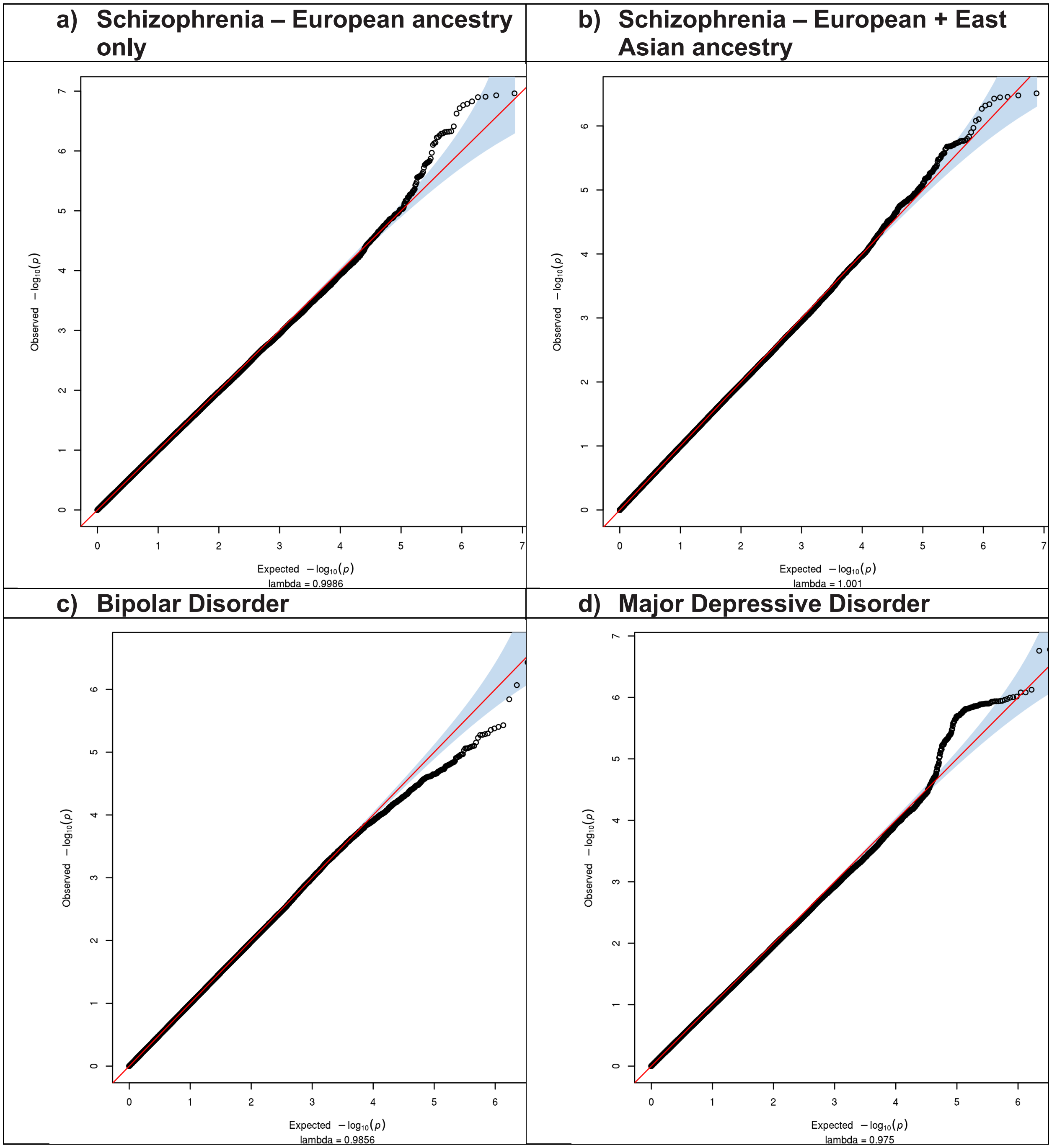

**e) Recurrent Major Depressive Disorder**

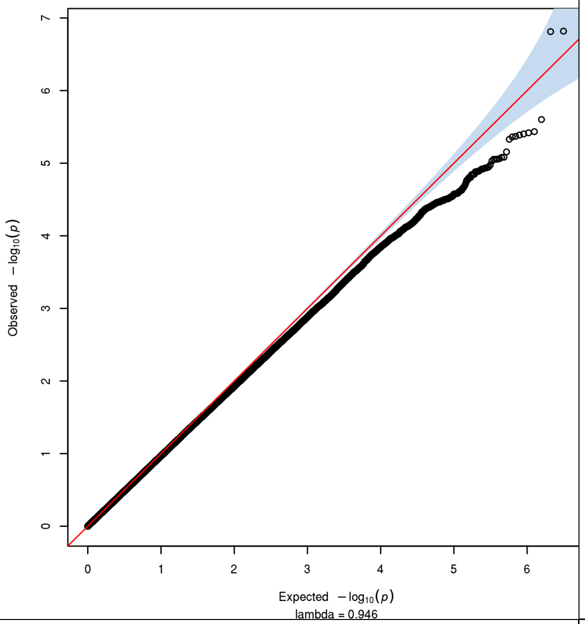


**f) Cross-Disorder SCZ-BIP-MDD – European ancestry only**

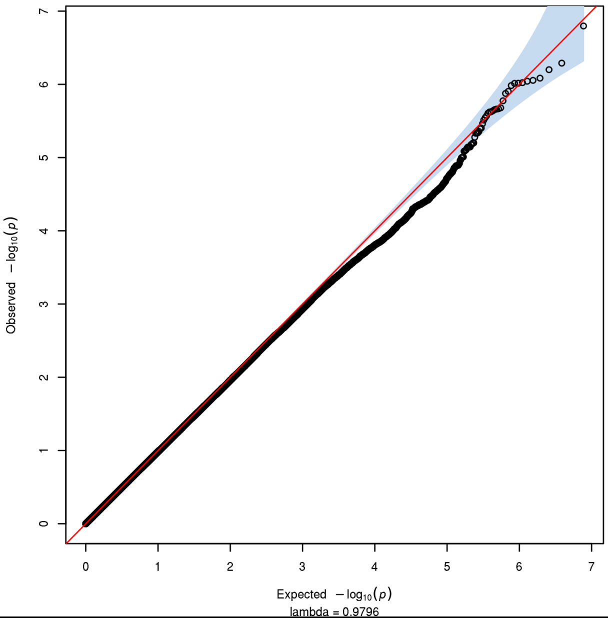


**g) Cross-Disorder SCZ-BIP-MDD – European + East Asian ancestry**

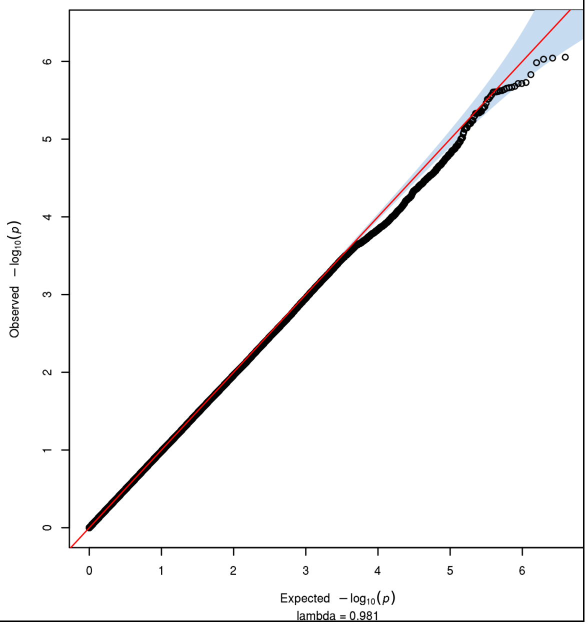

**h) Cross-Disorder SCZ-BIP-rMDD – European ancestry only**

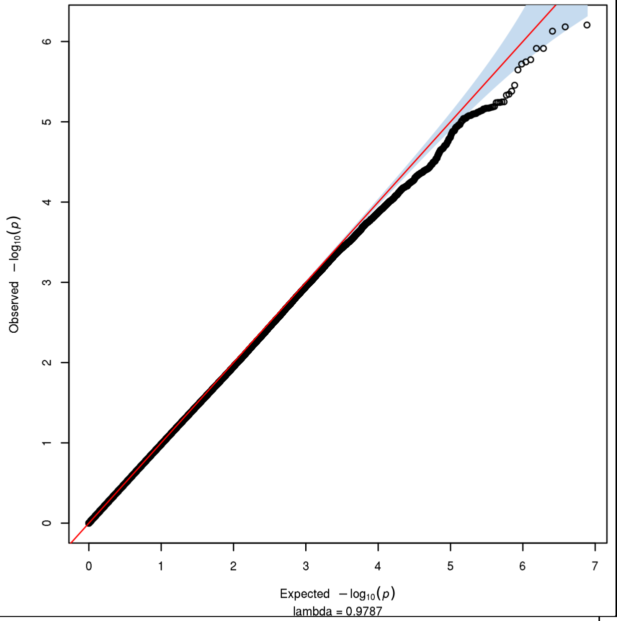


**i) Cross-Disorder SCZ-BIP-rMDD – European + East Asian ancestry**

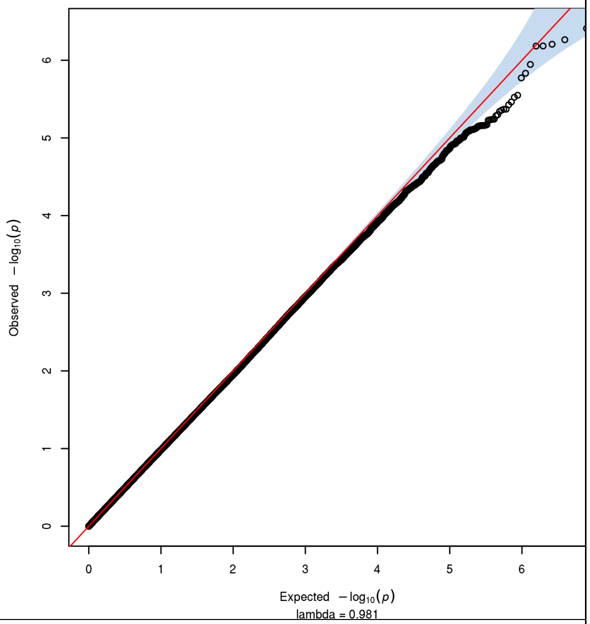


**j) Omnibus Test SCZ-BIP-MDD – European ancestry only**

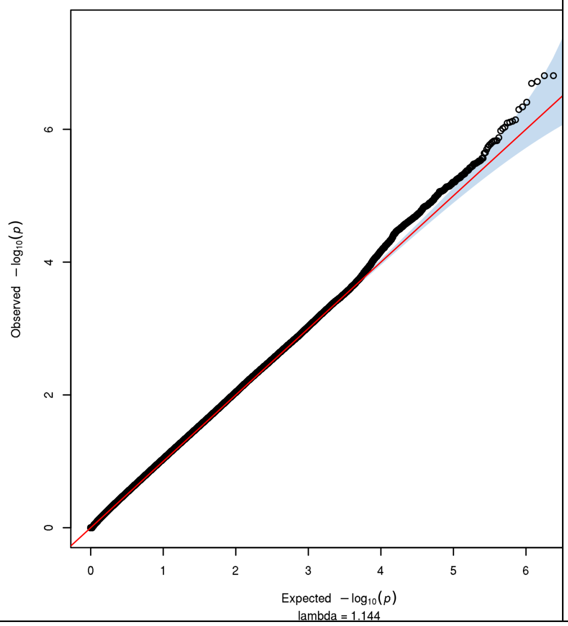


**k) Omnibus Test SCZ-BIP-MDD – European + East Asian ancestry**

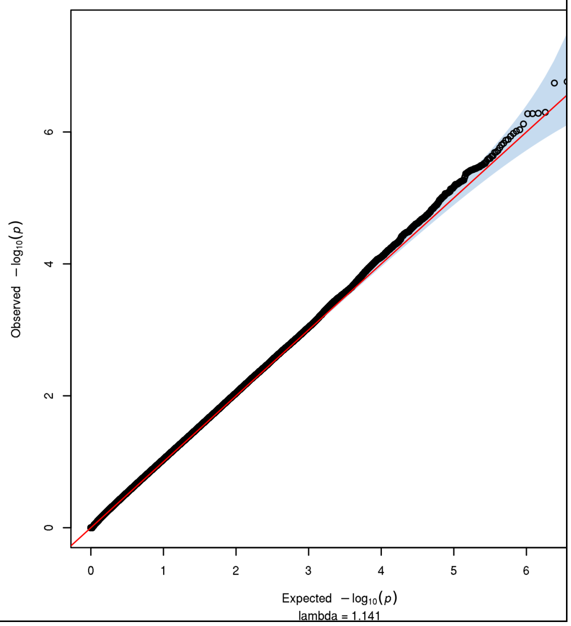

| l) Omnibus Test SCZ-BIP-rMDD – European ancestry only | m) Omnibus Test SCZ-BIP-rMDD – European + East Asian ancestry |
|---|---|

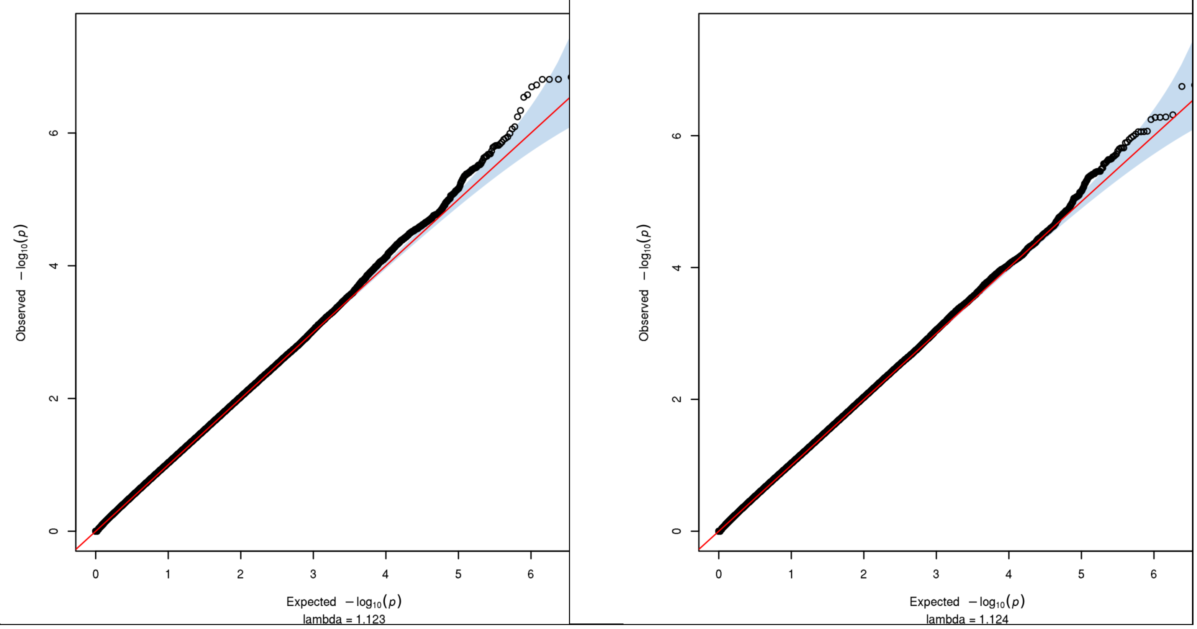

**Supplementary Figure 20. Manhattan plots of the GxS interaction GWAS in PGC**

Negative log10-transformed p-values for each variant (each dot) (y-axis) are plotted by chromosomal position (x-axis). The red and blue lines represent the thresholds for genome-wide significant association ($p = 5\times10^{-8}$) and suggestive association ($p = 1\times10^{-5}$), respectively. P-values for X chromosome (23) model B (alleles: females 0, 1, or 2; males 0 or 1) are included.

Abbreviations: BIP = Bipolar Disorder; (r)MDD = (recurrent) Major Depressive Disorder; SCZ = Schizophrenia

### a) Schizophrenia – European ancestry only

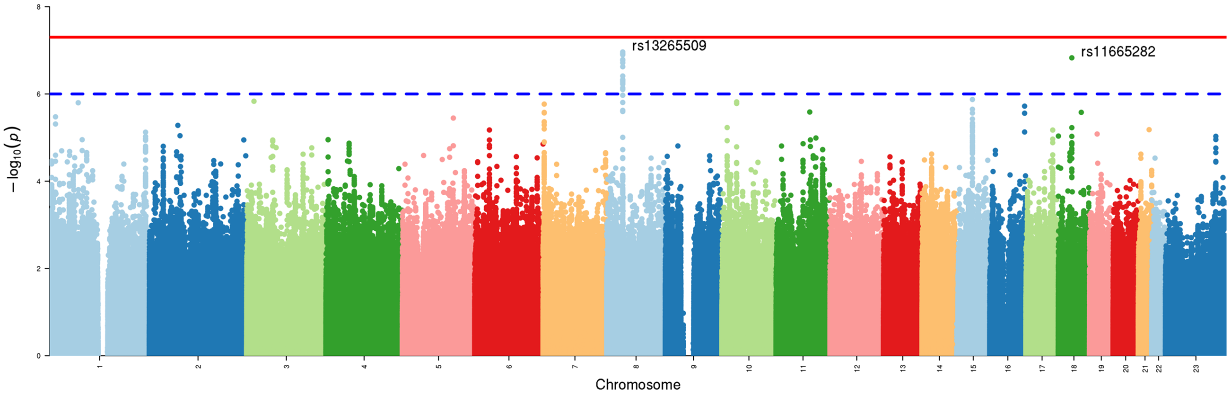


### b) Schizophrenia – European + East Asian ancestry

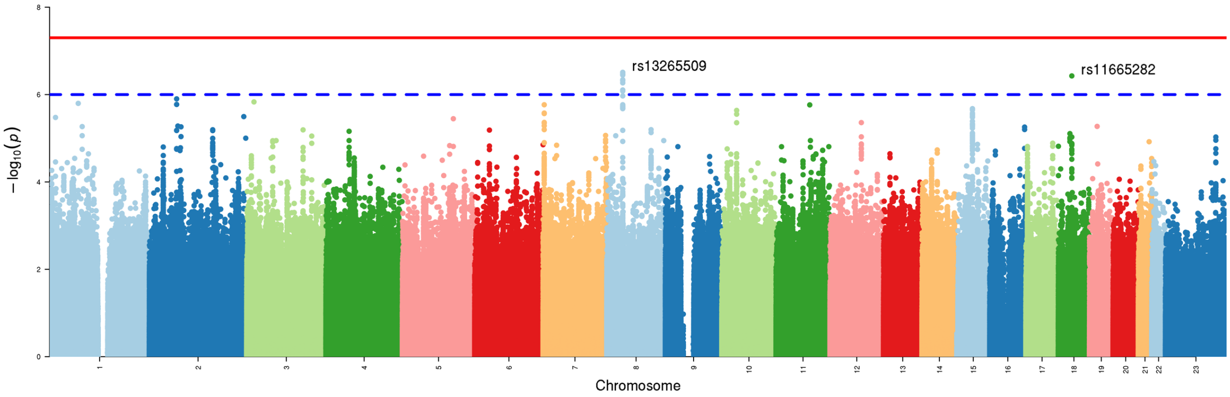

### c) Bipolar Disorder

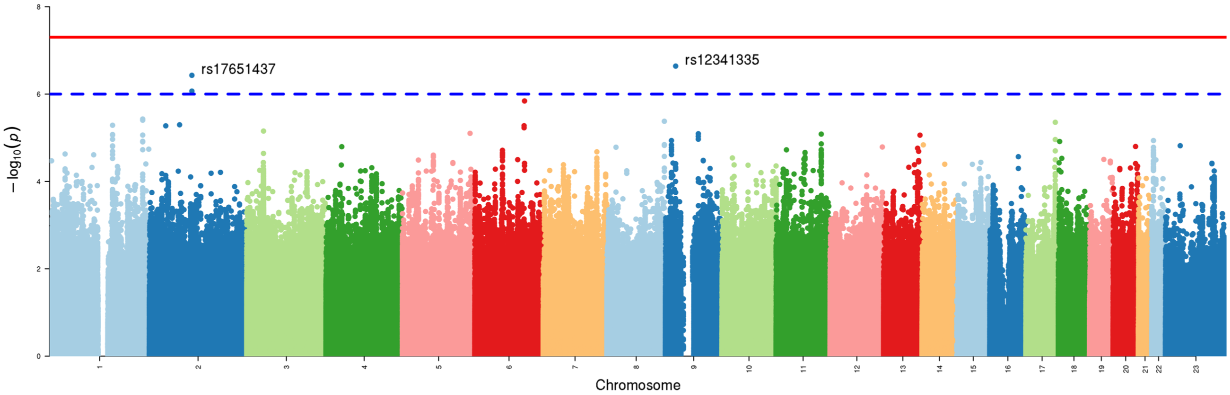


### d) Major Depressive Disorder

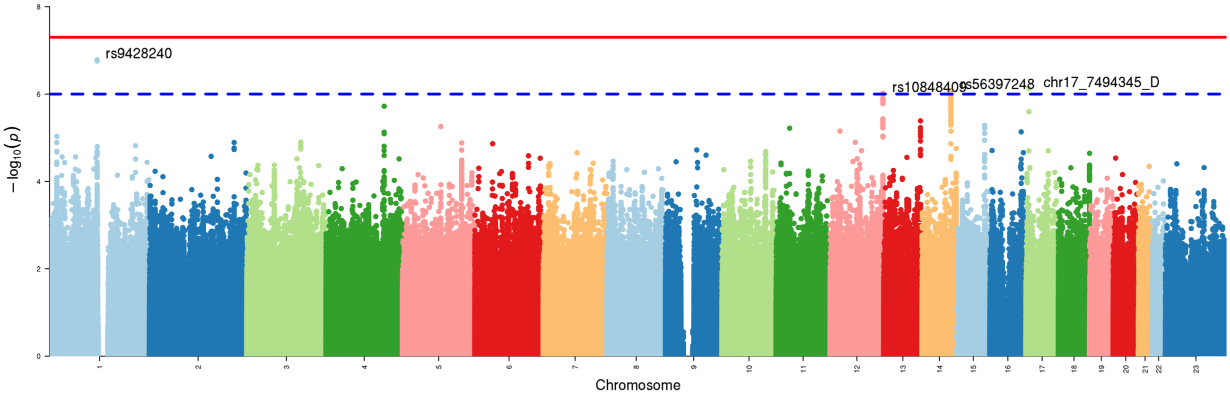


### e) Recurrent Major Depressive Disorder

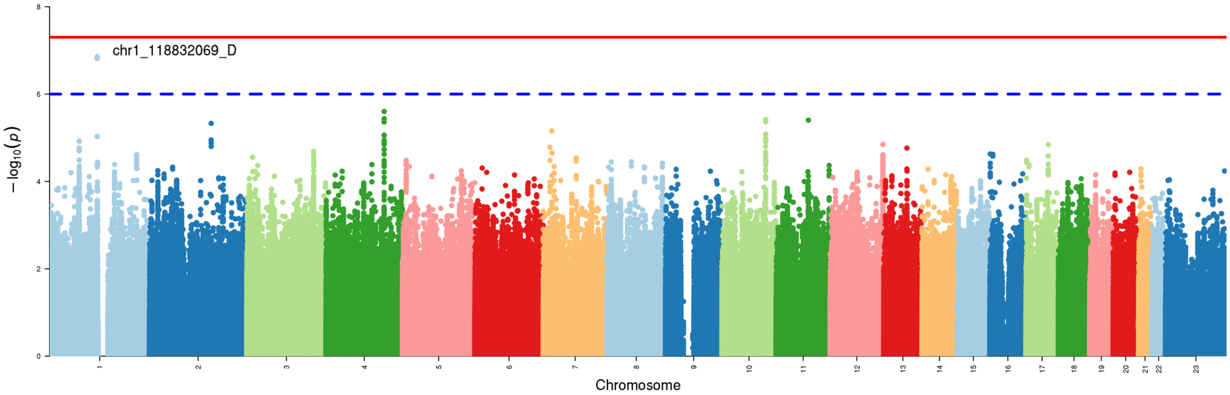

### f) Cross-Disorder SCZ-BIP-MDD – European ancestry only

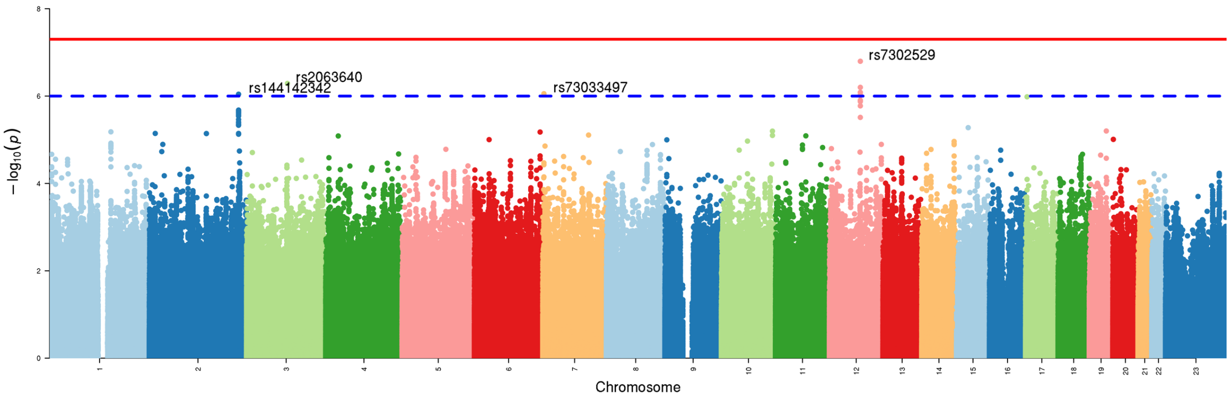


### g) Cross-Disorder SCZ-BIP-MDD – European + East Asian ancestry

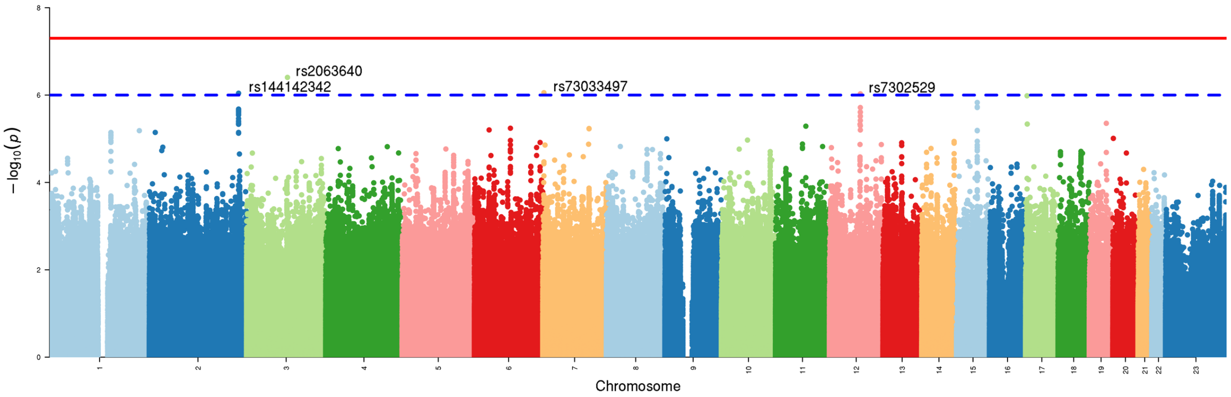


### h) Cross-Disorder SCZ-BIP-rMDD – European ancestry only

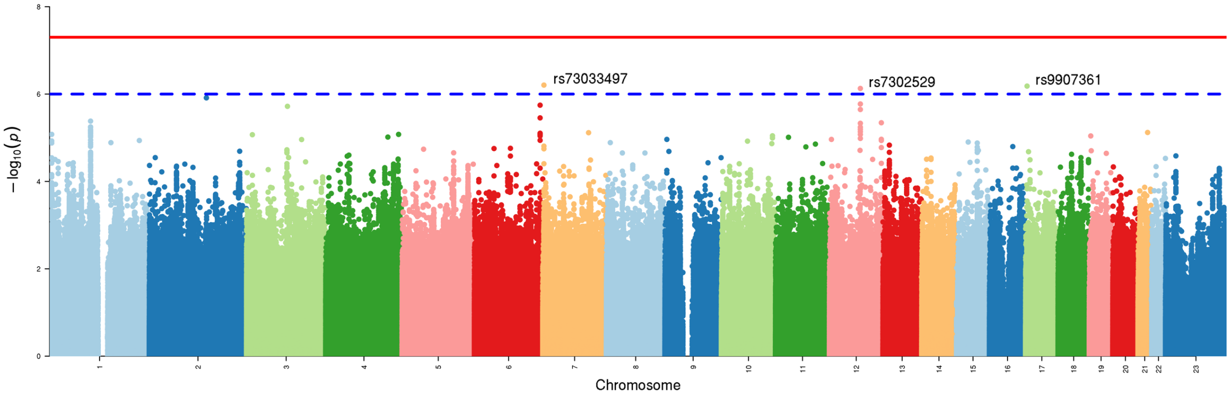

### i) Cross-Disorder SCZ-BIP-rMDD – European + East Asian ancestry

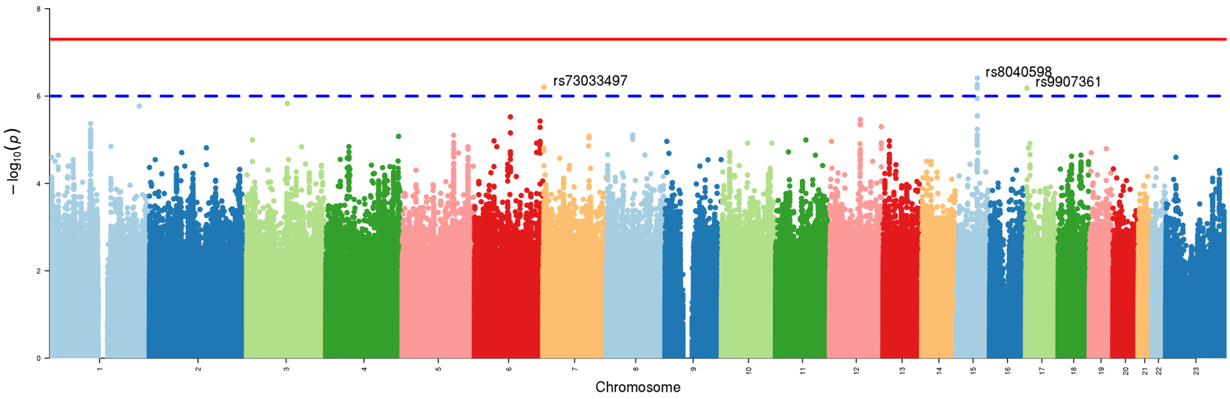


### j) Omnibus Test SCZ-BIP-MDD – European ancestry only

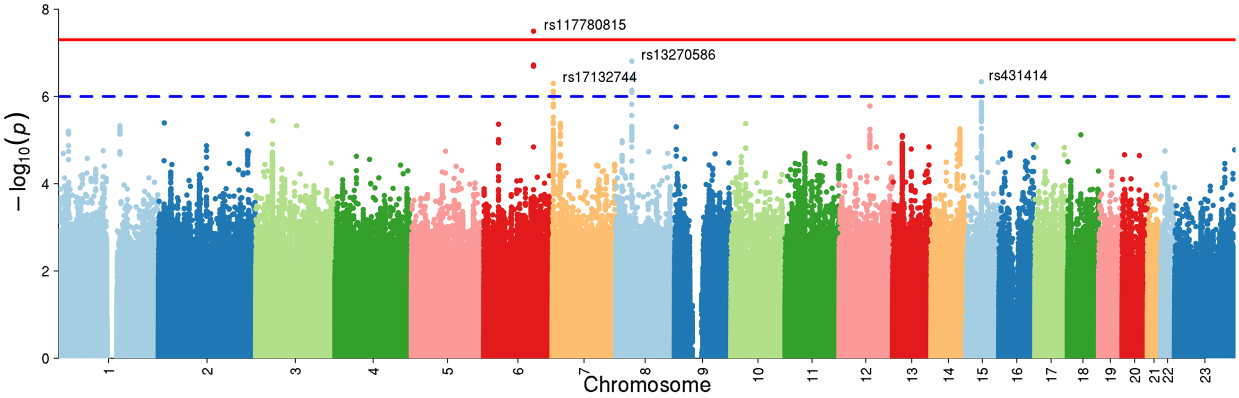


### k) Omnibus Test SCZ-BIP-MDD – European + East Asian ancestry

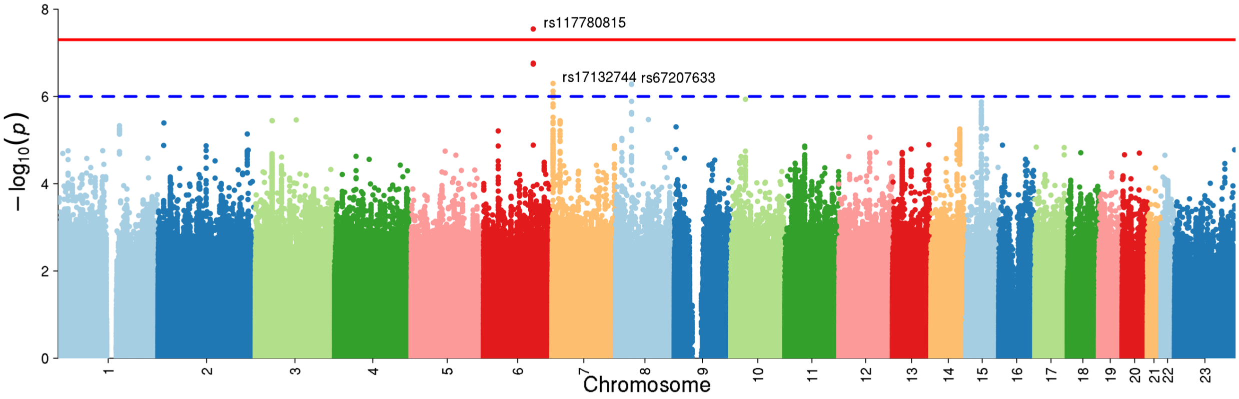

### l) Omnibus Test SCZ-BIP-rMDD – European ancestry only

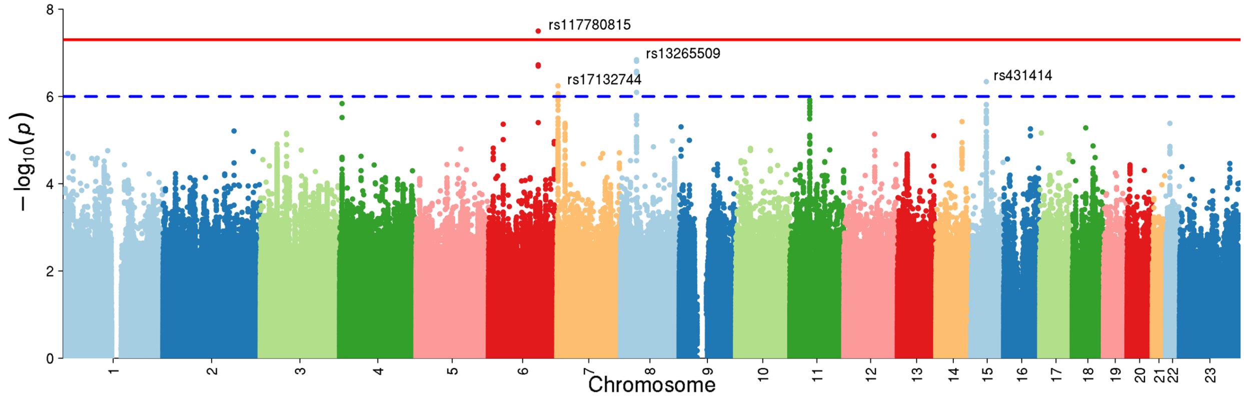


### m) Omnibus Test SCZ-BIP-rMDD – European + East Asian ancestry

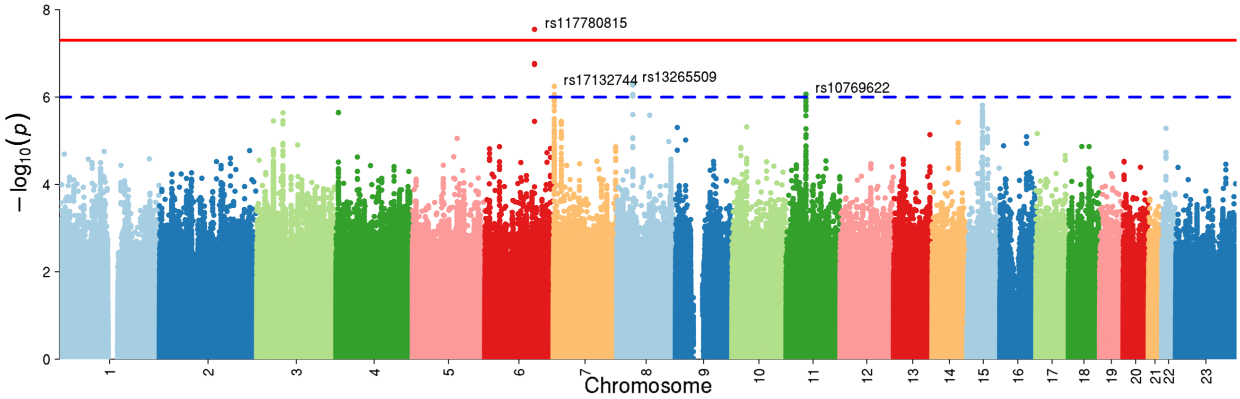

**Supplementary Figure 21. LocusZoom plots for loci with GxS interaction in PGC**

Plots were generated using the LocusZoom 1.4 Standalone application (49) for loci with GxS interaction $p < 1\times10^{-6}$.

Abbreviations: chr = chromosome; cM = centimorgans; Mb = megabases; $r^2$ = linkage disequilibrium level; BIP = Bipolar Disorder; (r)MDD = (recurrent) Major Depressive Disorder; SCZ = Schizophrenia

### a) Schizophrenia – European ancestry only

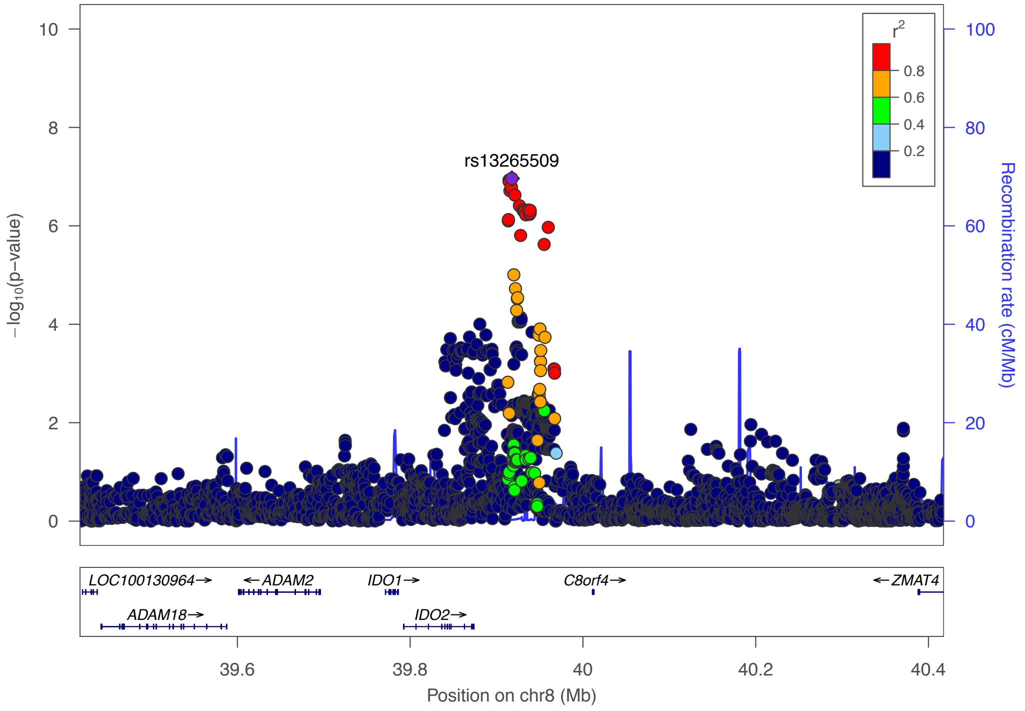

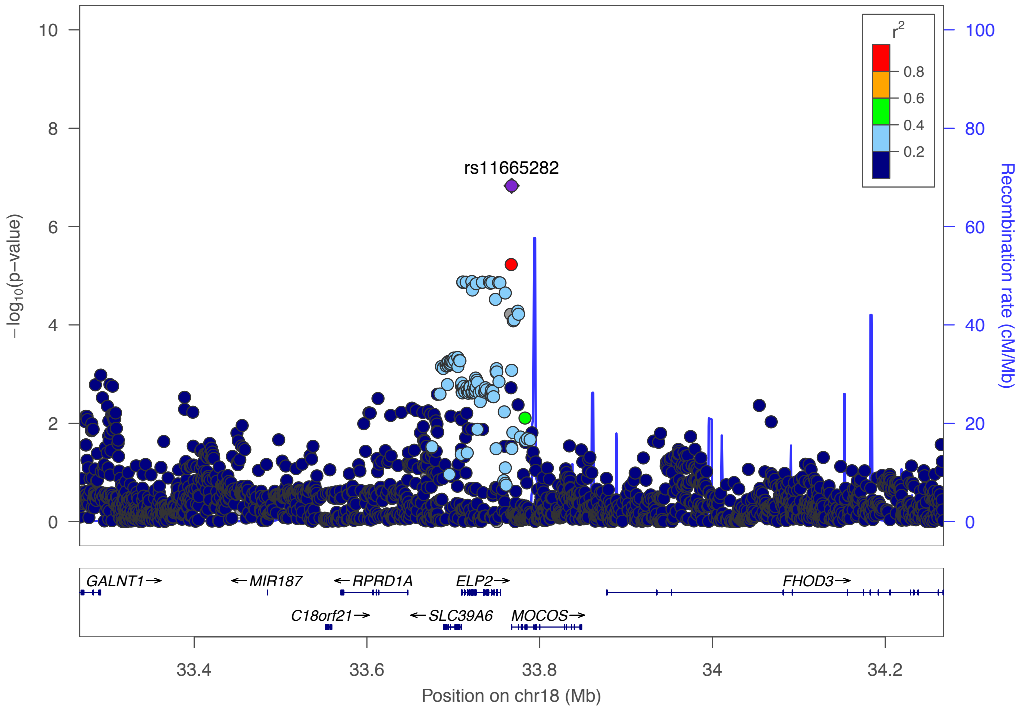


### b) Schizophrenia – European + East Asian ancestry

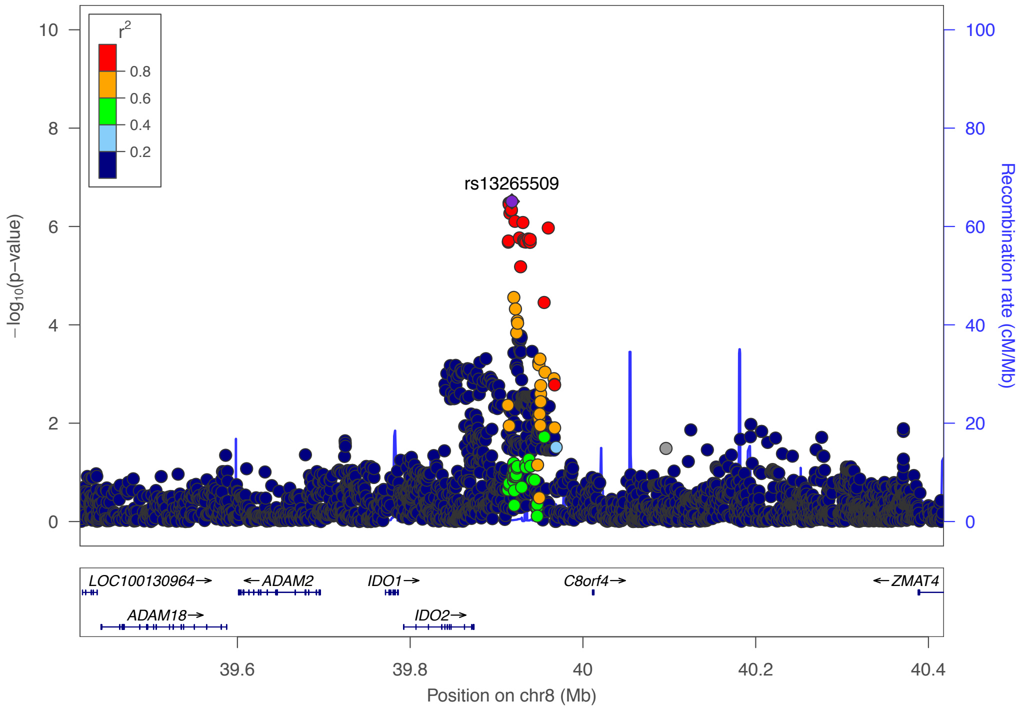

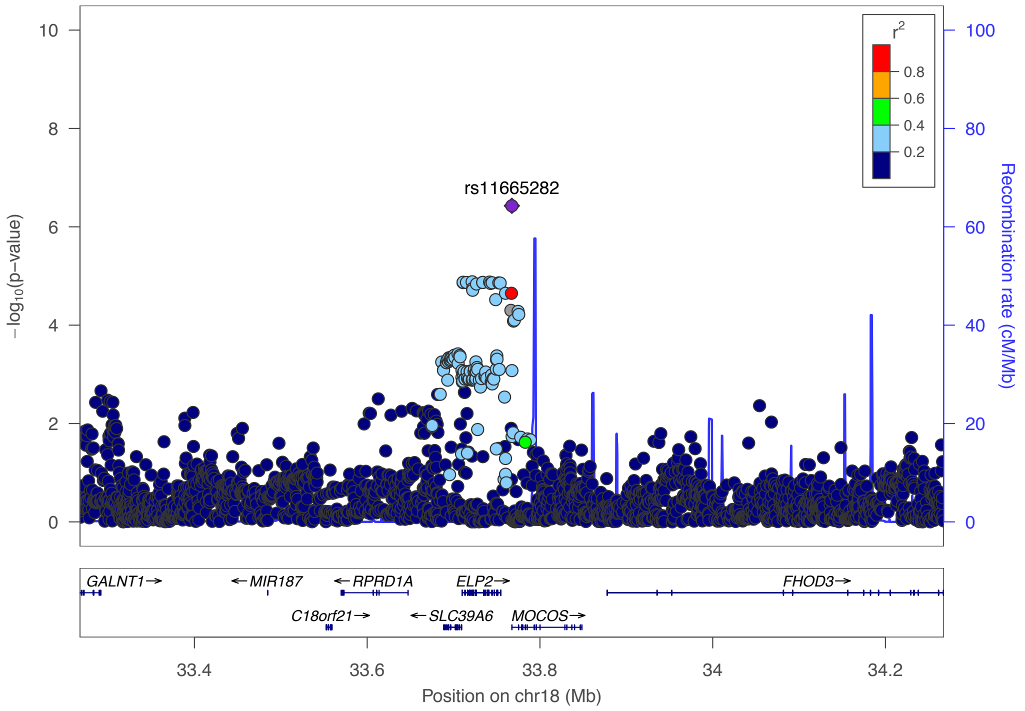


s

### c) Bipolar Disorder

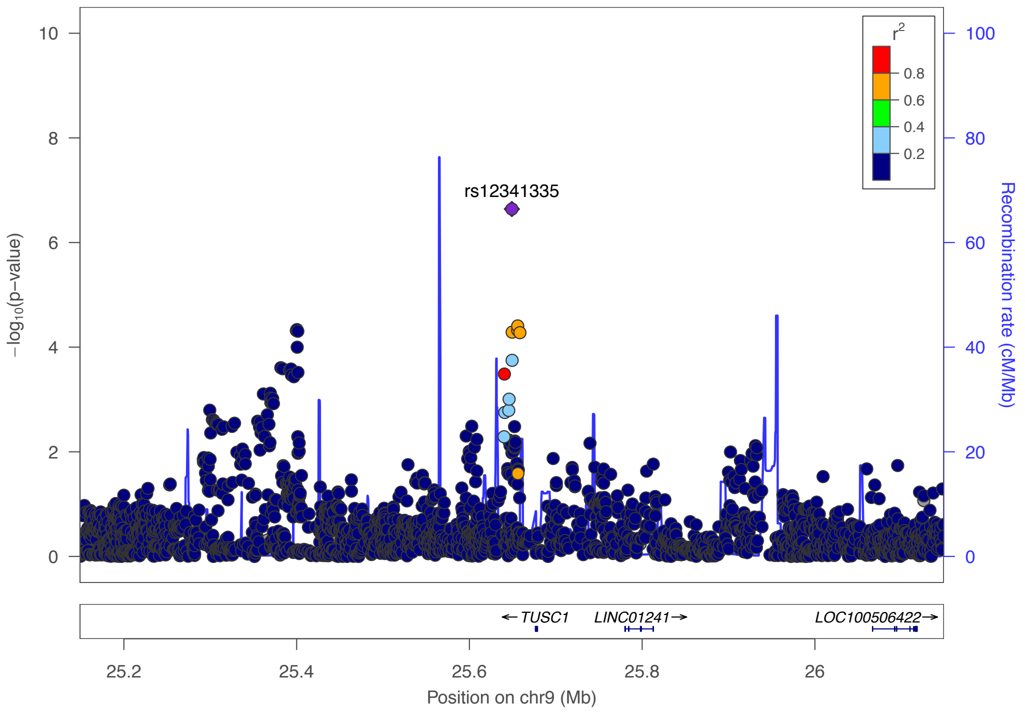

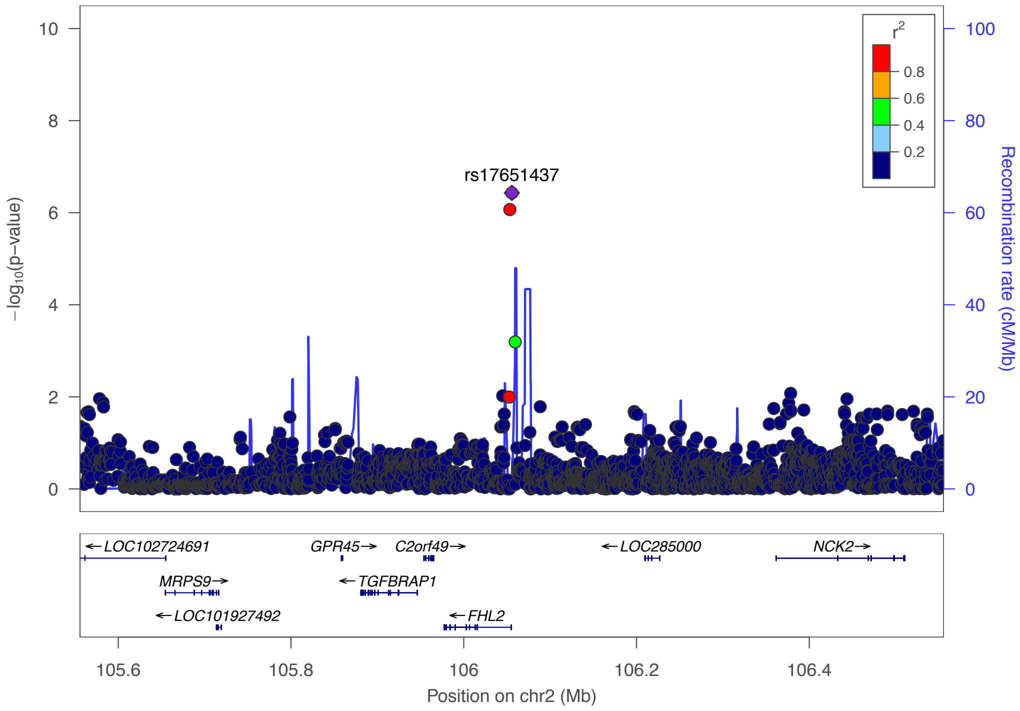


### d) Major Depressive Disorder

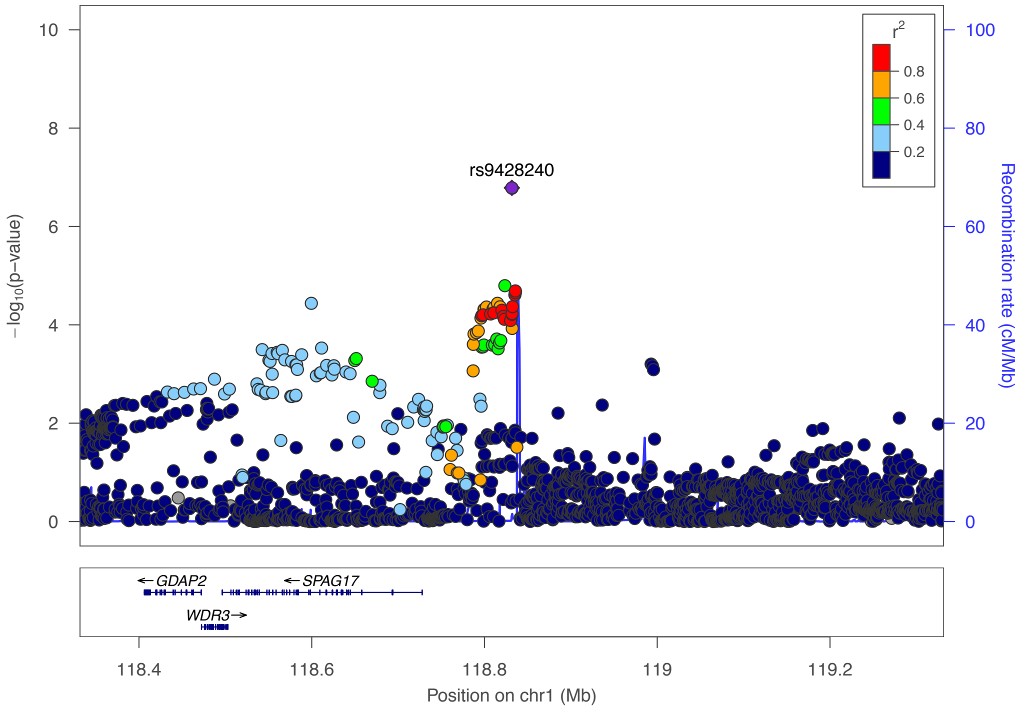

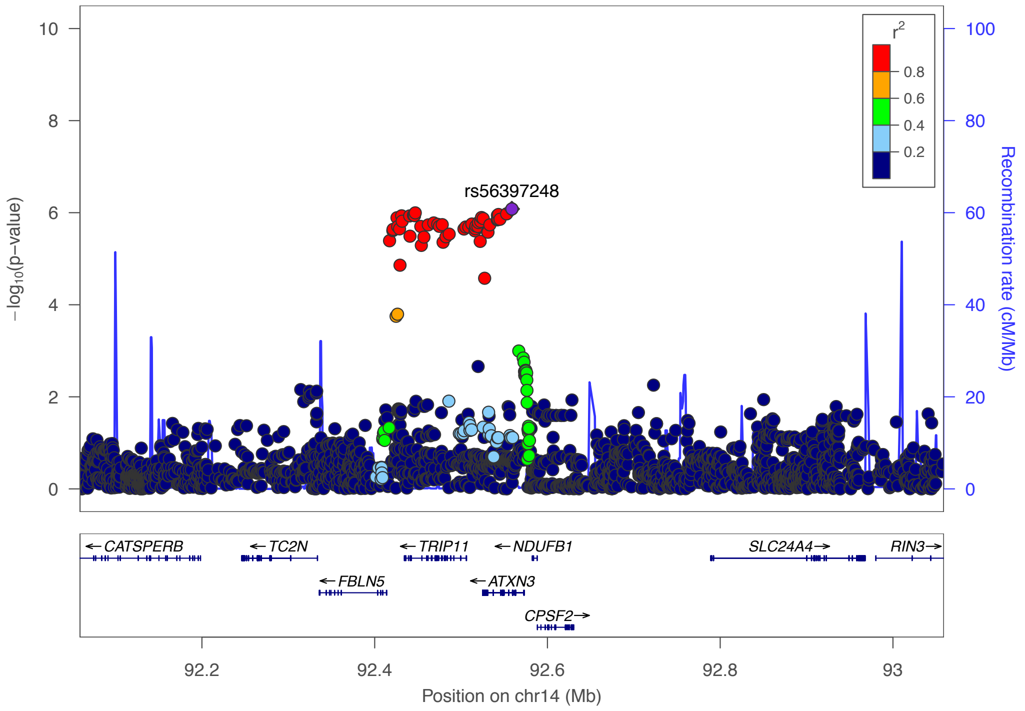


e) **Recurrent Major Depressive Disorder**

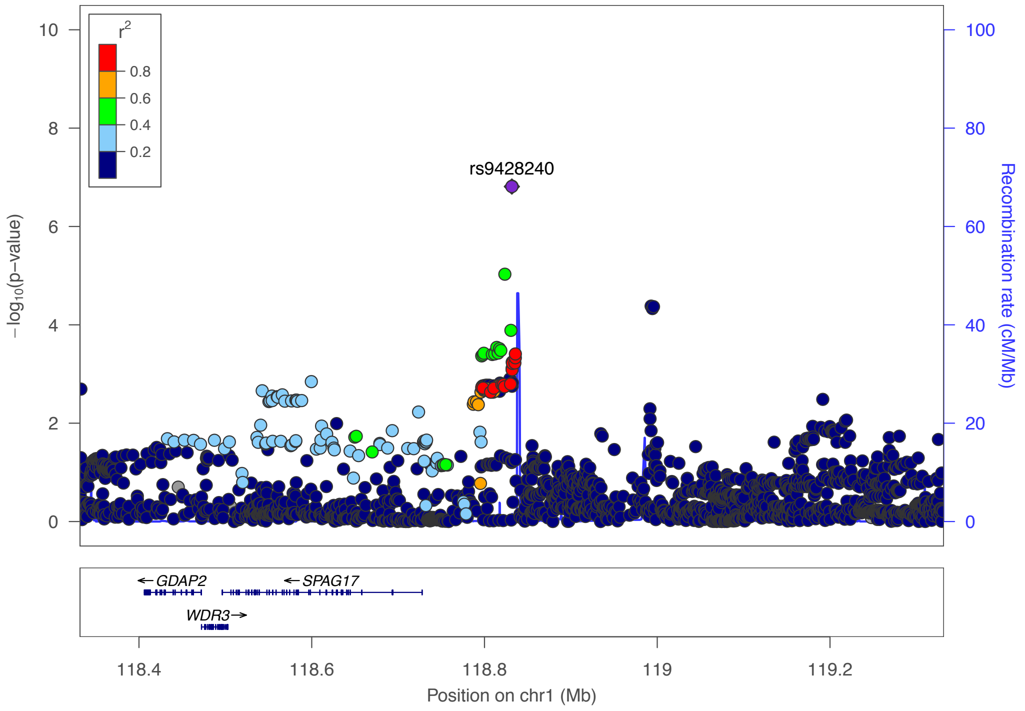

### f) Cross-Disorder SCZ-BIP-MDD – European ancestry only

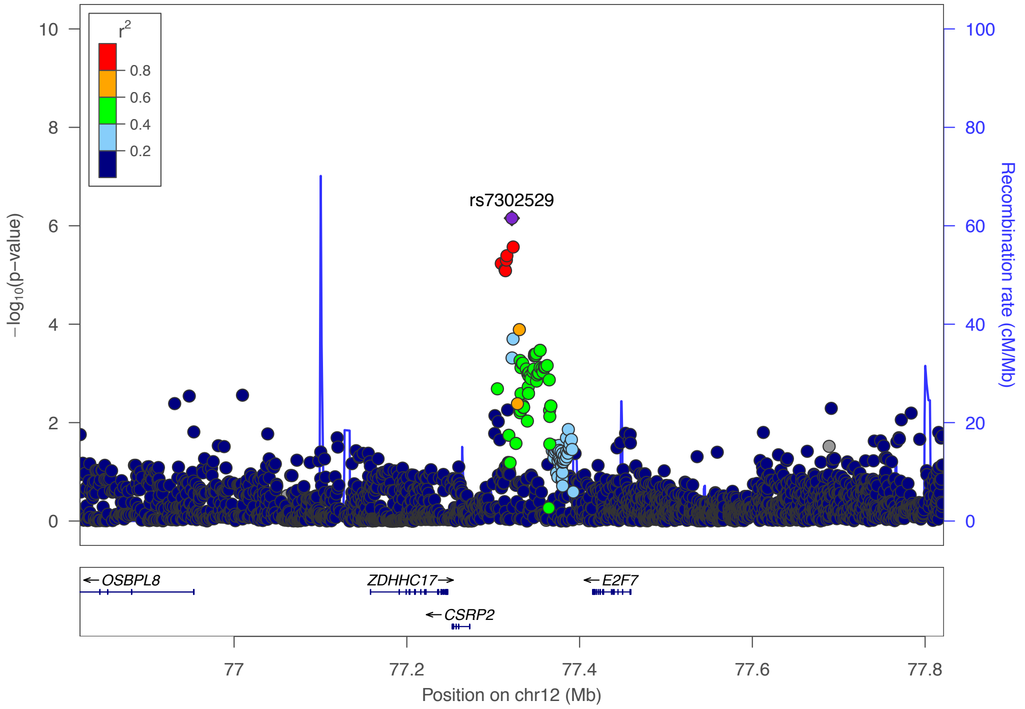


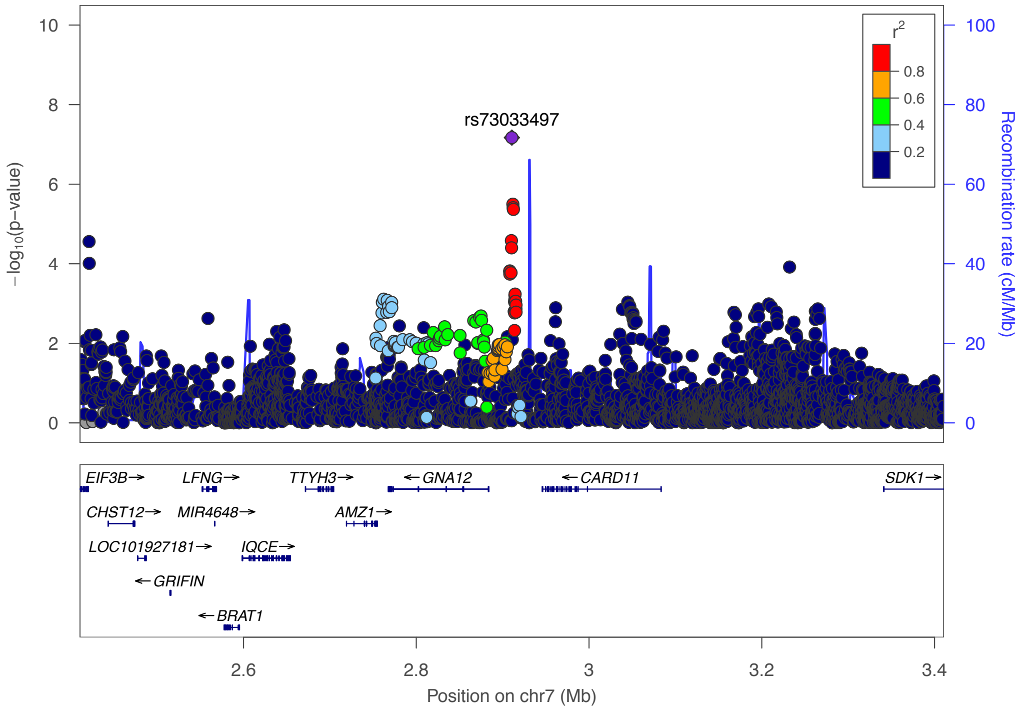

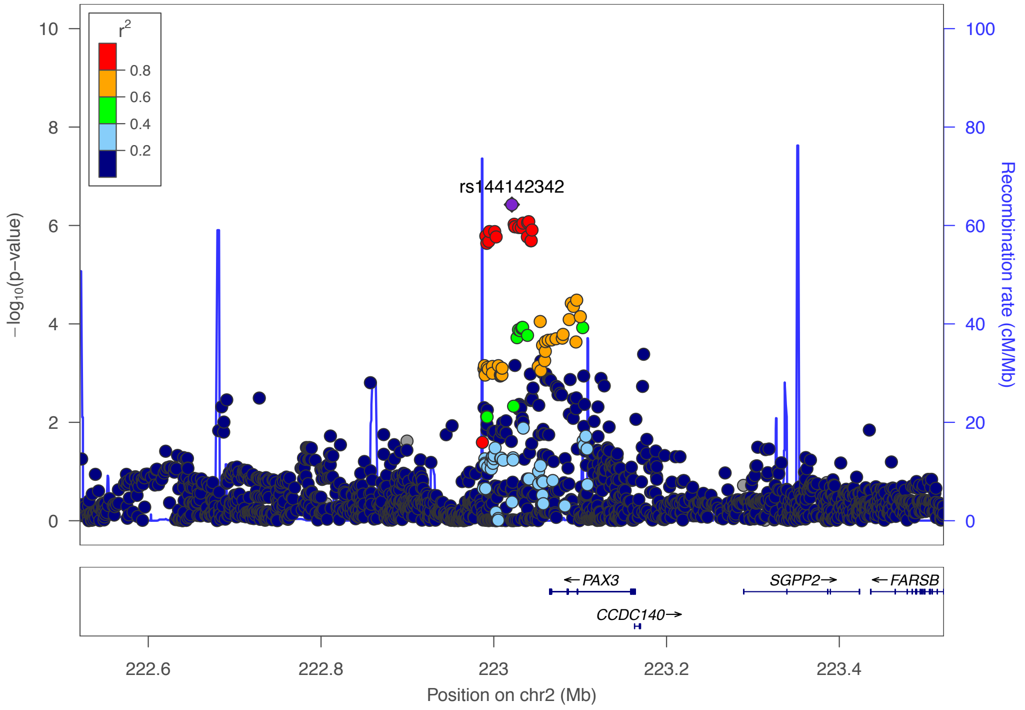


### g) Cross-Disorder SCZ-BIP-MDD – European + East Asian ancestry

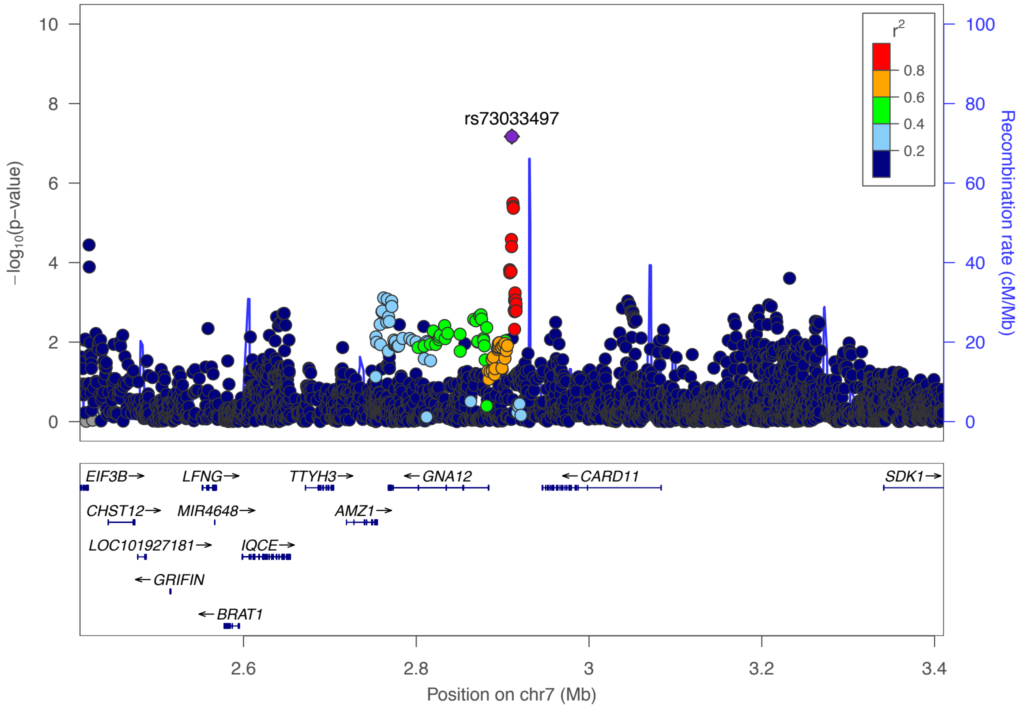

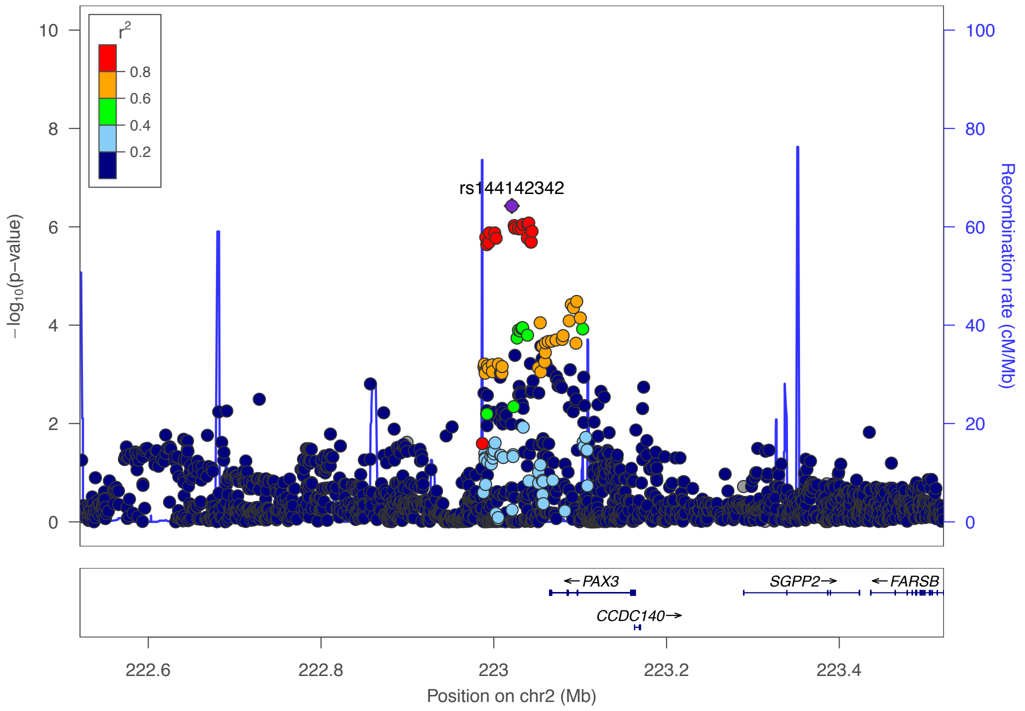
r2
0.8
0.6
0.4
0.2
rs144142342
−log10(p−value)
Recombination rate (cM/Mb)
PAX3
SGPP2
FARSB
CCDC140
222.6
222.8
223
223.2
223.4
Position on chr2 (Mb)

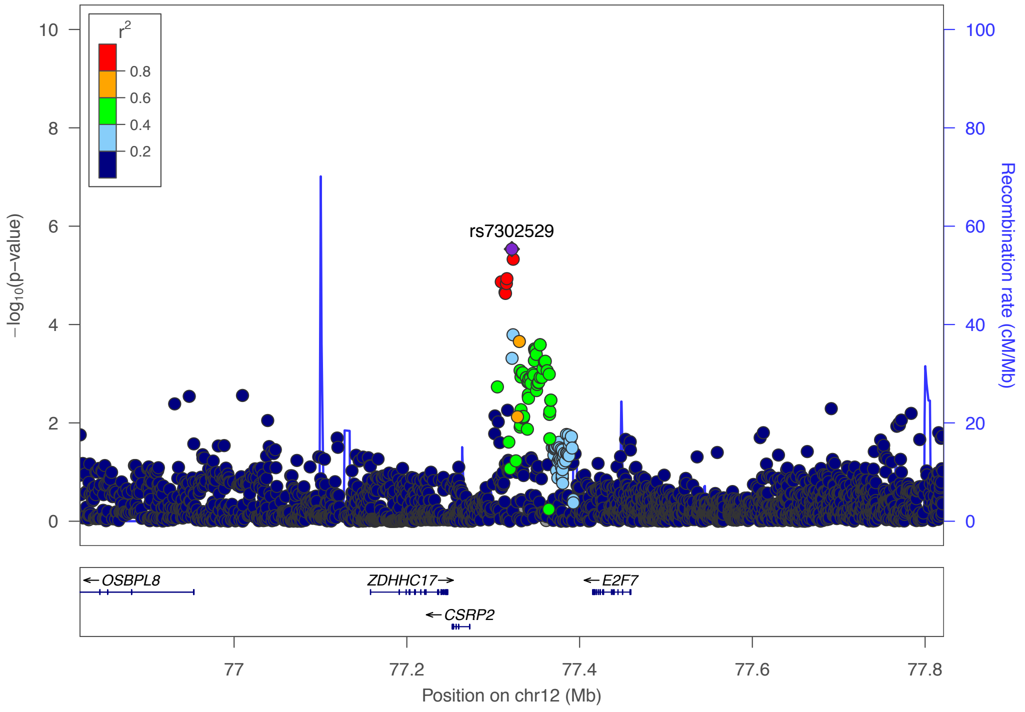
r2
0.8
0.6
0.4
0.2
rs7302529
−log10(p−value)
Recombination rate (cM/Mb)
OSBPL8
ZDHHC17
E2F7
CSRP2
77
77.2
77.4
77.6
77.8
Position on chr12 (Mb)

### h) Cross-Disorder SCZ-BIP-rMDD – European ancestry only

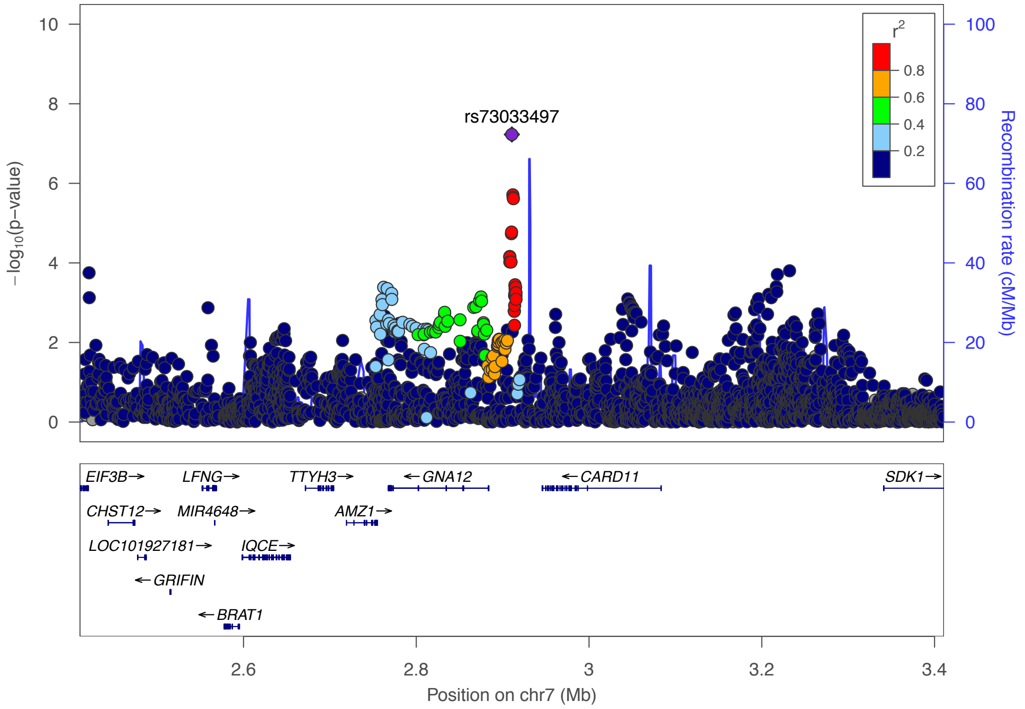


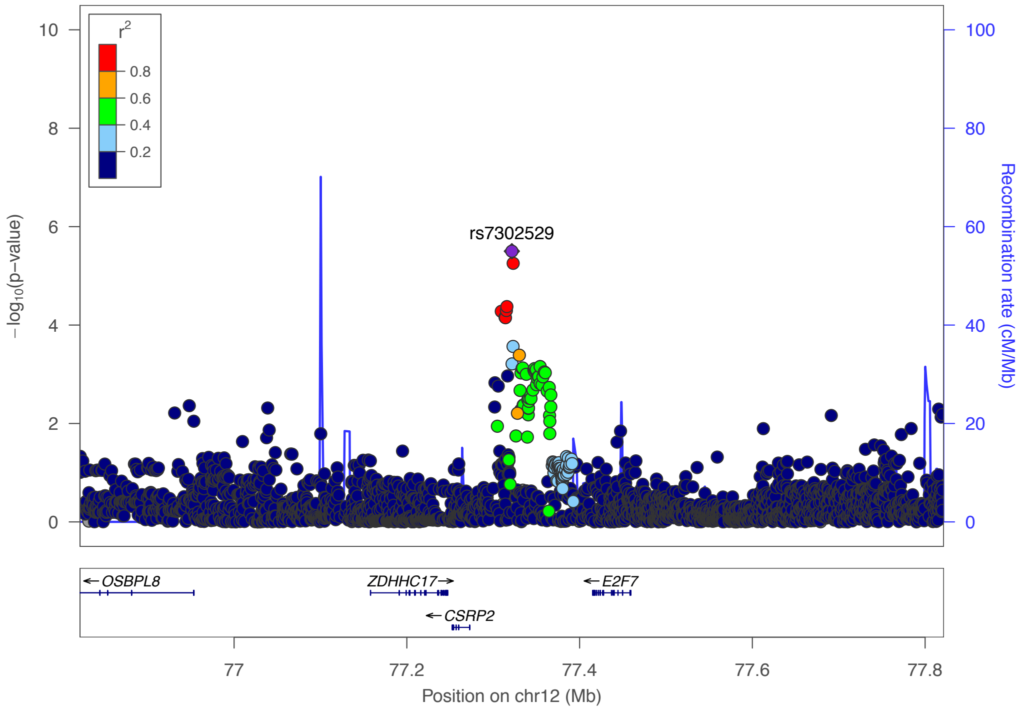

### i) Cross-Disorder SCZ-BIP-rMDD – European + East Asian ancestry

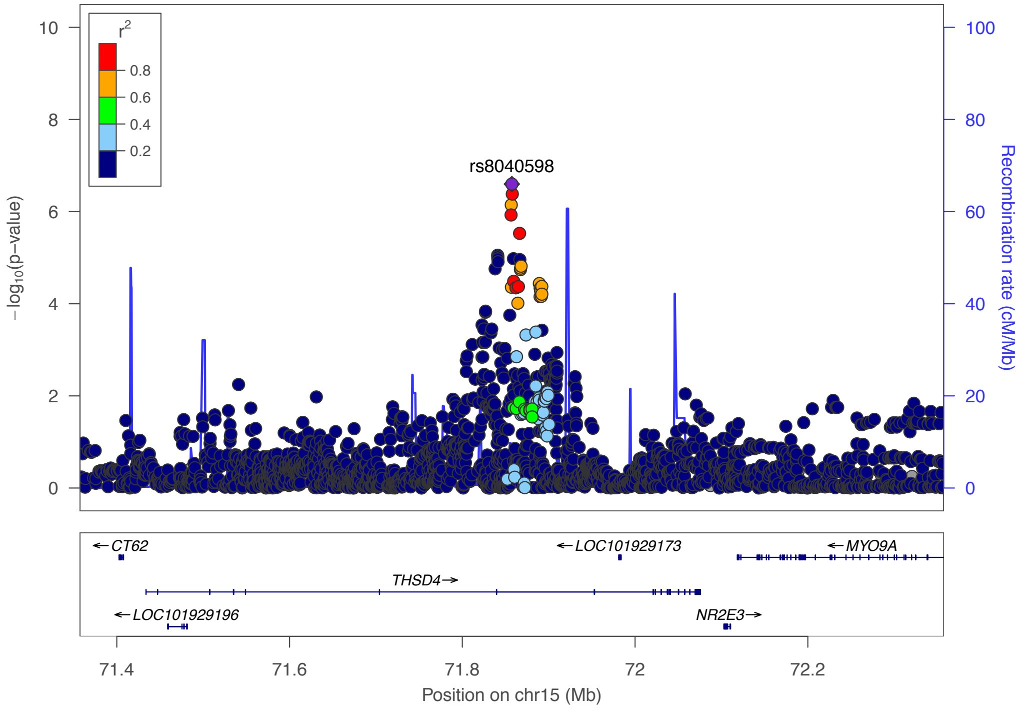


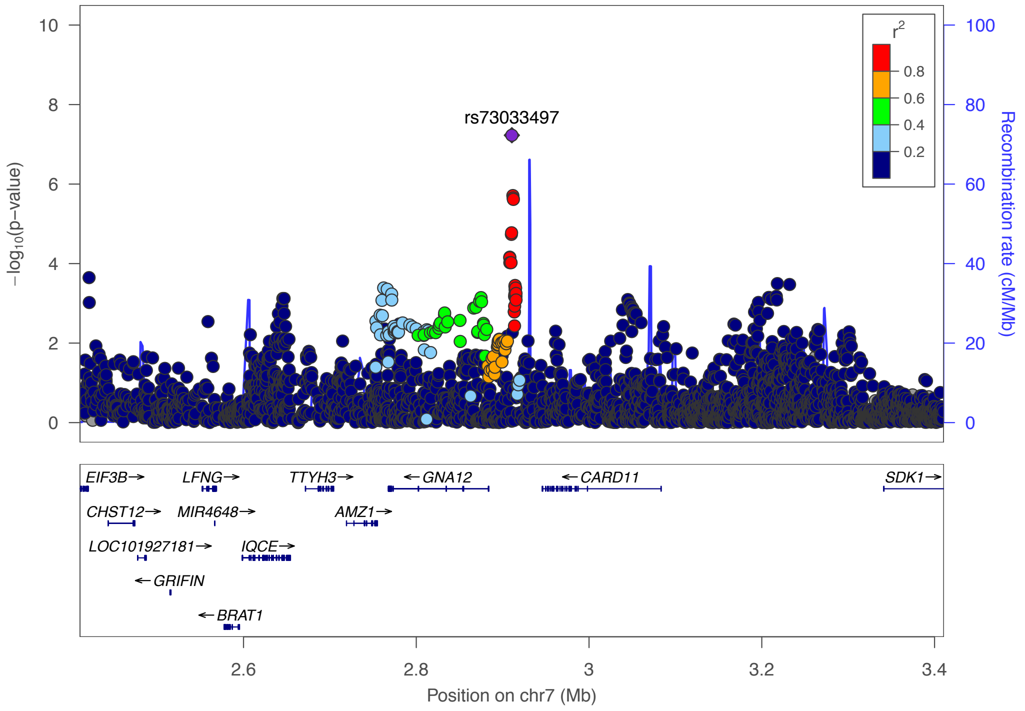

### j) Omnibus Test SCZ-BIP-MDD – European ancestry

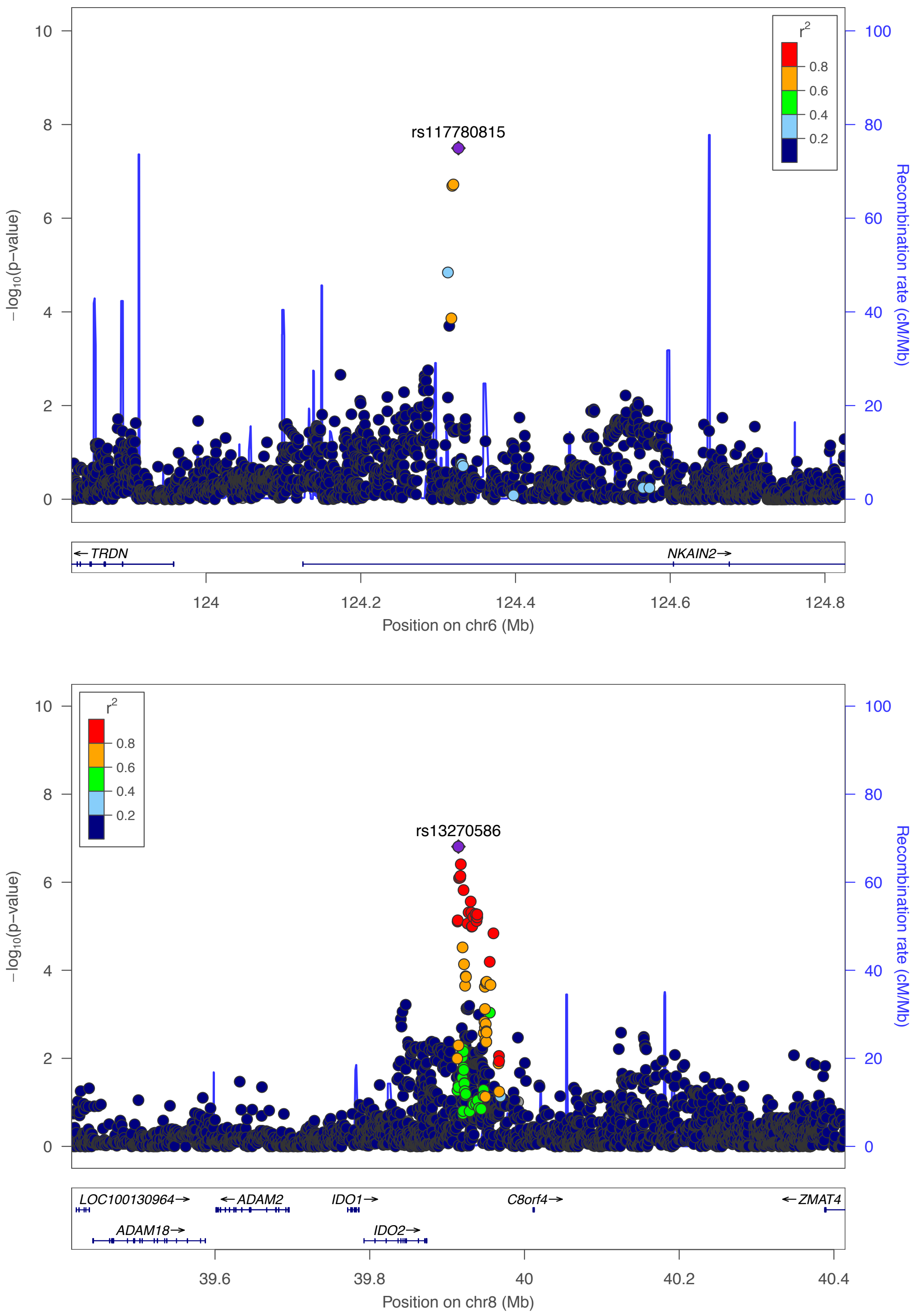

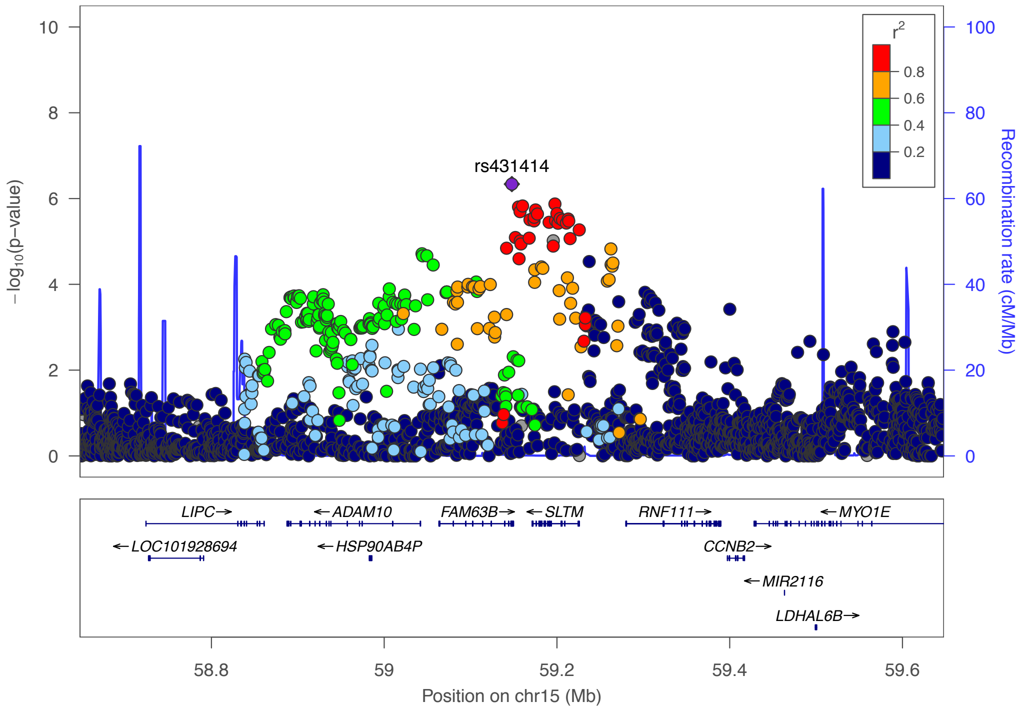
rs431414
−log10(p−value)
Recombination rate (cM/Mb)
r2
0.8
0.6
0.4
0.2
LIPC
LOC101928694
ADAM10
HSP90AB4P
FAM63B
SLTM
RNF111
CCNB2
MIR2116
LDHAL6B
MYO1E
58.8
59
59.2
59.4
59.6
Position on chr15 (Mb)

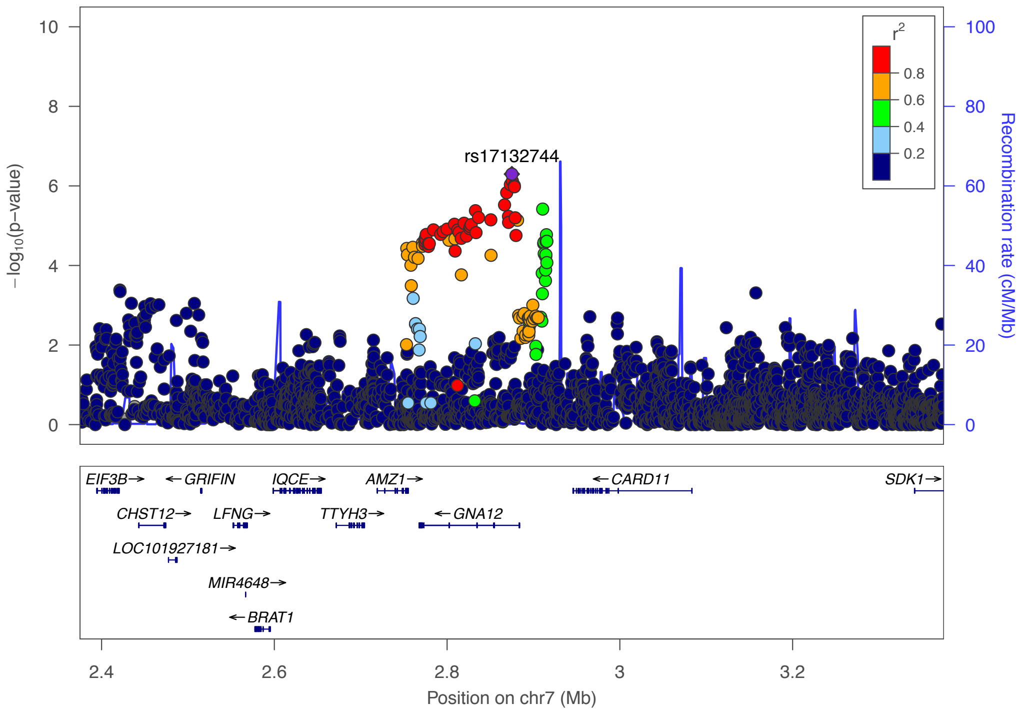
rs17132744
−log10(p−value)
Recombination rate (cM/Mb)
r2
0.8
0.6
0.4
0.2
EIF3B
GRIFIN
IQCE
AMZ1
CARD11
SDK1
CHST12
LFNG
TTYH3
GNA12
LOC101927181
MIR4648
BRAT1
2.4
2.6
2.8
3
3.2
Position on chr7 (Mb)

### k) Omnibus Test SCZ-BIP-MDD – European + East Asian ancestry

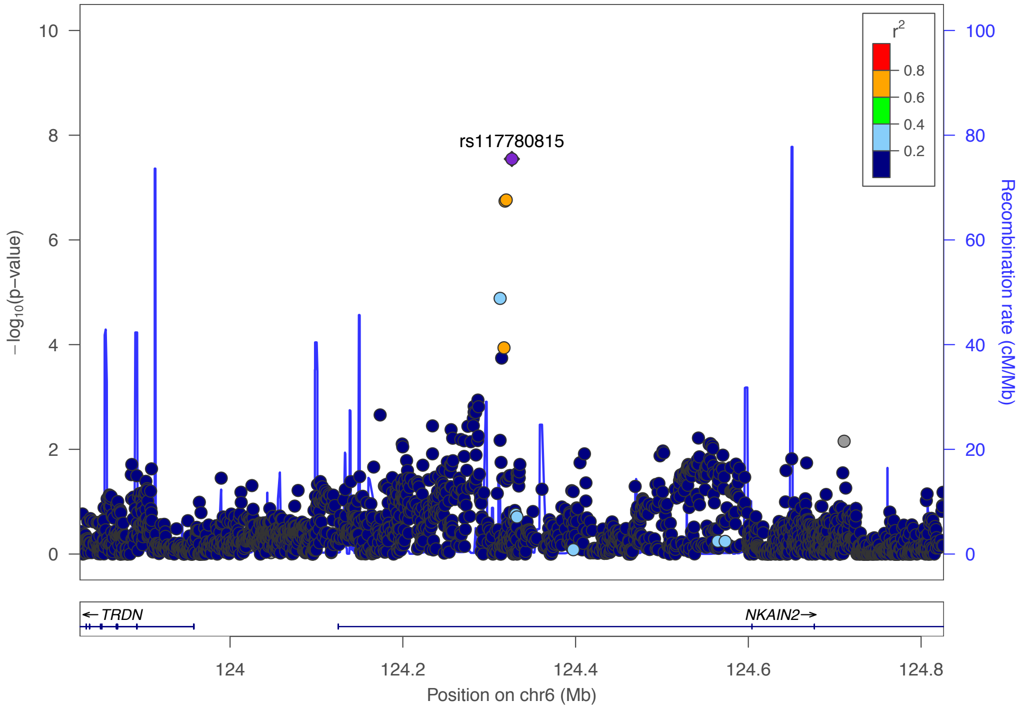


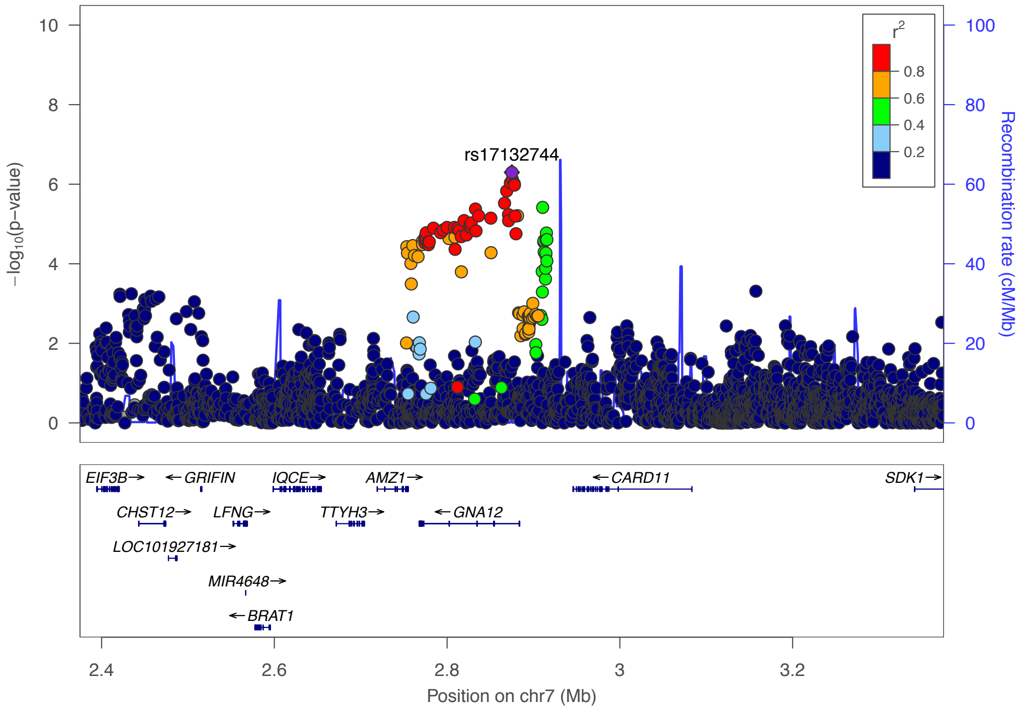

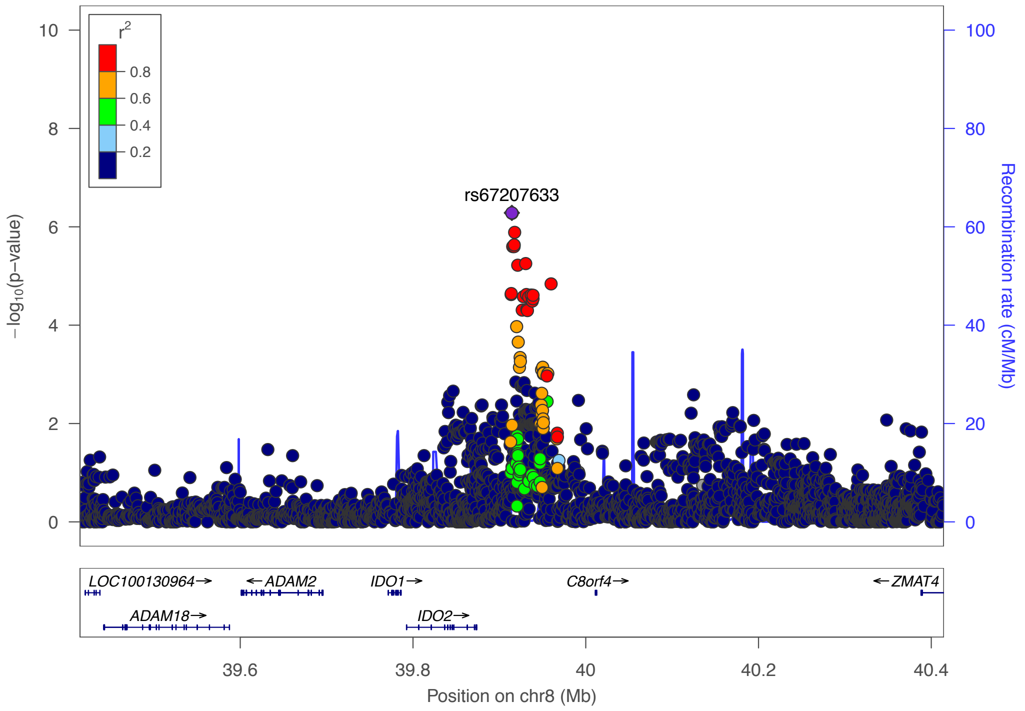


### l) Omnibus Test SCZ-BIP-rMDD – European ancestry

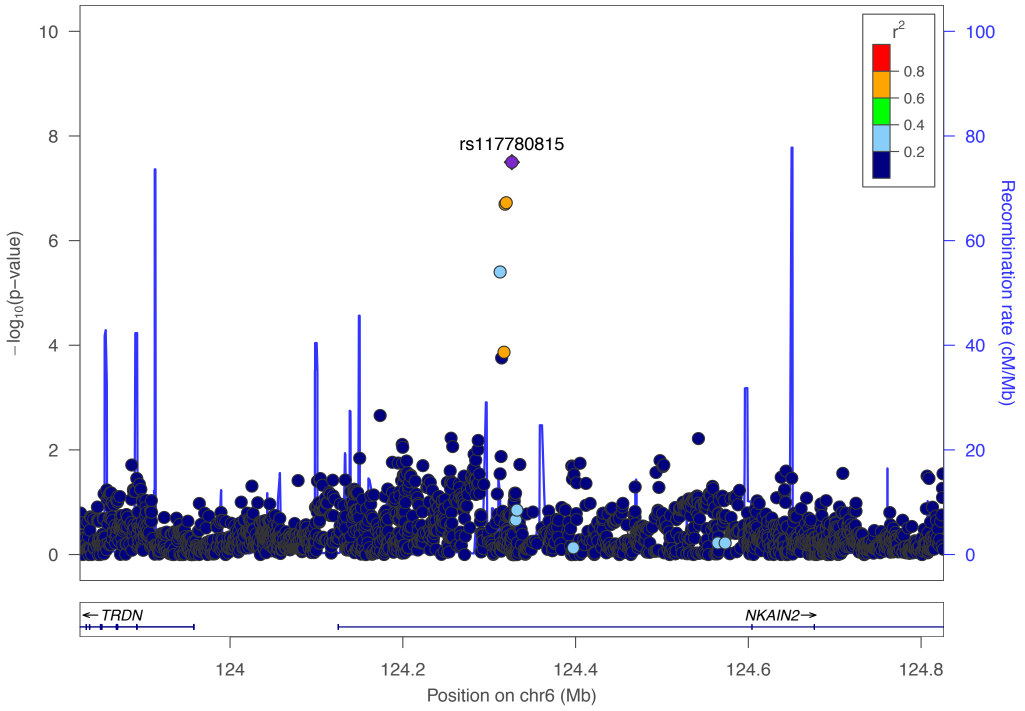

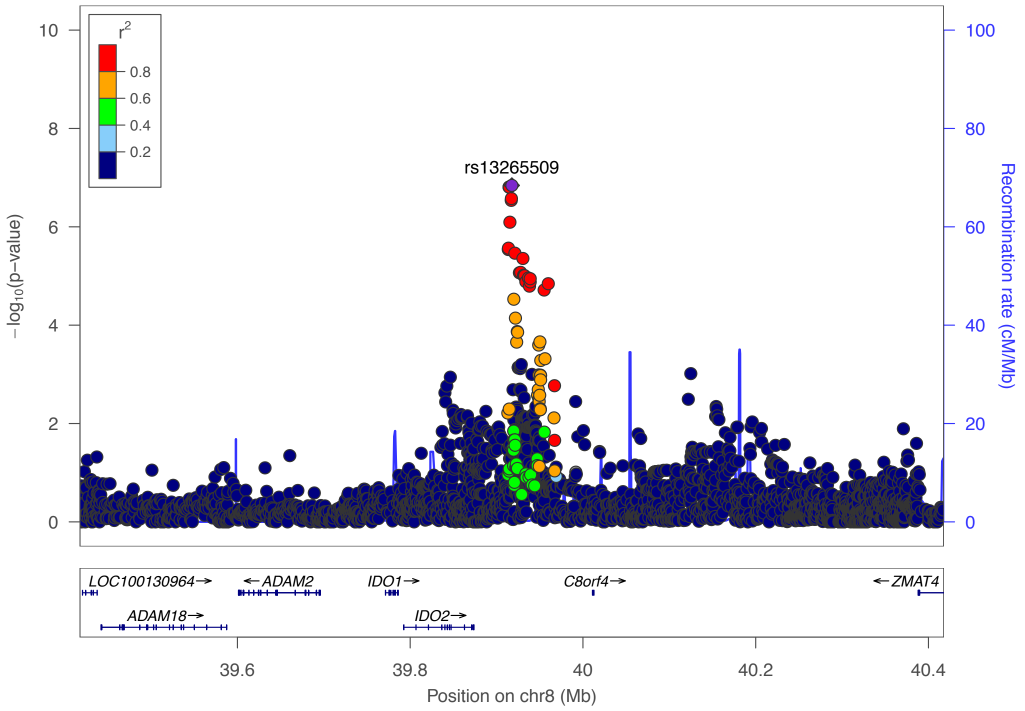
r2
0.8
0.6
0.4
0.2
rs13265509
−log10(p−value)
Recombination rate (cM/Mb)
LOC100130964→
←ADAM2
IDO1→
C8orf4→
←ZMAT4
ADAM18→
IDO2→
39.6
39.8
40
40.2
40.4
Position on chr8 (Mb)

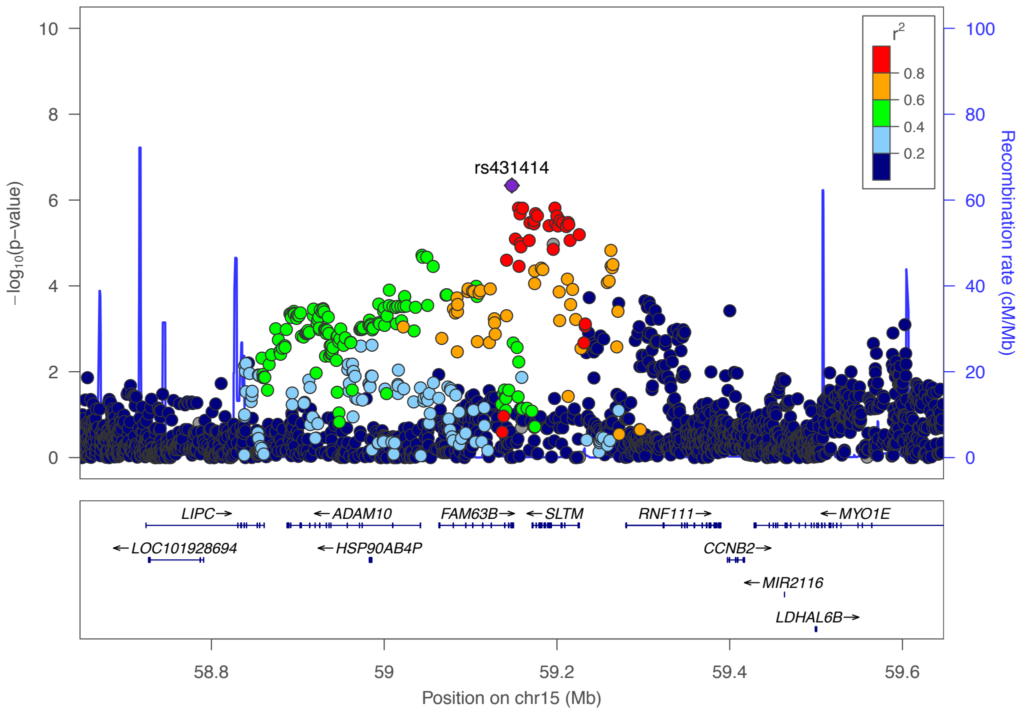
r2
0.8
0.6
0.4
0.2
rs431414
−log10(p−value)
Recombination rate (cM/Mb)
LIPC→
←ADAM10
FAM63B→
←SLTM
RNF111→
←MYO1E
←LOC101928694
←HSP90AB4P
CCNB2→
←MIR2116
LDHAL6B→
58.8
59
59.2
59.4
59.6
Position on chr15 (Mb)

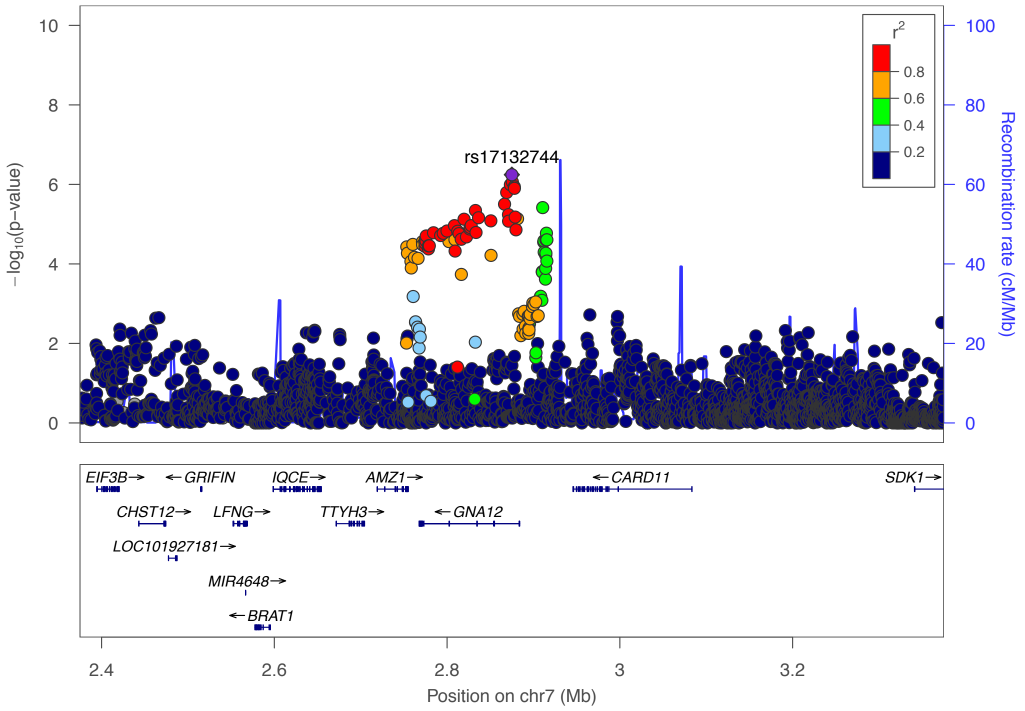


### m) Omnibus Test SCZ-BIP-rMDD – European + East Asian ancestry

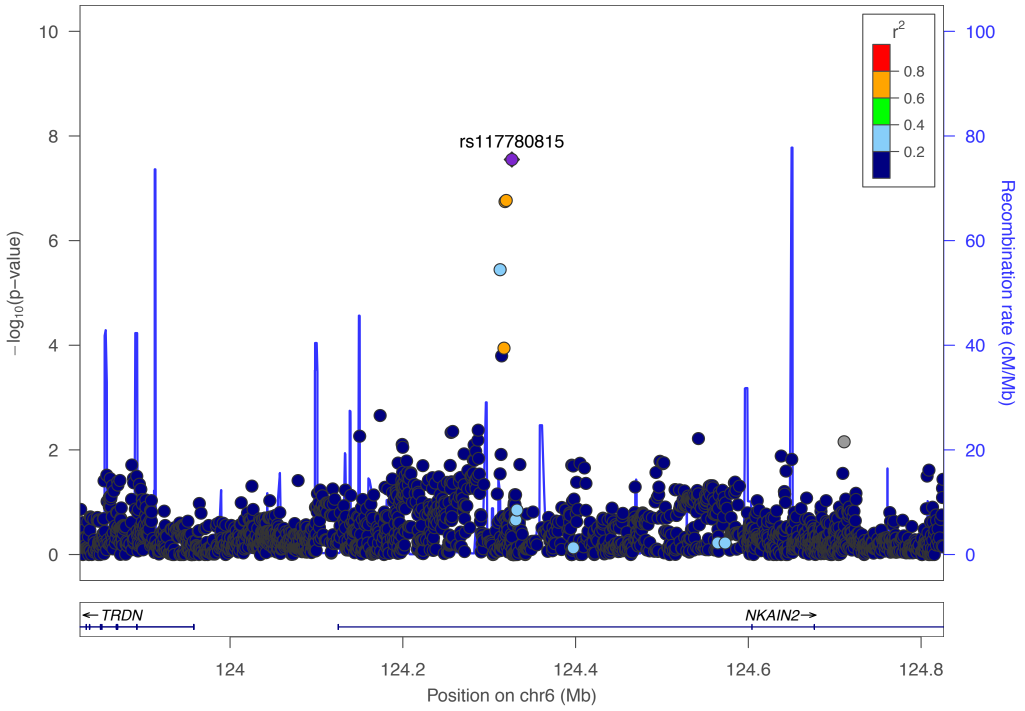

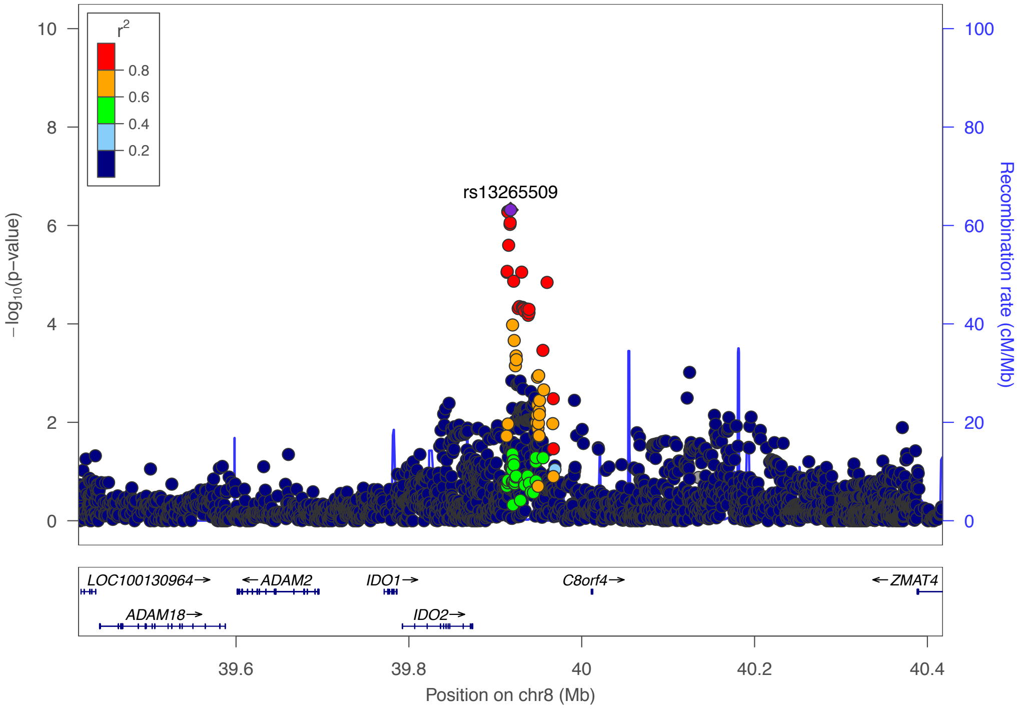
r2
0.8
0.6
0.4
0.2
rs13265509
−log10(p−value)
Recombination rate (cM/Mb)
10
8
6
4
2
0
100
80
60
40
20
0
LOC100130964→
←ADAM2
IDO1→
C8orf4→
←ZMAT4
ADAM18→
IDO2→
39.6
39.8
40
40.2
40.4
Position on chr8 (Mb)

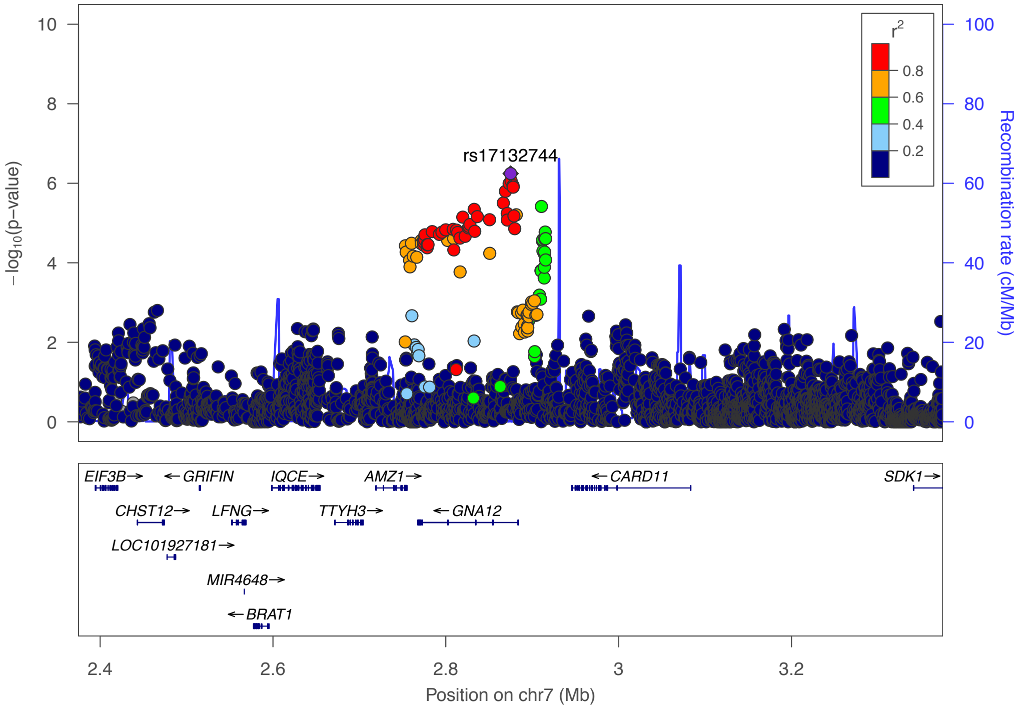
r2
0.8
0.6
0.4
0.2
rs17132744
−log10(p−value)
Recombination rate (cM/Mb)
10
8
6
4
2
0
100
80
60
40
20
0
EIF3B→
←GRIFIN
IQCE→
AMZ1→
←CARD11
SDK1→
CHST12→
LFNG→
TTYH3→
←GNA12
LOC101927181→
MIR4648→
←BRAT1
2.4
2.6
2.8
3
3.2
Position on chr7 (Mb)

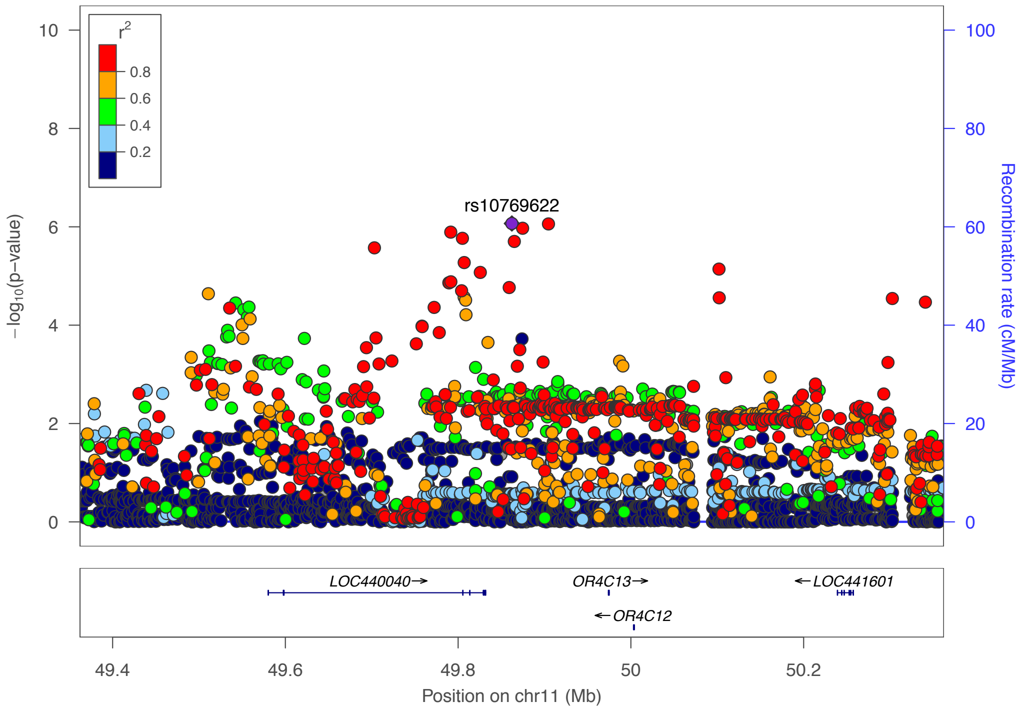
rs10769622
r2
0.8
0.6
0.4
0.2
10
8
6
4
2
0
−log10(p−value)
100
80
60
40
20
0
Recombination rate (cM/Mb)
LOC440040→
OR4C13→
←LOC441601
←OR4C12
49.4
49.6
49.8
50
50.2
Position on chr11 (Mb)

**Supplementary Figure 22. Forest plots for PGC**

Plots were generated using the rmeta package in R for loci (index SNPs) with GxS interaction $p < 1\times10^{-6}$.

Abbreviations: BIP = Bipolar Disorder; (r)MDD = (recurrent) Major Depressive Disorder; SCZ = Schizophrenia; HC F N = number of female healthy controls; HC M N = number of male healthy controls; PT F N = number of female patients; PT M N = number of male patients; Study = cohort abbreviation used by PGC; Meta = meta-analysis results

#### a) Schizophrenia – European ancestry only

rs13265509 (A/G)
Schizophrenia

| Study | HC F N | HC M N | PT F N | PT M N | p-value |
|---|---|---|---|---|---|
| aber | 241 | 436 | 178 | 526 | 9.56E-01 |
| ajsz | 412 | 1169 | 326 | 565 | 5.40E-01 |
| buls | 315 | 284 | 103 | 89 | 3.69E-02 |
| cati | 21 | 89 | 87 | 296 | 3.89E-01 |
| caws | 144 | 124 | 128 | 258 | 3.65E-01 |
| cims | 22 | 40 | 11 | 54 | 9.45E-01 |
| clm2 | 1960 | 1948 | 915 | 2420 | 1.47E-01 |
| clo3 | 876 | 1015 | 627 | 1426 | 7.58E-03 |
| cou3 | 339 | 329 | 203 | 318 | 8.03E-03 |
| denm | 189 | 258 | 192 | 259 | 5.69E-02 |
| dubl | 576 | 235 | 77 | 176 | 3.00E-02 |
| edin | 134 | 145 | 98 | 262 | 7.57E-01 |
| egcu | 819 | 295 | 168 | 62 | 6.31E-01 |
| ersw | 122 | 179 | 92 | 143 | 3.01E-01 |
| fii6 | 587 | 435 | 152 | 205 | 1.49E-01 |
| gras | 414 | 704 | 340 | 690 | 8.65E-01 |
| irwt | 438 | 544 | 427 | 856 | 7.10E-03 |
| lie2 | 141 | 125 | 37 | 95 | 8.18E-01 |
| mgs2 | 1265 | 1060 | 779 | 1768 | 1.63E-01 |
| msaf | 58 | 80 | 120 | 199 | 6.44E-01 |
| pewb | 902 | 861 | 141 | 392 | 7.29E-01 |
| pews | 95 | 138 | 62 | 87 | 5.20E-01 |
| s234 | 1073 | 1089 | 749 | 1129 | 8.72E-01 |
| swe1 | 102 | 100 | 93 | 115 | 8.32E-01 |
| swe5 | 1188 | 1299 | 689 | 1029 | 3.41E-01 |
| swe6 | 534 | 542 | 368 | 516 | 2.02E-01 |
| top8 | 201 | 201 | 160 | 212 | 9.67E-01 |
| umeb | 288 | 235 | 137 | 176 | 2.91E-01 |
| umes | 335 | 286 | 79 | 102 | 8.25E-01 |
| zhh1 | 95 | 90 | 63 | 126 | 6.62E-02 |
| **Meta** | **13864** | **14335** | **7590** | **14551** | **1.09E-07** |

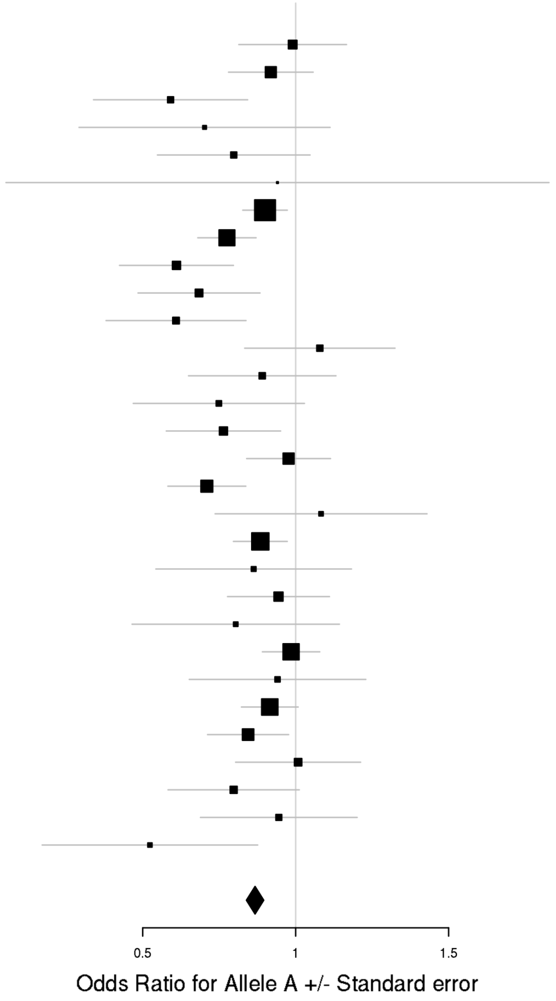

### b) Schizophrenia – European + East Asian ancestry

rs13265509 (A/G)
Schizophrenia

| Study | HC F N | HC M N | PT F N | PT M N | p-value |
|---|---|---|---|---|---|
| aber | 241 | 436 | 178 | 526 | 9.56E-01 |
| ajsz | 412 | 1169 | 326 | 565 | 5.40E-01 |
| buls | 315 | 284 | 103 | 89 | 3.69E-02 |
| cati | 21 | 89 | 87 | 296 | 3.89E-01 |
| caws | 144 | 124 | 128 | 258 | 3.65E-01 |
| cims | 22 | 40 | 11 | 54 | 9.45E-01 |
| clm2 | 1960 | 1948 | 915 | 2420 | 1.47E-01 |
| clo3 | 876 | 1015 | 627 | 1426 | 7.58E-03 |
| cou3 | 339 | 329 | 203 | 318 | 8.03E-03 |
| denm | 189 | 258 | 192 | 259 | 5.69E-02 |
| dubl | 576 | 235 | 77 | 176 | 3.00E-02 |
| edin | 134 | 145 | 98 | 262 | 7.57E-01 |
| egcu | 819 | 295 | 168 | 62 | 6.31E-01 |
| ersw | 122 | 179 | 92 | 143 | 3.01E-01 |
| fii6 | 587 | 435 | 152 | 205 | 1.49E-01 |
| gras | 414 | 704 | 340 | 690 | 8.65E-01 |
| irwt | 438 | 544 | 427 | 856 | 7.10E-03 |
| lie2 | 141 | 125 | 37 | 95 | 8.18E-01 |
| mgs2 | 1265 | 1060 | 779 | 1768 | 1.63E-01 |
| msaf | 58 | 80 | 120 | 199 | 6.44E-01 |
| pewb | 902 | 861 | 141 | 392 | 7.29E-01 |
| pews | 95 | 138 | 62 | 87 | 5.20E-01 |
| s234 | 1073 | 1089 | 749 | 1129 | 8.72E-01 |
| swe1 | 102 | 100 | 93 | 115 | 8.32E-01 |
| swe5 | 1188 | 1299 | 689 | 1029 | 3.41E-01 |
| swe6 | 534 | 542 | 368 | 516 | 2.02E-01 |
| top8 | 201 | 201 | 160 | 212 | 9.67E-01 |
| umeb | 288 | 235 | 137 | 176 | 2.91E-01 |
| umes | 335 | 286 | 79 | 102 | 8.25E-01 |
| zhh1 | 95 | 90 | 63 | 126 | 6.62E-02 |
| jpn1 | 215 | 212 | 237 | 252 | 3.70E-01 |
| **Meta** | **14079** | **14547** | **7827** | **14803** | **3.10E-07** |

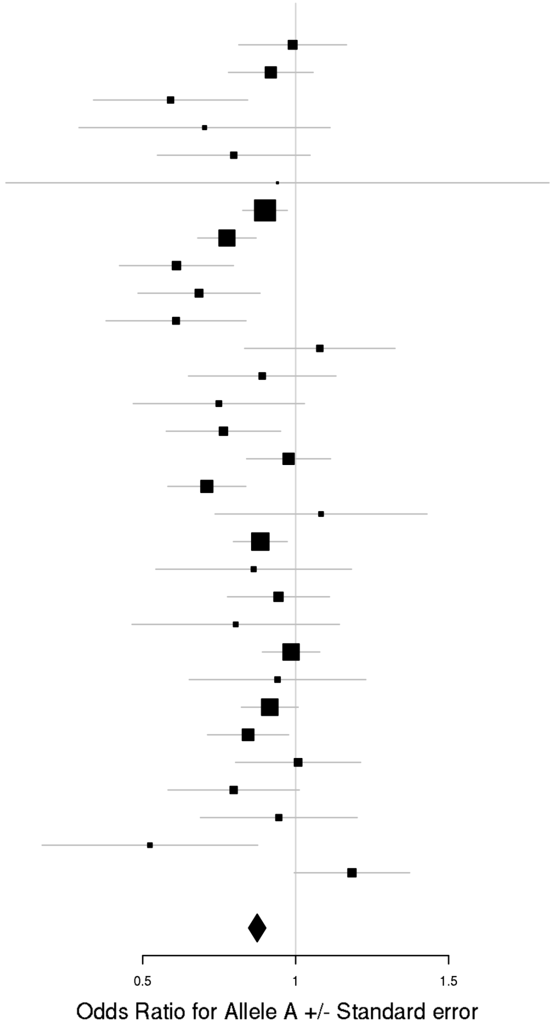

### c) Bipolar Disorder

rs12341335 (T/C)
Bipolar Disorder

| Study | HC F N | HC M N | PT F N | PT M N | p-value |
|---|---|---|---|---|---|
| bmau | 891 | 901 | 189 | 140 | 6.52E-02 |
| bmg2 | 211 | 232 | 117 | 64 | 8.20E-01 |
| bmg3 | 584 | 279 | 286 | 193 | 2.83E-01 |
| bonn | 593 | 607 | 343 | 321 | 1.04E-02 |
| gsk1 | 524 | 369 | 413 | 237 | 5.74E-01 |
| hal2 | 234 | 243 | 234 | 170 | 4.76E-01 |
| may1 | 377 | 377 | 560 | 374 | 3.92E-01 |
| mich | 153 | 135 | 9 | 3 | 9.11E-01 |
| rom3 | 107 | 86 | 134 | 96 | 2.23E-03 |
| swei | 1726 | 1846 | 786 | 470 | 9.50E-03 |
| top8 | 130 | 161 | 81 | 56 | 7.60E-02 |
| ucl2 | 244 | 444 | 397 | 324 | 1.50E-01 |
| ume4 | 291 | 257 | 350 | 211 | 4.39E-02 |
| usc2 | 680 | 472 | 633 | 661 | 8.94E-04 |
| **Meta** | **6745** | **6409** | **4532** | **3320** | **2.29E-07** |

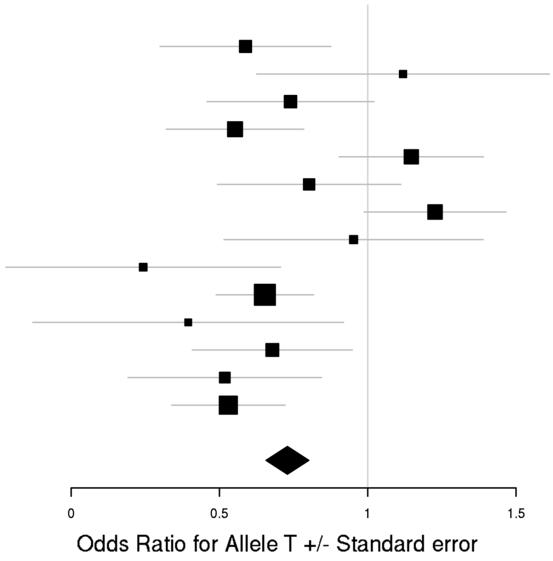

## rs17651437 (T/C)
### Bipolar Disorder

| Study | HC F N | HC M N | PT F N | PT M N | p-value |
|---|---|---|---|---|---|
| bmau | 889 | 898 | 185 | 140 | 1.48E-01 |
| bmg2 | 210 | 231 | 116 | 63 | 1.20E-01 |
| bmg3 | 587 | 281 | 282 | 194 | 2.78E-01 |
| bmpo | 207 | 471 | 249 | 159 | 5.41E-01 |
| bmsp | 143 | 143 | 137 | 111 | 4.24E-01 |
| bonn | 596 | 605 | 340 | 316 | 7.82E-01 |
| dub1 | 560 | 236 | 78 | 71 | 2.46E-02 |
| edi1 | 133 | 141 | 159 | 119 | 2.22E-01 |
| fat2 | 577 | 480 | 417 | 302 | 7.69E-01 |
| fran | 916 | 684 | 261 | 189 | 3.31E-01 |
| gain | 191 | 206 | 314 | 295 | 9.92E-02 |
| gsk1 | 527 | 369 | 412 | 239 | 7.32E-01 |
| hal2 | 233 | 242 | 231 | 170 | 7.61E-01 |
| icuk | 1661 | 1723 | 1710 | 757 | 5.31E-02 |
| may1 | 375 | 376 | 556 | 374 | 3.74E-01 |
| mich | 156 | 135 | 9 | 3 | 1.26E-01 |
| rom3 | 103 | 87 | 134 | 94 | 5.90E-01 |
| st2c | 551 | 632 | 361 | 272 | 9.17E-01 |
| stp1 | 378 | 406 | 502 | 417 | 1.56E-01 |
| swa2 | 1103 | 1122 | 516 | 361 | 1.53E-02 |
| swei | 1693 | 1820 | 779 | 460 | 1.58E-01 |
| top7 | 188 | 184 | 251 | 185 | 1.61E-01 |
| top8 | 130 | 159 | 78 | 55 | 5.23E-01 |
| ucl2 | 242 | 445 | 395 | 319 | 3.15E-01 |
| uclo | 283 | 211 | 242 | 172 | 3.51E-01 |
| wtcc | 499 | 456 | 977 | 576 | 5.24E-03 |
| **Meta** | **13131** | **12743** | **9691** | **6413** | **3.72E-07** |

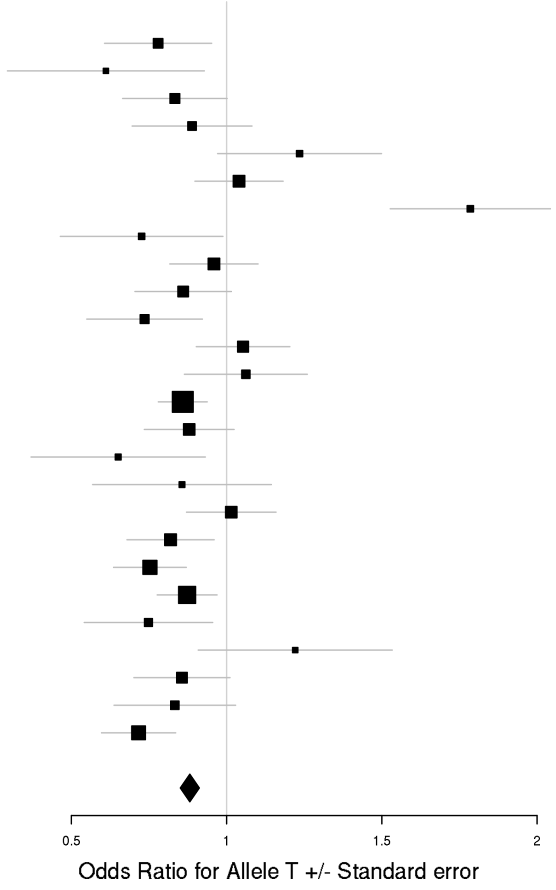

### d) Major Depressive Disorder

rs9428240 (T/C)
Major Depressive Disorder

| Study | HC F N | HC M N | PT F N | PT M N | p-value |
|---|---|---|---|---|---|
| boma | 530 | 536 | 377 | 210 | 4.96E-01 |
| col3 | 605 | 818 | 358 | 148 | 1.14E-02 |
| edi2 | 138 | 146 | 221 | 151 | 1.81E-01 |
| gens | 535 | 784 | 713 | 291 | 2.53E-01 |
| gep3 | 1346 | 1351 | 329 | 141 | 7.98E-01 |
| grdg | 292 | 176 | 365 | 106 | 6.33E-01 |
| grnd | 232 | 241 | 683 | 143 | 8.15E-01 |
| gsk2 | 577 | 276 | 563 | 284 | 1.93E-02 |
| i2b3 | 531 | 534 | 532 | 266 | 2.21E-01 |
| mmi2 | 277 | 227 | 314 | 267 | 6.01E-01 |
| mmo4 | 185 | 193 | 136 | 130 | 9.15E-01 |
| nes1 | 976 | 627 | 1021 | 471 | 1.06E-03 |
| qi6c | 364 | 223 | 346 | 151 | 8.67E-01 |
| rad3 | 820 | 557 | 1315 | 550 | 1.17E-03 |
| rage | 103 | 113 | 213 | 111 | 4.36E-01 |
| rai2 | 178 | 161 | 90 | 19 | 4.71E-01 |
| rau2 | 195 | 182 | 174 | 48 | 8.03E-01 |
| rde4 | 208 | 306 | 93 | 40 | 6.58E-01 |
| rot4 | 570 | 422 | 181 | 63 | 3.10E-01 |
| shp0 | 493 | 638 | 254 | 119 | 3.62E-03 |
| shpt | 223 | 276 | 119 | 46 | 9.23E-01 |
| stm2 | 422 | 503 | 551 | 368 | 1.06E-01 |
| twg2 | 1182 | 1417 | 773 | 321 | 2.33E-01 |
| **Meta** | **10982** | **10707** | **9721** | **4444** | **1.64E-07** |

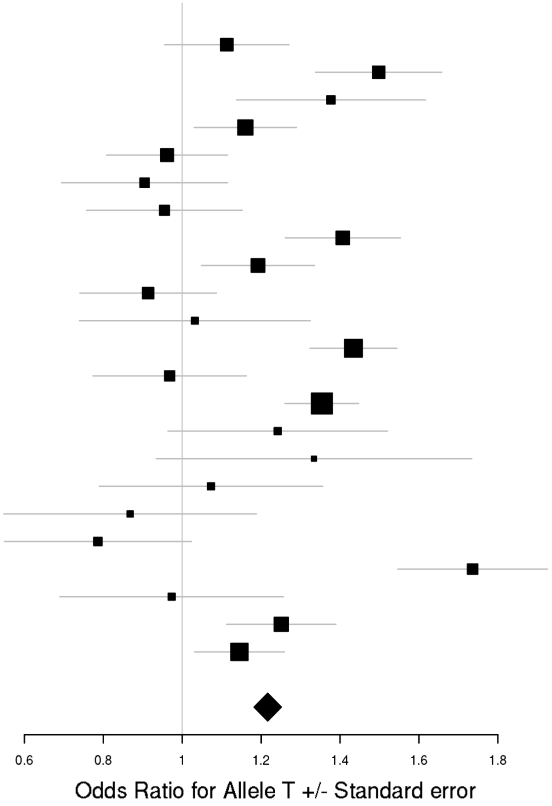

### e) Recurrent Major Depressive Disorder

chr1_118832069_D (D/I2)
Recurrent Major Depressive Disorder

| Study | HC F N | HC M N | PT F N | PT M N | p-value |
|---|---|---|---|---|---|
| boma | 530 | 536 | 297 | 149 | 4.47E-01 |
| col3 | 604 | 818 | 355 | 146 | 1.09E-02 |
| gens | 535 | 782 | 711 | 291 | 2.22E-01 |
| gep3 | 1345 | 1349 | 260 | 85 | 7.27E-01 |
| grnd | 231 | 238 | 673 | 140 | 6.86E-01 |
| gsk2 | 577 | 275 | 440 | 226 | 1.02E-02 |
| mmi2 | 276 | 227 | 229 | 174 | 8.85E-01 |
| mmo4 | 185 | 193 | 104 | 89 | 6.97E-01 |
| nes1 | 976 | 627 | 482 | 220 | 4.93E-04 |
| qi6c | 364 | 222 | 156 | 84 | 9.28E-01 |
| rad3 | 817 | 556 | 1016 | 419 | 1.19E-03 |
| rage | 103 | 113 | 182 | 75 | 3.21E-01 |
| rai2 | 178 | 161 | 89 | 19 | 5.07E-01 |
| rau2 | 195 | 182 | 175 | 47 | 8.59E-01 |
| rde4 | 208 | 306 | 41 | 18 | 1.34E-01 |
| rot4 | 570 | 422 | 58 | 14 | 8.87E-01 |
| shp0 | 493 | 638 | 128 | 52 | 5.52E-03 |
| **Meta** | **8187** | **7645** | **5396** | **2248** | **1.39E-07** |

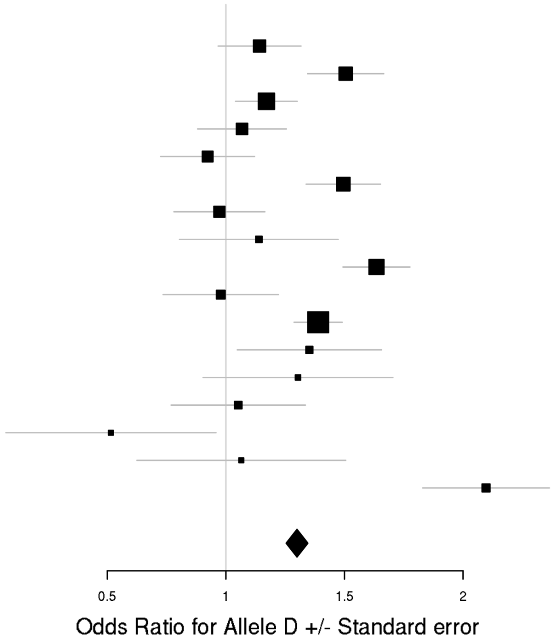

### f) Cross-Disorder SCZ-BIP-MDD – European ancestry only

rs7302529 (T/C)
Cross-Disorder SCZ-BIP-MDD

| Dx | Study | HC F N | HC M N | PT F N | PT M N | p-value |
|---|---|---|---|---|---|---|
| SCZ | aarh | 391 | 463 | 387 | 465 | 9.74E-03 |
| SCZ | ajsz | 414 | 1171 | 327 | 567 | 3.63E-01 |
| SCZ | asrb | 153 | 121 | 148 | 280 | 4.17E-02 |
| SCZ | boco | 160 | 160 | 761 | 952 | 3.97E-01 |
| SCZ | cims | 20 | 33 | 11 | 52 | 5.10E-01 |
| SCZ | clm2 | 671 | 657 | 908 | 2393 | 1.28E-01 |
| SCZ | clo3 | 684 | 818 | 627 | 1435 | 8.16E-02 |
| SCZ | denm | 188 | 259 | 193 | 266 | 2.61E-02 |
| SCZ | egcu | 828 | 299 | 169 | 62 | 1.20E-01 |
| SCZ | ersw | 120 | 175 | 92 | 141 | 9.94E-01 |
| SCZ | fi3m | 469 | 457 | 85 | 101 | 6.67E-01 |
| SCZ | fii6 | 585 | 427 | 152 | 207 | 1.85E-01 |
| SCZ | lacw | 23 | 21 | 51 | 104 | 9.77E-01 |
| SCZ | lie2 | 144 | 124 | 37 | 94 | 7.65E-01 |
| SCZ | lie5 | 204 | 182 | 126 | 363 | 3.83E-01 |
| SCZ | munc | 156 | 136 | 150 | 266 | 4.51E-01 |
| SCZ | swe5 | 585 | 654 | 690 | 1034 | 3.19E-01 |
| SCZ | swe6 | 259 | 283 | 360 | 511 | 9.70E-01 |
| SCZ | ucla | 278 | 282 | 170 | 522 | 9.44E-01 |
| SCZ | umeb | 183 | 146 | 109 | 154 | 2.63E-01 |
| SCZ | umes | 266 | 232 | 72 | 93 | 2.68E-01 |
| BIP | bmau | 895 | 895 | 184 | 140 | 8.66E-01 |
| BIP | bmg2 | 210 | 226 | 117 | 64 | 4.03E-01 |
| BIP | bmg3 | 158 | 54 | 288 | 197 | 7.26E-01 |
| BIP | bmpo | 213 | 476 | 250 | 160 | 9.67E-01 |
| BIP | bmsp | 144 | 147 | 138 | 114 | 1.62E-01 |
| BIP | bonn | 151 | 142 | 345 | 318 | 5.85E-01 |
| BIP | fran | 934 | 693 | 262 | 189 | 8.29E-01 |
| BIP | gsk1 | 408 | 299 | 415 | 239 | 9.25E-01 |
| BIP | icuk | 616 | 671 | 1691 | 741 | 8.58E-01 |
| BIP | may1 | 372 | 369 | 553 | 371 | 6.07E-02 |
| BIP | mich | 69 | 60 | 9 | 3 | 3.86E-01 |
| BIP | swei | 873 | 898 | 765 | 459 | 2.69E-03 |
| BIP | top8 | 126 | 152 | 80 | 55 | 8.16E-01 |
| BIP | ucl2 | 240 | 440 | 393 | 319 | 7.70E-01 |
| BIP | ume4 | 136 | 121 | 294 | 183 | 4.19E-01 |
| BIP | usc2 | 650 | 433 | 632 | 654 | 5.01E-02 |
| MDD | boma | 131 | 149 | 377 | 209 | 5.57E-02 |
| MDD | gep3 | 382 | 361 | 328 | 142 | 8.69E-01 |
| MDD | grdg | 290 | 176 | 365 | 106 | 1.11E-01 |
| MDD | grnd | 199 | 212 | 682 | 143 | 3.73E-01 |
| MDD | gsk2 | 134 | 65 | 561 | 285 | 7.44E-01 |
| MDD | mmi2 | 278 | 228 | 315 | 265 | 5.93E-02 |
| MDD | mmo4 | 181 | 189 | 135 | 129 | 7.39E-01 |
| MDD | rad3 | 694 | 485 | 1303 | 548 | 2.59E-01 |
| MDD | rage | 29 | 34 | 211 | 109 | 1.30E-01 |
| MDD | rai2 | 173 | 157 | 89 | 19 | 1.17E-02 |
| MDD | rau2 | 196 | 182 | 175 | 48 | 4.59E-01 |
| MDD | rot4 | 571 | 422 | 181 | 63 | 6.89E-01 |
| MDD | shpt | 224 | 276 | 119 | 46 | 9.51E-01 |
| MDD | twg2 | 1112 | 1336 | 763 | 317 | 1.71E-01 |
| | **Meta** | **17550** | **17448** | **17634** | **16697** | **1.60E-07** |

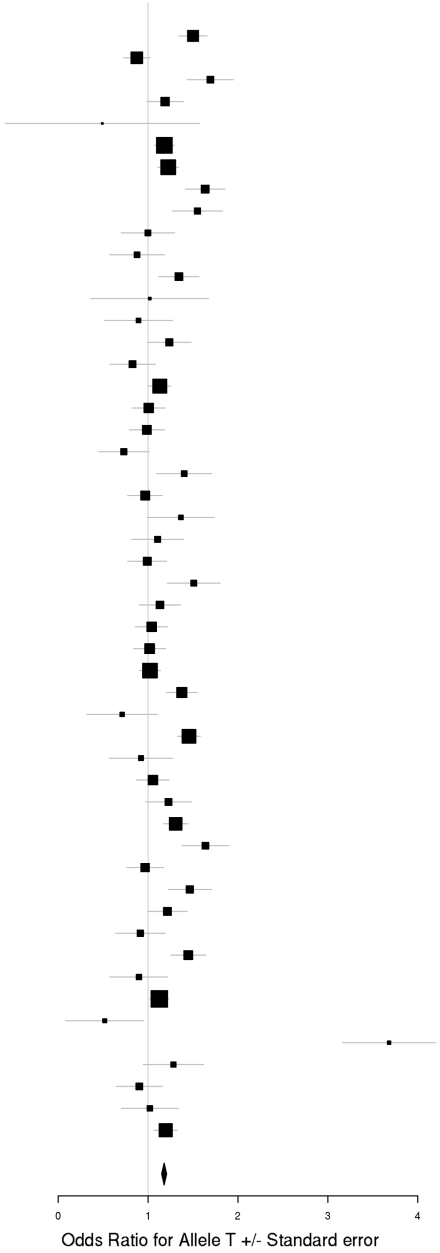

## rs73033497 (A/T)
### Cross-Disorder SCZ-BIP-MDD

| Dx | Study | HC F N | HC M N | PT F N | PT M N | p-value |
|---|---|---|---|---|---|---|
| SCZ | aarh | 383 | 455 | 380 | 454 | 3.47E-01 |
| SCZ | ajsz | 410 | 1155 | 320 | 561 | 6.30E-01 |
| SCZ | buls | 312 | 281 | 101 | 91 | 5.29E-01 |
| SCZ | cims | 20 | 33 | 11 | 52 | 1.69E-01 |
| SCZ | cou3 | 89 | 73 | 200 | 310 | 4.88E-01 |
| SCZ | dubl | 315 | 129 | 78 | 175 | 6.91E-02 |
| SCZ | edin | 37 | 37 | 98 | 262 | 4.76E-01 |
| SCZ | egcu | 815 | 294 | 165 | 61 | 3.88E-01 |
| SCZ | ersw | 120 | 178 | 91 | 140 | 6.08E-01 |
| SCZ | fi3m | 464 | 452 | 83 | 99 | 3.87E-01 |
| SCZ | irwt | 432 | 544 | 424 | 847 | 5.10E-02 |
| SCZ | s234 | 508 | 543 | 732 | 1109 | 1.46E-01 |
| SCZ | swe5 | 580 | 651 | 676 | 1015 | 5.72E-04 |
| SCZ | swe6 | 258 | 280 | 359 | 502 | 2.69E-01 |
| SCZ | top8 | 101 | 101 | 158 | 212 | 1.21E-01 |
| BIP | bmg3 | 154 | 53 | 282 | 194 | 4.10E-01 |
| BIP | dub1 | 280 | 112 | 76 | 68 | 3.99E-01 |
| BIP | gain | 84 | 64 | 311 | 288 | 1.09E-01 |
| BIP | may1 | 373 | 368 | 552 | 366 | 1.53E-01 |
| BIP | swei | 854 | 891 | 757 | 453 | 9.57E-01 |
| BIP | top8 | 127 | 150 | 79 | 55 | 5.81E-01 |
| BIP | ucl2 | 241 | 436 | 387 | 319 | 2.61E-01 |
| MDD | gens | 129 | 193 | 702 | 291 | 3.21E-01 |
| MDD | shp0 | 485 | 623 | 250 | 116 | 2.09E-01 |
| MDD | shpt | 223 | 273 | 118 | 46 | 4.03E-01 |
| MDD | twg2 | 1104 | 1321 | 756 | 309 | 3.76E-01 |
| | **Meta** | **8878** | **9690** | **8135** | **8395** | **8.82E-07** |

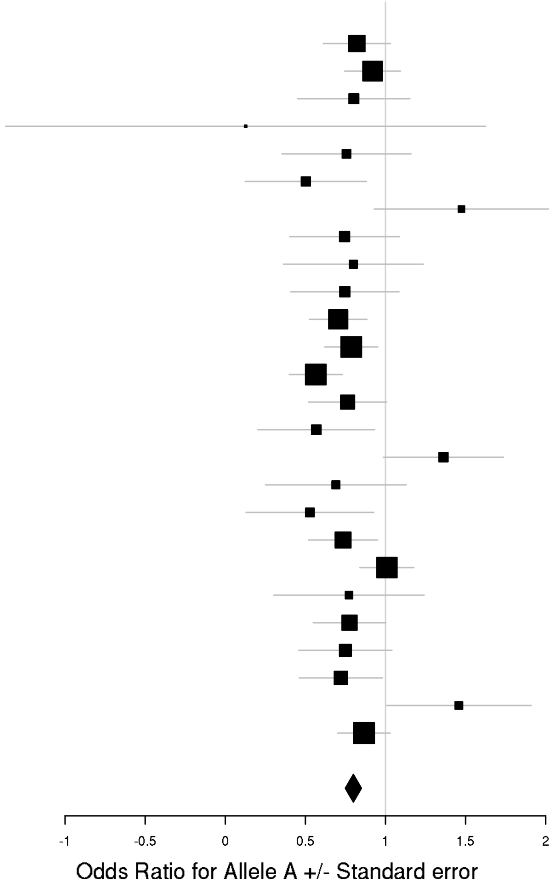

rs144142342 (C/G)
Cross-Disorder SCZ-BIP-MDD

| Dx | Study | HC F N | HC M N | PT F N | PT M N | p-value |
|---|---|---|---|---|---|---|
| SCZ | aarh | 390 | 462 | 386 | 464 | 7.57E-01 |
| SCZ | aber | 243 | 438 | 182 | 527 | 9.70E-01 |
| SCZ | ajsz | 414 | 1170 | 327 | 567 | 8.31E-02 |
| SCZ | asrb | 153 | 121 | 148 | 279 | 8.02E-01 |
| SCZ | boco | 160 | 160 | 759 | 952 | 1.77E-01 |
| SCZ | buls | 316 | 287 | 104 | 91 | 6.84E-01 |
| SCZ | cati | 7 | 44 | 90 | 304 | 3.37E-01 |
| SCZ | caws | 65 | 63 | 130 | 256 | 3.39E-01 |
| SCZ | cims | 20 | 33 | 11 | 52 | 5.84E-01 |
| SCZ | clm2 | 680 | 667 | 923 | 2448 | 2.44E-01 |
| SCZ | clo3 | 691 | 823 | 629 | 1440 | 7.96E-01 |
| SCZ | cou3 | 91 | 73 | 202 | 317 | 1.23E-01 |
| SCZ | denm | 188 | 259 | 193 | 265 | 5.14E-01 |
| SCZ | dubl | 319 | 129 | 79 | 176 | 3.18E-01 |
| SCZ | edin | 37 | 37 | 98 | 264 | 3.64E-01 |
| SCZ | egcu | 827 | 299 | 169 | 62 | 7.88E-01 |
| SCZ | ersw | 121 | 178 | 91 | 142 | 4.05E-01 |
| SCZ | fi3m | 468 | 457 | 85 | 100 | 7.07E-01 |
| SCZ | fii6 | 594 | 437 | 153 | 207 | 3.10E-01 |
| SCZ | gras | 413 | 710 | 342 | 694 | 3.46E-01 |
| SCZ | irwt | 435 | 546 | 425 | 857 | 8.76E-01 |
| SCZ | lacw | 23 | 21 | 51 | 104 | 8.17E-01 |
| SCZ | lie2 | 144 | 124 | 37 | 94 | 5.38E-01 |
| SCZ | lie5 | 204 | 181 | 126 | 362 | 2.84E-01 |
| SCZ | mgs2 | 525 | 379 | 784 | 1781 | 8.65E-02 |
| SCZ | msaf | 57 | 79 | 121 | 199 | 5.77E-01 |
| SCZ | munc | 156 | 136 | 150 | 264 | 6.57E-01 |
| SCZ | pewb | 880 | 854 | 140 | 394 | 7.95E-01 |
| SCZ | pews | 96 | 137 | 63 | 86 | 6.43E-01 |
| SCZ | port | 131 | 78 | 133 | 212 | 2.65E-01 |
| SCZ | s234 | 515 | 546 | 743 | 1118 | 2.94E-01 |
| SCZ | swe1 | 99 | 98 | 94 | 115 | 5.89E-01 |
| SCZ | swe5 | 583 | 653 | 690 | 1033 | 3.34E-02 |
| SCZ | swe6 | 259 | 283 | 360 | 511 | 9.74E-01 |
| SCZ | top8 | 101 | 100 | 159 | 212 | 7.24E-01 |
| SCZ | ucla | 278 | 282 | 170 | 522 | 3.93E-02 |
| SCZ | uclo | 143 | 95 | 144 | 358 | 9.14E-02 |
| SCZ | umeb | 185 | 149 | 110 | 156 | 3.08E-01 |
| SCZ | umes | 269 | 235 | 72 | 93 | 9.31E-01 |
| SCZ | zhh1 | 98 | 91 | 63 | 127 | 3.81E-01 |
| BIP | bmau | 894 | 894 | 184 | 140 | 5.59E-01 |
| BIP | bmg2 | 210 | 226 | 117 | 63 | 6.81E-01 |
| BIP | bmg3 | 158 | 54 | 288 | 197 | 5.99E-01 |
| BIP | bmpo | 213 | 476 | 250 | 159 | 3.65E-02 |
| BIP | bmsp | 143 | 147 | 138 | 114 | 6.67E-02 |
| BIP | bonn | 151 | 142 | 345 | 318 | 5.73E-01 |
| BIP | dub1 | 283 | 115 | 77 | 70 | 9.76E-01 |
| BIP | edi1 | 43 | 38 | 154 | 118 | 1.50E-01 |
| BIP | fat2 | 254 | 171 | 414 | 301 | 1.92E-01 |
| BIP | fran | 932 | 693 | 262 | 189 | 9.62E-01 |
| BIP | gain | 85 | 65 | 312 | 293 | 2.15E-01 |
| BIP | gsk1 | 407 | 299 | 415 | 239 | 3.28E-01 |
| BIP | hal2 | 65 | 73 | 234 | 171 | 4.01E-01 |
| BIP | icuk | 631 | 679 | 1725 | 757 | 4.56E-01 |
| BIP | may1 | 374 | 374 | 560 | 374 | 2.38E-01 |
| BIP | mich | 69 | 60 | 8 | 3 | 9.82E-01 |
| BIP | rom3 | 107 | 86 | 135 | 96 | 6.59E-01 |
| BIP | st2c | 399 | 497 | 359 | 269 | 3.20E-01 |
| BIP | stp1 | 286 | 312 | 502 | 415 | 4.55E-01 |
| BIP | swa2 | 578 | 569 | 498 | 348 | 7.63E-01 |
| BIP | swei | 873 | 896 | 764 | 457 | 5.81E-02 |
| BIP | top7 | 103 | 104 | 248 | 181 | 1.32E-01 |
| BIP | top8 | 128 | 155 | 81 | 56 | 5.36E-01 |
| BIP | ucl2 | 243 | 441 | 397 | 324 | 7.03E-01 |
| BIP | uclo | 138 | 110 | 239 | 170 | 5.29E-02 |
| BIP | ume4 | 139 | 121 | 299 | 184 | 5.50E-01 |
| BIP | usc2 | 659 | 438 | 635 | 659 | 7.11E-01 |
| BIP | wtcc | 204 | 185 | 970 | 573 | 1.57E-01 |
| MDD | boma | 131 | 149 | 377 | 208 | 2.94E-03 |
| MDD | cof3 | 61 | 65 | 74 | 46 | 6.66E-01 |
| MDD | col3 | 608 | 819 | 358 | 148 | 7.68E-01 |
| MDD | edi2 | 36 | 37 | 219 | 151 | 1.09E-01 |
| MDD | gens | 130 | 195 | 713 | 291 | 2.69E-01 |
| MDD | gep3 | 382 | 361 | 328 | 142 | 9.93E-01 |
| MDD | grdg | 290 | 176 | 364 | 106 | 2.72E-01 |
| MDD | grnd | 198 | 212 | 682 | 143 | 3.51E-01 |
| MDD | gsk2 | 134 | 65 | 561 | 285 | 6.59E-01 |
| MDD | i2b3 | 240 | 253 | 533 | 266 | 2.38E-01 |
| MDD | mmi2 | 276 | 228 | 314 | 265 | 1.30E-01 |
| MDD | mmo4 | 185 | 193 | 137 | 130 | 2.25E-01 |
| MDD | nes1 | 948 | 618 | 1018 | 470 | 5.72E-02 |
| MDD | qi3c | 368 | 208 | 498 | 361 | 4.51E-01 |
| MDD | qi6c | 363 | 220 | 347 | 152 | 7.17E-01 |
| MDD | qio2 | 301 | 223 | 402 | 160 | 5.23E-01 |
| MDD | rad3 | 694 | 485 | 1302 | 548 | 4.65E-01 |
| MDD | rage | 29 | 34 | 211 | 109 | 1.69E-01 |
| MDD | rai2 | 173 | 157 | 89 | 19 | 6.16E-01 |
| MDD | rau2 | 196 | 181 | 175 | 48 | 7.14E-01 |
| MDD | rde4 | 207 | 301 | 93 | 40 | 7.26E-01 |
| MDD | rot4 | 571 | 422 | 181 | 63 | 5.79E-01 |
| MDD | shp0 | 492 | 635 | 254 | 118 | 1.40E-01 |
| MDD | shpt | 224 | 275 | 118 | 46 | 3.06E-01 |
| MDD | stm2 | 279 | 356 | 551 | 367 | 1.34E-01 |
| MDD | twg2 | 1110 | 1336 | 763 | 316 | 5.76E-02 |
| | **Meta** | **28753** | **28538** | **30997** | **30441** | **9.06E-07** |

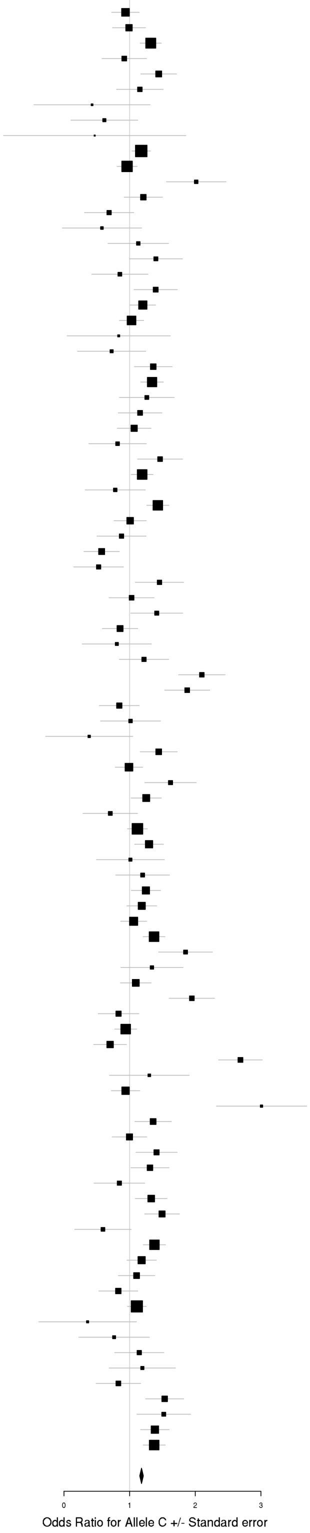

### g) Cross-Disorder SCZ-BIP-MDD – European + East Asian ancestry

rs73033497 (A/T)
Cross-Disorder SCZ-BIP-MDD

| Dx | Study | HC F N | HC M N | PT F N | PT M N | p-value |
|---|---|---|---|---|---|---|
| SCZ | aarh | 383 | 455 | 380 | 454 | 3.47E-01 |
| SCZ | ajsz | 410 | 1155 | 320 | 561 | 6.30E-01 |
| SCZ | buls | 312 | 281 | 101 | 91 | 5.29E-01 |
| SCZ | cims | 20 | 33 | 11 | 52 | 1.69E-01 |
| SCZ | cou3 | 89 | 73 | 200 | 310 | 4.88E-01 |
| SCZ | dubl | 315 | 129 | 78 | 175 | 6.91E-02 |
| SCZ | edin | 37 | 37 | 98 | 262 | 4.76E-01 |
| SCZ | egcu | 815 | 294 | 165 | 61 | 3.88E-01 |
| SCZ | ersw | 120 | 178 | 91 | 140 | 6.08E-01 |
| SCZ | fi3m | 464 | 452 | 83 | 99 | 3.87E-01 |
| SCZ | irwt | 432 | 544 | 424 | 847 | 5.10E-02 |
| SCZ | s234 | 508 | 543 | 732 | 1109 | 1.46E-01 |
| SCZ | swe5 | 580 | 651 | 676 | 1015 | 5.72E-04 |
| SCZ | swe6 | 258 | 280 | 359 | 502 | 2.69E-01 |
| SCZ | top8 | 101 | 101 | 158 | 212 | 1.21E-01 |
| BIP | bmg3 | 154 | 53 | 282 | 194 | 4.10E-01 |
| BIP | dub1 | 280 | 112 | 76 | 68 | 3.99E-01 |
| BIP | gain | 84 | 64 | 311 | 288 | 1.09E-01 |
| BIP | may1 | 373 | 368 | 552 | 366 | 1.53E-01 |
| BIP | swei | 854 | 891 | 757 | 453 | 9.57E-01 |
| BIP | top8 | 127 | 150 | 79 | 55 | 5.81E-01 |
| BIP | ucl2 | 241 | 436 | 387 | 319 | 2.61E-01 |
| MDD | gens | 129 | 193 | 702 | 291 | 3.21E-01 |
| MDD | shp0 | 485 | 623 | 250 | 116 | 2.09E-01 |
| MDD | shpt | 223 | 273 | 118 | 46 | 4.03E-01 |
| MDD | twg2 | 1104 | 1321 | 756 | 309 | 3.76E-01 |
| | **Meta** | **8878** | **9690** | **8135** | **8395** | **8.82E-07** |

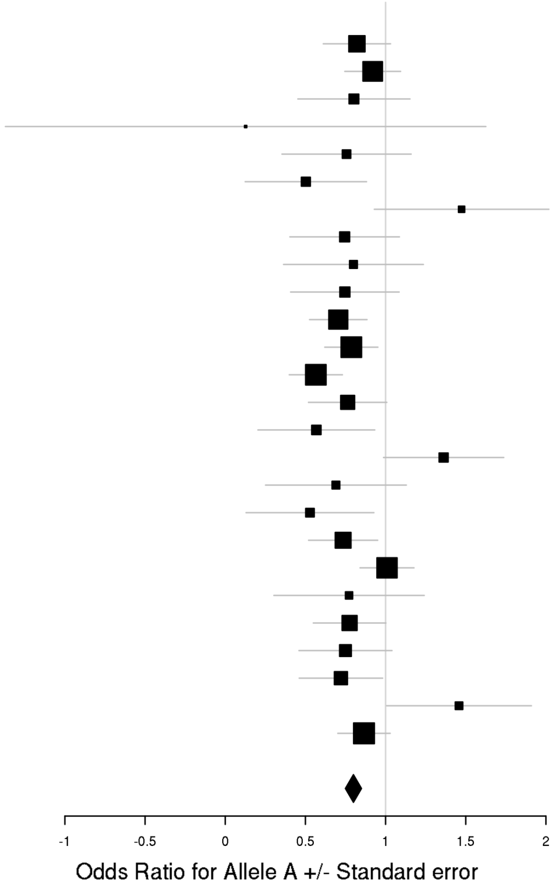

rs144142342 (C/G)
Cross-Disorder SCZ-BIP-MDD

| Dx | Study | HC F N | HC M N | PT F N | PT M N | p-value |
|---|---|---|---|---|---|---|
| SCZ | aarh | 390 | 462 | 386 | 464 | 7.57E-01 |
| SCZ | aber | 243 | 438 | 182 | 527 | 9.70E-01 |
| SCZ | ajsz | 414 | 1170 | 327 | 567 | 8.31E-02 |
| SCZ | asrb | 153 | 121 | 148 | 279 | 8.02E-01 |
| SCZ | boco | 160 | 160 | 759 | 952 | 1.77E-01 |
| SCZ | buls | 316 | 287 | 104 | 91 | 6.84E-01 |
| SCZ | cati | 7 | 44 | 90 | 304 | 3.37E-01 |
| SCZ | caws | 65 | 63 | 130 | 256 | 3.39E-01 |
| SCZ | cims | 20 | 33 | 11 | 52 | 5.84E-01 |
| SCZ | clm2 | 680 | 667 | 923 | 2448 | 2.44E-01 |
| SCZ | clo3 | 691 | 823 | 629 | 1440 | 7.96E-01 |
| SCZ | cou3 | 91 | 73 | 202 | 317 | 1.23E-01 |
| SCZ | denm | 188 | 259 | 193 | 265 | 5.14E-01 |
| SCZ | dubl | 319 | 129 | 79 | 176 | 3.18E-01 |
| SCZ | edin | 37 | 37 | 98 | 264 | 3.64E-01 |
| SCZ | egcu | 827 | 299 | 169 | 62 | 7.88E-01 |
| SCZ | ersw | 121 | 178 | 91 | 142 | 4.05E-01 |
| SCZ | fi3m | 468 | 457 | 85 | 100 | 7.07E-01 |
| SCZ | fii6 | 594 | 437 | 153 | 207 | 3.10E-01 |
| SCZ | gras | 413 | 710 | 342 | 694 | 3.46E-01 |
| SCZ | irwt | 435 | 546 | 425 | 857 | 8.76E-01 |
| SCZ | lacw | 23 | 21 | 51 | 104 | 8.17E-01 |
| SCZ | lie2 | 144 | 124 | 37 | 94 | 5.38E-01 |
| SCZ | lie5 | 204 | 181 | 126 | 362 | 2.84E-01 |
| SCZ | mgs2 | 525 | 379 | 784 | 1781 | 8.65E-02 |
| SCZ | msaf | 57 | 79 | 121 | 199 | 5.77E-01 |
| SCZ | munc | 156 | 136 | 150 | 264 | 6.57E-01 |
| SCZ | pewb | 880 | 854 | 140 | 394 | 7.95E-01 |
| SCZ | pews | 96 | 137 | 63 | 86 | 6.43E-01 |
| SCZ | port | 131 | 78 | 133 | 212 | 2.65E-01 |
| SCZ | s234 | 515 | 546 | 743 | 1118 | 2.94E-01 |
| SCZ | swe1 | 99 | 98 | 94 | 115 | 5.89E-01 |
| SCZ | swe5 | 583 | 653 | 690 | 1033 | 3.34E-02 |
| SCZ | swe6 | 259 | 283 | 360 | 511 | 9.74E-01 |
| SCZ | top8 | 101 | 100 | 159 | 212 | 7.24E-01 |
| SCZ | ucla | 278 | 282 | 170 | 522 | 3.93E-02 |
| SCZ | uclo | 143 | 95 | 144 | 358 | 9.14E-02 |
| SCZ | umeb | 185 | 149 | 110 | 156 | 3.08E-01 |
| SCZ | umes | 269 | 235 | 72 | 93 | 9.31E-01 |
| SCZ | zhh1 | 98 | 91 | 63 | 127 | 3.81E-01 |
| BIP | bmau | 894 | 894 | 184 | 140 | 5.59E-01 |
| BIP | bmg2 | 210 | 226 | 117 | 63 | 6.81E-01 |
| BIP | bmg3 | 158 | 54 | 288 | 197 | 5.99E-01 |
| BIP | bmpo | 213 | 476 | 250 | 159 | 3.65E-02 |
| BIP | bmsp | 143 | 147 | 138 | 114 | 6.67E-02 |
| BIP | bonn | 151 | 142 | 345 | 318 | 5.73E-01 |
| BIP | dub1 | 283 | 115 | 77 | 70 | 9.76E-01 |
| BIP | edi1 | 43 | 38 | 154 | 118 | 1.50E-01 |
| BIP | fat2 | 254 | 171 | 414 | 301 | 1.92E-01 |
| BIP | fran | 932 | 693 | 262 | 189 | 9.62E-01 |
| BIP | gain | 85 | 65 | 312 | 293 | 2.15E-01 |
| BIP | gsk1 | 407 | 299 | 415 | 239 | 3.28E-01 |
| BIP | hal2 | 65 | 73 | 234 | 171 | 4.01E-01 |
| BIP | icuk | 631 | 679 | 1725 | 757 | 4.56E-01 |
| BIP | may1 | 374 | 374 | 560 | 374 | 2.38E-01 |
| BIP | mich | 69 | 60 | 8 | 3 | 9.82E-01 |
| BIP | rom3 | 107 | 86 | 135 | 96 | 6.59E-01 |
| BIP | st2c | 399 | 497 | 359 | 269 | 3.20E-01 |
| BIP | stp1 | 286 | 312 | 502 | 415 | 4.55E-01 |
| BIP | swa2 | 578 | 569 | 498 | 348 | 7.63E-01 |
| BIP | swei | 873 | 896 | 764 | 457 | 5.81E-02 |
| BIP | top7 | 103 | 104 | 248 | 181 | 1.32E-01 |
| BIP | top8 | 128 | 155 | 81 | 56 | 5.36E-01 |
| BIP | ucl2 | 243 | 441 | 397 | 324 | 7.03E-01 |
| BIP | uclo | 138 | 110 | 239 | 170 | 5.29E-02 |
| BIP | ume4 | 139 | 121 | 299 | 184 | 5.50E-01 |
| BIP | usc2 | 659 | 438 | 635 | 659 | 7.11E-01 |
| BIP | wtcc | 204 | 185 | 970 | 573 | 1.57E-01 |
| MDD | boma | 131 | 149 | 377 | 208 | 2.94E-03 |
| MDD | cof3 | 61 | 65 | 74 | 46 | 6.66E-01 |
| MDD | col3 | 608 | 819 | 358 | 148 | 7.68E-01 |
| MDD | edi2 | 36 | 37 | 219 | 151 | 1.09E-01 |
| MDD | gens | 130 | 195 | 713 | 291 | 2.69E-01 |
| MDD | gep3 | 382 | 361 | 328 | 142 | 9.93E-01 |
| MDD | grdg | 290 | 176 | 364 | 106 | 2.72E-01 |
| MDD | grnd | 198 | 212 | 682 | 143 | 3.51E-01 |
| MDD | gsk2 | 134 | 65 | 561 | 285 | 6.59E-01 |
| MDD | i2b3 | 240 | 253 | 533 | 266 | 2.38E-01 |
| MDD | mmi2 | 276 | 228 | 314 | 265 | 1.30E-01 |
| MDD | mmo4 | 185 | 193 | 137 | 130 | 2.25E-01 |
| MDD | nes1 | 948 | 618 | 1018 | 470 | 5.72E-02 |
| MDD | qi3c | 368 | 208 | 498 | 361 | 4.51E-01 |
| MDD | qi6c | 363 | 220 | 347 | 152 | 7.17E-01 |
| MDD | qio2 | 301 | 223 | 402 | 160 | 5.23E-01 |
| MDD | rad3 | 694 | 485 | 1302 | 548 | 4.65E-01 |
| MDD | rage | 29 | 34 | 211 | 109 | 1.69E-01 |
| MDD | rai2 | 173 | 157 | 89 | 19 | 6.16E-01 |
| MDD | rau2 | 196 | 181 | 175 | 48 | 7.14E-01 |
| MDD | rde4 | 207 | 301 | 93 | 40 | 7.26E-01 |
| MDD | rot4 | 571 | 422 | 181 | 63 | 5.79E-01 |
| MDD | shp0 | 492 | 635 | 254 | 118 | 1.40E-01 |
| MDD | shpt | 224 | 275 | 118 | 46 | 3.06E-01 |
| MDD | stm2 | 279 | 356 | 551 | 367 | 1.34E-01 |
| MDD | twg2 | 1110 | 1336 | 763 | 316 | 5.76E-02 |
| | **Meta** | **28753** | **28538** | **30997** | **30441** | **9.06E-07** |

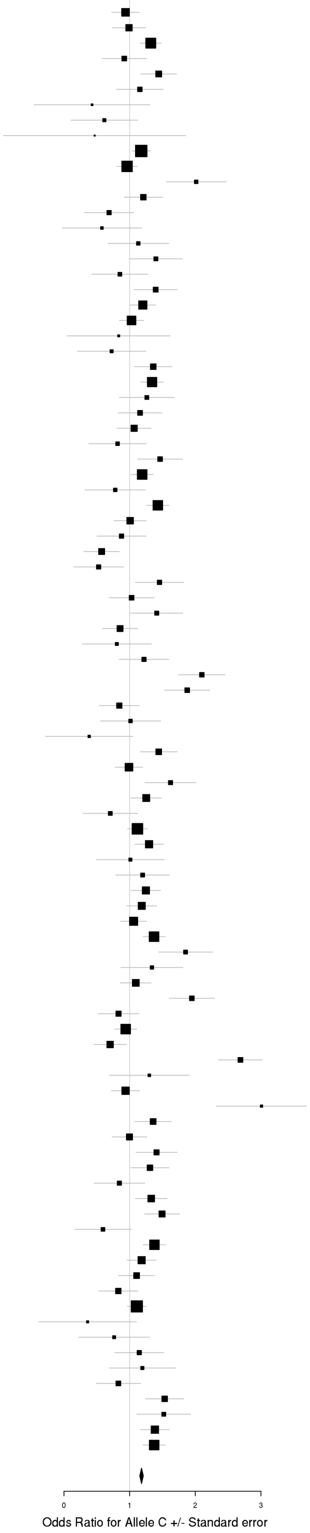

## rs7302529 (T/C)
### Cross-Disorder SCZ-BIP-MDD

| Dx | Study | HC F N | HC M N | PT F N | PT M N | p-value |
|---|---|---|---|---|---|---|
| SCZ | aarh | 391 | 463 | 387 | 465 | 9.74E-03 |
| SCZ | ajsz | 414 | 1171 | 327 | 567 | 3.63E-01 |
| SCZ | asrb | 153 | 121 | 148 | 280 | 4.17E-02 |
| SCZ | boco | 160 | 160 | 761 | 952 | 3.97E-01 |
| SCZ | cims | 20 | 33 | 11 | 52 | 5.10E-01 |
| SCZ | clm2 | 671 | 657 | 908 | 2393 | 1.28E-01 |
| SCZ | clo3 | 684 | 818 | 627 | 1435 | 8.16E-02 |
| SCZ | denm | 188 | 259 | 193 | 266 | 2.61E-02 |
| SCZ | egcu | 828 | 299 | 169 | 62 | 1.20E-01 |
| SCZ | ersw | 120 | 175 | 92 | 141 | 9.94E-01 |
| SCZ | fi3m | 469 | 457 | 85 | 101 | 6.67E-01 |
| SCZ | fii6 | 585 | 427 | 152 | 207 | 1.85E-01 |
| SCZ | lacw | 23 | 21 | 51 | 104 | 9.77E-01 |
| SCZ | lie2 | 144 | 124 | 37 | 94 | 7.65E-01 |
| SCZ | lie5 | 204 | 182 | 126 | 363 | 3.83E-01 |
| SCZ | munc | 156 | 136 | 150 | 266 | 4.51E-01 |
| SCZ | swe5 | 585 | 654 | 690 | 1034 | 3.19E-01 |
| SCZ | swe6 | 259 | 283 | 360 | 511 | 9.70E-01 |
| SCZ | ucla | 278 | 282 | 170 | 522 | 9.44E-01 |
| SCZ | umeb | 183 | 146 | 109 | 154 | 2.63E-01 |
| SCZ | umes | 266 | 232 | 72 | 93 | 2.68E-01 |
| SCZ | hok2 | 1344 | 668 | 154 | 322 | 2.33E-01 |
| SCZ | tcr1 | 336 | 570 | 289 | 563 | 6.13E-01 |
| BIP | bmau | 895 | 895 | 184 | 140 | 8.66E-01 |
| BIP | bmg2 | 210 | 226 | 117 | 64 | 4.03E-01 |
| BIP | bmg3 | 158 | 54 | 288 | 197 | 7.26E-01 |
| BIP | bmpo | 213 | 476 | 250 | 160 | 9.67E-01 |
| BIP | bmsp | 144 | 147 | 138 | 114 | 1.62E-01 |
| BIP | bonn | 151 | 142 | 345 | 318 | 5.85E-01 |
| BIP | fran | 934 | 693 | 262 | 189 | 8.29E-01 |
| BIP | gsk1 | 408 | 299 | 415 | 239 | 9.25E-01 |
| BIP | icuk | 616 | 671 | 1691 | 741 | 8.58E-01 |
| BIP | may1 | 372 | 369 | 553 | 371 | 6.07E-02 |
| BIP | mich | 69 | 60 | 9 | 3 | 3.86E-01 |
| BIP | swei | 873 | 898 | 765 | 459 | 2.69E-03 |
| BIP | top8 | 126 | 152 | 80 | 55 | 8.16E-01 |
| BIP | ucl2 | 240 | 440 | 393 | 319 | 7.70E-01 |
| BIP | ume4 | 136 | 121 | 294 | 183 | 4.19E-01 |
| BIP | usc2 | 650 | 433 | 632 | 654 | 5.01E-02 |
| MDD | boma | 131 | 149 | 377 | 209 | 5.57E-02 |
| MDD | gep3 | 382 | 361 | 328 | 142 | 8.69E-01 |
| MDD | grdg | 290 | 176 | 365 | 106 | 1.11E-01 |
| MDD | grnd | 199 | 212 | 682 | 143 | 3.73E-01 |
| MDD | gsk2 | 134 | 65 | 561 | 285 | 7.44E-01 |
| MDD | mmi2 | 278 | 228 | 315 | 265 | 5.93E-02 |
| MDD | mmo4 | 181 | 189 | 135 | 129 | 7.39E-01 |
| MDD | rad3 | 694 | 485 | 1303 | 548 | 2.59E-01 |
| MDD | rage | 29 | 34 | 211 | 109 | 1.30E-01 |
| MDD | rai2 | 173 | 157 | 89 | 19 | 1.17E-02 |
| MDD | rau2 | 196 | 182 | 175 | 48 | 4.59E-01 |
| MDD | rot4 | 571 | 422 | 181 | 63 | 6.89E-01 |
| MDD | shpt | 224 | 276 | 119 | 46 | 9.51E-01 |
| MDD | twg2 | 1112 | 1336 | 763 | 317 | 1.71E-01 |
| | **Meta** | **19230** | **18686** | **18077** | **17582** | **9.37E-07** |

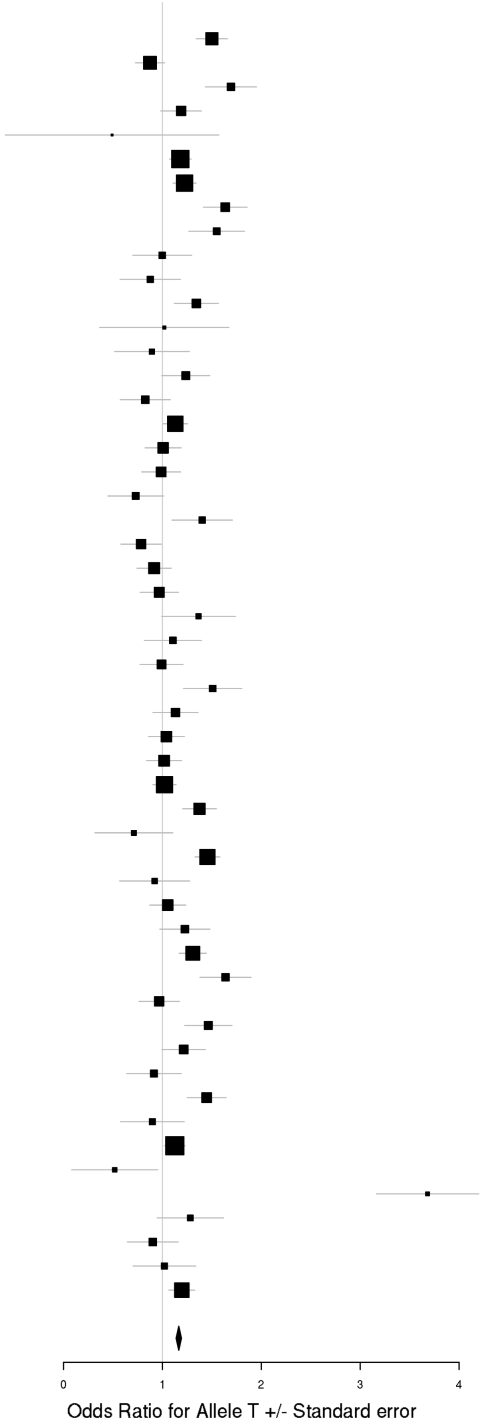

### h) Cross-Disorder SCZ-BIP-rMDD – European ancestry only

rs73033497 (A/T)
Cross-Disorder SCZ-BIP-RMDD

| Dx | Study | HC F N | HC M N | PT F N | PT M N | p-value |
|---|---|---|---|---|---|---|
| SCZ | aarh | 383 | 455 | 380 | 454 | 3.47E-01 |
| SCZ | ajsz | 410 | 1155 | 320 | 561 | 6.30E-01 |
| SCZ | buls | 312 | 281 | 101 | 91 | 5.29E-01 |
| SCZ | cims | 20 | 33 | 11 | 52 | 1.69E-01 |
| SCZ | cou3 | 89 | 73 | 200 | 310 | 4.88E-01 |
| SCZ | dubl | 315 | 129 | 78 | 175 | 6.91E-02 |
| SCZ | edin | 37 | 37 | 98 | 262 | 4.76E-01 |
| SCZ | egcu | 815 | 294 | 165 | 61 | 3.88E-01 |
| SCZ | ersw | 120 | 178 | 91 | 140 | 6.08E-01 |
| SCZ | fi3m | 464 | 452 | 83 | 99 | 3.87E-01 |
| SCZ | irwt | 432 | 544 | 424 | 847 | 5.10E-02 |
| SCZ | s234 | 508 | 543 | 732 | 1109 | 1.46E-01 |
| SCZ | swe5 | 580 | 651 | 676 | 1015 | 5.72E-04 |
| SCZ | swe6 | 258 | 280 | 359 | 502 | 2.69E-01 |
| SCZ | top8 | 101 | 101 | 158 | 212 | 1.21E-01 |
| BIP | bmg3 | 154 | 53 | 282 | 194 | 4.10E-01 |
| BIP | dub1 | 280 | 112 | 76 | 68 | 3.99E-01 |
| BIP | gain | 84 | 64 | 311 | 288 | 1.09E-01 |
| BIP | may1 | 373 | 368 | 552 | 366 | 1.53E-01 |
| BIP | swei | 854 | 891 | 757 | 453 | 9.57E-01 |
| BIP | top8 | 127 | 150 | 79 | 55 | 5.81E-01 |
| BIP | ucl2 | 241 | 436 | 387 | 319 | 2.61E-01 |
| rMDD | gens | 129 | 193 | 702 | 291 | 3.21E-01 |
| rMDD | shp0 | 485 | 623 | 125 | 51 | 2.25E-01 |
| | **Meta** | **7551** | **8096** | **7136** | **7975** | **6.22E-07** |

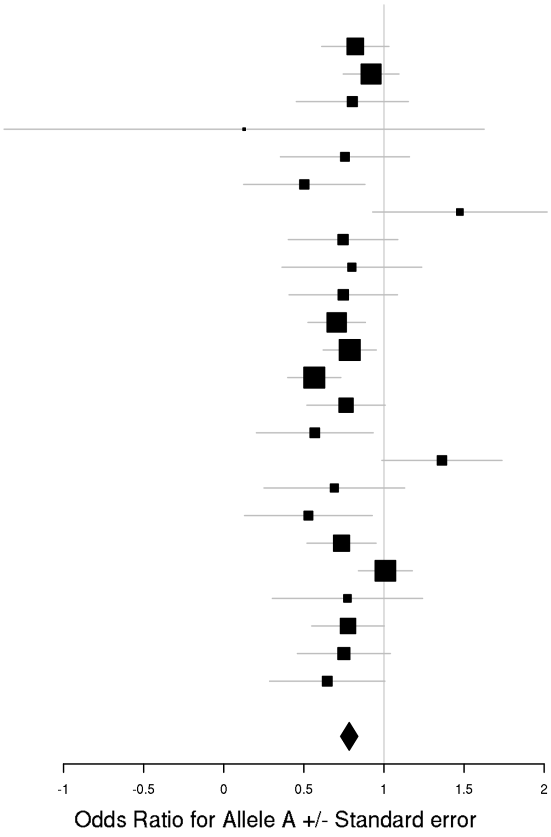

## rs7302529 (T/C)
### Cross-Disorder SCZ-BIP-RMDD

| Dx | Study | HC F N | HC M N | PT F N | PT M N | p-value |
|---|---|---|---|---|---|---|
| SCZ | aarh | 391 | 463 | 387 | 465 | 9.74E-03 |
| SCZ | ajsz | 414 | 1171 | 327 | 567 | 3.63E-01 |
| SCZ | asrb | 153 | 121 | 148 | 280 | 4.17E-02 |
| SCZ | boco | 160 | 160 | 761 | 952 | 3.97E-01 |
| SCZ | cims | 20 | 33 | 11 | 52 | 5.10E-01 |
| SCZ | clm2 | 671 | 657 | 908 | 2393 | 1.28E-01 |
| SCZ | clo3 | 684 | 818 | 627 | 1435 | 8.16E-02 |
| SCZ | denm | 188 | 259 | 193 | 266 | 2.61E-02 |
| SCZ | egcu | 828 | 299 | 169 | 62 | 1.20E-01 |
| SCZ | ersw | 120 | 175 | 92 | 141 | 9.94E-01 |
| SCZ | fi3m | 469 | 457 | 85 | 101 | 6.67E-01 |
| SCZ | fii6 | 585 | 427 | 152 | 207 | 1.85E-01 |
| SCZ | lacw | 23 | 21 | 51 | 104 | 9.77E-01 |
| SCZ | lie2 | 144 | 124 | 37 | 94 | 7.65E-01 |
| SCZ | lie5 | 204 | 182 | 126 | 363 | 3.83E-01 |
| SCZ | munc | 156 | 136 | 150 | 266 | 4.51E-01 |
| SCZ | swe5 | 585 | 654 | 690 | 1034 | 3.19E-01 |
| SCZ | swe6 | 259 | 283 | 360 | 511 | 9.70E-01 |
| SCZ | ucla | 278 | 282 | 170 | 522 | 9.44E-01 |
| SCZ | umeb | 183 | 146 | 109 | 154 | 2.63E-01 |
| SCZ | umes | 266 | 232 | 72 | 93 | 2.68E-01 |
| BIP | bmau | 895 | 895 | 184 | 140 | 8.66E-01 |
| BIP | bmg2 | 210 | 226 | 117 | 64 | 4.03E-01 |
| BIP | bmg3 | 158 | 54 | 288 | 197 | 7.26E-01 |
| BIP | bmpo | 213 | 476 | 250 | 160 | 9.67E-01 |
| BIP | bmsp | 144 | 147 | 138 | 114 | 1.62E-01 |
| BIP | bonn | 151 | 142 | 345 | 318 | 5.85E-01 |
| BIP | fran | 934 | 693 | 262 | 189 | 8.29E-01 |
| BIP | gsk1 | 408 | 299 | 415 | 239 | 9.25E-01 |
| BIP | icuk | 616 | 671 | 1691 | 741 | 8.58E-01 |
| BIP | may1 | 372 | 369 | 553 | 371 | 6.07E-02 |
| BIP | mich | 69 | 60 | 9 | 3 | 3.86E-01 |
| BIP | swei | 873 | 898 | 765 | 459 | 2.69E-03 |
| BIP | top8 | 126 | 152 | 80 | 55 | 8.16E-01 |
| BIP | ucl2 | 240 | 440 | 393 | 319 | 7.70E-01 |
| BIP | ume4 | 136 | 121 | 294 | 183 | 4.19E-01 |
| BIP | usc2 | 650 | 433 | 632 | 654 | 5.01E-02 |
| rMDD | boma | 131 | 149 | 299 | 148 | 1.20E-01 |
| rMDD | gep3 | 382 | 361 | 260 | 85 | 8.82E-01 |
| rMDD | grnd | 199 | 212 | 677 | 140 | 3.29E-01 |
| rMDD | gsk2 | 134 | 65 | 438 | 227 | 7.43E-01 |
| rMDD | mmi2 | 278 | 228 | 230 | 173 | 2.70E-02 |
| rMDD | mmo4 | 181 | 189 | 104 | 89 | 4.70E-01 |
| rMDD | rad3 | 694 | 485 | 1005 | 417 | 2.52E-01 |
| rMDD | rage | 29 | 34 | 180 | 73 | 2.86E-01 |
| rMDD | rai2 | 173 | 157 | 88 | 19 | 1.05E-02 |
| rMDD | rau2 | 196 | 182 | 175 | 47 | 5.17E-01 |
| rMDD | rot4 | 571 | 422 | 57 | 14 | 2.25E-01 |
| | **Meta** | **15924** | **15660** | **15543** | **15700** | **7.43E-07** |

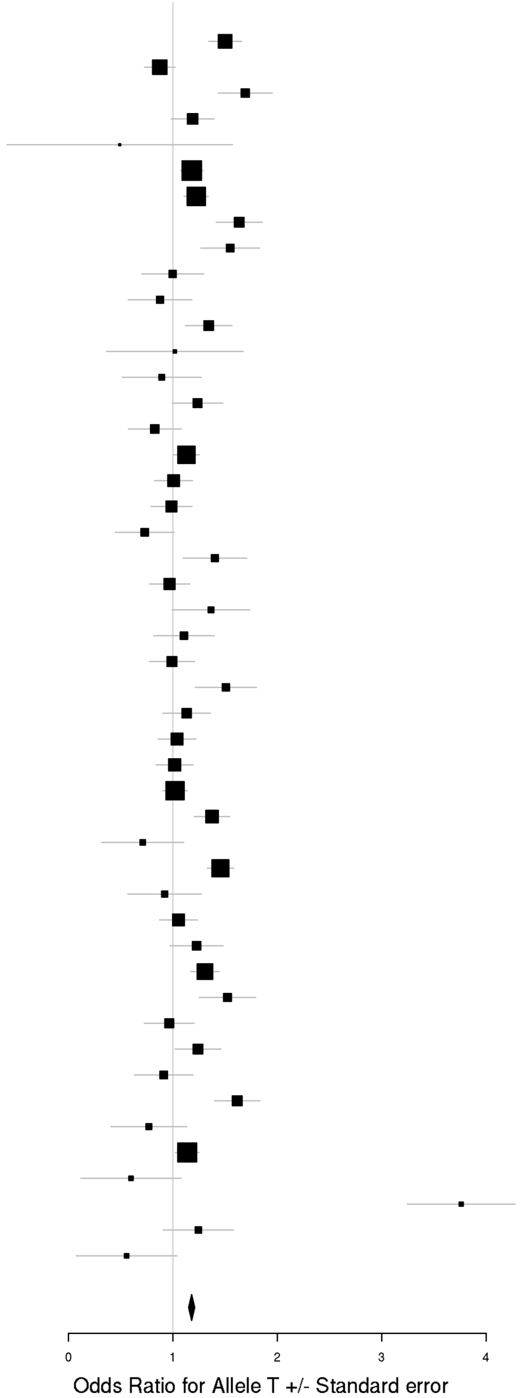

### i) Cross-Disorder SCZ-BIP-rMDD – European + East Asian ancestry

rs8040598 (A/G)
Cross-Disorder SCZ-BIP-RMDD

| Dx | Study | HC F N | HC M N | PT F N | PT M N | p-value |
|---|---|---|---|---|---|---|
| SCZ | aarh | 391 | 463 | 387 | 465 | 1.79E-02 |
| SCZ | ajsz | 414 | 1171 | 327 | 567 | 9.91E-01 |
| SCZ | asrb | 153 | 121 | 148 | 280 | 5.70E-01 |
| SCZ | boco | 160 | 160 | 761 | 952 | 8.52E-01 |
| SCZ | buls | 315 | 288 | 104 | 91 | 2.83E-01 |
| SCZ | cims | 20 | 33 | 11 | 52 | 3.20E-01 |
| SCZ | clm2 | 681 | 668 | 923 | 2450 | 7.24E-01 |
| SCZ | clo3 | 692 | 824 | 629 | 1442 | 7.56E-01 |
| SCZ | denm | 188 | 259 | 193 | 266 | 2.63E-01 |
| SCZ | dubl | 318 | 129 | 79 | 176 | 4.90E-01 |
| SCZ | edin | 37 | 37 | 98 | 264 | 8.31E-01 |
| SCZ | egcu | 828 | 299 | 169 | 62 | 6.33E-01 |
| SCZ | ersw | 121 | 178 | 92 | 142 | 4.48E-02 |
| SCZ | fi3m | 469 | 457 | 85 | 101 | 7.04E-01 |
| SCZ | fii6 | 594 | 437 | 153 | 207 | 9.44E-02 |
| SCZ | gras | 415 | 707 | 342 | 693 | 2.76E-01 |
| SCZ | irwt | 434 | 546 | 424 | 855 | 3.93E-01 |
| SCZ | lacw | 23 | 21 | 51 | 104 | 8.93E-01 |
| SCZ | lie2 | 144 | 124 | 37 | 94 | 8.01E-02 |
| SCZ | lie5 | 204 | 182 | 126 | 363 | 1.15E-01 |
| SCZ | mgs2 | 527 | 381 | 783 | 1777 | 6.40E-01 |
| SCZ | msaf | 57 | 79 | 121 | 198 | 9.62E-01 |
| SCZ | munc | 156 | 136 | 150 | 266 | 7.86E-01 |
| SCZ | pewb | 885 | 856 | 140 | 396 | 2.76E-01 |
| SCZ | pews | 96 | 138 | 63 | 87 | 9.01E-01 |
| SCZ | s234 | 517 | 548 | 744 | 1131 | 3.86E-02 |
| SCZ | swe5 | 585 | 654 | 690 | 1034 | 7.45E-01 |
| SCZ | swe6 | 259 | 283 | 360 | 511 | 1.69E-01 |
| SCZ | top8 | 102 | 101 | 160 | 214 | 5.05E-01 |
| SCZ | ucla | 278 | 282 | 170 | 522 | 1.12E-01 |
| SCZ | umeb | 185 | 150 | 110 | 156 | 7.40E-01 |
| SCZ | umes | 269 | 235 | 72 | 93 | 1.49E-02 |
| SCZ | hok2 | 1344 | 668 | 154 | 322 | 1.30E-02 |
| SCZ | tcr1 | 336 | 570 | 289 | 563 | 7.50E-02 |
| BIP | bmau | 895 | 895 | 184 | 140 | 3.38E-02 |
| BIP | bmg2 | 210 | 226 | 117 | 64 | 1.41E-01 |
| BIP | bmg3 | 158 | 54 | 288 | 197 | 8.90E-01 |
| BIP | bmpo | 213 | 476 | 250 | 160 | 5.69E-01 |
| BIP | bmsp | 144 | 147 | 138 | 114 | 8.53E-01 |
| BIP | bonn | 151 | 142 | 345 | 318 | 1.58E-01 |
| BIP | dub1 | 281 | 115 | 77 | 70 | 6.79E-01 |
| BIP | fran | 934 | 693 | 262 | 189 | 4.42E-01 |
| BIP | gain | 85 | 65 | 314 | 294 | 9.12E-01 |
| BIP | gsk1 | 408 | 299 | 415 | 239 | 3.77E-01 |
| BIP | may1 | 375 | 374 | 560 | 374 | 8.14E-01 |
| BIP | mich | 69 | 60 | 9 | 3 | 4.69E-01 |
| BIP | rom3 | 107 | 87 | 135 | 96 | 8.33E-01 |
| BIP | swa2 | 581 | 572 | 496 | 350 | 2.06E-01 |
| BIP | swei | 873 | 898 | 765 | 459 | 8.96E-02 |
| BIP | top8 | 128 | 155 | 81 | 56 | 2.13E-01 |
| BIP | ucl2 | 243 | 442 | 397 | 324 | 5.34E-01 |
| BIP | ume4 | 139 | 121 | 299 | 184 | 5.56E-01 |
| BIP | usc2 | 659 | 438 | 635 | 659 | 2.29E-01 |
| rMDD | boma | 131 | 149 | 299 | 148 | 6.11E-01 |
| rMDD | cof3 | 60 | 64 | 53 | 37 | 4.09E-01 |
| rMDD | gens | 130 | 195 | 711 | 290 | 2.29E-01 |
| rMDD | gep3 | 382 | 361 | 260 | 85 | 1.00E+ |
| rMDD | grnd | 199 | 212 | 677 | 140 | 3.53E-01 |
| rMDD | gsk2 | 134 | 65 | 438 | 227 | 7.99E-01 |
| rMDD | mmi2 | 278 | 228 | 230 | 173 | 3.05E-01 |
| rMDD | mmo4 | 185 | 193 | 105 | 89 | 5.09E-01 |
| rMDD | qi3c | 369 | 209 | 326 | 253 | 8.20E-01 |
| rMDD | qi6c | 363 | 221 | 156 | 85 | 3.74E-01 |
| rMDD | rad3 | 694 | 485 | 1005 | 417 | 5.90E-01 |
| rMDD | rage | 29 | 34 | 180 | 73 | 9.60E-01 |
| rMDD | rai2 | 172 | 154 | 87 | 19 | 3.01E-01 |
| rMDD | rau2 | 196 | 182 | 175 | 47 | 3.80E-01 |
| rMDD | rde4 | 208 | 303 | 41 | 18 | 4.18E-01 |
| rMDD | rot4 | 571 | 422 | 57 | 14 | 4.12E-01 |
| rMDD | shp0 | 494 | 637 | 128 | 52 | 8.48E-01 |
| | **Meta** | **23425** | **22558** | **19829** | **23353** | **3.90E-07** |

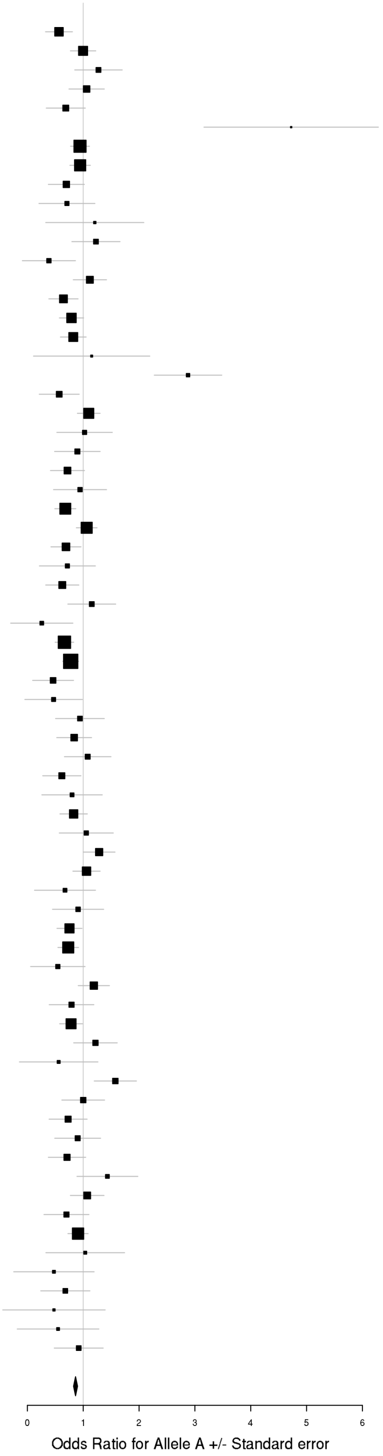

rs73033497 (A/T)
Cross-Disorder SCZ-BIP-RMDD

| Dx | Study | HC F N | HC M N | PT F N | PT M N | p-value |
|---|---|---|---|---|---|---|
| SCZ | aarh | 383 | 455 | 380 | 454 | 3.47E-01 |
| SCZ | ajsz | 410 | 1155 | 320 | 561 | 6.30E-01 |
| SCZ | buls | 312 | 281 | 101 | 91 | 5.29E-01 |
| SCZ | cims | 20 | 33 | 11 | 52 | 1.69E-01 |
| SCZ | cou3 | 89 | 73 | 200 | 310 | 4.88E-01 |
| SCZ | dubl | 315 | 129 | 78 | 175 | 6.91E-02 |
| SCZ | edin | 37 | 37 | 98 | 262 | 4.76E-01 |
| SCZ | egcu | 815 | 294 | 165 | 61 | 3.88E-01 |
| SCZ | ersw | 120 | 178 | 91 | 140 | 6.08E-01 |
| SCZ | fi3m | 464 | 452 | 83 | 99 | 3.87E-01 |
| SCZ | irwt | 432 | 544 | 424 | 847 | 5.10E-02 |
| SCZ | s234 | 508 | 543 | 732 | 1109 | 1.46E-01 |
| SCZ | swe5 | 580 | 651 | 676 | 1015 | 5.72E-04 |
| SCZ | swe6 | 258 | 280 | 359 | 502 | 2.69E-01 |
| SCZ | top8 | 101 | 101 | 158 | 212 | 1.21E-01 |
| BIP | bmg3 | 154 | 53 | 282 | 194 | 4.10E-01 |
| BIP | dub1 | 280 | 112 | 76 | 68 | 3.99E-01 |
| BIP | gain | 84 | 64 | 311 | 288 | 1.09E-01 |
| BIP | may1 | 373 | 368 | 552 | 366 | 1.53E-01 |
| BIP | swei | 854 | 891 | 757 | 453 | 9.57E-01 |
| BIP | top8 | 127 | 150 | 79 | 55 | 5.81E-01 |
| BIP | ucl2 | 241 | 436 | 387 | 319 | 2.61E-01 |
| rMDD | gens | 129 | 193 | 702 | 291 | 3.21E-01 |
| rMDD | shp0 | 485 | 623 | 125 | 51 | 2.25E-01 |
| | **Meta** | **7551** | **8096** | **7136** | **7975** | **6.22E-07** |

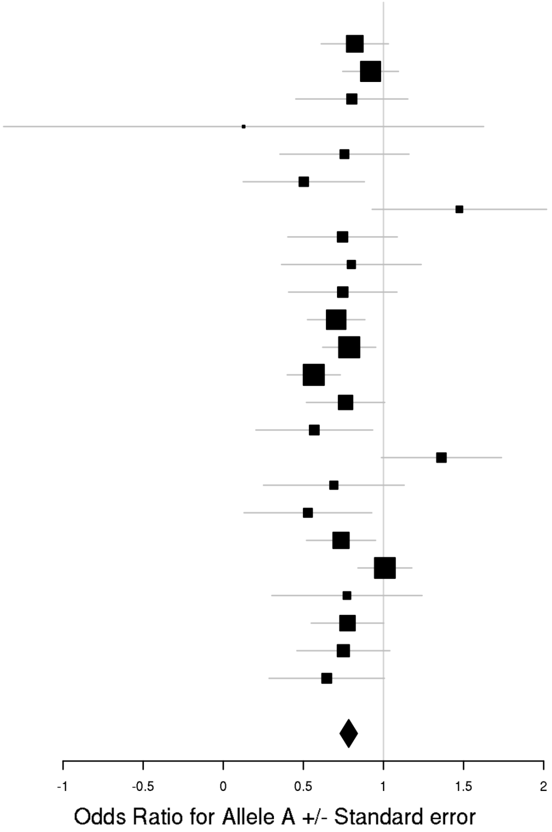

**Supplementary Figure 23. X chromosome model comparisons in PGC**

Abbreviations: BIP = Bipolar Disorder; (r)MDD = (recurrent) Major Depressive Disorder; SCZ = Schizophrenia; HC F N = number of female healthy controls; HC M N = number of male healthy controls; PT F N = number of female patients; PT M N = number of male patients; Study = cohort abbreviation used by PGC; Meta = meta-analysis results

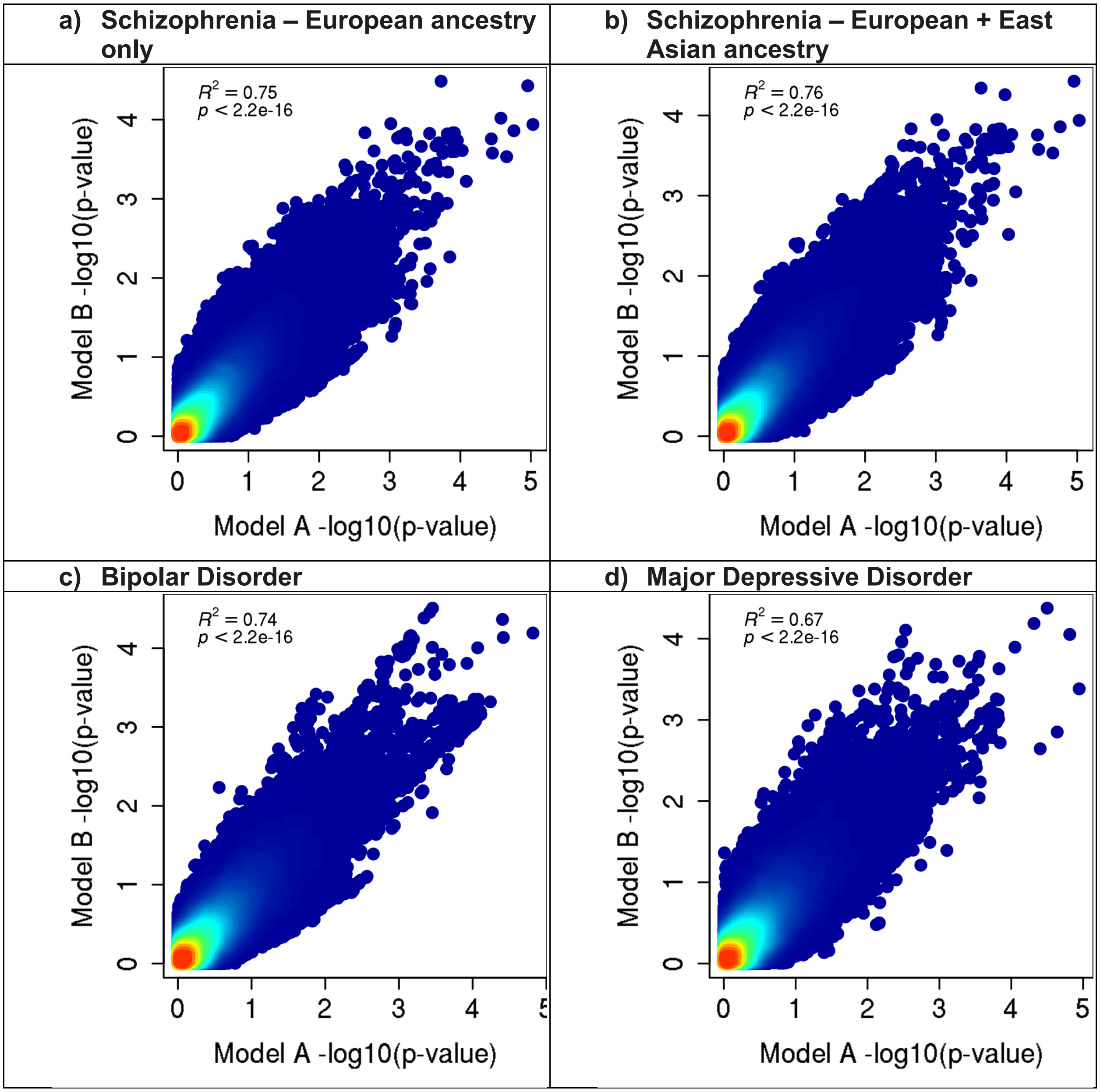

**e) Recurrent Major Depressive Disorder**

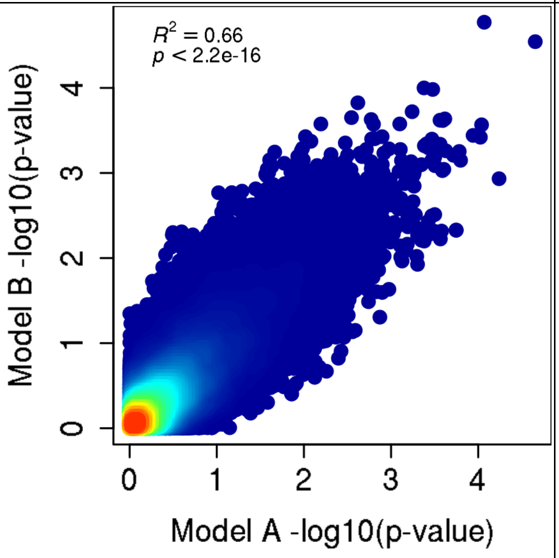


**f) Cross-Disorder SCZ-BIP-MDD – European ancestry only**

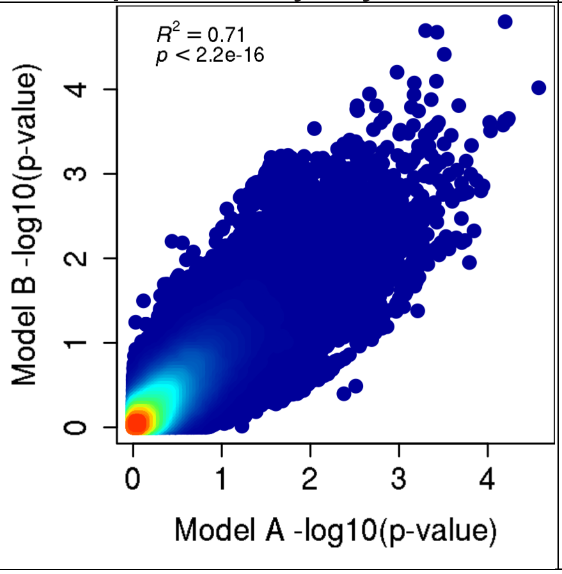


**g) Cross-Disorder SCZ-BIP-MDD – European + East Asian ancestry**

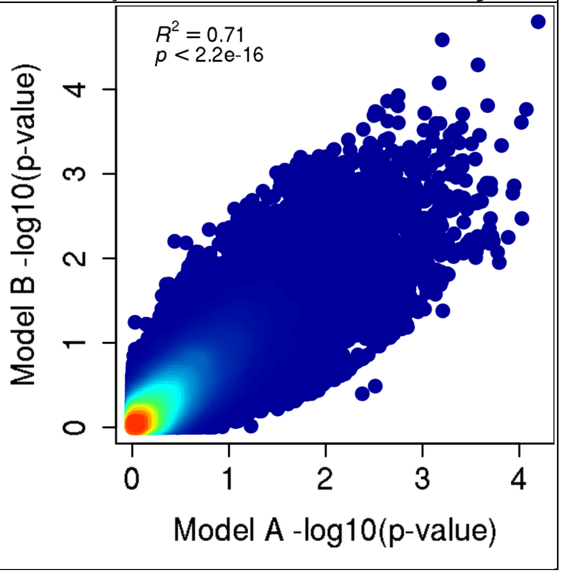

| h) Cross-Disorder SCZ-BIP-rMDD – European ancestry only | i) Cross-Disorder SCZ-BIP-rMDD – European + East Asian ancestry |
| --- | --- |
| 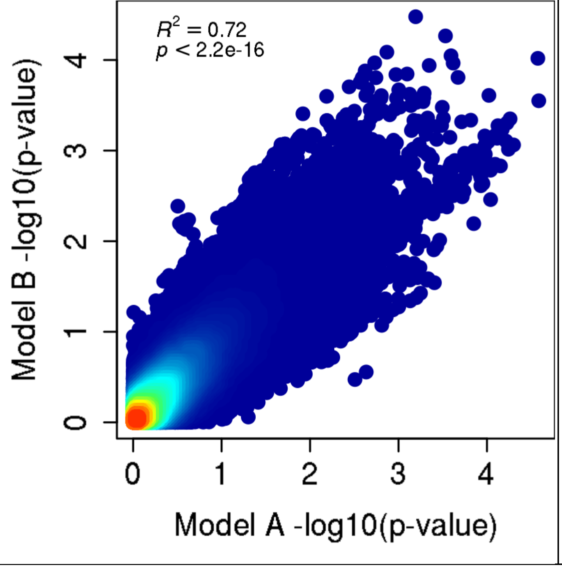 | 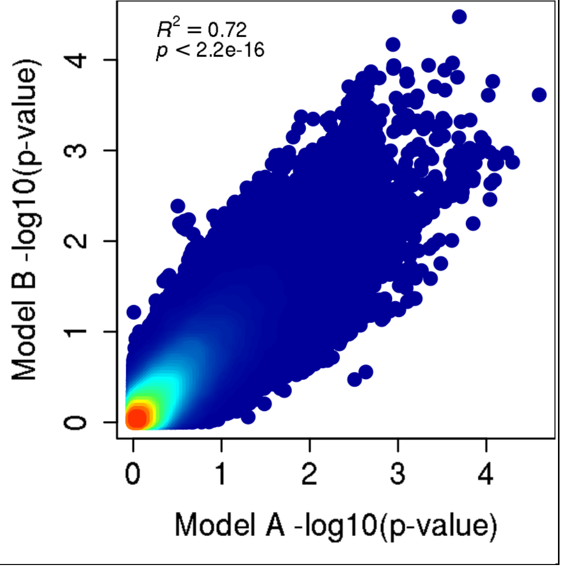 |

**Supplementary Figure 24. Manhattan plots for gene-based GxS tests in PGC**

These analyses were carried out in MAGMA on the genomic control output with INFO score > 0.6, *European ancestry only*, and autosomal SNPs only, with the MHC region included.

Negative log10-transformed p-values for each gene (y-axis) are plotted by chromosomal position (x-axis). Each dot represents a gene, and the solid red and dotted blue horizontal lines represent the thresholds for genome-wide significant association ($p = 2.57{\times}10^{-6}$) and suggestive association ($p = 1{\times}10^{-5}$), respectively.

Plots were generated using the plot package in R.

Abbreviations: BIP = Bipolar Disorder; (r)MDD = (recurrent) Major Depressive Disorder; SCZ = Schizophrenia; SLTM = SAFB Like Transcription Modulator

**a) Schizophrenia**

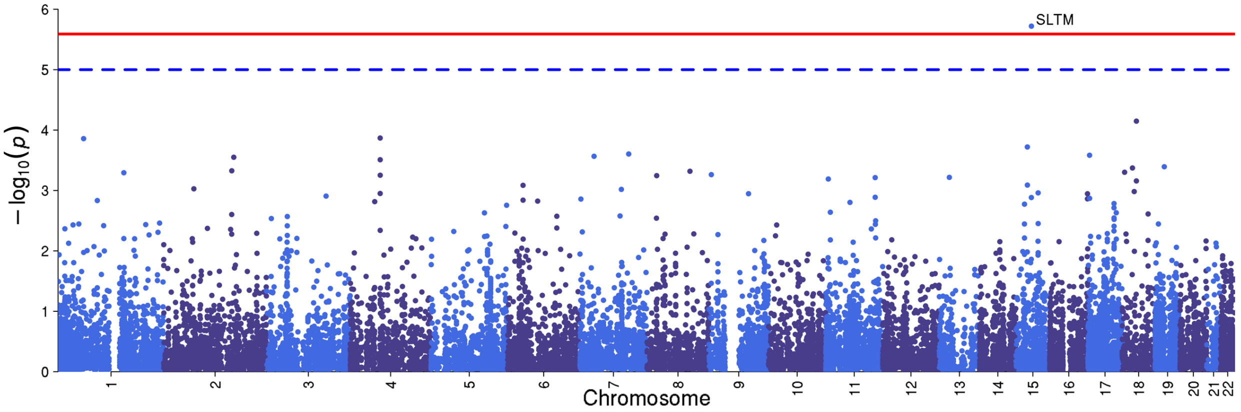


**b) Bipolar Disorder**

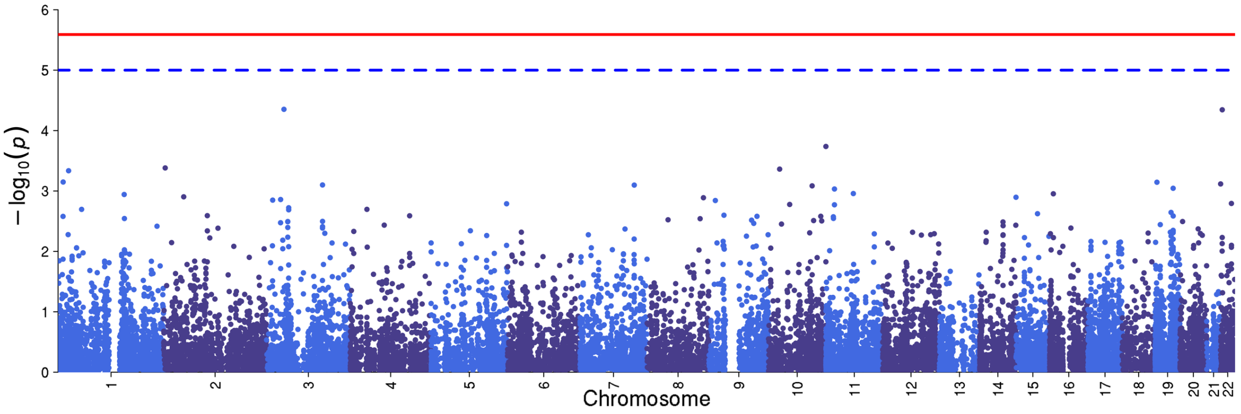

c) Major Depressive Disorder

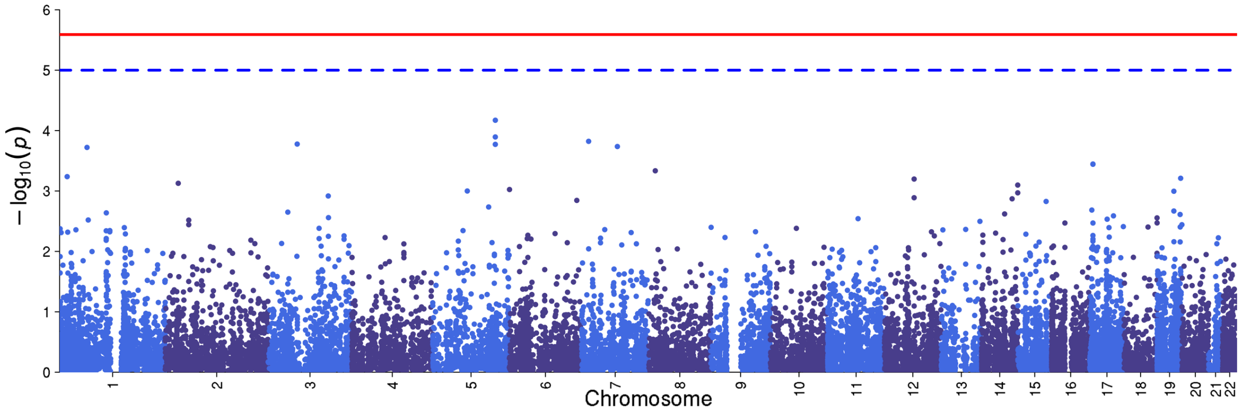


d) Recurrent Major Depressive Disorder

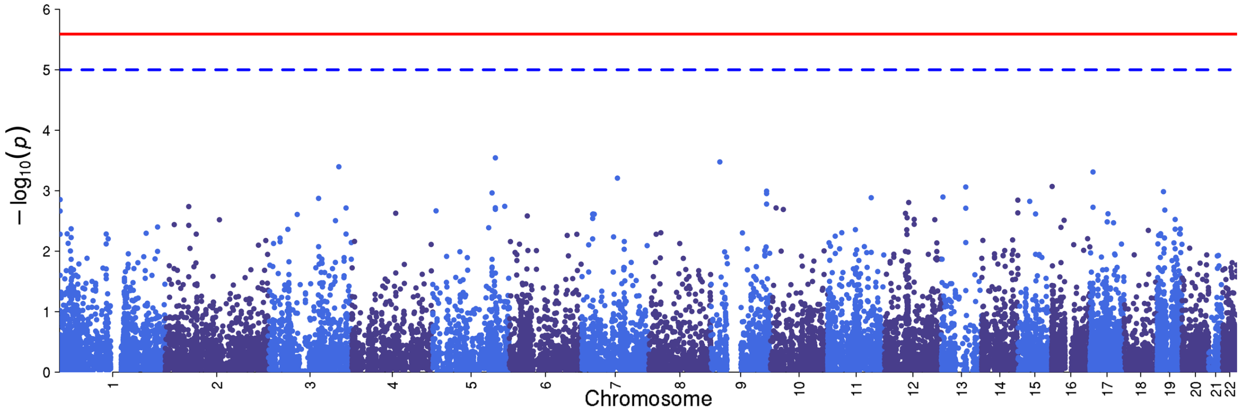


e) Cross-Disorder SCZ-BIP-MDD

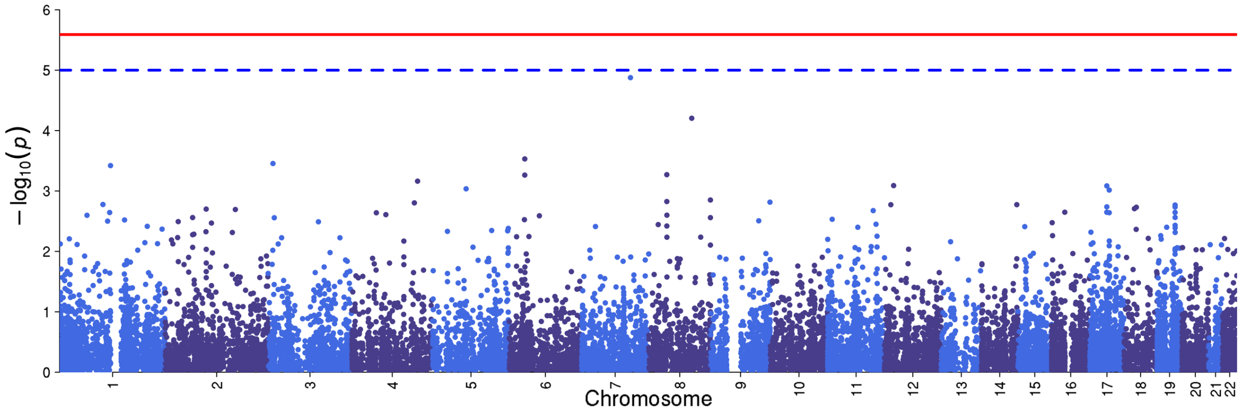

### f) Cross-Disorder SCZ-BIP-rMDD

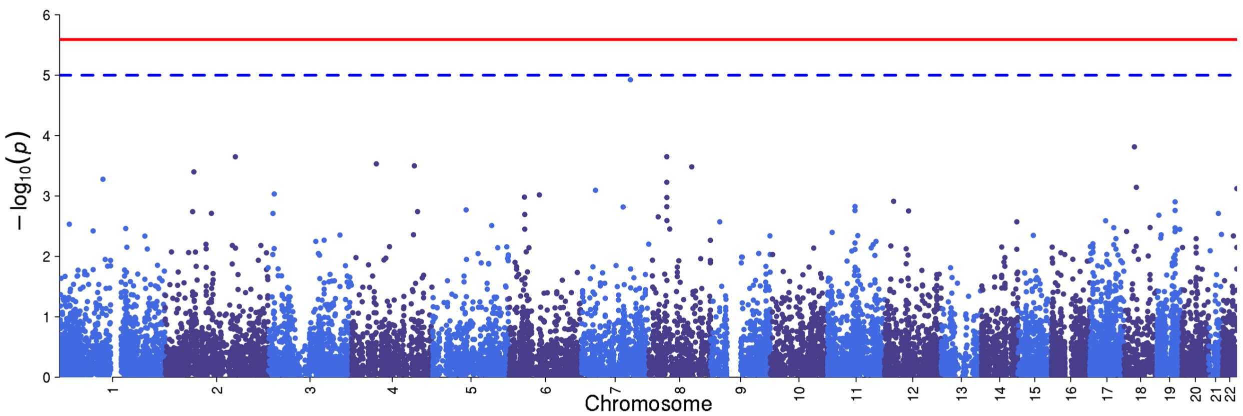


### g) Omnibus Test SCZ-BIP-MDD

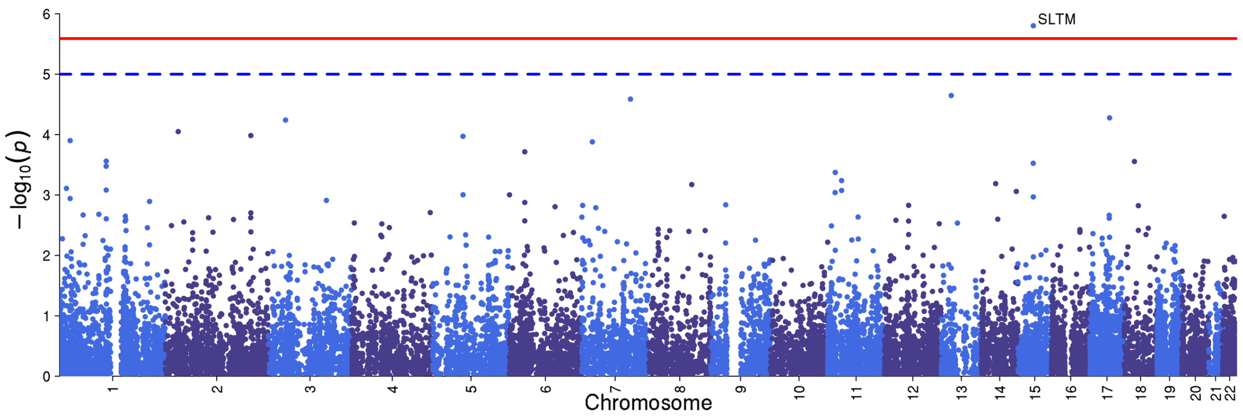


### h) Omnibus Test SCZ-BIP-rMDD

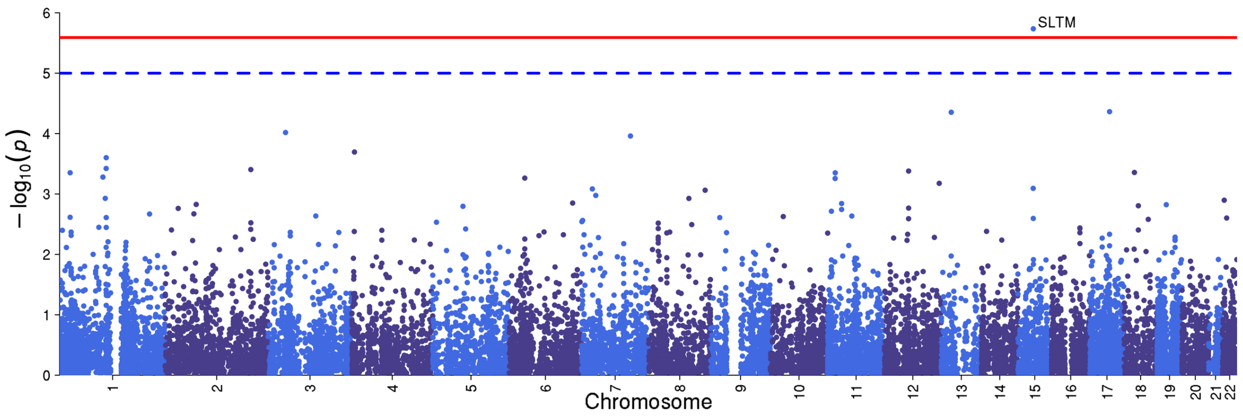

**Acknowledgements**

The PGC data were analyzed on the Genetic Cluster Computer (GCC) (http://www.geneticcluster.org), which is supported by a Netherlands Organisation for Scientific Research 'Medium Investment' grant (480-05-003) to Prof Danielle Posthuma, by the VU University Amsterdam, and by the Dutch Brain Foundation to Prof Roel Ophoff. The GCC is hosted by the Dutch National Computing and Networking Services.

This study makes use of data generated by the Wellcome Trust Case-Control Consortium.(50-52) Data from (53) were excluded. A full list of the investigators who contributed to the generation of the data is available from www.wtccc.org.uk. Funding for the project was provided by the Wellcome Trust under award 076113, 085475 and 090355.

The Finnish schizophrenia data used for the research were obtained from the THL Biobank. We thank all study participants for their generous participation in the THL Psychiatric Family Collections, the National FINRISK Study, Health 2000, and Northern Finland Birth Cohorts studies.

Data have also been provided by the Study of Health in Pomerania (SHIP) from the Community Medicine Research Alliance of the Medical Faculty at the Ernst Moritz Arndt University of Greifswald. Funding for SHIP was provided by BMBF – Federal Ministry for Education and Research under subsidy identification codes 01ZZ9603, 01ZZ0103, and 01ZZ0701.

Funding support for the companion studies, Genome-Wide Association Study of Schizophrenia (GAIN) and Molecular Genetics of Schizophrenia - nonGAIN Sample (MGS_nonGAIN), was provided by Genomics Research Branch at National Institute of Mental Health (NIMH) (see below) and the genotyping and analysis of samples was provided through the Genetic Association Information Network (GAIN) and under the MGS U01 MH79469 (PV Gejman) and U01 MH79470 (DF Levinson), and R01 MH81800 (PV Gejman). Assistance with data cleaning was provided by the National Center for Biotechnology Information. The GAIN and MGS dataset(s) used for the analyses described in this manuscript were obtained from the database of Genotype and Phenotype (dbGaP) found at http://www.ncbi.nlm.nih.gov/gap through dbGaP accession numbers **phs000021.v2.p1** (GAIN) and **phs000167.v1.p1** (MGS_nonGAIN). Samples and associated phenotype data for the Genome-Wide Association of Schizophrenia Study were provided by the Molecular Genetics of Schizophrenia Collaboration (PI: Pablo V. Gejman, Evanston Northwestern Healthcare (ENH) and Northwestern University, Evanston, Illinois, USA). Samples and associated phenotype data for the MGS GWAS study were collected under the following grants: NIMH Schizophrenia Genetics Initiative U01s: MH46276 (CR Cloninger), MH46289 (C Kaufmann), and MH46318 (MT Tsuang); and MGS Part 1 (MGS1) and Part 2 (MGS2) R01s: MH67257 (NG Buccola), MH59588 (BJ Mowry), MH59571 (PV Gejman), MH59565 (Robert Freedman), MH59587 (F Amin), MH60870 (WF Byerley), MH59566 (DW Black), MH59586 (JM Silverman), MH61675 (DF Levinson), and MH60879 (CR Cloninger). Further details of collection sites, individuals, and institutions may be found in data supplement Table 1 of Sanders et al. (2008; PMID: 18198266) and at the study dbGaP pages.

Funding support for the Whole Genome Association Study of Bipolar Disorder was provided by the NIMH and the genotyping of samples was provided through the GAIN. The datasets used for the analyses described in this manuscript were obtained from the dbGaP found at http://www.ncbi.nlm.nih.gov/gap through dbGaP accession number **phs000017.v1.p1**. Samples and associated phenotype data for the Collaborative Genomic Study of Bipolar Disorder were provided by the The NIMH Genetics Initiative for Bipolar Disorder. Data and biomaterials were collected in four projects that participated in NIMH Bipolar Disorder Genetics Initiative. From 1991-98, the Principal Investigators and Co-Investigators were: Indiana University, Indianapolis, IN, U01 MH46282, John Nurnberger, M.D., Ph.D., Marvin Miller, M.D., and Elizabeth Bowman, M.D.; Washington University, St. Louis, MO, U01 MH46280, Theodore Reich, M.D., Allison Goate, Ph.D., and John Rice, Ph.D.; Johns Hopkins University, Baltimore, MD U01 MH46274, J. Raymond DePaulo, Jr., M.D., Sylvia Simpson, M.D., MPH, and Colin Stine, Ph.D.; NIMH Intramural Research Program, Clinical Neurogenetics Branch, Bethesda, MD, Elliot Gershon, M.D., Diane Kazuba, B.A., and Elizabeth Maxwell, M.S.W. Data and biomaterials were collected as part of ten projects that participated in the NIMH Bipolar Disorder Genetics Initiative. From 1999-03, the Principal Investigators and Co-Investigators were: Indiana University, Indianapolis, IN, R01 MH59545, John Nurnberger, M.D., Ph.D., Marvin J. Miller, M.D., Elizabeth S. Bowman, M.D., N. Leela Rau, M.D., P. Ryan Moe, M.D., Nalini Samavedy, M.D., Rif El-Mallakh, M.D. (at University of Louisville), Husseini Manji, M.D. (at Wayne State University), Debra A. Glitz, M.D. (at Wayne State University), Eric T. Meyer, M.S., Carrie Smiley, R.N., Tatiana Foroud, Ph.D., Leah Flury, M.S., Danielle M. Dick, Ph.D., Howard Edenberg, Ph.D.; Washington University, St. Louis, MO, R01 MH059534, John Rice, Ph.D, Theodore Reich, M.D., Allison Goate, Ph.D., Laura Bierut, M.D. ; Johns Hopkins University, Baltimore, MD, R01 MH59533, Melvin McInnis M.D. , J. Raymond DePaulo, Jr., M.D., Dean F. MacKinnon, M.D., Francis M. Mondimore, M.D., James B. Potash, M.D., Peter P. Zandi, Ph.D, Dimitrios Avramopoulos, and Jennifer Payne; University of Pennsylvania, PA, R01 MH59553, Wade Berrettini M.D.,Ph.D. ; University of California at Irvine, CA, R01 MH60068, William Byerley M.D., and Mark Vawter M.D. ; University of Iowa, IA, R01 MH059548, William Coryell M.D. , and Raymond Crowe M.D.; University of Chicago, IL, R01 MH59535, Elliot Gershon, M.D., Judith Badner Ph.D. , Francis McMahon M.D. , Chunyu Liu Ph.D., Alan Sanders M.D., Maria Caserta, Steven Dinwiddie M.D., Tu Nguyen, Donna Harakal; University of California at San Diego, CA, R01 MH59567, John Kelsoe, M.D., Rebecca McKinney, B.A.; Rush University, IL, R01 MH059556, William Scheftner M.D. , Howard M. Kravitz, D.O., M.P.H., Diana Marta, B.S., Annette Vaughn- Brown, MSN, RN, and Laurie Bederow, MA; NIMH Intramural Research Program, Bethesda, MD, 1Z01MH002810-01, Francis J. McMahon, M.D., Layla Kassem, PsyD, Sevilla Detera-Wadleigh, Ph.D, Lisa Austin,Ph.D, Dennis L. Murphy, M.D.

### Consortium Membership

#### Schizophrenia Working Group of the Psychiatric Genomics Consortium

Stephan Ripke[1,2], Benjamin M. Neale[1,2,3,4], Aiden Corvin[5], James T. R. Walters[6], Kai-How Farh[1], Peter A. Holmans[6,7], Phil Lee[1,2,4], Brendan Bulik-Sullivan[1,2], David A. Collier[8,9], Hailiang Huang[1,3], Tune H. Pers[3,10,11], Ingrid Agartz[12,13,14], Esben Agerbo[15,16,17], Margot Albus[18], Madeline Alexander[19], Farooq Amin[20,21], Silviu A. Bacanu[22], Martin Begemann[23], Richard A. Belliveau Jr[2], Judit Bene[24,25], Sarah E. Bergen[2,26], Elizabeth Bevilacqua[2], Tim B. Bigdeli[22], Donald W. Black[27], Richard Bruggeman[28], Nancy G. Buccola[29], Randy L. Buckner[30,31,32], William Byerley[33], Wiepke Cahn[34], Guiqing Cai[35,36], Dominique Campion[37], Rita M. Cantor[38], Vaughan J. Carr[39,40], Noa Carrera[6], Stanley V. Catts[39,41], Kimberly D. Chambert[2], Raymond C. K. Chan[42], Ronald Y. L. Chen[43], Eric Y. H. Chen[43,44], Wei Cheng[45], Eric F. C. Cheung[46], Siow Ann Chong[47], C. Robert Cloninger[48], David Cohen[49], Nadine Cohen[50], Paul Cormican[5], Nick Craddock[6,7], James J. Crowley[51], David Curtis[52,53], Michael Davidson[54], Kenneth L. Davis[36], Franziska Degenhardt[55,56], Jurgen Del Favero[57], Ditte Demontis[17,58,59], Dimitris Dikeos[60], Timothy Dinan[61], Srdjan Djurovic[14,62], Gary Donohoe[5,63], Elodie Drapeau[36], Jubao Duan[64,65], Frank Dudbridge[66], Naser Durmishi[67], Peter Eichhammer[68], Johan Eriksson[69,70,71], Valentina Escott-Price[6], Laurent Essioux[72], Ayman H. Fanous[73,74,75,76], Martilias S. Farrell[51], Josef Frank[77], Lude Franke[78], Robert Freedman[79], Nelson B. Freimer[80], Marion Friedl[81], Joseph I. Friedman[36], Menachem Fromer[1,2,4,82], Giulio Genovese[2], Lyudmila Georgieva[6], Ina Giegling[81,83], Paola Giusti-Rodríguez[51], Stephanie Godard[84], Jacqueline I. Goldstein[1,3], Vera Golimbet[85], Srihari Gopal[86], Jacob Gratten[87], Lieuwe de Haan[88], Christian Hammer[23], Marian L. Hamshere[6], Mark Hansen[89], Thomas Hansen[17,90], Vahram Haroutunian[36,91,92], Annette M. Hartmann[81], Frans A. Henskens[39,93,94], Stefan Herms[55,56,95], Joel N. Hirschhorn[3,11,96], Per Hoffmann[55,56,95], Andrea Hofman[55,56], Mads V. Hollegaard[97], David M. Hougaard[97], Masashi Ikeda[98], Inge Joa[99], Antonio Julià[100], René S. Kahn[34], Luba Kalaydjieva[101,102], Sena Karachanak-Yankova[103], Juha Karjalainen[78], David Kavanagh[6], Matthew C. Keller[104], James L. Kennedy[105,106,107], Andrey Khrunin[108], Yunjung Kim[51], Janis Klovins[109], James A. Knowles[110], Bettina Konte[81], Vaidutis Kucinskas[111], Zita Ausrele Kucinskiene[111], Hana Kuzelova-Ptackova[112], Anna K. Kähler[26], Claudine Laurent[19,113], Jimmy Lee Chee Keong[47,114], S. Hong Lee[87], Sophie E. Legge[6], Bernard Lerer[115], Miaoxin Li[43,44,116], Tao Li[117], Kung-Yee Liang[118], Jeffrey Lieberman[119], Svetlana Limborska[108], Carmel M. Loughland[39,120], Jan Lubinski[121], Jouko Lönnqvist[122], Milan Macek Jr[112], Patrik K. E. Magnusson[26], Brion S. Maher[123], Wolfgang Maier[124], Jacques Mallet[125], Sara Marsal[100], Manuel Mattheisen[17,58,59,126], Morten Mattingsdal[14,127], Robert W. McCarley[128,129,†], Colm McDonald[130], Andrew M. McIntosh[131,132], Sandra Meier[77], Carin J. Meijer[88], Bela Melegh[24,25], Ingrid Melle[14,133], Raquelle I. Mesholam-Gately[128,134], Andres Metspalu[135], Patricia T. Michie[39,136], Lili Milani[135], Vihra Milanova[137], Younes Mokrab[8], Derek W. Morris[5,63], Ole Mors[17,58,138], Kieran C. Murphy[139], Robin M. Murray[140], Inez Myin-Germeys[141], Bertram Müller-Myhsok[142,143,144], Mari Nelis[135], Igor Nenadic[145], Deborah A. Nertney[146], Gerald Nestadt[147], Kristin K. Nicodemus[148], Liene Nikitina-Zake[109], Laura Nisenbaum[149], Annelie Nordin[150], Eadbhard O'Callaghan[151], Colm O'Dushlaine[2], F. Anthony O'Neill[152], Sang-Yun Oh[153], Ann Olincy[79], Line Olsen[17,90], Jim Van Os[141,154], Psychosis Endophenotypes International Consortium[155], Christos Pantelis[39,156], George N. Papadimitriou[60], Sergi Papiol[23], Elena Parkhomenko[36], Michele T. Pato[110], Tiina Paunio[157,158], Milica Pejovic-Milovancevic[159], Diana O. Perkins[160], Olli Pietiläinen[158,161,] Jonathan Pimm[53], Andrew J. Pocklington[6], John Powell[140], Alkes Price[3,162], Ann E. Pulver[147], Shaun M. Purcell[82], Digby Quested[163], Henrik B. Rasmussen[17,90], Abraham Reichenberg[36], Mark A. Reimers[164], Alexander L. Richards[6], Joshua L.

Roffman[30,32], Panos Roussos[82,165], Douglas M. Ruderfer[6,82], Veikko Salomaa[71], Alan R. Sanders[64,65], Ulrich Schall[39,120], Christian R. Schubert[166], Thomas G. Schulze[77,167], Sibylle G. Schwab[168], Edward M. Scolnick[2], Rodney J. Scott[39,169,170], Larry J. Seidman[128,134,†,] Jianxin Shi[171], Engilbert Sigurdsson[172], Teimuraz Silagadze[173], Jeremy M. Silverman[36,174], Kang Sim[47], Petr Slominsky[108], Jordan W. Smoller[2,4], Hon-Cheong So[43], Chris C. A. Spencer[175], Eli A. Stahl[3,82], Hreinn Stefansson[176], Stacy Steinberg[176], Elisabeth Stogmann[177], Richard E. Straub[178], Eric Strengman[179,34], Jana Strohmaier[77], T. Scott Stroup[119], Mythily Subramaniam[47], Jaana Suvisaari[122], Dragan M. Svrakic[48], Jin P. Szatkiewicz[51], Erik Söderman[12], Srinivas Thirumalai[180], Draga Toncheva[103], Sarah Tosato[181], Juha Veijola[182,183], John Waddington[184], Dermot Walsh[185], Dai Wang[86], Qiang Wang[117], Bradley T. Webb[22], Mark Weiser[54], Dieter B. Wildenauer[186], Nigel M. Williams[6], Stephanie Williams[51], Stephanie H. Witt[77], Aaron R. Wolen[164], Emily H. M. Wong[43], Brandon K. Wormley[22], Hualin Simon Xi[187], Clement C. Zai[105,106], Xuebin Zheng[188], Fritz Zimprich[177], Naomi R. Wray[87], Kari Stefansson[176], Peter M. Visscher[87], Wellcome Trust Case-Control Consortium 2189, Rolf Adolfsson[150], Ole A. Andreassen[14,133], Douglas H. R. Blackwood[132], Elvira Bramon[190], Joseph D. Buxbaum[35,36,91,191], Anders D. Børglum[17,58,59,138], Sven Cichon[55,56,95,192], Ariel Darvasi[193,†,] Enrico Domenici[194], Hannelore Ehrenreich[23], Tõnu Esko[3,11,96,135], Pablo V. Gejman[64,65], Michael Gill[5], Hugh Gurling[53], Christina M. Hultman[26], Nakao Iwata[98], Assen V. Jablensky[39,102,186,195], Erik G. Jönsson[12,14], Kenneth S. Kendler[196], George Kirov[6], Jo Knight[105,106,107], Todd Lencz[197,198,199], Douglas F. Levinson[19], Qingqin S. Li[86], Jianjun Liu[188,200], Anil K. Malhotra[197,198,199], Steven A. McCarroll[2,96], Andrew McQuillin[53], Jennifer L. Moran[2], Preben B. Mortensen[15,16,17], Sathish Periyasamy[87,209], Murray J. Cairns[210,211,212], Paul A. Tooney[210,211,212], Jing Qin Wu[210,211], Brian Kelly[212], Bryan J. Mowry[87,201], Markus M. Nöthen[55,56], Roel A. Ophoff[38,80,34], Michael J. Owen[6,7], Aarno Palotie[2,4,161], Carlos N. Pato[110], Tracey L. Petryshen[2,128,202], Danielle Posthuma[203,204,205], Marcella Rietschel[77], Brien P. Riley[196], Dan Rujescu[81,83], Pak C. Sham[43,44,116], Pamela Sklar[82,91,165], David St Clair[206], Daniel R. Weinberger[178,207], Jens R. Wendland[166], Thomas Werge[17,90,208], Mark J. Daly[1,2,3], Patrick F. Sullivan[26,51,160], Michael C. O'Donovan[6,7].
† deceased

*Wellcome Trust Case-Control Consortium 2*

Management Committee: Peter Donnelly[180,217], Ines Barroso[218], Jenefer M Blackwell[219,220], Elvira Bramon[196], Matthew A Brown[221], Juan P Casas[222,223], Aiden Corvin[5], Panos Deloukas[218], Audrey Duncanson[224], Janusz Jankowski[225], Hugh S Markus[226], Christopher G Mathew[227], Colin N A Palmer[228], Robert Plomin[9], Anna Rautanen[180], Stephen J Sawcer[229], Richard C Trembath[227], Ananth C Viswanathan[230,231], Nicholas W Wood[232]. Data and Analysis Group: Chris C A Spencer[180], Gavin Band[180], Céline Bellenguez[180], Peter Donnelly[180,217], Colin Freeman[180], Eleni Giannoulatou[180], Garrett Hellenthal[180], Richard Pearson[180], Matti Pirinen[180], Amy Strange[180], Zhan Su[180], Damjan Vukcevic[180]. DNA, Genotyping, Data QC, and Informatics: Cordelia Langford[218], Ines Barroso[218], Hannah Blackburn[218], Suzannah J Bumpstead[218], Panos Deloukas[218], Serge Dronov[218], Sarah Edkins[218], Matthew Gillman[218], Emma Gray[218], Rhian Gwilliam[218], Naomi Hammond[218], Sarah E Hunt[218], Alagurevathi Jayakumar[218], Jennifer Liddle[218], Owen T McCann[218], Simon C Potter[218], Radhi Ravindrarajah[218], Michelle Ricketts[218], Avazeh Tashakkori-Ghanbaria[218], Matthew Waller[218], Paul Weston[218], Pamela Whittaker[218], Sara Widaa[218]. Publications Committee: Christopher G Mathew[227], Jenefer M Blackwell[219,220], Matthew A Brown[221], Aiden Corvin[5], Mark I McCarthy[233], Chris C A Spencer[180].

*Psychosis Endophenotype International Consortium*
Maria J Arranz[156,234], Steven Bakker[101], Stephan Bender[235,236], Elvira Bramon[156,237,238], David A Collier[8,9], Benedicto Crespo-Facorro[239,240], Jeremy Hall[134], Conrad Iyegbe[156], Assen V Jablensky[241], René S Kahn[101], Luba Kalaydjieva[102,242], Stephen Lawrie[134], Cathryn M Lewis[156], Kuang Lin[156], Don H Linszen[243], Ignacio Mata[239,240], Andrew M McIntosh[134], Robin M Murray[142], Roel A Ophoff[80], Jim Van Os[143,156], John Powell[156], Dan Rujescu[81,83], Muriel Walshe[156], Matthias Weisbrod[236], Durk Wiersma[244].

**Affiliations:**
[1] Analytic and Translational Genetics Unit, Massachusetts General Hospital, Boston, Massachusetts 02114, USA.
[2] Stanley Center for Psychiatric Research, Broad Institute of MIT and Harvard, Cambridge, Massachusetts 02142, USA.
[3] Medical and Population Genetics Program, Broad Institute of MIT and Harvard, Cambridge, Massachusetts 02142, USA.
[4] Psychiatric and Neurodevelopmental Genetics Unit, Massachusetts General Hospital, Boston, Massachusetts 02114, USA.
[5] Neuropsychiatric Genetics Research Group, Department of Psychiatry, Trinity College Dublin, Dublin 8, Ireland.
[6] MRC Centre for Neuropsychiatric Genetics and Genomics, Institute of Psychological Medicine and Clinical Neurosciences, School of Medicine, Cardiff University, Cardiff CF24 4HQ, UK.
[7] National Centre for Mental Health, Cardiff University, Cardiff CF244HQ, UK.
[8] Eli Lilly and Company Limited, Erl Wood Manor, Sunninghill Road, Windlesham, Surrey GU20 6PH, UK.
[9] Social, Genetic and Developmental Psychiatry Centre, Institute of Psychiatry, King's College London, London SE5 8AF, UK.
[10] Center for Biological Sequence Analysis, Department of Systems Biology, Technical University of Denmark, 2800 Kongens Lyngby, Denmark.
[11] Division of Endocrinology and Center for Basic and Translational Obesity Research, Boston Children's Hospital, Boston, Massachusetts 02115, USA.
[12] Department of Clinical Neuroscience, Psychiatry Section, Karolinska Institutet, SE-17176 Stockholm, Sweden.
[13] Department of Psychiatry, Diakonhjemmet Hospital, 0319 Oslo, Norway.
[14] NORMENT, KG Jebsen Centre for Psychosis Research, Institute of Clinical Medicine, University of Oslo, 0424 Oslo, Norway.
[15] Centre for Integrative Register-based Research, CIRRAU, Aarhus University, 8210 Aarhus, Denmark.
[16] National Centre for Register-based Research, Aarhus University, 8210 Aarhus, Denmark.
[17] The Lundbeck Foundation Initiative for Integrative Psychiatric Research, iPSYCH, Denmark.
[18] State Mental Hospital, 85540 Haar, Germany.
[19] Department of Psychiatry and Behavioral Sciences, Stanford University, Stanford, California 94305, USA.
[20] Department of Psychiatry and Behavioral Sciences, Atlanta Veterans Affairs Medical Center, Atlanta, Georgia 30033, USA.
[21] Department of Psychiatry and Behavioral Sciences, Emory University, Atlanta, Georgia 30322, USA.

[22] Virginia Institute for Psychiatric and Behavioral Genetics, Department of Psychiatry, Virginia Commonwealth University, Richmond, Virginia 23298, USA.
[23] Clinical Neuroscience, Max Planck Institute of Experimental Medicine, Göttingen 37075, Germany.
[24] Department of Medical Genetics, University of Pécs, Pécs H-7624, Hungary.
[25] Szentagothai Research Center, University of Pécs, Pécs H-7624, Hungary.
[26] Department of Medical Epidemiology and Biostatistics, Karolinska Institutet, Stockholm SE-17177, Sweden.
[27] Department of Psychiatry, University of Iowa Carver College of Medicine, Iowa City, Iowa 52242, USA.
[28] University Medical Center Groningen, Department of Psychiatry, University of Groningen NL-9700 RB, The Netherlands.
[29] School of Nursing, Louisiana State University Health Sciences Center, New Orleans, Louisiana 70112, USA.
[30] Athinoula A. Martinos Center for Biomedical Imaging, Massachusetts General Hospital, Boston, Massachusetts 02129, USA.
[31] Center for Brain Science, Harvard University, Cambridge, Massachusetts 02138, USA.
[32] Department of Psychiatry, Massachusetts General Hospital, Boston, Massachusetts 02114, USA.
[33] Department of Psychiatry, University of California at San Francisco, San Francisco, California 94143, USA.
[34] University Medical Center Utrecht, Department of Psychiatry, Rudolf Magnus Institute of Neuroscience, 3584 Utrecht, The Netherlands.
[35] Department of Human Genetics, Icahn School of Medicine at Mount Sinai, New York, New York 10029, USA.
[36] Department of Psychiatry, Icahn School of Medicine at Mount Sinai, New York, New York 10029, USA.
[37] Centre Hospitalier du Rouvray and INSERM U1079 Faculty of Medicine, 76301 Rouen, France.
[38] Department of Human Genetics, David Geffen School of Medicine, University of California, Los Angeles, California 90095, USA.
[39] Schizophrenia Research Institute, Sydney, New South Wales 2010, Australia.
[40] School of Psychiatry, University of New South Wales, Sydney, New South Wales 2031, Australia.
[41] Royal Brisbane and Women's Hospital, University of Queensland, Brisbane (St Lucia), Queensland 4072, Australia.
[42] Institute of Psychology, Chinese Academy of Science, Beijing 100101, China.
[43] Department of Psychiatry, Li Ka Shing Faculty of Medicine, The University of Hong Kong, Hong Kong, SAR China.
[44] State Key Laboratory for Brain and Cognitive Sciences, Li Ka Shing Faculty of Medicine, The University of Hong Kong, Hong Kong, SAR China.
[45] Department of Computer Science, University of North Carolina, Chapel Hill, North Carolina 27514, USA.
[46] Castle Peak Hospital, Hong Kong, SAR China.
[47] Institute of Mental Health, Singapore 539747, Singapore.
[48] Department of Psychiatry, Washington University, St. Louis, Missouri 63110, USA.

[49] Department of Child and Adolescent Psychiatry, Assistance Publique Hopitaux de Paris, Pierre and Marie Curie Faculty of Medicine and Institute for Intelligent Systems and Robotics, Paris 75013, France.
[50] Blue Note Biosciences, Princeton, New Jersey 08540, USA
[51] Department of Genetics, University of North Carolina, Chapel Hill, North Carolina 27599-7264, USA.
[52] Department of Psychological Medicine, Queen Mary University of London, London E1 1BB, UK.
[53] Molecular Psychiatry Laboratory, Division of Psychiatry, University College London, London WC1E6JJ, UK.
[54] Sheba Medical Center, Tel Hashomer 52621, Israel.
[55] Department of Genomics, Life and Brain Center, D-53127 Bonn, Germany.
[56] Institute of Human Genetics, University of Bonn, D-53127 Bonn, Germany.
[57] Applied Molecular Genomics Unit, VIB Department of Molecular Genetics, University of Antwerp, B-2610 Antwerp, Belgium.
[58] Centre for Integrative Sequencing, iSEQ, Aarhus University, 8000 Aarhus C, Denmark.
[59] Department of Biomedicine, Aarhus University,8000 Aarhus C, Denmark.
[60] First Department of Psychiatry, University of Athens Medical School, Athens 11528, Greece.
[61] Department of Psychiatry, University College Cork, Co. Cork, Ireland.
[62] Department of Medical Genetics, Oslo University Hospital, 0424 Oslo, Norway.
[63] Cognitive Genetics and Therapy Group, School of Psychology and Discipline of Biochemistry, National University of Ireland Galway, Co. Galway, Ireland.
[64] Department of Psychiatry and Behavioral Neuroscience, University of Chicago, Chicago, Illinois 60637, USA.
[65] Department of Psychiatry and Behavioral Sciences, NorthShore University Health System, Evanston, Illinois 60201, USA.
[66] Department of Non-Communicable Disease Epidemiology, London School of Hygiene and Tropical Medicine, London WC1E 7HT, UK.
[67] Department of Child and Adolescent Psychiatry, University Clinic of Psychiatry, Skopje 1000, Republic of Macedonia.
[68] Department of Psychiatry, University of Regensburg, 93053 Regensburg, Germany.
[69] Department of General Practice, Helsinki University Central Hospital, University of Helsinki P.O. Box 20, Tukholmankatu 8 B, FI-00014, Helsinki, Finland
[70] Folkhälsan Research Center, Helsinki, Finland, Biomedicum Helsinki 1, Haartmaninkatu 8, FI-00290, Helsinki, Finland.
[71] National Institute for Health and Welfare, P.O. Box 30, FI-00271 Helsinki, Finland.
[72] Translational Technologies and Bioinformatics, Pharma Research and Early Development, F. Hoffman-La Roche, CH-4070 Basel, Switzerland.
[73] Department of Psychiatry, Georgetown University School of Medicine, Washington DC 20057, USA.
[74] Department of Psychiatry, Keck School of Medicine of the University of Southern California, Los Angeles, California 90033, USA.
[75] Department of Psychiatry, Virginia Commonwealth University School of Medicine, Richmond, Virginia 23298, USA.
[76] Mental Health Service Line, Washington VA Medical Center, Washington DC 20422, USA.

[77] Department of Genetic Epidemiology in Psychiatry, Central Institute of Mental Health, Medical Faculty Mannheim, University of Heidelberg, Heidelberg, D-68159 Mannheim, Germany.
[78] Department of Genetics, University of Groningen, University Medical Centre Groningen, 9700 RB Groningen, The Netherlands.
[79] Department of Psychiatry, University of Colorado Denver, Aurora, Colorado 80045, USA.
[80] Center for Neurobehavioral Genetics, Semel Institute for Neuroscience and Human Behavior, University of California, Los Angeles, California 90095, USA.
[81] Department of Psychiatry, University of Halle, 06112 Halle, Germany.
[82] Division of Psychiatric Genomics, Department of Psychiatry, Icahn School of Medicine at Mount Sinai, New York, New York, New York 10029, USA.
[83] Department of Psychiatry, University of Munich, 80336, Munich, Germany.
[84] Departments of Psychiatry and Human and Molecular Genetics, INSERM, Institut de Myologie, Hôpital de la Pitiè-Salpêtrière, Paris 75013, France.
[85] Mental Health Research Centre, Russian Academy of Medical Sciences, 115522 Moscow, Russia.
[86] Neuroscience Therapeutic Area, Janssen Research and Development, Raritan, New Jersey 08869, USA.
[87] Queensland Brain Institute, The University of Queensland, Brisbane, Queensland, 4072, Australia.
[88] Academic Medical Centre University of Amsterdam, Department of Psychiatry, 1105 AZ Amsterdam, The Netherlands.
[89] Illumina, La Jolla, California, California 92122, USA.
[90] Institute of Biological Psychiatry, Mental Health Centre Sct. Hans, Mental Health Services Copenhagen, Denmark-4000, Denmark.
[91] Friedman Brain Institute, Icahn School of Medicine at Mount Sinai, New York, New York 10029, USA.
[92] J. J. Peters VA Medical Center, Bronx, New York, New York 10468, USA.
[93] Priority Research Centre for Health Behaviour, University of Newcastle, Newcastle, New South Wales 2308, Australia.
[94] School of Electrical Engineering and Computer Science, University of Newcastle, Newcastle, New South Wales 2308, Australia.
[95] Division of Medical Genetics, Department of Biomedicine, University of Basel, Basel CH-4058, Switzerland.
[96] Department of Genetics, Harvard Medical School, Boston, Massachusetts, Massachusetts 02115, USA.
[97] Section of Neonatal Screening and Hormones, Department of Clinical Biochemistry, Immunology and Genetics, Statens Serum Institut, Copenhagen DK-2300, Denmark.
[98] Department of Psychiatry, Fujita Health University School of Medicine, Toyoake, Aichi, 470-1192, Japan.
[99] Regional Centre for Clinical Research in Psychosis, Department of Psychiatry, Stavanger University Hospital, 4011 Stavanger, Norway.
[100] Rheumatology Research Group, Vall d'Hebron Research Institute, Barcelona 08035, Spain.
[101] Centre for Medical Research, The University of Western Australia, Perth, Western Australia 6009, Australia.

[102] The Perkins Institute for Medical Research, The University of Western Australia, Perth, Western Australia 6009, Australia.
[103] Department of Medical Genetics, Medical University, Sofia 1431, Bulgaria.
[104] Department of Psychology, University of Colorado Boulder, Boulder, Colorado 80309, USA.
[105] Campbell Family Mental Health Research Institute, Centre for Addiction and Mental Health, Toronto, Ontario M5T 1R8, Canada.
[106] Department of Psychiatry, University of Toronto, Toronto, Ontario M5T 1R8, Canada.
[107] Institute of Medical Science, University of Toronto, Toronto, Ontario M5S 1A8, Canada.
[108] Institute of Molecular Genetics, Russian Academy of Sciences, Moscow 123182, Russia.
[109] Latvian Biomedical Research and Study Centre, Riga, LV-1067, Latvia.
[110] Department of Psychiatry and Zilkha Neurogenetics Institute, Keck School of Medicine at University of Southern California, Los Angeles, California 90089, USA.
[111] Faculty of Medicine, Vilnius University, LT-01513 Vilnius, Lithuania.
[112] Department of Biology and Medical Genetics, 2nd Faculty of Medicine and University Hospital Motol, 150 06 Prague, Czech Republic.
[113] Department of Child and Adolescent Psychiatry, Pierre and Marie Curie Faculty of Medicine, Paris 75013, France.
[114] Duke-National University of Singapore Graduate Medical School, Singapore 169857.
[115] Department of Psychiatry, Hadassah-Hebrew University Medical Center, Jerusalem 91120, Israel.
[116] Centre for Genomic Sciences, The University of Hong Kong, Hong Kong, SAR China.
[117] Mental Health Centre and Psychiatric Laboratory, West China Hospital, Sichuan University, Chengdu, 610041 Sichuan, China.
[118] Department of Biostatistics, Johns Hopkins University Bloomberg School of Public Health, Baltimore, Maryland 21205, USA.
[119] Department of Psychiatry, Columbia University, New York, New York 10032, USA.
[120] Priority Centre for Translational Neuroscience and Mental Health, University of Newcastle, Newcastle NSW2300, Australia.
[121] Department of Genetics and Pathology, International Hereditary Cancer Center, Pomeranian Medical University in Szczecin, 70-453 Szczecin, Poland.
[122] Department of Mental Health and Substance Abuse Services; National Institute for Health and Welfare, P.O. BOX30, FI-00271 Helsinki, Finland.
[123] Department of Mental Health, Bloomberg School of Public Health, Johns Hopkins University, Baltimore, Maryland 21205, USA.
[124] Department of Psychiatry, University of Bonn, D-53127 Bonn, Germany.
[125] Centre National de la Recherche Scientifique, Laboratoire de Génétique Moléculaire de la Neurotransmission et des Processus Neurodégénératifs, Hôpital de la Pitié Salpêtrière, 75013 Paris, France.
[126] Department of Genomics Mathematics, University of Bonn, D-53127 Bonn, Germany.
[127] Research Unit, Sørlandet Hospital, 4604 Kristiansand, Norway.
[128] Department of Psychiatry, Harvard Medical School, Boston, Massachusetts 02115, USA.
[129] VA Boston Health Care System, Brockton, Massachusetts 02301, USA.
[130] Department of Psychiatry, National University of Ireland Galway, Co. Galway, Ireland.
[131] Centre for Cognitive Ageing and Cognitive Epidemiology, University of Edinburgh, Edinburgh EH16 4SB, UK.
[132] Division of Psychiatry, University of Edinburgh, Edinburgh EH16 4SB, UK.

[133] Division of Mental Health and Addiction, Oslo University Hospital, 0424 Oslo, Norway.
[134] Massachusetts Mental Health Center Public Psychiatry Division of the Beth Israel Deaconess Medical Center, Boston, Massachusetts 02114, USA.
[135] Estonian Genome Center, University of Tartu, Tartu 50090, Estonia.
[136] School of Psychology, University of Newcastle, Newcastle, New South Wales 2308, Australia.
[137] First Psychiatric Clinic, Medical University, Sofia 1431, Bulgaria.
[138] Department of Psychiatry, Aarhus University Hospital, Denmark-8240 Risskov, Denmark.
[139] Department of Psychiatry, Royal College of Surgeons in Ireland, Dublin 2, Ireland.
[140] King's College London, London SE5 8AF, UK.
[141] Maastricht University Medical Centre, South Limburg Mental Health Research and Teaching Network, EURON, 6229 HX Maastricht, The Netherlands.
[142] Institute of Translational Medicine, University of Liverpool, Liverpool L69 3BX, UK.
[143] Max Planck Institute of Psychiatry, 80336 Munich, Germany.
[144] Munich Cluster for Systems Neurology (SyNergy), 80336 Munich, Germany.
[145] Department of Psychiatry and Psychotherapy, Jena University Hospital, 07743 Jena, Germany.
[146] Department of Psychiatry, Queensland Brain Institute and Queensland Centre for Mental Health Research, University of Queensland, Brisbane (St Lucia), Queensland, 4072, Australia.
[147] Department of Psychiatry and Behavioral Sciences, Johns Hopkins University School of Medicine, Baltimore, Maryland 21205, USA.
[148] Department of Psychiatry, Trinity College Dublin, Dublin 2, Ireland.
[149] Eli Lilly and Company, Lilly Corporate Center, Indianapolis, 46285 Indiana, USA.
[150] Department of Clinical Sciences, Psychiatry, Umeå University, SE-901 87 Umeå, Sweden.
[151] DETECT Early Intervention Service for Psychosis, Blackrock, Co. Dublin, Ireland.
[152] Centre for Public Health, Institute of Clinical Sciences, Queen's University Belfast, Belfast BT12 6AB, UK.
[153] Lawrence Berkeley National Laboratory, University of California at Berkeley, Berkeley, California 94720, USA.
[154] Institute of Psychiatry, King's College London, London SE5 8AF, UK.
[155] A list of authors and affiliations appear below.
[156] Melbourne Neuropsychiatry Centre, University of Melbourne & Melbourne Health, Melbourne, Victoria 3053, Australia.
[157] Department of Psychiatry, University of Helsinki, P.O. Box 590, FI-00029 HUS, Helsinki, Finland.
[158] Public Health Genomics Unit, National Institute for Health and Welfare, P.O. BOX 30, FI-00271 Helsinki, Finland
[159] Medical Faculty, University of Belgrade, 11000 Belgrade, Serbia.
[160] Department of Psychiatry, University of North Carolina, Chapel Hill, North Carolina 27599-7160, USA.
[161] Institute for Molecular Medicine Finland, FIMM, University of Helsinki, P.O. Box 20FI-00014, Helsinki, Finland
[162] Department of Epidemiology, Harvard School of Public Health, Boston, Massachusetts 02115, USA.
[163] Department of Psychiatry, University of Oxford, Oxford, OX3 7JX, UK.

[164] Virginia Institute for Psychiatric and Behavioral Genetics, Virginia Commonwealth University, Richmond, Virginia 23298, USA.
[165] Institute for Multiscale Biology, Icahn School of Medicine at Mount Sinai, New York, New York 10029, USA.
[166] PharmaTherapeutics Clinical Research, Pfizer Worldwide Research and Development, Cambridge, Massachusetts 02139, USA.
[167] Department of Psychiatry and Psychotherapy, University of Gottingen, 37073 Göttingen, Germany.
[168] Psychiatry and Psychotherapy Clinic, University of Erlangen, 91054 Erlangen, Germany.
[169] Hunter New England Health Service, Newcastle, New South Wales 2308, Australia.
[170] School of Biomedical Sciences, University of Newcastle, Newcastle, New South Wales 2308, Australia.
[171] Division of Cancer Epidemiology and Genetics, National Cancer Institute, Bethesda, Maryland 20892, USA.
[172] University of Iceland, Landspitali, National University Hospital, 101 Reykjavik, Iceland.
[173] Department of Psychiatry and Drug Addiction, Tbilisi State Medical University (TSMU), N33, 0177 Tbilisi, Georgia.
[174] Research and Development, Bronx Veterans Affairs Medical Center, New York, New York 10468, USA.
[175] Wellcome Trust Centre for Human Genetics, Oxford OX3 7BN, UK.
[176] deCODE Genetics, 101 Reykjavik, Iceland.
[177] Department of Clinical Neurology, Medical University of Vienna, 1090 Wien, Austria.
[178] Lieber Institute for Brain Development, Baltimore, Maryland 21205, USA.
[179] Department of Medical Genetics, University Medical Centre Utrecht, Universiteitsweg 100, 3584 CG, Utrecht, The Netherlands.
[180] Berkshire Healthcare NHS Foundation Trust, Bracknell RG12 1BQ, UK.
[181] Section of Psychiatry, University of Verona, 37134 Verona, Italy.
[182] Department of Psychiatry, University of Oulu, P.O. Box 5000, 90014, Finland.
[183] University Hospital of Oulu, P.O. Box 20, 90029 OYS, Finland.
[184] Molecular and Cellular Therapeutics, Royal College of Surgeons in Ireland, Dublin 2, Ireland.
[185] Health Research Board, Dublin 2, Ireland.
[186] School of Psychiatry and Clinical Neurosciences, The University of Western Australia, Perth Western Australia 6009, Australia.
[187] Computational Sciences CoE, Pfizer Worldwide Research and Development, Cambridge, Massachusetts 02139, USA.
[188] Human Genetics, Genome Institute of Singapore, A*STAR, Singapore 138672.
[189] A list of authors and affiliations appear below.
[190] University College London, London WC1E 6BT, UK.
[191] Department of Neuroscience, Icahn School of Medicine at Mount Sinai, New York, New York 10029, USA.
[192] Institute of Neuroscience and Medicine (INM-1), Research Center Jülich, 52428 Jülich, Germany.
[193] Department of Genetics, The Hebrew University of Jerusalem, 91905 Jerusalem, Israel.
[194] Neuroscience Discovery and Translational Area, Pharma Research and Early Development, F. Hoffman-La Roche, CH-4070 Basel, Switzerland.

[195] Centre for Clinical Research in Neuropsychiatry, School of Psychiatry and Clinical Neurosciences, The University of Western Australia, Medical Research Foundation Building, Perth Western Australia 6000, Australia.
[196] Virginia Institute for Psychiatric and Behavioral Genetics, Departments of Psychiatry and Human and Molecular Genetics, Virginia Commonwealth University, Richmond, Virginia 23298, USA.
[197] The Feinstein Institute for Medical Research, Manhasset, New York 11030, USA.
[198] The Zucker School of Medicine at Hofstra/Northwell, Hempstead, New York 11549, USA.
[199] The Zucker Hillside Hospital, Glen Oaks, New York 11004, USA.
[200] Saw Swee Hock School of Public Health, National University of Singapore, Singapore 117597, Singapore.
[201] Queensland Centre for Mental Health Research, University of Queensland, Brisbane 4076, Queensland, Australia.
[202] Center for Human Genetic Research and Department of Psychiatry, Massachusetts General Hospital, Boston, Massachusetts 02114, USA.
[203] Department of Child and Adolescent Psychiatry, Erasmus University Medical Centre, Rotterdam 3000, The Netherlands.
[204] Department of Complex Trait Genetics, Neuroscience Campus Amsterdam, VU University Medical Center Amsterdam, Amsterdam 1081, The Netherlands.
[205] Department of Functional Genomics, Center for Neurogenomics and Cognitive Research, Neuroscience Campus Amsterdam, VU University, Amsterdam 1081, The Netherlands.
[206] University of Aberdeen, Institute of Medical Sciences, Aberdeen AB25 2ZD, UK.
[207] Departments of Psychiatry, Neurology, Neuroscience and Institute of Genetic Medicine, Johns Hopkins School of Medicine, Baltimore, Maryland 21205, USA.
[208] Department of Clinical Medicine, University of Copenhagen, Copenhagen 2200, Denmark.
[209] Queensland Centre for Mental Health Research, The Park - Centre for Mental Health, Wacol, Queensland, Australia 4076
[210] Schizophrenia Research Institute, Sydney, New South Wales 2010, Australia.
[211] School of Biomedical Sciences and Pharmacy, University of Newcastle, Callaghan, New South Wales 2308, Australia.
[212] Priority Centre for Translational Neuroscience and Mental Health, University of Newcastle, Newcastle, New South Wales 2300, Australia.
[211] School of Biomedical Sciences and Pharmacy, University of Newcastle, Callaghan, New South Wales 2308, Australia.
[210] Schizophrenia Research Institute, Sydney, New South Wales 2010, Australia.
[212] Priority Centre for Translational Neuroscience and Mental Health, University of Newcastle, Newcastle, New South Wales 2300, Australia.
[211] School of Biomedical Sciences and Pharmacy, University of Newcastle, Callaghan, New South Wales 2308, Australia.
[210] Schizophrenia Research Institute, Sydney, New South Wales 2010, Australia.
[212] Priority Centre for Translational Neuroscience and Mental Health, University of Newcastle, Newcastle, New South Wales 2300, Australia.
[217] Department of Statistics, University of Oxford, Oxford, UK.
[218] Wellcome Trust Sanger Institute, Wellcome Trust Genome Campus, Hinxton, Cambridge, UK.

[219] Cambridge Institute for Medical Research, University of Cambridge School of Clinical Medicine, Cambridge, UK.
[220] Telethon Institute for Child Health Research, Centre for Child Health Research, University of Western Australia, Subiaco, Western Australia, Australia.
[221] Diamantina Institute of Cancer, Immunology and Metabolic Medicine, Princess Alexandra Hospital, University of Queensland, Brisbane, Queensland, Australia.
[222] Department of Epidemiology and Population Health, London School of Hygiene and Tropical Medicine, London, UK.
[223] Department of Epidemiology and Public Health, University College London, London, UK.
[224] Molecular and Physiological Sciences, The Wellcome Trust, London, UK.
[225] Peninsula School of Medicine and Dentistry, Plymouth University, Plymouth, UK.
[226] Clinical Neurosciences, St George's University of London, London, UK.
[227] Department of Medical and Molecular Genetics, School of Medicine, King's College London, Guy's Hospital, London, UK.
[228] Biomedical Research Centre, Ninewells Hospital and Medical School, Dundee, UK.
[229] Department of Clinical Neurosciences, University of Cambridge, Addenbrooke's Hospital, Cambridge, UK.
[230] Institute of Ophthalmology, University College London, London, UK.
[231] National Institute for Health Research, Biomedical Research Centre at Moorfields Eye Hospital, National Health Service Foundation Trust, London, UK.
[232] Department of Molecular Neuroscience, Institute of Neurology, London, UK.
[233] Oxford Centre for Diabetes, Endocrinology and Metabolism, Churchill Hospital, Oxford, UK.
[234] Fundació de Docència i Recerca Mútua de Terrassa, Universitat de Barcelona, Spain.
[235] Child and Adolescent Psychiatry, University of Technology Dresden, Dresden, Germany.
[236] Section for Experimental Psychopathology, General Psychiatry, Heidelberg, Germany.
[237] Institute of Cognitive Neuroscience, University College London, London, UK.
[238] Mental Health Sciences Unit, University College London, London, UK.
[239] Centro Investigación Biomédica en Red Salud Mental, Madrid, Spain.
[240] University Hospital Marqués de Valdecilla, Instituto de Formación e Investigación Marqués de Valdecilla, University of Cantabria, Santander, Spain.
[241] Centre for Clinical Research in Neuropsychiatry, The University of Western Australia, Perth, Western Australia, Australia.
[242] Western Australian Institute for Medical Research, The University of Western Australia, Perth, Western Australia, Australia.
[243] Department of Psychiatry, Academic Medical Center, University of Amsterdam, Amsterdam, The Netherlands.
[244] Department of Psychiatry, University Medical Center Groningen, University of Groningen, The Netherlands.

**Bipolar Disorder Working Group of the Psychiatric Genomics Consortium**

Eli A Stahl[1,2,3,] Gerome Breen[4,5,] Andreas J Forstner[6,7,8,9,10,] Andrew McQuillin[11,] Stephan Ripke[12,13,14,] Vassily Trubetskoy[13,] Manuel Mattheisen[15,16,17,18,19,] Yunpeng Wang[20,21,] Jonathan R I Coleman[4,5,] Héléna A Gaspar[4,5,] Christiaan A de Leeuw[22,] Stacy Steinberg[23,] Jennifer M Whitehead Pavlides[24,] Maciej Trzaskowski[25,] Enda M Byrne[25,] Tune H Pers[3,26,] Peter A Holmans[27,] Alexander L Richards[27,] Liam Abbott[12,] Esben Agerbo[19,28,29,] Huda Akil[30,] Diego Albani[31,] Ney Alliey-Rodriguez[32,] Thomas D Als[15,16,19,] Adebayo Anjorin[33,] Verneri Antilla[14,]

Swapnil Awasthi[13,] Judith A Badner[34,] Marie Bækvad-Hansen[19,35,] Jack D Barchas[36,] Nicholas Bass[11,] Michael Bauer[37,] Richard Belliveau[12,] Sarah E Bergen[38,] Carsten Bøcker Pedersen[19,28,29,] Erlend Bøen[39,] Marco P. Boks[40,] James Boocock[41,] Monika Budde[42,] William Bunney[43,] Margit Burmeister[44,] Jonas Bybjerg-Grauholm[19,35,] William Byerley[45,] Miquel Casas[46,47,48,49,] Felecia Cerrato[12,] Pablo Cervantes[50,] Kimberly Chambert[12,] Alexander W Charney[2,] Danfeng Chen[12,] Claire Churchhouse[12,14,] Toni-Kim Clarke[51,] William Coryell[52,] David W Craig[53,] Cristiana Cruceanu[50,54,] David Curtis[55,56,] Piotr M Czerski[57,] Anders M Dale[58,59,60,61,] Simone de Jong[4,5,] Franziska Degenhardt[8,] Jurgen Del-Favero[62,] J Raymond DePaulo[63,] Srdjan Djurovic[64,65,] Amanda L Dobbyn[1,2,] Ashley Dumont[12,] Torbjørn Elvsåshagen[66,67,] Valentina Escott-Price[27,] Chun Chieh Fan[61,] Sascha B Fischer[6,10,] Matthew Flickinger[68,] Tatiana M Foroud[69,] Liz Forty[27,] Josef Frank[70,] Christine Fraser[27,] Nelson B Freimer[71,] Katrin Gade[42,75,] Diane Gage[12,] Julie Garnham[76,] Claudia Giambartolomei[206,] Marianne Giørtz Pedersen[19,28,29,] Jaqueline Goldstein[12,] Scott D Gordon[77,] Katherine Gordon-Smith[78,] Elaine K Green[79,] Melissa J Green[80,133,] Tiffany A Greenwood[60,] Jakob Grove[15,16,19,81,] Weihua Guan[82,] José Guzman-Parra[83,] Marian L Hamshere[27,] Martin Hautzinger[84,] Urs Heilbronner[42,] Stefan Herms[6,8,10,] Maria Hipolito[85,] Per Hoffmann[6,8,10,] Dominic Holland[58,86,] Laura Huckins[1,2,] Stéphane Jamain[87,88,] Jessica S Johnson[1,2,] Radhika Kandaswamy[4,] Robert Karlsson[38,] James L Kennedy[89,90,91,92,] Sarah Kittel-Schneider[93,] James A Knowles[94,95,] Manolis Kogevinas[96,] Anna C Koller[8,] Ralph Kupka[97,98,99,] Catharina Lavebratt[72,] Jacob Lawrence[100,] William B Lawson[85,] Markus Leber[101,] Phil H Lee[12,14,102,] Shawn E Levy[103,] Jun Z Li[104,] Chunyu Liu[105,] Susanne Lucae[106,] Anna Maaser[8,] Donald J MacIntyre[107,108,] Pamela B Mahon[63,109,] Wolfgang Maier[110,] Lina Martinsson[73,] Steve McCarroll[12,111,] Peter McGuffin[4,] Melvin G McInnis[112,] James D McKay[113,] Helena Medeiros[95,] Sarah E Medland[77,] Fan Meng[30,112,] Lili Milani[114,] Grant W Montgomery[25,] Derek W Morris[115,116,] Thomas W Mühleisen[6,117,] Niamh Mullins[4,] Hoang Nguyen[1,2,] Caroline M Nievergelt[60,118,] Annelie Nordin Adolfsson[119,] Evaristus A Nwulia[85,] Claire O'Donovan[76,] Loes M Olde Loohuis[71,] Anil P S Ori[71,] Lilijana Oruc[120,] Urban Ösby[121,] Roy H Perlis[122,123,] Amy Perry[78,] Andrea Pfennig[37,] James B Potash[63,] Shaun M Purcell[2,109,] Eline J Regeer[124,] Andreas Reif[93,] Céline S Reinbold[6,10,] John P Rice[125,] Fabio Rivas[83,] Margarita Rivera[4,126,] Panos Roussos[1,2,127,] Douglas M Ruderfer[128,] Euijung Ryu[129,] Cristina Sánchez-Mora[46,47,49,] Alan F Schatzberg[130,] William A Scheftner[131,] Nicholas J Schork[132,] Cynthia Shannon Weickert[80,133,] Tatyana Shehktman[60,] Paul D Shilling[60,] Engilbert Sigurdsson[134,] Claire Slaney[76,] Olav B Smeland[135,136,] Janet L Sobell[137,] Christine Søholm Hansen[19,35,] Anne T Spijker[138,] David St Clair[139,] Michael Steffens[140,] John S Strauss[91,141,] Fabian Streit[70,] Jana Strohmaier[70,] Szabolcs Szelinger[142,] Robert C Thompson[112,] Thorgeir E Thorgeirsson[23,] Jens Treutlein[70,] Helmut Vedder[143,] Weiqing Wang[1,2,] Stanley J Watson[112,] Thomas W Weickert[80,133,] Stephanie H Witt[70,] Simon Xi[144,] Wei Xu[145,146,] Allan H Young[147,] Peter Zandi[148,] Peng Zhang[149,] Sebastian Zöllner[112,] eQTLGen Consortium, BIOS Consortium, Rolf Adolfsson[119,] Ingrid Agartz[17,39,150,] Martin Alda[76,151,] Lena Backlund[73,] Bernhard T Baune[152,158,] Frank Bellivier[153,154,155,156,] Wade H Berrettini[157,] Joanna M Biernacka[129,] Douglas H R Blackwood[51,] Michael Boehnke[68,] Anders D Børglum[15,16,19,] Aiden Corvin[116,] Nicholas Craddock[27,] Mark J Daly[12,14,] Udo Dannlowski[158,] Tõnu Esko[3,111,114,159,] Bruno Etain[153,155,156,160,] Mark Frye[161,] Janice M Fullerton[133,162,] Elliot S Gershon[32,163,] Michael Gill[116,] Fernando Goes[63,] Maria Grigoroiu-Serbanescu[164,] Joanna Hauser[57,] David M Hougaard[19,35,] Christina M Hultman[38,] Ian Jones[27,] Lisa A Jones[78,] René S Kahn[2,40,] George Kirov[27,] Mikael Landén[38,165,] Marion Leboyer[88,153,166,] Cathryn M Lewis[4,5,167,] Qingqin S Li[168,] Jolanta Lissowska[169,] Nicholas G Martin[77,170,] Fermin Mayoral[83,] Susan L McElroy[171,] Andrew M McIntosh[51,172,] Francis J McMahon[173,] Ingrid Melle[174,175,] Andres Metspalu[114,176,] Philip B Mitchell[80,] Gunnar Morken[177,178,] Ole Mors[19,179,] Preben Bo

Mortensen[15,19,28,29], Bertram Müller-Myhsok[54,180,181], Richard M Myers[103], Benjamin M Neale[3,12,14], Vishwajit Nimgaonkar[182], Merete Nordentoft[19,183], Markus M Nöthen[8], Michael C O'Donovan[27], Ketil J Oedegaard[184,185], Michael J Owen[27], Sara A Paciga[186], Carlos Pato[95,187], Michele T Pato[95], Danielle Posthuma[22,188], Josep Antoni Ramos-Quiroga[46,47,48,49], Marta Ribasés[46,47,49], Marcella Rietschel[70], Guy A Rouleau[189,190], Martin Schalling[72], Peter R Schofield[133,162], Thomas G Schulze[42,63,70,75,173], Alessandro Serretti[191], Jordan W Smoller[12,192,193], Hreinn Stefansson[23], Kari Stefansson[23,194], Eystein Stordal[195,196], Patrick F Sullivan[38,197,198], Gustavo Turecki[199], Arne E Vaaler[200], Eduard Vieta[201], John B Vincent[141], Thomas Werge[19,202,203], John I Nurnberger[204], Naomi R Wray[24,25], Arianna Di Florio[27,198], Howard J Edenberg[205], Sven Cichon[6,8,10,117], Roel A Ophoff[40,41,71], Laura J Scott[68], Ole A Andreassen[135,136], John Kelsoe[60], Pamela Sklar[1,2,†]

† deceased

**Affiliations:**

[1] Department of Genetics and Genomic Sciences, Icahn School of Medicine at Mount Sinai, New York, New York, USA

[2] Department of Psychiatry, Icahn School of Medicine at Mount Sinai, New York, New York, USA

[3] Medical and Population Genetics, Broad Institute, Cambridge, Massachusetts, USA

[4] Social, Genetic and Developmental Psychiatry Centre, King's College London, London, UK

[5] NIHR BRC for Mental Health, King's College London, London, UK

[6] Department of Biomedicine, University of Basel, Basel, Switzerland

[7] Department of Psychiatry (UPK), University of Basel, Basel, Switzerland

[8] Institute of Human Genetics, University of Bonn, School of Medicine & University Hospital Bonn, Bonn, Germany

[9] Centre for Human Genetics, University of Marburg, Marburg, Germany

[10] Institute of Medical Genetics and Pathology, University Hospital Basel, Basel, Switzerland

[11] Molecular Psychiatry Laboratory, Division of Psychiatry, University College London, London WC1E6JJ, UK

[12] Stanley Center for Psychiatric Research, Broad Institute, Cambridge, Massachusetts, USA

[13] Department of Psychiatry and Psychotherapy, Charité - Universitätsmedizin, Berlin, Germany

[14] Analytic and Translational Genetics Unit, Massachusetts General Hospital, Boston, Massachusetts, USA

[15] iSEQ, Center for Integrative Sequencing, Aarhus University, Aarhus, Denmark

[16] Department of Biomedicine - Human Genetics, Aarhus University, Aarhus, Denmark

[17] Department of Clinical Neuroscience, Centre for Psychiatry Research, Karolinska Institutet, Stockholm, Sweden

[18] Department of Psychiatry, Psychosomatics and Psychotherapy, Center of Mental Health, University Hospital Würzburg, Würzburg, Germany

[19] iPSYCH, The Lundbeck Foundation Initiative for Integrative Psychiatric Research, Denmark

[20] Institute of Biological Psychiatry, Mental Health Centre Sct. Hans, Copenhagen, Denmark

[21] Institute of Clinical Medicine, University of Oslo, Oslo, Norway

[22] Department of Complex Trait Genetics, Center for Neurogenomics and Cognitive Research, Amsterdam Neuroscience, Vrije Universiteit Amsterdam, Amsterdam, Netherlands

[23] deCODE Genetics / Amgen, Reykjavik, Iceland

[24] Queensland Brain Institute, The University of Queensland, Brisbane, Queensland, Australia

[25] Institute for Molecular Bioscience, The University of Queensland, Brisbane, Queensland, Australia
[26] Division of Endocrinology and Center for Basic and Translational Obesity Research, Boston Children's Hospital, Boston, Massachusetts, USA
[27] MRC Centre for Neuropsychiatric Genetics and Genomics, Institute of Psychological Medicine and Clinical Neurosciences, School of Medicine, Cardiff University, Cardiff CF24 4HQ, UK
[28] National Centre for Register-Based Research, Aarhus University, Aarhus, Denmark
[29] Centre for Integrated Register-based Research, Aarhus University, Aarhus, Denmark
[30] Molecular & Behavioral Neuroscience Institute, University of Michigan, Ann Arbor, MI, USA
[31] Department of Neuroscience, IRCCS - Istituto Di Ricerche Farmacologiche Mario Negri, Milan, Italy
[32] Department of Psychiatry and Behavioral Neuroscience, University of Chicago, Chicago, Illinois, USA
[33] Psychiatry, Berkshire Healthcare NHS Foundation Trust, Bracknell, UK
[34] Psychiatry, Rush University Medical Center, Chicago, Illinois, USA
[35] Center for Neonatal Screening, Department for Congenital Disorders, Statens Serum Institut, Copenhagen, Denmark
[36] Department of Psychiatry, Weill Cornell Medical College, New York, New York, USA
[37] Department of Psychiatry and Psychotherapy, University Hospital Carl Gustav Carus, Technische Universität Dresden, Dresden, Germany
[38] Department of Medical Epidemiology and Biostatistics, Karolinska Institutet, Stockholm, Sweden
[39] Department of Psychiatric Research, Diakonhjemmet Hospital, Oslo, Norway
[40] Psychiatry, UMC Utrecht Brain Center Rudolf Magnus, Utrecht, Netherlands
[41] Human Genetics, University of California Los Angeles, Los Angeles, California, USA
[42] Institute of Psychiatric Phenomics and Genomics (IPPG), University Hospital, LMU Munich, Munich, Germany
[43] Department of Psychiatry and Human Behavior, University of California, Irvine, Irvine, California, USA
[44] Molecular & Behavioral Neuroscience Institute and Department of Computational Medicine & Bioinformatics, University of Michigan, Ann Arbor, MI, USA
[45] Psychiatry, University of California San Francisco, San Francisco, California, USA
[46] Instituto de Salud Carlos III, Biomedical Network Research Centre on Mental Health (CIBERSAM), Madrid, Spain
[47] Department of Psychiatry, Hospital Universitari Vall d´Hebron, Barcelona, Spain
[48] Department of Psychiatry and Forensic Medicine, Universitat Autònoma de Barcelona, Barcelona, Spain
[49] Psychiatric Genetics Unit, Group of Psychiatry Mental Health and Addictions, Vall d´Hebron Research Institut (VHIR), Universitat Autònoma de Barcelona, Barcelona, Spain
[50] Department of Psychiatry, Mood Disorders Program, McGill University Health Center, Montreal, Quebec, Canada
[51] Division of Psychiatry, University of Edinburgh, Edinburgh, UK
[52] University of Iowa Hospitals and Clinics, Iowa City, Iowa, USA
[53] Translational Genomics, USC, Phoenix, Arizona, USA

[54] Department of Translational Research in Psychiatry, Max Planck Institute of Psychiatry, Munich, Germany
[55] Centre for Psychiatry, Queen Mary University of London, London, UK
[56] UCL Genetics Institute, University College London, London, UK
[57] Department of Psychiatry, Laboratory of Psychiatric Genetics, Poznan University of Medical Sciences, Poznan, Poland
[58] Department of Neurosciences, University of California San Diego, La Jolla, California, USA
[59] Department of Radiology, University of California San Diego, La Jolla, California, USA
[60] Department of Psychiatry, University of California San Diego, La Jolla, California, USA
[61] Department of Cognitive Science, University of California San Diego, La Jolla, California, USA
[62] Applied Molecular Genomics Unit, VIB Department of Molecular Genetics, University of Antwerp, Antwerp, Belgium
[63] Department of Psychiatry and Behavioral Sciences, Johns Hopkins University School of Medicine, Baltimore, MD, USA
[64] Department of Medical Genetics, Oslo University Hospital Ullevål, Oslo, Norway
[65] NORMENT, KG Jebsen Centre for Psychosis Research, Department of Clinical Science, University of Bergen, Bergen, Norway
[66] Department of Neurology, Oslo University Hospital, Oslo, Norway
[67] NORMENT, KG Jebsen Centre for Psychosis Research, Oslo University Hospital, Oslo, Norway
[68] Center for Statistical Genetics and Department of Biostatistics, University of Michigan, Ann Arbor, MI, USA
[69] Department of Medical & Molecular Genetics, Indiana University, Indianapolis, Indiana, USA
[70] Department of Genetic Epidemiology in Psychiatry, Central Institute of Mental Health, Medical Faculty Mannheim, Heidelberg University, Mannheim, Germany
[71] Center for Neurobehavioral Genetics, University of California Los Angeles, Los Angeles, California, USA
[72] Department of Molecular Medicine and Surgery, Karolinska Institutet and Center for Molecular Medicine, Karolinska University Hospital, Stockholm, Sweden
[75] Department of Psychiatry and Psychotherapy, University Medical Center Göttingen, Göttingen, Germany
[76] Department of Psychiatry, Dalhousie University, Halifax, Nova Scotia, Canada
[77] Genetics and Computational Biology, QIMR Berghofer Medical Research Institute, Brisbane, Queensland, Australia
[78] Department of Psychological Medicine, University of Worcester, Worcester, UK
[79] School of Biomedical Sciences, Plymouth University Peninsula Schools of Medicine and Dentistry, University of Plymouth, Plymouth, UK
[80] School of Psychiatry, University of New South Wales, Sydney, New South Wales, Australia
[81] Bioinformatics Research Centre, Aarhus University, Aarhus, Denmark
[82] Biostatistics, University of Minnesota System, Minneapolis, Minnesota, USA
[83] Mental Health Department, University Regional Hospital, Biomedicine Institute (IBIMA), Málaga, Spain
[84] Department of Psychology, Eberhard Karls Universität Tübingen, Tubingen, Germany
[85] Department of Psychiatry and Behavioral Sciences, Howard University Hospital, Washington, DC, USA

[86] Center for Multimodal Imaging and Genetics, University of California San Diego, La Jolla, California, USA
[87] Psychiatrie Translationnelle, Inserm U955, Créteil, France
[88] Faculté de Médecine, Université Paris Est, Créteil, France
[89] Campbell Family Mental Health Research Institute, Centre for Addiction and Mental Health, Toronto, Ontario, Canada
[90] Neurogenetics Section, Centre for Addiction and Mental Health, Toronto, Ontario, Canada
[91] Department of Psychiatry, University of Toronto, Toronto, Ontario, Canada
[92] Institute of Medical Sciences, University of Toronto, Toronto, Ontario, Canada
[93] Department of Psychiatry, Psychosomatic Medicine and Psychotherapy, University Hospital Frankfurt, Frankfurt am Main, Germany
[94] Cell Biology, SUNY Downstate Medical Center College of Medicine, Brooklyn, New York, USA
[95] Institute for Genomic Health, SUNY Downstate Medical Center College of Medicine, Brooklyn, New York, USA
[96] ISGlobal, Barcelona, Spain
[97] Psychiatry, Altrecht, Utrecht, Netherlands
[98] Psychiatry, GGZ inGeest, Amsterdam, Netherlands
[99] Psychiatry, VU medisch centrum, Amsterdam, Netherlands
[100] Psychiatry, North East London NHS Foundation Trust, Ilford, UK
[101] Clinic for Psychiatry and Psychotherapy, University Hospital Cologne, Cologne, Germany
[102] Psychiatric and Neurodevelopmental Genetics Unit, Massachusetts General Hospital, Boston, Massachusetts, USA
[103] Hudson Alpha Institute for Biotechnology, Huntsville, Alabama, USA
[104] Department of Human Genetics, University of Michigan, Ann Arbor, Michigan, USA
[105] Psychiatry, University of Illinois at Chicago College of Medicine, Chicago, Illinois, USA
[106] Max Planck Institute of Psychiatry, Munich, Germany
[107] Mental Health, NHS 24, Glasgow, UK
[108] Division of Psychiatry, Centre for Clinical Brain Sciences, University of Edinburgh, Edinburgh, UK
[109] Psychiatry, Brigham and Women's Hospital, Boston, Massachusetts, USA
[110] Department of Psychiatry and Psychotherapy, University of Bonn, Bonn, Germany
[111] Department of Genetics, Harvard Medical School, Boston, Massachusetts, USA
[112] Department of Psychiatry, University of Michigan, Ann Arbor, Michigan, USA
[113] Genetic Cancer Susceptibility Group, International Agency for Research on Cancer, Lyon, France
[114] Estonian Genome Center, University of Tartu, Tartu, Estonia
[115] Discipline of Biochemistry, Neuroimaging and Cognitive Genomics (NICOG) Centre, National University of Ireland, Galway, Galway, Ireland
[116] Neuropsychiatric Genetics Research Group, Dept of Psychiatry and Trinity Translational Medicine Institute, Trinity College Dublin, Dublin, Ireland
[117] Institute of Neuroscience and Medicine (INM-1), Research Centre Jülich, Jülich, Germany
[118] Research/Psychiatry, Veterans Affairs San Diego Healthcare System, San Diego, California, USA
[119] Department of Clinical Sciences, Psychiatry, Umeå University Medical Faculty, Umeå, Sweden

[120] Department of Clinical Psychiatry, Psychiatry Clinic, Clinical Center University of Sarajevo, Sarajevo, Bosnia and Herzegovina
[121] Department of Neurobiology, Care sciences, and Society, Karolinska Institutet and Center for Molecular Medicine, Karolinska University Hospital, Stockholm, Sweden
[122] Psychiatry, Harvard Medical School, Boston, Massachusetts, USA
[123] Division of Clinical Research, Massachusetts General Hospital, Boston, Massachusetts, USA
[124] Outpatient Clinic for Bipolar Disorder, Altrecht, Utrecht, Netherlands
[125] Department of Psychiatry, Washington University in Saint Louis, Saint Louis, Missouri, USA
[126] Department of Biochemistry and Molecular Biology II, Institute of Neurosciences, Center for Biomedical Research, University of Granada, Granada, Spain
[127] Department of Neuroscience, Icahn School of Medicine at Mount Sinai, New York, New York, USA
[128] Medicine, Psychiatry, Biomedical Informatics, Vanderbilt University Medical Center, Nashville, Tennessee, USA
[129] Department of Health Sciences Research, Mayo Clinic, Rochester, Minnesota, USA
[130] Psychiatry and Behavioral Sciences, Stanford University School of Medicine, Stanford, California, USA
[131] Rush University Medical Center, Chicago, Illinois, USA
[132] Scripps Translational Science Institute, La Jolla, California, USA
[133] Neuroscience Research Australia, Sydney, New South Wales, Australia
[134] Faculty of Medicine, Department of Psychiatry, School of Health Sciences, University of Iceland, Reykjavik, Iceland
[135] Division of Mental Health and Addiction, Oslo University Hospital, Oslo, Norway
[136] NORMENT, University of Oslo, Oslo, Norway
[137] Psychiatry and the Behavioral Sciences, University of Southern California, Los Angeles, California, USA
[138] Mood Disorders, PsyQ, Rotterdam, Netherlands
[139] Institute for Medical Sciences, University of Aberdeen, Aberdeen, UK
[140] Research Division, Federal Institute for Drugs and Medical Devices (BfArM), Bonn, Germany
[141] Centre for Addiction and Mental Health, Toronto, Ontario, Canada
[142] Neurogenomics, TGen, Los Angeles, Arizona, USA
[143] Psychiatry, Psychiatrisches Zentrum Nordbaden, Wiesloch, Germany
[144] Computational Sciences Center of Emphasis, Pfizer Global Research and Development, Cambridge, Massachusetts, USA
[145] Department of Biostatistics, Princess Margaret Cancer Centre, Toronto, Ontario, Canada
[146] Dalla Lana School of Public Health, University of Toronto, Toronto, Ontario, Canada
[147] Psychological Medicine, Institute of Psychiatry, Psychology & Neuroscience, King's College London, London, UK
[148] Department of Mental Health, Johns Hopkins University Bloomberg School of Public Health, Baltimore, Maryland, USA
[149] Institute of Genetic Medicine, Johns Hopkins University School of Medicine, Baltimore, Maryland, USA
[150] NORMENT, KG Jebsen Centre for Psychosis Research, Division of Mental Health and Addiction, Institute of Clinical Medicine and Diakonhjemmet Hospital, University of Oslo, Oslo, Norway

[151] National Institute of Mental Health, Klecany, Czech Republic
[152] Department of Psychiatry, University of Melbourne, Melbourne, Victoria, Australia
[153] Department of Psychiatry and Addiction Medicine, Assistance Publique - Hôpitaux de Paris, Paris, France
[154] Paris Bipolar and TRD Expert Centres, FondaMental Foundation, Paris, France
[155] UMR-S1144 Team 1: Biomarkers of relapse and therapeutic response in addiction and mood disorders, INSERM, Paris, France
[156] Psychiatry, Université Paris Diderot, Paris, France
[157] Psychiatry, University of Pennsylvania, Philadelphia, Pennsylvania, USA
[158] Department of Psychiatry, University of Münster, Münster, Germany
[159] Division of Endocrinology, Children's Hospital Boston, Boston, Massachusetts, USA
[160] Centre for Affective Disorders, Institute of Psychiatry, Psychology and Neuroscience, London, UK
[161] Department of Psychiatry & Psychology, Mayo Clinic, Rochester, Minnesota, USA
[162] School of Medical Sciences, University of New South Wales, Sydney, New South Wales, Australia
[163] Department of Human Genetics, University of Chicago, Chicago, Illinois, USA
[164] Biometric Psychiatric Genetics Research Unit, Alexandru Obregia Clinical Psychiatric Hospital, Bucharest, Romania
[165] Institute of Neuroscience and Physiology, University of Gothenburg, Gothenburg, Sweden
[166] INSERM, Paris, France
[167] Department of Medical & Molecular Genetics, King's College London, London, UK
[168] Neuroscience Therapeutic Area, Janssen Research and Development, LLC, Titusville, New Jersey, USA
[169] Cancer Epidemiology and Prevention, M. Sklodowska-Curie Cancer Center and Institute of Oncology, Warsaw, Poland
[170] School of Psychology, The University of Queensland, Brisbane, Queensland, Australia
[171] Research Institute, Lindner Center of HOPE, Mason, Ohio, USA
[172] Centre for Cognitive Ageing and Cognitive Epidemiology, University of Edinburgh, Edinburgh, UK
[173] Human Genetics Branch, Intramural Research Program, National Institute of Mental Health, Bethesda, Maryland, USA
[174] Division of Mental Health and Addiction, Oslo University Hospital, Oslo, Norway
[175] Division of Mental Health and Addiction, University of Oslo, Institute of Clinical Medicine, Oslo, Norway
[176] Institute of Molecular and Cell Biology, University of Tartu, Tartu, Estonia
[177] Mental Health, Faculty of Medicine and Health Sciences, Norwegian University of Science and Technology - NTNU, Trondheim, Norway
[178] Psychiatry, St Olavs University Hospital, Trondheim, Norway
[179] Psychosis Research Unit, Aarhus University Hospital, Risskov, Denmark
[180] Munich Cluster for Systems Neurology (SyNergy), Munich, Germany
[181] University of Liverpool, Liverpool, UK
[182] Psychiatry and Human Genetics, University of Pittsburgh, Pittsburgh, Pennsylvania, USA
[183] Mental Health Services in the Capital Region of Denmark, Mental Health Center Copenhagen, University of Copenhagen, Copenhagen, Denmark
[184] Division of Psychiatry, Haukeland Universitetssjukehus, Bergen, Norway

[185] Faculty of Medicine and Dentistry, University of Bergen, Bergen, Norway
[186] Human Genetics and Computational Biomedicine, Pfizer Global Research and Development, Groton, Connecticut, USA
[187] College of Medicine Institute for Genomic Health, SUNY Downstate Medical Center College of Medicine, Brooklyn, New York, USA
[188] Department of Clinical Genetics, Amsterdam Neuroscience, Vrije Universiteit Medical Center, Amsterdam, Netherlands
[189] Department of Neurology and Neurosurgery, McGill University, Faculty of Medicine, Montreal, Quebec, Canada
[190] Montreal Neurological Institute and Hospital, Montreal, Quebec, Canada
[191] Department of Biomedical and NeuroMotor Sciences, University of Bologna, Bologna, Italy
[192] Department of Psychiatry, Massachusetts General Hospital, Boston, Massachusetts, USA
[193] Psychiatric and Neurodevelopmental Genetics Unit (PNGU), Massachusetts General Hospital, Boston, Massachusetts, USA
[194] Faculty of Medicine, University of Iceland, Reykjavik, Iceland
[195] Department of Psychiatry, Hospital Namsos, Namsos, Norway
[196] Department of Neuroscience, Norges Teknisk Naturvitenskapelige Universitet Fakultet for naturvitenskap og teknologi, Trondheim, Norway
[197] Department of Genetics, University of North Carolina at Chapel Hill, Chapel Hill, North Carolina, USA
[198] Department of Psychiatry, University of North Carolina at Chapel Hill, Chapel Hill, North Carolina, USA
[199] Department of Psychiatry, McGill University, Montreal, Quebec, Canada
[200] Dept of Psychiatry, Sankt Olavs Hospital Universitetssykehuset i Trondheim, Trondheim, Norway
[201] Clinical Institute of Neuroscience, Hospital Clinic, University of Barcelona, IDIBAPS, CIBERSAM, Barcelona, Spain
[202] Institute of Biological Psychiatry, MHC Sct. Hans, Mental Health Services Copenhagen, Roskilde, Denmark
[203] Department of Clinical Medicine, University of Copenhagen, Copenhagen, Denmark
[204] Psychiatry, Indiana University School of Medicine, Indianapolis, Indiana, USA
[205] Biochemistry and Molecular Biology, Indiana University School of Medicine, Indianapolis, Indiana, USA
[206] Department of Pathology and Laboratory Medicine, University of California Los Angeles, Los Angeles, California, USA

**Major Depressive Disorder Working Group of the Psychiatric Genomics Consortium**

Naomi R Wray[1,2], Stephan Ripke[3,4,5], Manuel Mattheisen [6,7,8], Maciej Trzaskowski[1], Enda M Byrne[1], Abdel Abdellaoui[9], Mark J Adams[10], Esben Agerbo[11,12,13], Tracy M Air[14], Till F M Andlauer[15,16], Silviu-Alin Bacanu[17], Marie Bækvad-Hansen[13,18], Aartjan T F Beekman[19], Tim B Bigdeli[17,20], Elisabeth B Binder[15,21], Julien Bryois[22], Henriette N Buttenschøn[13,23,24], Jonas Bybjerg-Grauholm[13,18], Na Cai[25,26,] Enrique Castelao[27], Jane Hvarregaard Christensen[8,13,24], Toni-Kim Clarke[10], Jonathan R I Coleman[28], Lucía Colodro-Conde[29], Baptiste Couvy-Duchesne[2,30], Nick Craddock[31], Gregory E Crawford[32,33], Gail Davies[34], Ian J Deary[34], Franziska Degenhardt[35], Eske M Derks[29], Nese Direk[36,37], Conor V Dolan[9], Erin C Dunn[38,39,40], Thalia C Eley[28], Valentina Escott-Price[41], Farnush Farhadi Hassan Kiadeh[42], Hilary K Finucane[43,44], Jerome C Foo[45],

Andreas J Forstner[35,46,47,48], Josef Frank[45], Héléna A Gaspar[28], Michael Gill[49], Fernando S Goes[50], Scott D Gordon[29], Jakob Grove[8,13,24,51], Lynsey S Hall[10,52], Christine Søholm Hansen[13,18], Thomas F Hansen[53,54,55], Stefan Herms[35,47], Ian B Hickie[56], Per Hoffmann[35,47], Georg Homuth[57], Carsten Horn[58], Jouke-Jan Hottenga[9], David M Hougaard[13,18], David M Howard[10,28], Marcus Ising[59], Rick Jansen[19], Ian Jones[60], Lisa A Jones[61], Eric Jorgenson[62], James A Knowles[63], Isaac S Kohane[64,65,66], Julia Kraft[4], Warren W. Kretzschmar[67], Zoltán Kutalik[68,69], Yihan Li[67], Penelope A Lind[29], Donald J MacIntyre[70,71], Dean F MacKinnon[50], Robert M Maier[2], Wolfgang Maier[72], Jonathan Marchini[73], Hamdi Mbarek[9], Patrick McGrath[74], Peter McGuffin[28], Sarah E Medland[29], Divya Mehta[2,75], Christel M Middeldorp[9,76,77], Evelin Mihailov[78], Yuri Milaneschi[19], Lili Milani[78], Francis M Mondimore[50], Grant W Montgomery[1], Sara Mostafavi[79,80], Niamh Mullins[28], Matthias Nauck[81,82], Bernard Ng[80], Michel G Nivard[9], Dale R Nyholt[83], Paul F O'Reilly[28], Hogni Oskarsson[84], Michael J Owen[60], Jodie N Painter[29], Carsten Bøcker Pedersen[11,12,13], Marianne Giørtz Pedersen[11,12,13], Roseann E Peterson[17,85], Wouter J Peyrot[19], Giorgio Pistis[27], Danielle Posthuma[86,87], Jorge A Quiroz[88], Per Qvist[8,13,24], John P Rice[89], Brien P. Riley[17], Margarita Rivera[28,90], Saira Saeed Mirza[36], Robert Schoevers[91], Eva C Schulte[92,93], Ling Shen[62], Jianxin Shi[94], Stanley I Shyn[95], Engilbert Sigurdsson[96], Grant C B Sinnamon[97], Johannes H Smit[19], Daniel J Smith[98], Hreinn Stefansson[99], Stacy Steinberg[99], Fabian Streit[45], Jana Strohmaier[45], Katherine E Tansey[100], Henning Teismann[101], Alexander Teumer[102], Wesley Thompson[13,54,103,104], Pippa A Thomson[105], Thorgeir E Thorgeirsson[99], Matthew Traylor[106], Jens Treutlein[45], Vassily Trubetskoy[4], André G Uitterlinden[107], Daniel Umbricht[108], Sandra Van der Auwera[109], Albert M van Hemert[110], Alexander Viktorin[22], Peter M Visscher[1,2], Yunpeng Wang[13,54,104], Bradley T. Webb[111], Shantel Marie Weinsheimer[13,54], Jürgen Wellmann[101], Gonneke Willemsen[9], Stephanie H Witt[45], Yang Wu[1], Hualin S Xi[112], Jian Yang[2,113], Futao Zhang[1], Volker Arolt[114], Bernhard T Baune[114,115,116], Klaus Berger[101], Dorret I Boomsma[9], Sven Cichon[35,47,117,118], Udo Dannlowski[114], Eco JC de Geus[9,119], J Raymond DePaulo[50], Enrico Domenici[120], Katharina Domschke[121,122], Tõnu Esko[5,78], Hans J Grabe[109], Steven P Hamilton[123], Caroline Hayward[124], Andrew C Heath[89], Kenneth S Kendler[17], Stefan Kloiber[59,125,126], Glyn Lewis[127], Qingqin S Li[128], Susanne Lucae[59], Pamela AF Madden[89], Patrik K Magnusson[22], Nicholas G Martin[29], Andrew M McIntosh[10,34], Andres Metspalu[78,129], Ole Mors[13,130], Preben Bo Mortensen[11,12,13,24], Bertram Müller-Myhsok[15,131,132], Merete Nordentoft[13,133], Markus M Nöthen[35], Michael C O'Donovan[60], Sara A Paciga[134], Nancy L Pedersen[22], Brenda WJH Penninx[19], Roy H Perlis[38,135], David J Porteous[105], James B Potash[136], Martin Preisig[27], Marcella Rietschel[45], Catherine Schaefer[62], Thomas G Schulze[45, 93,137,138,139], Jordan W Smoller[38,39,40], Kari Stefansson[99,140], Henning Tiemeier[36,141,142], Rudolf Uher[143], Henry Völzke[102], Myrna M Weissman[74,144], Thomas Werge[13,54,145], Cathryn M Lewis[28,146], Douglas F Levinson[147], Gerome Breen[28,148], Anders D Børglum[8,13,24], Patrick F Sullivan[22,149,150].

**Affiliations:**

[1] Institute for Molecular Bioscience, The University of Queensland, Brisbane, Queensland, Australia

[2] Queensland Brain Institute, The University of Queensland, Brisbane, Queensland, Australia

[3] Analytic and Translational Genetics Unit, Massachusetts General Hospital, Boston, Massachusetts, USA

[4] Department of Psychiatry and Psychotherapy, Universitätsmedizin Berlin Campus Charité Mitte, Berlin, Germany

[5] Medical and Population Genetics, Broad Institute, Cambridge, Massachusetts, USA

[6] Department of Psychiatry, Psychosomatics and Psychotherapy, University of Wurzburg, Wurzburg, Germany
[7] Centre for Psychiatry Research, Department of Clinical Neuroscience, Karolinska Institutet, Stockholm, Sweden
[8] Department of Biomedicine, Aarhus University, Aarhus, Denmark
[9] Dept of Biological Psychology & EMGO+ Institute for Health and Care Research, Vrije Universiteit Amsterdam, Amsterdam, Netherlands
[10] Division of Psychiatry, University of Edinburgh, Edinburgh, UK
[11] Centre for Integrated Register-based Research, Aarhus University, Aarhus, Denmark
[12] National Centre for Register-Based Research, Aarhus University, Aarhus, Denmark
[13] iPSYCH, The Lundbeck Foundation Initiative for Integrative Psychiatric Research, Denmark
[14] Discipline of Psychiatry, University of Adelaide, Adelaide, South Australia, Australia
[15] Department of Translational Research in Psychiatry, Max Planck Institute of Psychiatry, Munich, Germany
[16] Department of Neurology, Klinikum rechts der Isar, Technical University of Munich, Munich, Germany
[17] Virginia Institute for Psychiatric and Behavioral Genetics, Departments of Psychiatry and Human and Molecular Genetics, Virginia Commonwealth University, Richmond, Virginia 23298, USA
[18] Center for Neonatal Screening, Department for Congenital Disorders, Statens Serum Institut, Copenhagen, Denmark
[19] Department of Psychiatry, Vrije Universiteit Medical Center and GGZ inGeest, Amsterdam, Netherlands
[20] Virginia Institute for Psychiatric and Behavior Genetics, Richmond, Virginia, USA
[21] Department of Psychiatry and Behavioral Sciences, Emory University School of Medicine, Atlanta, Georgia, USA
[22] Department of Medical Epidemiology and Biostatistics, Karolinska Institutet, Stockholm, Sweden
[23] Department of Clinical Medicine, Translational Neuropsychiatry Unit, Aarhus University, Aarhus, Denmark
[24] iSEQ, Centre for Integrative Sequencing, Aarhus University, Aarhus, Denmark
[25] Human Genetics, Wellcome Trust Sanger Institute, Cambridge, UK
[26] Statistical genomics and systems genetics, European Bioinformatics Institute (EMBL-EBI), Cambridge, UK
[27] Department of Psychiatry, Lausanne University Hospital and University of Lausanne, Lausanne, Switzerland
[28] Social, Genetic and Developmental Psychiatry Centre, King's College London, London, UK
[29] Genetics and Computational Biology, QIMR Berghofer Medical Research Institute, Brisbane, Queensland, Australia
[30] Centre for Advanced Imaging, The University of Queensland, Brisbane, Queensland, Australia
[31] Psychological Medicine, Cardiff University, Cardiff, UK
[32] Center for Genomic and Computational Biology, Duke University, Durham, North Carolina, USA
[33] Department of Pediatrics, Division of Medical Genetics, Duke University, Durham, North Carolina, USA

[34] Centre for Cognitive Ageing and Cognitive Epidemiology, University of Edinburgh, Edinburgh, UK
[35] Institute of Human Genetics, University of Bonn, School of Medicine & University Hospital Bonn, Bonn, Germany
[36] Epidemiology, Erasmus MC, Rotterdam, Zuid-Holland, Netherlands
[37] Psychiatry, Dokuz Eylul University School Of Medicine, Izmir, Turkey
[38] Department of Psychiatry, Massachusetts General Hospital, Boston, Massachusetts, USA
[39] Psychiatric and Neurodevelopmental Genetics Unit (PNGU), Massachusetts General Hospital, Boston, Massachusetts, USA
[40] Stanley Center for Psychiatric Research, Broad Institute, Cambridge, Massachusetts, USA
[41] Neuroscience and Mental Health, Cardiff University, Cardiff, UK
[42] Bioinformatics, University of British Columbia, Vancouver, BC, Canada
[43] Department of Epidemiology, Harvard T.H. Chan School of Public Health, Boston, Massachusetts, USA
[44] Department of Mathematics, Massachusetts Institute of Technology, Cambridge, Massachusetts, USA
[45] Department of Genetic Epidemiology in Psychiatry, Central Institute of Mental Health, Medical Faculty Mannheim, Heidelberg University, Mannheim, Baden-Württemberg, Germany
[46] Department of Psychiatry (UPK), University of Basel, Basel, Switzerland
[47] Department of Biomedicine, University of Basel, Basel, Switzerland
[48] Centre for Human Genetics, University of Marburg, Marburg, Germany
[49] Department of Psychiatry, Trinity College Dublin, Dublin, Ireland
[50] Psychiatry & Behavioral Sciences, Johns Hopkins University, Baltimore, Maryland, USA
[51] Bioinformatics Research Centre, Aarhus University, Aarhus, Denmark
[52] Institute of Genetic Medicine, Newcastle University, Newcastle upon Tyne, UK
[53] Danish Headache Centre, Department of Neurology, Rigshospitalet, Glostrup, Denmark
[54] Institute of Biological Psychiatry, Mental Health Center Sct. Hans, Mental Health Services Capital Region of Denmark, Copenhagen, Denmark
[55] iPSYCH, The Lundbeck Foundation Initiative for Psychiatric Research, Copenhagen, Denmark
[56] Brain and Mind Centre, University of Sydney, Sydney, New South Wales, Australia
[57] Interfaculty Institute for Genetics and Functional Genomics, Department of Functional Genomics, University Medicine and Ernst Moritz Arndt University Greifswald, Greifswald, Mecklenburg-Vorpommern, Germany
[58] Roche Pharmaceutical Research and Early Development, Pharmaceutical Sciences, Roche Innovation Center Basel, F. Hoffmann-La Roche Ltd, Basel, Switzerland
[59] Max Planck Institute of Psychiatry, Munich, Germany
[60] MRC Centre for Neuropsychiatric Genetics and Genomics, Cardiff University, Cardiff, UK
[61] Department of Psychological Medicine, University of Worcester, Worcester, UK
[62] Division of Research, Kaiser Permanente Northern California, Oakland, California, USA
[63] Psychiatry & The Behavioral Sciences, University of Southern California, Los Angeles, California, USA
[64] Department of Biomedical Informatics, Harvard Medical School, Boston, Massachusetts, USA
[65] Department of Medicine, Brigham and Women's Hospital, Boston, Massachusetts, USA
[66] Informatics Program, Boston Children's Hospital, Boston, Massachusetts, USA
[67] Wellcome Trust Centre for Human Genetics, University of Oxford, Oxford, UK

[68] Institute of Social and Preventive Medicine (IUMSP), Lausanne University Hospital and University of Lausanne, Lausanne, Switzerland
[69] Swiss Institute of Bioinformatics, Lausanne, Switzerland
[70] Division of Psychiatry, Centre for Clinical Brain Sciences, University of Edinburgh, Edinburgh, UK
[71] Mental Health, NHS 24, Glasgow, UK
[72] Department of Psychiatry and Psychotherapy, University of Bonn, Bonn, Germany
[73] Statistics, University of Oxford, Oxford, UK
[74] Psychiatry, Columbia University College of Physicians and Surgeons, New York, New York, USA
[75] School of Psychology and Counseling, Queensland University of Technology, Brisbane, Queensland, Australia
[76] Child and Youth Mental Health Service, Children's Health Queensland Hospital and Health Service, South Brisbane, Queensland, Australia
[77] Child Health Research Centre, University of Queensland, Brisbane, Queensland, Australia
[78] Estonian Genome Center, University of Tartu, Tartu, Estonia
[79] Medical Genetics, University of British Columbia, Vancouver, British Columbia, Canada
[80] Statistics, University of British Columbia, Vancouver, British Columbia, Canada
[81] DZHK (German Centre for Cardiovascular Research), Partner Site Greifswald, University Medicine, University Medicine Greifswald, Greifswald, Mecklenburg-Vorpommern, Germany
[82] Institute of Clinical Chemistry and Laboratory Medicine, University Medicine Greifswald, Greifswald, Mecklenburg-Vorpommern, Germany
[83] Institute of Health and Biomedical Innovation, Queensland University of Technology, Brisbane, Queensland, Australia
[84] Humus, Reykjavik, Iceland
[85] Virginia Institute for Psychiatric & Behavioral Genetics, Virginia Commonwealth University, Richmond, VA, USA
[86] Clinical Genetics, Vrije Universiteit Medical Center, Amsterdam, Netherlands
[87] Complex Trait Genetics, Vrije Universiteit Amsterdam, Amsterdam, Netherlands
[88] Solid Biosciences, Boston, Massachusetts, USA
[89] Department of Psychiatry, Washington University in Saint Louis School of Medicine, Saint Louis, Missouri, USA
[90] Department of Biochemistry and Molecular Biology II, Institute of Neurosciences, Biomedical Research Center (CIBM), University of Granada, Granada, Spain
[91] Department of Psychiatry, University of Groningen, University Medical Center Groningen, Groningen, Netherlands
[92] Department of Psychiatry and Psychotherapy, University Hospital, Ludwig Maximilian University Munich, Munich, Germany
[93] Institute of Psychiatric Phenomics and Genomics (IPPG), University Hospital, Ludwig Maximilian University Munich, Munich, Germany
[94] Division of Cancer Epidemiology and Genetics, National Cancer Institute, Bethesda, Maryland, USA
[95] Behavioral Health Services, Kaiser Permanente Washington, Seattle, Washington, USA
[96] Faculty of Medicine, Department of Psychiatry, University of Iceland, Reykjavik, Iceland
[97] School of Medicine and Dentistry, James Cook University, Townsville, Queensland, Australia
[98] Institute of Health and Wellbeing, University of Glasgow, Glasgow, UK

[99] deCODE Genetics / Amgen, Reykjavik, Iceland
[100] College of Biomedical and Life Sciences, Cardiff University, Cardiff, UK
[101] Institute of Epidemiology and Social Medicine, University of Münster, Münster, Nordrhein-Westfalen, Germany
[102] Institute for Community Medicine, University Medicine Greifswald, Greifswald, Mecklenburg-Vorpommern, Germany
[103] Department of Psychiatry, University of California, San Diego, San Diego, California, USA
[104] KG Jebsen Centre for Psychosis Research, Norway Division of Mental Health and Addiction, Oslo University Hospital, Oslo, Norway
[105] Medical Genetics Section, CGEM, IGMM, University of Edinburgh, Edinburgh, UK
[106] Clinical Neurosciences, University of Cambridge, Cambridge, UK
[107] Internal Medicine, Erasmus Medical Center, Rotterdam, Zuid-Holland, Netherlands
[108] Roche Pharmaceutical Research and Early Development, Neuroscience, Ophthalmology and Rare Diseases Discovery & Translational Medicine Area, Roche Innovation Center Basel, F. Hoffmann-La Roche Ltd, Basel, Switzerland
[109] Department of Psychiatry and Psychotherapy, University Medicine Greifswald, Greifswald, Mecklenburg-Vorpommern, Germany
[110] Department of Psychiatry, Leiden University Medical Center, Leiden, Netherlands
[111] Virginia Institute for Psychiatric & Behavioral Genetics, Virginia Commonwealth University, Richmond, Virginia, USA
[112] Computational Sciences Center of Emphasis, Pfizer Global Research and Development, Cambridge, Massachusetts, USA
[113] Institute for Molecular Bioscience; Queensland Brain Institute, The University of Queensland, Brisbane, Queensland, Australia
[114] Department of Psychiatry, University of Münster, Münster, Nordrhein-Westfalen, Germany
[115] Department of Psychiatry, Melbourne Medical School, University of Melbourne, Melbourne, Australia
[116] Florey Institute for Neuroscience and Mental Health, University of Melbourne, Melbourne, Australia
[117] Institute of Medical Genetics and Pathology, University Hospital Basel, University of Basel, Basel, Switzerland
[118] Institute of Neuroscience and Medicine (INM-1), Research Center Juelich, Juelich, Germany
[119] Amsterdam Public Health Institute, Vrije Universiteit Medical Center, Amsterdam, Netherlands
[120] Centre for Integrative Biology, Università degli Studi di Trento, Trento, Trentino-Alto Adige, Italy
[121] Department of Psychiatry and Psychotherapy, Medical Center - University of Freiburg, Faculty of Medicine, University of Freiburg, Freiburg, Germany
[122] Center for NeuroModulation, Faculty of Medicine, University of Freiburg, Freiburg, Germany
[123] Psychiatry, Kaiser Permanente Northern California, San Francisco, California, USA
[124] Medical Research Council Human Genetics Unit, Institute of Genetics and Molecular Medicine, University of Edinburgh, Edinburgh, UK
[125] Department of Psychiatry, University of Toronto, Toronto, Ontario, Canada
[126] Centre for Addiction and Mental Health, Toronto, Ontario, Canada
[127] Division of Psychiatry, University College London, London, UK

[128] Neuroscience Therapeutic Area, Janssen Research and Development, LLC, Titusville, New Jersey, USA
[129] Institute of Molecular and Cell Biology, University of Tartu, Tartu, Estonia
[130] Psychosis Research Unit, Aarhus University Hospital, Risskov, Aarhus, Denmark
[131] Munich Cluster for Systems Neurology (SyNergy), Munich, Germany
[132] University of Liverpool, Liverpool, UK
[133] Mental Health Center Copenhagen, Copenhagen Universtity Hospital, Copenhagen, Denmark
[134] Human Genetics and Computational Biomedicine, Pfizer Global Research and Development, Groton, Connecticut, USA
[135] Psychiatry, Harvard Medical School, Boston, Massachusetts, USA
[136] Psychiatry, University of Iowa, Iowa City, Iowa, USA
[137] Department of Psychiatry and Behavioral Sciences, Johns Hopkins University, Baltimore, Maryland, USA
[138] Department of Psychiatry and Psychotherapy, University Medical Center Göttingen, Goettingen, Niedersachsen, Germany
[139] Human Genetics Branch, NIMH Division of Intramural Research Programs, Bethesda, Maryland, USA
[140] Faculty of Medicine, University of Iceland, Reykjavik, Iceland
[141] Child and Adolescent Psychiatry, Erasmus Medical Center, Rotterdam, Zuid-Holland, Netherlands
[142] Psychiatry, Erasmus Medical Center, Rotterdam, Zuid-Holland, Netherlands
[143] Psychiatry, Dalhousie University, Halifax, NS, Canada
[144] Division of Translational Epidemiology, New York State Psychiatric Institute, New York, New York, USA
[145] Department of Clinical Medicine, University of Copenhagen, Copenhagen, Denmark
[146] Department of Medical & Molecular Genetics, King's College London, London, UK
[147] Psychiatry & Behavioral Sciences, Stanford University, Stanford, California, USA
[148] NIHR Maudsley Biomedical Research Centre, King's College London, London, UK
[149] Department of Genetics, University of North Carolina at Chapel Hill, Chapel Hill, North Carolina, USA
[150] Department of Psychiatry, University of North Carolina at Chapel Hill, Chapel Hill, North Carolina, USA

**Sex differences cross-disorder analysis group of the Psychiatric Genomics Consortium**

Martin Alda[1,2], Gabriëlla A. M. Blokland[3,4,5,6], Anders D. Børglum[7,8,9], Marco Bortolato[10], Janita Bratlen[11], Gerome Breen[12,13], Cynthia M. Bulik[14,15,16], Christie L. Burton[17], Enda M. Byrne[18], Caitlin E. Carey[6,19], Jonathan R. I. Coleman[12,13], Lea K. Davis[20], Ditte Demontis[21,22], Laramie E. Duncan[23], Howard J. Edenberg[24], Lauren Erdman[25], Steven V. Faraone[26], Jill M. Goldstein[27,28,29,30], Slavina B. Goleva[31], Jakob Grove[7,8,9,32], Wei Guo[33], Christopher Hübel[12,15], Laura M. Huckins[34], Ekaterina A. Khramtsova[35,36], Phil H. Lee[37], Joanna Martin[38], Carol A. Mathews[39], Manuel Mattheisen[7,22,40,41,42], Benjamin M. Neale[43], Roseann E. Peterson[44], Tracey L. Petryshen[4,5,6,45], Elise Robinson[6,43,46], Jordan W. Smoller[4,5,6], Eli A. Stahl[47,48], Barbara E. Stranger[35,49], Michela Traglia[50,51,52], Raymond K. Walters[43,53], Lauren A. Weiss[50,51,52], Thomas Werge[54,55,56,57], Stacey J. Winham[58], Naomi Wray[59], Yin Yao[60]

**Affiliations**

[1]Department of Psychiatry, Dalhousie University, Halifax, Nova Scotia, Canada
[2]National Institute of Mental Health, Klecany, Czech Republic
[3]Department of Psychiatry and Neuropsychology, School for Mental Health and Neuroscience, Faculty of Health, Medicine and Life Sciences, Maastricht University, Maastricht, The Netherlands
[4]Psychiatric and Neurodevelopmental Genetics Unit, Department of Psychiatry and Center for Genomic Medicine, Massachusetts General Hospital, Boston, Massachusetts, USA
[5]Department of Psychiatry, Harvard Medical School, Boston, Massachusetts, USA
[6]Stanley Center for Psychiatric Research, Broad Institute of MIT and Harvard, Cambridge, Massachusetts, USA
[7]Department of Biomedicine, Aarhus University, Aarhus, Denmark
[8]The Lundbeck Foundation Initiative for Integrative Psychiatric Research (iPSYCH), Copenhagen, Denmark
[9]Center for Genome Analysis and Personalized Medicine, Aarhus, Denmark
[10]University of Utah College of Pharmacy, Salt Lake City, Utah, USA
[11]Department of Human Genetics, Radboud university medical center, Nijmegen, the Netherlands
[12]Social, Genetic & Developmental Psychiatry Centre, Institute of Psychiatry, Psychology & Neuroscience, King's College London, London, UK
[13]National Institute of Health Research Maudsley Biomedical Research Centre, South London and Maudsley NHS Trust, London, UK
[14]Department of Psychiatry, University of North Carolina at Chapel Hill, Chapel Hill, North Carolina, USA
[15]Department of Medical Epidemiology and Biostatistics, Karolinska Institutet, Stockholm, Sweden
[16]Department of Nutrition, University of North Carolina at Chapel Hill, Chapel Hill, North Carolina, USA
[17]Hospital for Sick Children, Toronto, Ontario, Canada
[18]Institute for Molecular Bioscience, The University of Queensland, Brisbane, Queensland, Australia
[19]Analytic and Translational Genetics Unit, Center for Genomic Medicine, Massachusetts General Hospital, Boston, Massachusetts, USA
[20]Department of Medicine; Department of Psychiatry and Behavioral Sciences, Division of Genetic Medicine, Vanderbilt University Medical Center, Nashville, Tennessee, USA
[21]The Lundbeck Foundation Initiative for Integrative Psychiatric Research, iPSYCH, Denmark
[22]Department of Biomedicine; Center for Genomics and Personalized Medicine, Aarhus University, Aarhus, Denmark
[23]Department of Psychiatry and Behavioral Sciences, Stanford University, Stanford, California, USA
[24]Department of Biochemistry and Molecular Biology, Indiana University School of Medicine, Indianapolis, Indiana, USA
[25]University of Toronto/Hospital for Sick Children Genetics and Genome Biology Program, Toronto, Ontario, Canada
[26]Departments of Psychiatry and of Neuroscience and Physiology, SUNY Upstate Medical University, Syracuse, New York, USA
[27]Department of Psychiatry and Vincent Department of Obstetrics, Gynecology & Reproductive

Biology, Massachusetts General Hospital, Boston, Massachusetts, USA
[28]MGH-MIT-HMS Athinoula A. Martinos Center for Biomedical Imaging, Charlestown, Massachusetts, USA
[29]Innovation Center on Sex Differences in Medicine (ICON), Massachusetts General Hospital, Boston, Massachusetts, USA
[30]Departments of Psychiatry and Medicine, Harvard Medical School, Boston, Massachusetts, USA
[31]Department of Molecular Physiology and Biophysics, Vanderbilt University, Nashville, Tennessee, USA
[32]Bioinformatics Research Centre (BiRC), Aarhus, Denmark
[33]Genetic Epidemiology Research Branch National Institute of Mental Health, National Institutes of Health, Bethesda, MD, National Institutes of Health, Bethesda, Maryland, USA
[34]Pamela Sklar Division of Psychiatric Genomics, Icahn School of Medicine at Mount Sinai, New York, New York, USA
[35]Section of Genetic Medicine, Department of Medicine and Institute for Genomics and Systems Biology, University of Chicago, Chicago, Illinois, USA
[36]Computational Sciences, Janssen Pharmaceuticals, Spring House, Pennsylvania, USA
[37]Department of Psychiatry, Massachusetts General Hospital, Harvard Medical School, Boston, Massachusetts, USA
[38]MRC Centre for Neuropsychiatric Genetics and Genomics, Division of Psychological Medicine and Clinical Neurosciences, Cardiff University, Cardiff, UK
[39]Department of Psychiatry and Genetics Institute, University of Florida, Gainesville, Florida, USA
[40]Julius-Maximilians-Universität Würzburg, Wurzburg, Bayern, Germany
[41]Aarhus University, Aarhus, Denmark
[42]Centre for Psychiatry Research, Department of Clinical Neuroscience, Karolinska Institutet, Stockholm, Sweden
[43]Analytic and Translational Genetics Unit, Massachusetts General Hospital, Harvard Medical School, Boston, Massachusetts, USA
[44]Department of Psychiatry, Virginia Institute for Psychiatric and Behavioral Genetics, Virginia Commonwealth University, Richmond, Virginia, USA
[45]Concert Pharmaceuticals, Inc., Lexington, Massachusetts, USA
[46]Department of Epidemiology, Harvard T.H. Chan School of Public Health, Boston, Massachusetts, USA
[47]Department of Genetics and Genomic Sciences, Icahn School of Medicine at Mount Sinai, New York, New York, USA
[48]Department of Psychiatry, Icahn School of Medicine at Mount Sinai, New York, New York, USA
[49]Center for Genetic Medicine, Department of Pharmacology, Northwestern University, Chicago, Illinois, USA
[50]Department of Psychiatry, University of California San Francisco, San Francisco, California, USA
[51]Institute for Human Genetics, University of California San Francisco, San Francisco, California, USA
[52]Weill Institute for Neurosciences, University of California San Francisco, San Francisco, California, USA

[53]Broad Institute of MIT and Harvard, Stanley Center for Psychiatric Research, Cambridge, Massachusetts, USA
[54]Institute of Biological Psychiatry, MHC Sct. Hans, Mental Health Services Copenhagen, Copenhagen, Denmark
[55]Department of Clinical Medicine, Faculty of Health, University of Copenhagen, Copenhagen, Denmark
[56]Center for GeoGenetics, GLOBE Institute, University of Copenhagen, Copenhagen, Denmark
[57]The Lundbeck Foundation Initiative for Integrative Psychiatric Research, iPSYCH, Copenhagen, Denmark
[58]Department of Health Sciences Research, Division of Biomedical Statistics and Informatics, Mayo Clinic, Rochester, Minnesota, USA
[59]Institute for Molecular Bioscience, University of Queensland, Brisbane, Australia
[60]Fudan University, School of Life Sciences, Shanghai, China

**iPSYCH**

*Management Group:* Anders D. Børglum[1,2,3], David M. Hougaard[1,4], Merete Nordentoft[1,5,6], Ole Mors[1,7], Preben Bo Mortensen[1,3,8,9], Thomas Werge[1,6,10], Kristjar Skajaa[1,6]; *Advisory Board:* Markus Nöthen[11], Michael Owen[12], Robert H. Yolken[13], Niels Plath[14], Jonathan Mill[15], Daniel Geschwind[16]

**Affiliations:**

[1]The Lundbeck Foundation Initiative for Integrative Psychiatric Research (iPSYCH), Copenhagen, Denmark
[2]Department of Biomedicine - Human Genetics, Aarhus University, Aarhus, Denmark
[3]Centre for Integrative Sequencing (iSEQ), Aarhus University, Aarhus, Denmark
[4]Center for Neonatal Screening, Department for Congenital Disorders, Statens Serum Institut, Copenhagen, Denmark
[5]Copenhagen Mental Health Center, Mental Health Services Capital Region of Denmark Copenhagen, Copenhagen, Denmark
[6]Department of Clinical Medicine, Faculty of Health and Medical Sciences, University of Copenhagen, Copenhagen, Denmark
[7]Psychosis Research Unit, Aarhus University Hospital, Risskov, Denmark
[8]National Centre for Register-Based Research (NCCR), Aarhus University, Aarhus, Denmark
[9]Centre for Integrated Register-based Research (CIRRAU), Aarhus University, Aarhus, Denmark
[10]Institute of Biological Psychiatry, Mental Health Center Sct. Hans, Mental Health Services Copenhagen, Roskilde, Denmark
[11]Institute for Human Genetics, University of Bonn, Germany
[12]MRC Centre for Neuropsychiatric Genetics & Genomics, Cardiff University, Cardiff, UK
[13]Stanley Division of Development Neurovirology, Johns Hopkins University, Baltimore, Maryland, USA
[14]H. Lundbeck A/S, Copenhagen, Denmark
[15]University of Exeter Medical School & Institute of Psychiatry, King's College London, London, UK
[16]David Geffen School of Medicine, University of California Los Angeles, Los Angeles, California, USA

### Supplementary References

1. Chang CC, Chow CC, Tellier LC, Vattikuti S, Purcell SM, Lee JJ (2015): Second-generation PLINK: Rising to the challenge of larger and richer datasets. *Gigascience*. 4:7.
2. Khramtsova EA, Heldman R, Derks EM, Yu D, Tourette Syndrome/Obsessive-Compulsive Disorder Working Group of the Psychiatric Genomics C, Davis LK, et al. (2018): Sex differences in the genetic architecture of obsessive-compulsive disorder. *Am J Med Genet B Neuropsychiatr Genet*. 180:351-364.
3. Pedersen CB, Mors O, Bertelsen A, Waltoft BL, Agerbo E, McGrath JJ, et al. (2014): A comprehensive nationwide study of the incidence rate and lifetime risk for treated mental disorders. *JAMA Psychiatry*. 71:573-581.
4. Bulik-Sullivan BK, Loh PR, Finucane HK, Ripke S, Yang J, Schizophrenia Working Group of the Psychiatric Genomics C, et al. (2015): LD Score regression distinguishes confounding from polygenicity in genome-wide association studies. *Nat Genet*. 47:291-295.
5. Bulik-Sullivan BK, Finucane HK, Anttila V, Gusev A, Day FR, Loh PR, et al. (2015): An atlas of genetic correlations across human diseases and traits. *Nat Genet*. 47:1236-1241.
6. Martin J, Khramtsova EA, Goleva SB, Blokland GAM, Traglia M, Walters RK, et al. (2020): Examining sex-differentiated genetic effects across neuropsychiatric and behavioral traits. *bioRxiv*. https://doi.org/10.1101/2020.05.04.076042.
7. Willer CJ, Li Y, Abecasis GR (2010): METAL: Fast and efficient meta-analysis of genomewide association scans. *Bioinformatics*. 26:2190-2191.
8. Keller MC (2014): Gene x environment interaction studies have not properly controlled for potential confounders: The problem and the (simple) solution. *Biol Psychiatry*. 75:18-24.
9. Loley C, Alver M, Assimes TL, Bjonnes A, Goel A, Gustafsson S, et al. (2016): No Association of Coronary Artery Disease with X-Chromosomal Variants in Comprehensive International Meta-Analysis. *Sci Rep*. 6:35278.
10. Chow JC, Yen Z, Ziesche SM, Brown CJ (2005): Silencing of the mammalian X chromosome. *Annu Rev Genomics Hum Genet*. 6:69-92.
11. König IR, Loley C, Erdmann J, Ziegler A (2014): How to include chromosome X in your genome-wide association study. *Genet Epidemiol*. 38:97-103.
12. Loley C, Ziegler A, Konig IR (2011): Association tests for X-chromosomal markers--a comparison of different test statistics. *Hum Hered*. 71:23-36.
13. Cross-Disorder Group of the Psychiatric Genomics Consortium, Lee SH, Ripke S, Neale BM, Faraone SV, Purcell SM, et al. (2013): Genetic relationship between five psychiatric disorders estimated from genome-wide SNPs. *Nat Genet*. 45:984-994.
14. Cross-Disorder Group of the Psychiatric Genomics Consortium, Smoller JW, Craddock N, Kendler K, Lee PH, Neale BM, et al. (2013): Identification of risk loci with shared effects on five major psychiatric disorders: a genome-wide analysis. *Lancet*. 381:1371-1379.
15. Lee PH, Bergen SE, Perlis RH, Sullivan PF, Sklar P, Smoller JW, et al. (2011): Modifiers and subtype-specific analyses in whole-genome association studies: a likelihood framework. *Hum Hered*. 72:10-20.
16. Bhattacharjee S, Rajaraman P, Jacobs KB, Wheeler WA, Melin BS, Hartge P, et al. (2012): A subset-based approach improves power and interpretation for the combined analysis of genetic association studies of heterogeneous traits. *Am J Hum Genet*. 90:821-835.
17. Benner C, Spencer CC, Havulinna AS, Salomaa V, Ripatti S, Pirinen M (2016): FINEMAP: Efficient variable selection using summary data from genome-wide association studies. *Bioinformatics*. 32:1493-1501.

18. Hormozdiari F, Kostem E, Kang EY, Pasaniuc B, Eskin E (2014): Identifying causal variants at loci with multiple signals of association. *Genetics*. 198:497-508.
19. de Leeuw CA, Mooij JM, Heskes T, Posthuma D (2015): MAGMA: Generalized gene-set analysis of GWAS data. *PLoS computational biology*. 11:e1004219.
20. Gene Ontology Consortium (2015): Gene Ontology Consortium: going forward. *Nucleic Acids Res*. 43:D1049-1056.
21. Gene Ontology Consortium, Blake JA, Dolan M, Drabkin H, Hill DP, Li N, et al. (2013): Gene Ontology annotations and resources. *Nucleic Acids Res*. 41:D530-535.
22. Liberzon A, Birger C, Thorvaldsdottir H, Ghandi M, Mesirov JP, Tamayo P (2015): The Molecular Signatures Database (MSigDB) hallmark gene set collection. *Cell Syst*. 1:417-425.
23. Network & Pathway Analysis Subgroup of Psychiatric Genomics Consortium (2015): Psychiatric genome-wide association study analyses implicate neuronal, immune and histone pathways. *Nat Neurosci*. 18:199-209.
24. Pardiñas AF, Holmans P, Pocklington AJ, Escott-Price V, Ripke S, Carrera N, et al. (2018): Common schizophrenia alleles are enriched in mutation-intolerant genes and in regions under strong background selection. *Nat Genet*.
25. Watanabe K, Taskesen E, van Bochoven A, Posthuma D (2017): Functional mapping and annotation of genetic associations with FUMA. *Nat Commun*. 8:1826.
26. Benjamini Y, Hochberg Y (1995): Controlling the false discovery rate: a practical and powerful approach to multiple testing. *Journal of the Royal Statistical Society, Series B*. 57:289–300.
27. GTEx Consortium (2013): The Genotype-Tissue Expression (GTEx) project. *Nat Genet*. 45:580-585.
28. GTEx Consortium (2015): Human genomics. The Genotype-Tissue Expression (GTEx) pilot analysis: multitissue gene regulation in humans. *Science*. 348:648-660.
29. Kang HJ, Kawasawa YI, Cheng F, Zhu Y, Xu X, Li M, et al. (2011): Spatio-temporal transcriptome of the human brain. *Nature*. 478:483-489.
30. Pletikos M, Sousa AM, Sedmak G, Meyer KA, Zhu Y, Cheng F, et al. (2014): Temporal specification and bilaterality of human neocortical topographic gene expression. *Neuron*. 81:321-332.
31. Sunkin SM, Ng L, Lau C, Dolbeare T, Gilbert TL, Thompson CL, et al. (2013): Allen Brain Atlas: An integrated spatio-temporal portal for exploring the central nervous system. *Nucleic Acids Res*. 41:D996-D1008.
32. Cahoy JD, Emery B, Kaushal A, Foo LC, Zamanian JL, Christopherson KS, et al. (2008): A transcriptome database for astrocytes, neurons, and oligodendrocytes: a new resource for understanding brain development and function. *The Journal of neuroscience : the official journal of the Society for Neuroscience*. 28:264-278.
33. Zhang Y, Chen K, Sloan SA, Bennett ML, Scholze AR, O'Keeffe S, et al. (2014): An RNA-sequencing transcriptome and splicing database of glia, neurons, and vascular cells of the cerebral cortex. *The Journal of neuroscience : the official journal of the Society for Neuroscience*. 34:11929-11947.
34. Gandal MJ, Zhang P, Hadjimichael E, Walker RL, Chen C, Liu S, et al. (2018): Transcriptome-wide isoform-level dysregulation in ASD, schizophrenia, and bipolar disorder. *Science*. 362.
35. Wang D, Liu S, Warrell J, Won H, Shi X, Navarro FCP, et al. (2018): Comprehensive functional genomic resource and integrative model for the human brain. *Science*. 362.

36. Hoffman GE, Bendl J, Voloudakis G, Montgomery KS, Sloofman L, Wang YC, et al. (2019): CommonMind Consortium provides transcriptomic and epigenomic data for Schizophrenia and Bipolar Disorder. *Sci Data*. 6:180.
37. Jaffe AE, Straub RE, Shin JH, Tao R, Gao Y, Collado-Torres L, et al. (2018): Developmental and genetic regulation of the human cortex transcriptome illuminate schizophrenia pathogenesis. *Nat Neurosci*. 21:1117-1125.
38. Collado-Torres L, Burke EE, Peterson A, Shin J, Straub RE, Rajpurohit A, et al. (2019): Regional Heterogeneity in Gene Expression, Regulation, and Coherence in the Frontal Cortex and Hippocampus across Development and Schizophrenia. *Neuron*. 103:203-216 e208.
39. Nievergelt CM, Maihofer AX, Klengel T, Atkinson EG, Chen CY, Choi KW, et al. (2019): International meta-analysis of PTSD genome-wide association studies identifies sex- and ancestry-specific genetic risk loci. *Nat Commun*. 10:4558.
40. Martin J, Walters RK, Demontis D, Mattheisen M, Lee SH, Robinson E, et al. (2018): A genetic investigation of sex bias in the prevalence of attention-deficit/hyperactivity disorder. *Biol Psychiatry*. 83:1044-1053.
41. Mitra I, Tsang K, Ladd-Acosta C, Croen LA, Aldinger KA, Hendren RL, et al. (2016): Pleiotropic mechanisms indicated for sex differences in autism. *PLoS Genet*. 12:e1006425.
42. Hyde CL, Nagle MW, Tian C, Chen X, Paciga SA, Wendland JR, et al. (2016): Identification of 15 genetic loci associated with risk of major depression in individuals of European descent. *Nat Genet*. 48:1031-1036.
43. Walters R, Abbott L, Bryant S, Churchhouse C, Palmer D, Neale B (2018): Heritability of >2,000 traits and disorders in the UK Biobank. http://wwwnealelabis/uk-biobank/.
44. Cross-Disorder Group of the Psychiatric Genomics Consortium (2019): Genomic relationships, novel loci, and pleiotropic mechanisms across eight psychiatric disorders. *Cell*. 179:1469-1482 e1411.
45. Major Depressive Disorder Working Group of the Psychiatric GWAS Consortium, Ripke S, Wray NR, Lewis CM, Hamilton SP, Weissman MM, et al. (2013): A mega-analysis of genome-wide association studies for major depressive disorder. *Mol Psychiatry*. 18:497-511.
46. Psychiatric Genomics Consortium Schizophrenia Working Group (2014): Biological insights from 108 schizophrenia-associated genetic loci. *Nature*. 511:421-427.
47. Psychiatric GWAS Consortium Bipolar Disorder Working Group (2011): Large-scale genome-wide association analysis of bipolar disorder identifies a new susceptibility locus near ODZ4. *Nat Genet*. 43:977-983.
48. Pedersen CB, Bybjerg-Grauholm J, Pedersen MG, Grove J, Agerbo E, Baekvad-Hansen M, et al. (2018): The iPSYCH2012 case-cohort sample: New directions for unravelling genetic and environmental architectures of severe mental disorders. *Mol Psychiatry*. 23:6-14.
49. Pruim RJ, Welch RP, Sanna S, Teslovich TM, Chines PS, Gliedt TP, et al. (2010): LocusZoom: regional visualization of genome-wide association scan results. *Bioinformatics*. 26:2336-2337.
50. Irish Schizophrenia Genomics C, the Wellcome Trust Case Control C (2012): Genome-wide association study implicates HLA-C*01:02 as a risk factor at the major histocompatibility complex locus in schizophrenia. *Biol Psychiatry*. 72:620-628.
51. Psychosis Endophenotypes International Consortium, Wellcome Trust Case-Control Consortium, Bramon E, Pirinen M, Strange A, Lin K, et al. (2014): A genome-wide association analysis of a broad psychosis phenotype identifies three loci for further investigation. *Biol Psychiatry*. 75:386-397.

52. Wellcome Trust Case Control C (2007): Genome-wide association study of 14,000 cases of seven common diseases and 3,000 shared controls. *Nature*. 447:661-678.
53. Levinson DF, Shi J, Wang K, Oh S, Riley B, Pulver AE, et al. (2012): Genome-wide association study of multiplex schizophrenia pedigrees. *Am J Psychiatry*. 169:963-973.
